# Single-molecule sequencing of long DNA molecules allows high contiguity *de novo* genome assembly for the fungus fly, *Sciara coprophila*

**DOI:** 10.1101/2020.02.24.963009

**Authors:** John M. Urban, Michael S. Foulk, Jacob E. Bliss, C. Michelle Coleman, Nanyan Lu, Reza Mazloom, Susan J. Brown, Allan C. Spradling, Susan A. Gerbi

**Affiliations:** Brown University Division of Biology and Medicine, Department of Molecular Biology, Cell Biology and Biochemistry, Providence, Rhode Island 02912, USA; Carnegie Institution for Science, Department of Embryology, 3520 San Martin Drive, Baltimore, Maryland 21218, USA; Mercyhurst University, Department of Biology, Erie, PA 16546, USA; Kansas State University Division of Biology, KSU Bioinformatics Center, Ackert Hall, Manhattan, Kansas 66502, USA

**Keywords:** genome assembly, single molecule sequencing, long reads, optical maps, nanopore sequencing, DNA modifications, non-model organism, emerging model system, insect genomes, fungus fly *Sciara* (*Bradysia*) coprophila

## Abstract

The lower Dipteran fungus fly, *Sciara coprophila*, has many unique biological features. For example, *Sciara* undergoes paternal chromosome elimination and maternal X chromosome nondisjunction during spermatogenesis, paternal X elimination during embryogenesis, intrachromosomal DNA amplification of DNA puff loci during larval development, and germline-limited chromosome elimination from all somatic cells. Paternal chromosome elimination in *Sciara* was the first observation of imprinting, though the mechanism remains a mystery. Here, we present the first draft genome sequence for *Sciara coprophila* to take a large step forward in aiding these studies. We approached assembling the *Sciara* genome using multiple sequencing technologies: PacBio, Oxford Nanopore MinION, and Illumina. To find an optimal assembly using these datasets, we generated 44 Illumina assemblies using 7 short-read assemblers and 50 long-read assemblies of PacBio and MinION sequence data using 6 long-read assemblers. We ranked assemblies using a battery of reference-free metrics, and scaffolded a subset of the highest-ranking assemblies using BioNano Genomics optical maps. RNA-seq datasets from multiple life stages and both sexes facilitated genome annotation. Moreover, we anchored nearly half of the *Sciara* genome sequence into chromosomes. Finally, we used the signal level of both the PacBio and Oxford Nanopore data to explore the presence or absence of DNA modifications in the *Sciara* genome since DNA modifications may play a role in imprinting in *Sciara*, as they do in mammals. These data serve as the foundation for future research by the growing community studying the unique features of this emerging model system.

## INTRODUCTION

The fungus gnat, *Sciara coprophila* (also known as *Bradysia coprophila*), is a Dipteran fly that is both an old and emerging model system rich with opportunities for studying fundamental biology, especially chromosomal biology due to its dynamic genome. In contrast to the rule that the amount of nuclear DNA is constant in all cells of an organism (Boivin et al. 1948), the nuclear DNA in *Sciara* cells exhibits copy number differences at the levels of loci, chromosomes, and the genome. Genomic copy numbers vary from canonical haploid and diploid tissues to the endocycling larval salivary glands that result in cells with over 8000 copies of each chromosome held closely together to form giant polytene chromosomes (Rasch 1970b). Locus-specific copy number regulation occurs at the “DNA puff” loci in polytene chromosomes where site-specific re-replication results in intrachromosomal DNA amplification (Rasch 1970a; Gerbi et al. 2002). Whole chromosome copy number gains and losses are seen in spermatogenesis, fertilization, in somatic cells of early embryos, and in the germ-line during development (Gerbi 1986).

The chromosome cycle of *Sciara* gives rise to numerous research opportunities not found in *Drosophila*, the standard Dipteran model organism. In *Sciara*, there are “L” chromosomes limited to the germ-line of both sexes (Gerbi 1986). Whereas oogenesis has orthodox chromosome movements, they are unusual in spermatogenesis leading to sperm cells that are haploid for each autosome, diploid for the X, and variable for the L with 0-4 copies. X diploidy in sperm is due to developmentally programmed X chromosome nondisjunction in male meiosis (Gerbi 1986). Fertilization ultimately produces zygotes and early embryos that are temporarily triploid for the X chromosome, and variable for the L. The fates of the X and L chromosomes in early embryonic nuclei are subsequently determined by whether a cell is somatic or germline, and by whether it is male or female. All L chromosomes are eliminated from somatic cell nuclei in early embryos. As part of the sex determination pathway, X diploidy is restored in female somatic cells (XX) by the elimination of one X, but the elimination of two X chromosomes in male somatic cells (XO) leads to X haploidy (Gerbi 1986). Diploidy for the X and L is restored in the germline through elimination events later in development (Gerbi 1986).

The X chromosomes eliminated during early embryo development are always paternally derived. Moreover, all paternally derived chromosomes, except L, are eliminated in the first meiotic division of spermatogenesis in the only known case of a naturally occurring monopolar spindle (Gerbi 1986). The ability to differentiate between the maternal and paternal chromosomes gave rise to the term “imprinting” (Crouse 1960) and was the first description of this phenomenon in any system. L chromosomes apparently escape imprinting in *Sciara* as maternal and paternal copies are both eliminated from all nuclei destined to become somatic cells (Crouse et al. 1971), and they are not eliminated with the paternal cohort during male meiosis. The mechanism for imprinting in *Sciara* remains unknown. It is of interest to learn if DNA modifications occur in the *Sciara* genome, since imprinting in mammals utilizes DNA methylation (Li et al. 1993).

This black fungus gnat and its unusual chromosomal features are part of one of the most interesting yet little-studied groups of Dipteran flies, the suborder Nematocera. The group of Nematocerans contains agricultural pests as well as disease vectors, such as mosquitoes (Matthews et al 2018). Nematocera diverged from higher Dipteran flies, the suborder Brachycera that includes the fruit fly *Drosophila melanogaster*, ∼200 million years ago (Wiegmann et al. 2011). *Bradysia (Sciara) coprophila* is classified as part of the infraorder Bibionomorpha in the Sciaroidea super family, which also comprises the family Cecidomyiidae (gall midges) and the Hessian fly in particular, a notorious wheat pest (Stuart et al 2012). Sciarid flies also include the Mycetophilidae, a fungus gnat family where members have been shown to withstand freezing and thawing (Sformo et al 2009). Indeed, we also have unpublished observations that *Sciara coprophila* embryos and larvae can be stored in the cold from a few months to over a year in a diapause-like state before returning to room temperature and resuming development. Despite flies making up at least 10% of all metazoan diversity, there are only 157 Dipteran genomes described (i5k.github.io/arthropod_genomes_at_ncbi), most of which are highly fragmented assemblies, and the majority of which are from the higher Dipteran order and limited to only two suborders therein (Muscomorpha, Stratiomyomorpha). Thus, there is a real need for high quality genomes across the Dipteran tree, and particularly for the lower Dipteran suborder that includes *Sciara*.

The complete *Sciara* genome comprises three autosomes (chromosomes II, III and IV), an X chromosome, and the germ-line limited L chromosome (Figure 1; Gerbi 1986). L chromosomes are eliminated from nuclei destined to become somatic cells in the 5th or 6th nuclear division, ∼3 hours after egg deposition (Gerbi 1986). *Sciara* lacks a Y chromosome, and sex is determined by whether or not the mother carries a variant of the X, called X’, that has a long paracentric inversion. Females that are XX have only sons, whereas X’X females have only daughters. The XX or X’X genotype of adult females can be determined by phenotypic wing markers (Figure 1). The *Sciara* genome has ∼38% GC content (Gerbi 1971) and is ∼280 Mb in somatic cells and ∼363 Mb in germ cells that contain L chromosomes (Rasch 2006) (Supplemental Table S1A-D).

**Figure 1:**
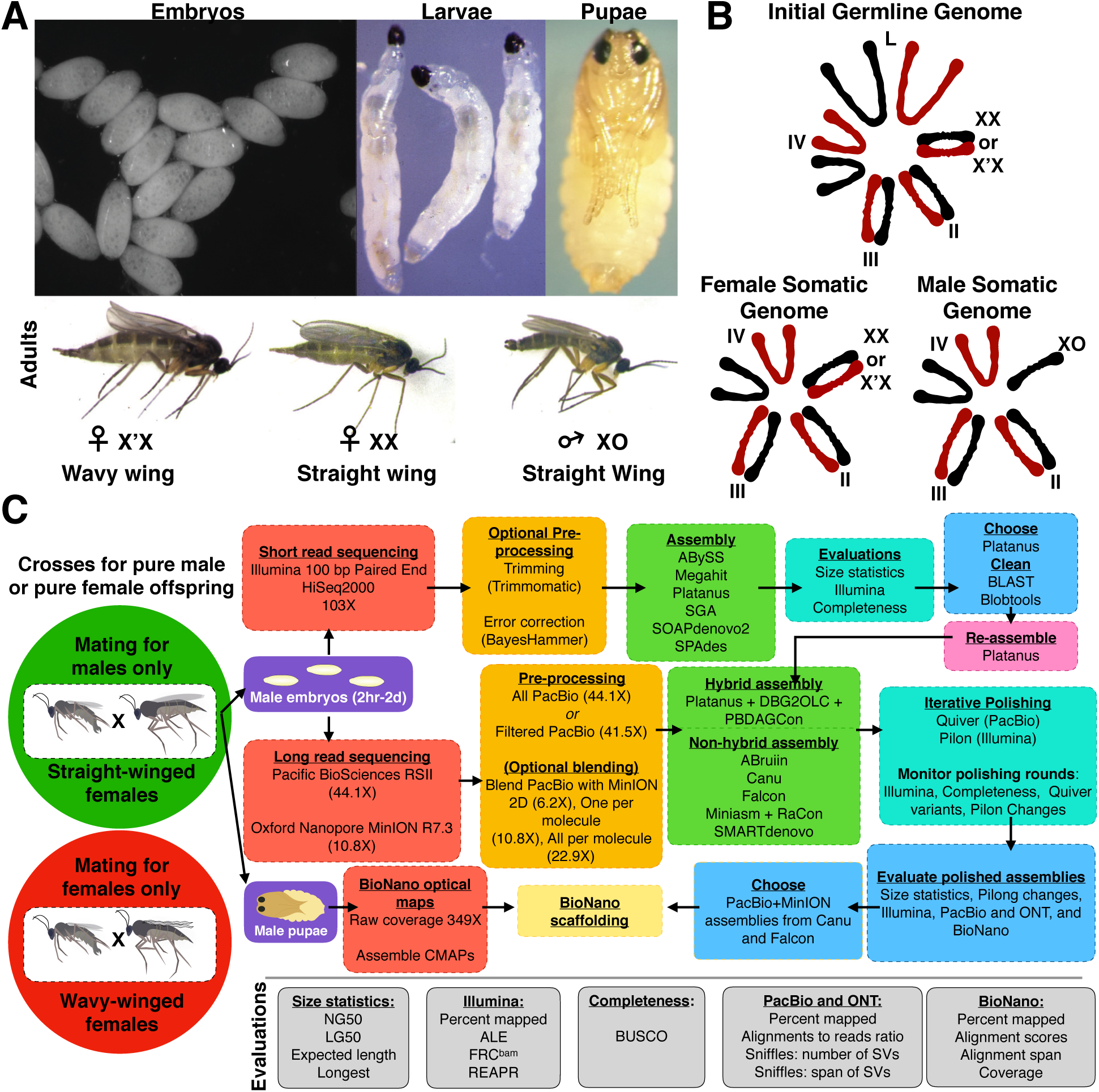
Genome sequencing and assembly strategy for *Sciara coprophila*. **(A)** Images of different lifecycle stages of *Sciara coprophila*: embryos, larvae, pupae, and adults. The adult figures show a male and the two different types of females that can be distinguished based on the Wavy wing phenotype that marks the X’ chromosome. **(B)** Examples of different chromosome compositions in *Sciara* cells. We focused on the male somatic genome. Red chromosomes are paternal, black are maternal. **(C)** The genome assembly and evaluation workflow up until BioNano scaffolding. The workflow begins by highlighting that crosses can be conducted to generate only male (green) or only female (red) offspring using the Wavy wing phenotypic marker. We used only matings for males to obtain genomic DNA for sequencing, illustrated by the arrows from the green circle that point to subsequent steps in the pipeline. Both male (green) and female (red) offspring were used for transcriptomes.

There are many ways to assemble a genome, but no universal recipe of sequencing technologies, pre-assembly practices (e.g. quality filtering, error correction), assembly algorithms, parameter tuning, and post-assembly steps exists that guarantees the best assembly for a given genome. Therefore, to maximize contiguity and quality, we sequenced the *Sciara* genome with multiple technologies, including 100 bp Illumina paired-end reads, long reads from Pacific Biosciences (PacBio) (Eid et al. 2009) and the Oxford Nanopore MinION (Ip et al. 2015), and generated optical maps from the BioNano Genomics Irys platform (Lam et al. 2012). We produced assemblies using combinations of these technologies with multiple algorithms and ranked each using a battery of reference-free metrics. Highly contiguous assemblies that were most complete in expected gene content and which were judged to be most consistent with our Illumina, PacBio, MinION, BioNano, and RNA-seq datasets were identified. These evaluations allowed us to monitor steps (e.g. polishing), to choose a few assemblies for BioNano scaffolding, and to make a final selection for the *Sciara* draft genome.

We report here the first draft genome assembly for *Sciara coprophila*, and its accompanying gene and repeat annotations. The *Sciara* genome sequence will be a valuable resource for future comparative genomics analyses, as one of the highest-quality Nematoceran genome sequences available, as the only sequenced member of the Sciaridae family, and due to its phylogenetic position at the gateway between lower and higher Dipterans. More than half of the *Sciara* genome is contained on contigs ≥1.9 Mb and scaffolds ≥6.8 Mb. This exceeds the contiguity of ∼90% of all Dipteran genome assemblies (i5k.github.io/arthropod_genomes_at_ncbi). More specifically, the contig sizes in this release of the *Sciara* genome are longer than 42 of the 43 Nematoceran genome assemblies described, only outshined by the assembly for the mosquito, *Aedes aegypti* (Matthews et al 2018). The megabase-scale contigs and scaffolds will aid in efforts to improve the contiguity of more fragmented assemblies of related species by synteny. The genome annotation contains >97% of expected gene content. Up to 49% of the *Sciara* genome sequence was anchored into specific loci of chromosomes X, II, III, and IV; and 100% was classified as either X or autosomal, allowing an analysis of dosage compensation of the single male X. A *Rickettsia* genome was co-assembled with the *Sciara* genome, suggesting it may be an endosymbiont. The signal data from both PacBio and MinION both suggest the presence of DNA modifications in the *Sciara* genome. Finally, candidate L sequences were briefly explored. Sequencing, assembly, and annotation of the *Sciara* genome reported here serves as the foundation for future studies of the many unique features of this emerging model organism.

## RESULTS

### Data collection

Using wing phenotypic markers, XX *Sciara* adult females were crossed with XO males to produce only male progeny (Figure 1). DNA isolated from purely male embryos was used for sequencing (Illumina, PacBio, MinION), thereby avoiding assembly complications from the heteromorphic X’ chromosome found in female-producing females (Figure 1B), as well as minimizing possible complications from later life stages due to polytenization and contamination from the gut microbiome. Moreover, although early embryos were included to potentially capture sequences from chromosome L, the somatic genome is over-represented in these samples and we do not expect L sequences from the germline genome to be well-represented. Separate preparations of male embryo genomic DNA were made for 100 bp paired-end Illumina, and long-read PacBio and Oxford Nanopore MinION sequencing resulting in 103X, 50-55X, and 10-11X coverage, respectively (Table 1). We used male pupae to collect nearly 350X coverage from a third single molecule technology: optical maps from the BioNano Genomics (BNG) Irys (Lam et al. 2012) (Table 1). Finally, to facilitate gene annotation, we acquired sex- and stage-specific 100 bp paired-end RNA-seq datasets from whole embryos, larvae, pupae, and adults using the appropriate crosses for only males (XX × XO) or only females (X’X × XO) (Supplemental Table S2).

**Table 1:** Genome sequencing datasets for Sciara coprophila

|  | **Illumina**<br>**HiSeq 2000** | **PacBio RSII** | **Oxford**<br>**Nanopore**<br>**MinION MkI** | **BioNano**<br>**Genomics**<br>**Irys** |
| --- | --- | --- | --- | --- |
| **Source** | Male Embryos | Male Embryos | Male Embryos* | Male pupae |
| **Library** | Paired-End* | SMRTBell | MAP002-006<br>(2D) | IrysPrep |
| **Details** | - | P5-C3 | Pores R7.3-R7.3 70bps<br>6mer | BssSI |
| **Read Length N50**<br>**(kb)** | 0.1 | 9.681 | 9.934 | 132.613 |
| **Mean Read**<br>**Length (kb)** | 0.1 | 6.607 | 5.883 | 62.531 |
| **Count** | 301,513,554 | 1,949,427 | 532,714 | 1,628,681 |
| **Span (Gb)** | 30.15 | 12.88 | 3.15 | 101.84 |
| **Coverage >0 kb** | 103.26 | 44.11 | 10.77 | 348.78 |
| **>20 kb** | 0 | 1.28 | 2.91 | 330.22 |
| **>30 kb** | 0 | 0.01 | 1.72 | 323.31 |
| **>50 kb** | 0 | 0 | 0.71 | 303.02 |
| **>100 kb** | 0 | 0 | 0.28 | 226.1 |
| **>150 kb** | 0 | 0 | 0.2 | 148.5 |

\*A minority of the MinION data came from male adults (see Methods).

**Table 2:** Anchoring into chromosomes using previously known sequences

| Sequence | Location | Canu contig size | Falcon contig size | Reference |
| --- | --- | --- | --- | --- |
| DNA puff II/9A | Chr II locus 9A | 13.1 Mb | 28.5 Mb | DiBartolomeis and Gerbi 1989; Bienz-Tadmor et al. 1991; Urnov et al 2002; Foulk et al. 2006 |
| RNA Puff III/9B | Chr III locus 9B | 5.4 Mb | 12.5 Mb | Wu et al 1993; Foulk et al. 2006 |
| Ecdysone receptor | Chr IV locus 12A | 3.8 Mb | 9.6 Mb* | Foulk et al 2013 |
| Ultraspiracle | Chr IV locus 10A | 9.3 Mb | 5.5 Mb | Foulk et al 2013 |
| Hsp70 | Chr IV locus 4A or 12C | 5.4 Mb | 13 Mb | Mok et al. 2000 |
| Hsp70 | Chr IV locus 4A or 12C | 6.8 Mb | 2.6 Mb | Mok et al. 2000 |
| ScoHet1 | Chr IV locus 5A | 15.2 Mb | (9.6 Mb)* | Greciano et al. 2009 |
| ScoHet2 | Chr IV locus 12C-13A | 5.9 Mb | 4 Mb | Greciano et al. 2009 |
| rDNA | End of Chr X | 5 primary contigs and 11 associated contigs ($\Sigma$ 1.3 Mb) | 2 primary contigs and 41 associated contigs ($\Sigma$ 1.7 Mb) | Pardue et al. 1970; Gerbi 1971; Crouse et al. 1977; Kerrebrock et al. 1989 |
| Microclone B4 | End of Chr X | 69.8 kb | 59 kb | Escribá et al. 2011 |
| Microclone F7 | Near centromere of Chr X**, non-centromeric Chr IV, L chromosomes | 3 associated contigs ($\Sigma$ 66.8 kb) | 1 primary and 1 associated contig ($\Sigma$ 161.6 kb) | Escribá et al. 2011 |
| Microclone G2 (Sccr) | Centromeres of all chromosomes | 20 primary and 85 associated contigs ($\Sigma$ 1.3 Mb) | 6 primary and 42 associated contigs ($\Sigma$ 604 kb) | Escribá et al. 2011 |

\*Ecdysone receptor (EcR) and ScoHet1 identified the same 9.6 Mb contig in Falcon. The locus inconsistency may represent a misassembly in Falcon or misannotation from Greciano et al (2009). Nevertheless, both EcR and ScoHet1 results agree it is from chr IV.
\*\* Coverage analyses confirm contigs with F7 as chromosome X sequence.

### Short-read assemblies

Using the Illumina dataset, both with and without quality filtering and/or error-correction steps, we generated 44 assemblies using 7 popular short-read genome assemblers (Figures 1C and 2A). The assemblies ranged from ∼226-348 Mb in size (Supplemental Table S3), with a mean assembly size of ∼280 Mb, exactly the expected somatic genome size. Evaluating these assemblies with several reference-free evaluation tools (see Methods) allowed us to determine the highest quality assemblies (Figures 1C, 2A-D, Supplemental Figure S1). Rankings from these metrics were generally correlated with each other (Figure 2A, Supplemental Figure S2A). Platanus and ABySS assemblies most consistently returned the best rankings across metrics with Platanus assemblies having higher mean ranks overall (Figure 2A and Supplemental Figure S1). All Illumina assemblies did moderately well in terms of gene content, most containing between 80-85% of the expected Arthropod BUSCOs (Figure 2B). Nonetheless, all of the Illumina assemblies were highly fragmented, containing up to hundreds of thousands of contigs mostly less than 1 kb in length. The NG50 values ranged from 2.5-7.3 kb (Figure 2C, Supplemental Table S3). Although some scaffolds in the assemblies reached up to the Mb range, they were all bacterial, a common observation for assemblies from whole animals (Supplemental Figure S3). Of recognizable bacterial sequence, at the genus level, ∼90% was characterized as *Delftia* and ∼5% as *Rickettsia*. Amongst all Illumina assemblies, the longest scaffolds of apparent insect origin were 50-60 kb. Filtering for bacterial contamination and re-assembling with the filtered data did not improve the contiguity (Supplemental Table S3). Although short-read-only assemblies were not pursued further, the highest quality Platanus assembly was used in hybrid assemblies with long reads, and the high accuracy Illumina short reads were useful for polishing long-read assemblies.

**Figure 2:**
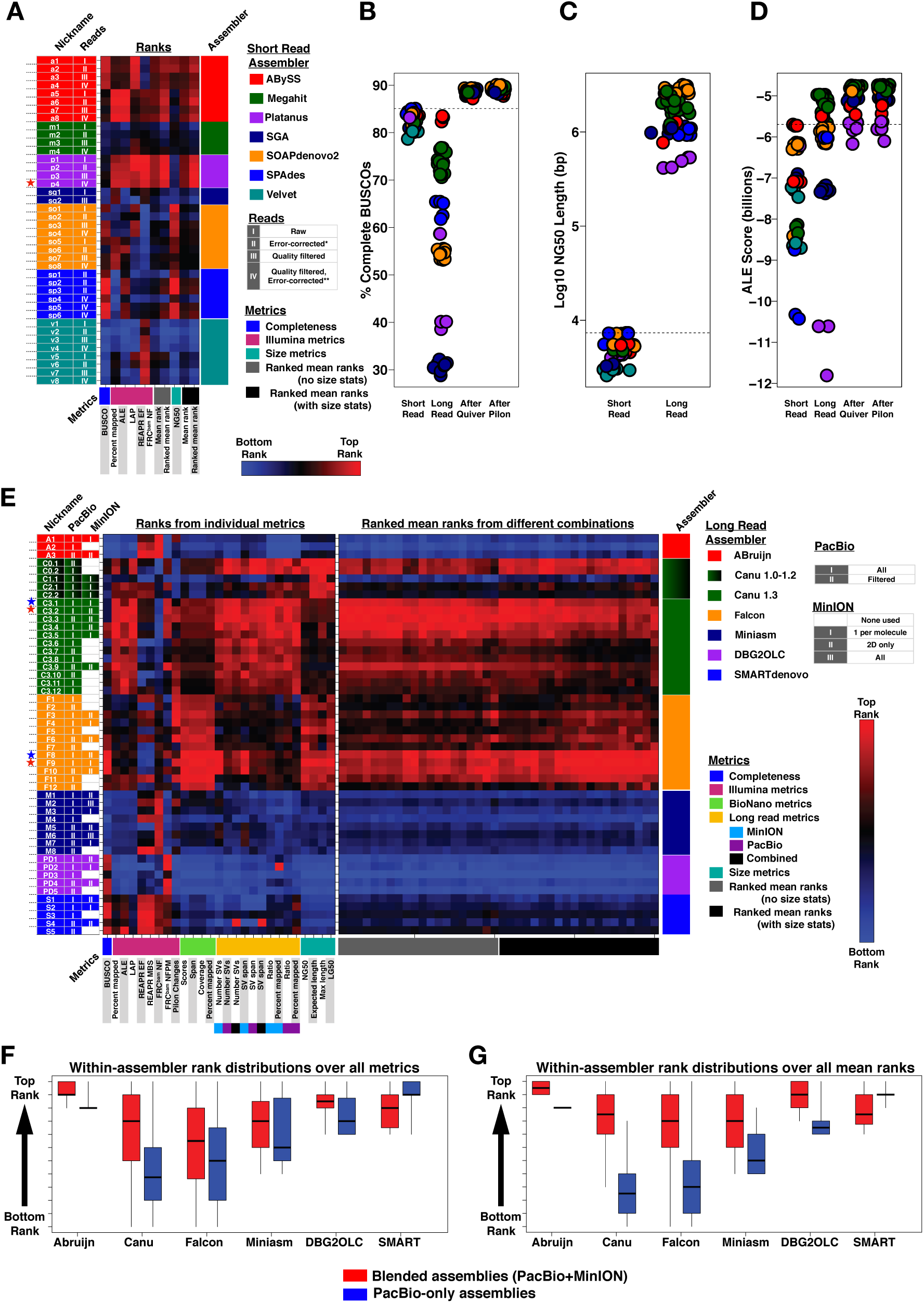
Assembly evaluations. **(A)** Rank matrix for 40 Illumina assemblies. Each column corresponds to a metric. Each row corresponds to an assembly. The columns and rows are organized by metric class and assembler, respectively. Multiple assemblies were generated for each assembler differing by the input reads, parameters used, or both. Assembly nicknames allow finding the assemblies in supplementary tables and methods. Assembly ranks for each metric span from lowest (blue) to highest (red) in each column. Assemblies (rows) that do well across the metrics tend to be mostly shades of red. The red star marks the Platanus assembly that performed best overall and was used as the input for hybrid assemblies. **(B-D)** Use the short-read assembly color scheme from (A) and the long-read color scheme from (E) to visualize **(B)** percent of complete BUSCOs found, **(C)** Log10 NG50 lengths, and **(D)** ALE scores for short-read and long-read assemblies. B and D show the long-read scores before and after polishing steps. The dotted lines in (C) represent the maximum NG50 from short-read assemblies. **(E)** Rank matrix for 50 long-read assemblies organized as described in A. Red and blue stars mark assemblies brought into BioNano scaffolding. Red stars represent the scaffolded assemblies that were chosen after BioNano scaffolding. **(F-G)** Box and whisker plots of within-assembler rank distributions comparing blended (red) to PacBio-only (blue) inputs to each assembler. The boxplots are not comparable between assemblers. The boxes show the 25^th^-75^th^ percentile, the black line is the median, and the whiskers span the range (min to max). Assemblies from a given assembler were ranked either using (F) all individual metrics from E or (G) all ranked mean ranks from different combinations of metric ranks from E. The ranks were then partitioned into those from blended versus PacBio-only assemblies. In both cases (F-G), blended assemblies from all assemblers except SMARTdenovo had significantly higher ranks by Wilcoxon Rank Sum Test than PacBio-only assemblies from the same assembler.

### Long-read datasets and assemblies

A route to obtaining more contiguous assemblies is incorporating data from single molecule, long-read technologies, such as Single Molecule Real Time (SMRT) sequencing from Pacific Biosciences (PacBio) and nanopore sequencing with the MinION from Oxford Nanopore Technologies (ONT). These technologies are more error-prone than Illumina, but the errors are approximately randomly distributed allowing high quality consensus sequences with enough coverage (Eid et al. 2009; Ip et al. 2015; Loman et al. 2015). Both long-read technologies produced read lengths that exceeded the scaffold lengths in the Illumina short-read assemblies, particularly MinION reads obtained using our modified protocols (Supplemental Figure S4; Urban et al 2015). Thus, even before attempting to assemble the long reads, we had a richer source of long-distance information than the short-read assemblies provided.

The majority of long-read coverage (50-55X total) was from PacBio (44.1X; Table 1; Supplemental Figure S4), and we were able to produce high quality assemblies using PacBio reads alone. However, despite having four times lower coverage, the MinION data (10.77X) had in excess of two times more coverage from molecules greater than 20 kb and over a hundred times more coverage from molecules exceeding 30 kb than the PacBio data (Table 1). Over 10% of the MinION data was from molecules that surpassed the longest PacBio read length of 36 kb, approximately a third of which came from high quality 2D reads (Table 1, Supplemental Figure S4). Validation of the MinION reads on assemblies generated from the PacBio data alone showed many high quality 1D and 2D reads (Supplemental Figure S5). These included hundreds of 2D reads exceeding 50 kb and several >100 kb that aligned across their full lengths with percent identities up to 94.6%. One notable 131 kb MinION 2D read aligned with 91.1% accuracy to the PacBio data. This gave us an opportunity to test whether even a small amount of ultra-long MinION reads could improve upon the PacBio assemblies. Therefore, we also generated assemblies from a blend of both single-molecule technologies, referred to here as “blended assemblies” to differentiate them from “hybrid assemblies” that refers to combining short-read and long-read technologies (Figure 1C).

In total, we generated 50 assemblies using long reads (Figure 1C), including hybrid assemblies that started from Illumina contigs. We evaluated the long-read assemblies with the same metrics used to rank the short-read assemblies (Figure 2 B-D, Supplemental Figure S1). Before polishing, ABruijn and Canu assemblies rose highest in most rankings (Figure 2 B-D, Supplemental Figure S1), perhaps because these assemblers had the best consensus sequence modules. Even before polishing, most long-read assemblies outperformed short-read assemblies for percent error-free bases (REAPR) and had comparable or better scores in other metrics (e.g. LAP, ALE, FRC). However, most underperformed in terms of gene content with fewer than 80% BUSCOs detected (Figure 2 B-D, Supplemental Figure S1).

### Long-read assembly polishing and monitoring

To ensure that the assembly evaluations primarily reflected the structural integrity of each assembly rather than differences in consensus quality, we employed extensive post-assembly polishing using Quiver (Chin et al 2013) and Pilon (Walker et al 2014) (Figure 1C). We monitored the outputs from each round of polishing using the metrics discussed above as well as the number of variants detected and changes made by the polishing algorithms (Figure 1C). The assemblies started out with up to millions of Quiver variants and converged to just a few thousand, and evaluations improved across Quiver rounds, with the biggest impact occurring in the first round (Supplemental Figure S6). After Quiver polishing, Canu assemblies continued to take many of the highest ranks whereas ABruijn assemblies lost their lead (Figure 2 B-D, Supplemental Figure S1). Quiver polishing also closed the gaps between the highest and lowest scoring assemblies in each metric. For example, whereas the percent of BUSCOs detected ranged from 30-83% prior to Quiver polishing, ∼90% were detected in all assemblies after (Figure 2B). Moreover, all polished long-read assemblies outperformed the best scoring short-read assemblies in each metric, with the exception of the hybrid assemblies that still underperformed on the ALE metric (Figure 2D). The Illumina-based metrics favored non-hybrid long-read assemblies over both the short-read and the hybrid assemblies that were constructed from the same Illumina data. This speaks to the structural and consensus quality of the contig sequences derived from long reads alone (Figure 2D, Supplemental Figure S1; “After Quiver”). Nevertheless, Illumina-polishing with Pilon improved the consensus further, fixing 19.2-25.8 thousand base and small indel errors (∼60-90 errors/Mb) in the first round, and 0.9-2.4 thousand (∼3-8 errors/Mb) in the second. The small number of corrections introduced in the final round indicates long stretches (hundreds of kb) of high-quality consensus sequences between any remaining errors in the final assemblies. Accordingly, Pilon tended to improve evaluations modestly over what Quiver had already accomplished (Figure 2B, 2D, Supplemental Figure S1; “After Pilon”). For example, it resulted in detecting up to an additional 1.05% of BUSCOs (0.63% on average).

### Selecting assemblies for BioNano scaffolding

After polishing, the number of variants or genes detected and other metrics that reflect consensus sequence quality converged to similar scores across assemblies. This allowed us to focus on the size and long-range integrity of contigs when making selections for scaffolding with optical maps. We used an expanded battery of reference-free metrics to guide our choice of which assemblies to scaffold (Figures 1C and 2E). The additional metrics were based on long reads and optical maps (see Methods). There was general agreement on assembly rankings among metrics from the four orthogonal technologies (Supplemental Figure S2B).

Long-read assembly sizes ranged from 281.5-306.6 Mb (Supplemental Table S4), close to the expected *Sciara* male somatic genome size of 280 Mb. All long-read assemblers produced assemblies that were orders of magnitude more contiguous than short-read assemblies. NG50s were typically in the Mb range and all exceeded 100 kb (Figure 2C, Supplemental Figure S1F, Supplemental Table S4). For all size metrics, assemblies from Canu and Falcon ranked highest (Figure 2C, 2E), with the largest NG50s of 3.08 Mb and 3.17 Mb, respectively (Figure 2C “Long Read”, Supplemental Table S4). Canu and Falcon assemblies also had the lowest LG50s containing 50% of the expected genome size on just 21 and 23 contigs, respectively (Supplemental Figure S1F, Supplemental Table S4). The highest normalized expected contig sizes (Salzberg et al. 2012) for assemblies from Canu and Falcon exceeded 5 Mb and the longest contigs from each exceeded 20 Mb (Supplemental Figure S1F, Supplemental Table S4).

Longer contigs can simply be a consequence of more aggressively joining reads at the cost of more misjoins. Therefore, we interrogated whether Canu and Falcon assemblies, which had the longest contigs, suffered from higher error rates. However, in direct opposition, Canu and Falcon assemblies were consistent rank leaders in our battery of evaluations (Figure 2E). Canu assemblies led most Illumina-based and long-read metrics. Falcon assemblies led BioNano metrics and gene content (Figure 2E; Supplemental Figure S1), although differences in gene content were negligible (Figure 2B). Canu and Falcon assembles had fewer putative mis-assemblies than others as proxied, for example, by long-read detection of structural variants (Supplemental Figure S7J). They also had apparently higher long-range integrity according to BioNano map alignments, which spanned a range of 237-252 Mb in Falcon and Canu assemblies, but only 181-230 Mb in others (Supplemental Figure S7H, S7J, S7L). In sum, Canu and Falcon assemblies had longer contigs and ranked higher than other assemblies in most metrics (Figure 2E), the latter arguing against more misjoins.

To select a final subset of assemblies for BioNano hybrid scaffolding, we sorted the assemblies by taking mean ranks across 40 combinations of the 27 metrics (Figure 2E). In general, blended assemblies tended to rank higher than their PacBio-only counterparts for 5 of the 6 long-read assemblers, although this often reflected modest improvements in the actual scores. The largest variation amongst scores tended to reflect the assembler used (Figure 2 F-G, Supplemental Figure S7). Blended assemblies from Canu and Falcon were the clear rank leaders again in this final analysis (Figure 2 E-G), and two assemblies from each were chosen for BioNano hybrid scaffolding. The chosen assemblies were constructed from all 44X PacBio data and either only 2D MinION reads (6.2X) or 1D and 2D reads (10.8X).

### Scaffolding with optical maps

We obtained BioNano Irys optical map data from male pupae (Figure 3, Table 1). The raw molecule N50 was 214.1 kb for molecules >150 kb. The genomic consensus maps (CMAPs) produced from them had a map N50 of 712 kb and a cumulative length of 325.5 Mb. Thus, the genome length estimated from the CMAPs was between the expected sizes of the somatic and germline genomes. The CMAPs spanned 266-278 Mb of the Canu and Falcon contigs. The CMAPs and sequence assemblies were used to produce the hybrid scaffold maps. Both the CMAPs and sequence contigs had similar spans across the hybrid scaffold maps of approximately 275-280 Mb. We found that the hybrid scaffolds derived from both Canu assemblies and from both Falcon assemblies were nearly identical as determined by evaluation metrics and whole genome alignments (Supplemental Figures S8-S9, Supplemental Table S5). Therefore, we chose the single scaffold set from each pair that was evaluated to be slightly better, hereafter referred to as “Canu” and “Falcon”. Hybrid scaffolding approximately tripled the contiguity of the assemblies (Figure 4A, Supplementary Tables S6, S7). Throughout the following text, Canu assembly statistics will be described with corresponding Falcon statistics in parentheses. The total numbers of sequences in the Canu (Falcon) assembly decreased from 1044 to 857 (713 to 608) while increasing the NG50 of 2.3 Mb to 6.7 Mb (3.5 Mb to 10 Mb). The assembly size also increased from 302 Mb to 311 Mb (296 Mb to 303 Mb) (Figure 4A-C). The Canu (Falcon) scaffolds had 187 (105) gaps summing to 8.7 Mb (6.7 Mb) with a maximum gap size of 677 kb (965 kb) and median of 20.8 kb (30.5 kb) (Supplemental Table S8).

**FIGURE 3:**
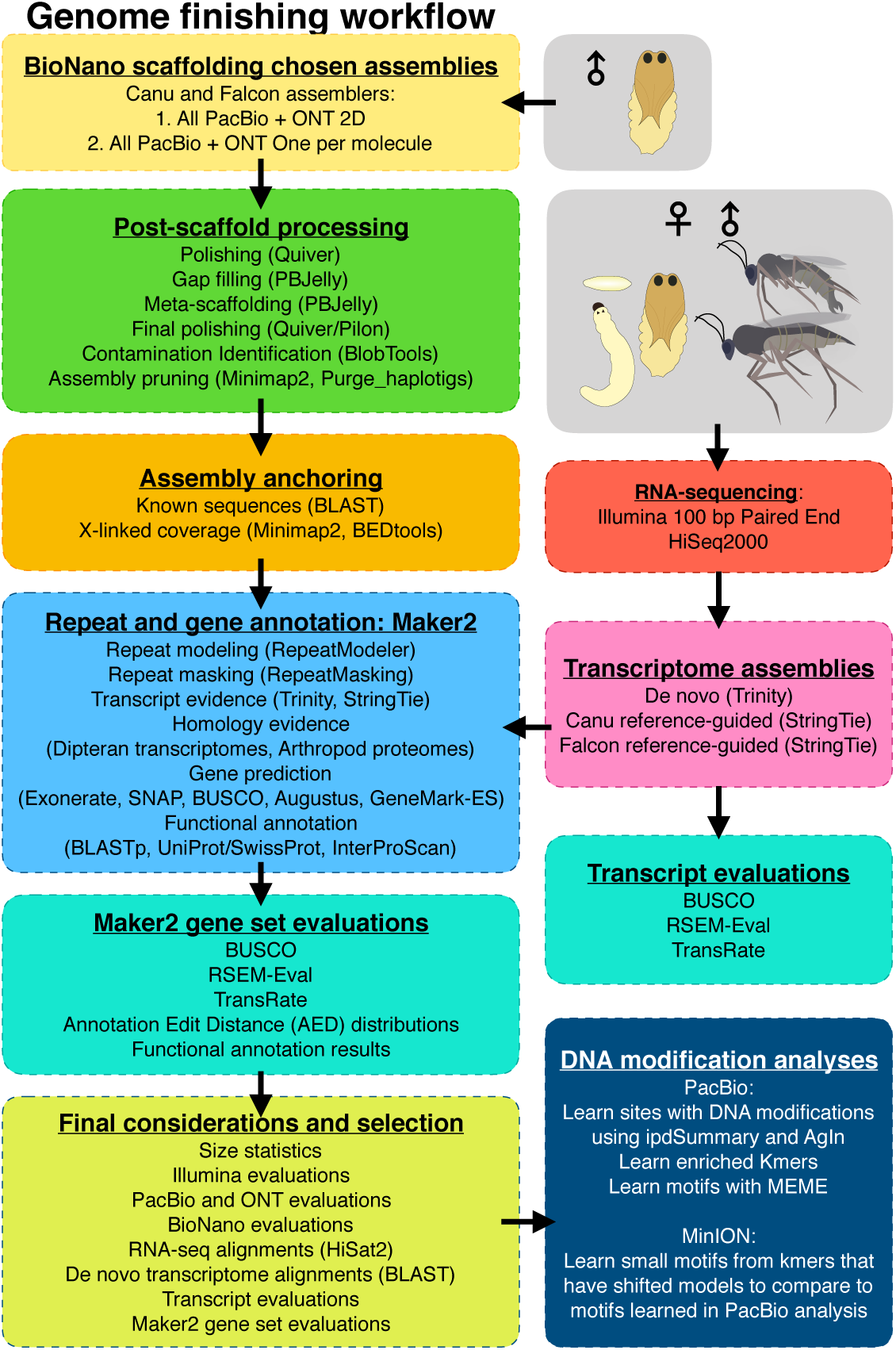
Post-assembly work flow: Workflow starting after selecting assemblies for BioNano scaffolding. Chosen assemblies were scaffolded, polished, gap-filled, filtered for contamination, anchored into chromosomes by sequences with known chromosomal addresses, and anchored to the X or autosomes by haploid or diploid coverage. Repeats were characterized and RNA-seq was used to facilitate transcriptome assembly and gene annotation. The single-molecule datasets were re-used to investigate DNA modifications.

**Figure 4:**
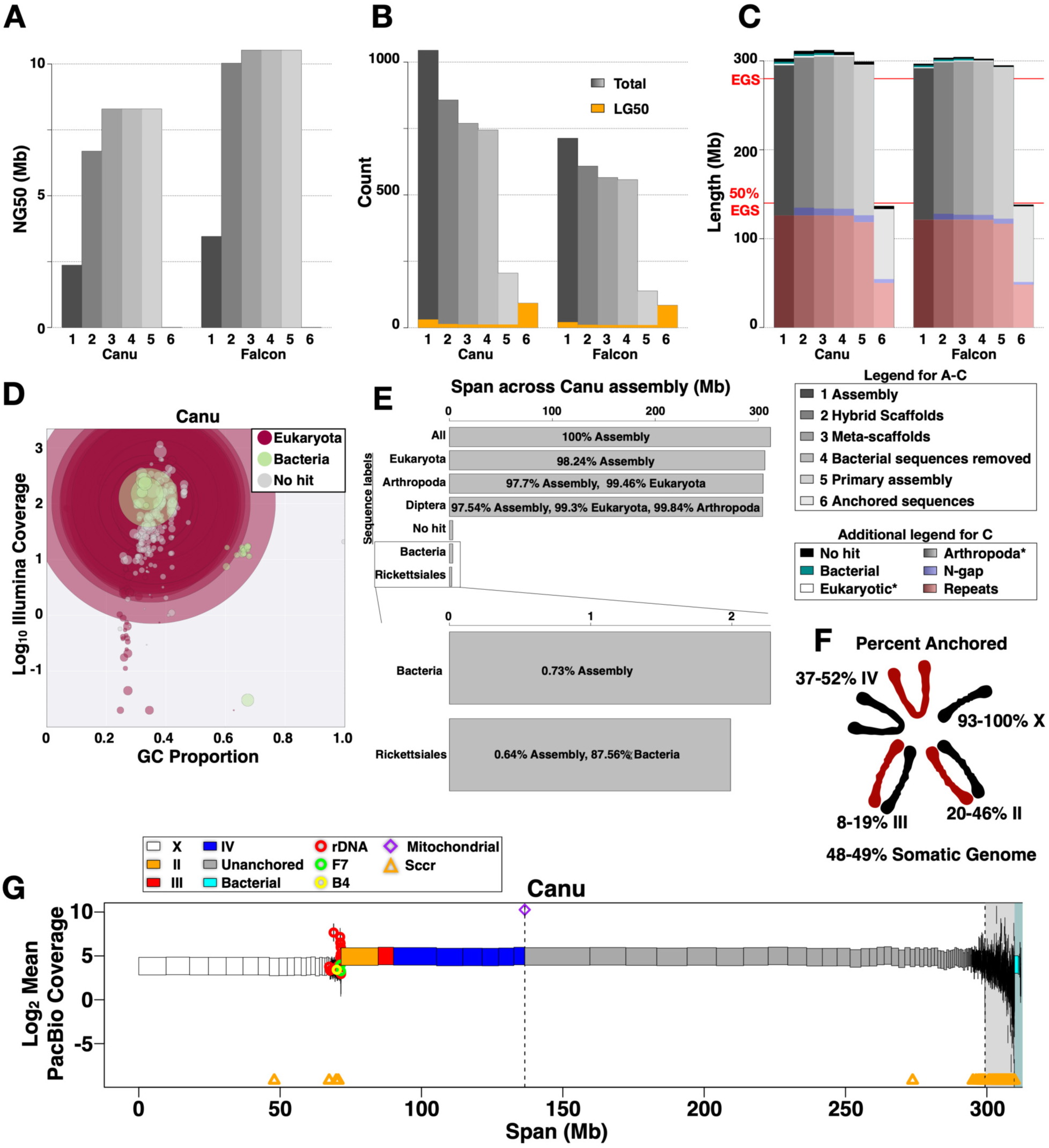
Assembly scaffolding and anchoring. **(A)** NG50 of the assembly at different stages 1-6 as defined in “Legend for A-C” within the figure. **(B)** Number of sequences in the assembly at different stages 1-6 as in A. The orange portions are LG50 counts, or the number of the longest sequences in each set needed to reach 50% of the <u>e</u>stimated <u>g</u>enome <u>s</u>ize (EGS = 280 Mb) for the somatic genome. **(C)** The total length of the assembly at different stages 1-6 as in A. The “Additional legend for C” defines colored portions of the bars. *The length of the Eukayotic and Arthropod labeled sequences includes everything up through that color. **(D)** Log10 Illumina coverage versus GC content over the Canu assembly (similar results for Falcon), colored by taxonomy information, and with circle sizes proportional to the contig sizes they represent. **(E)** The proportion of the assembly taxonomically labeled as Eukaryotic, Arthropoda, Diptera, Bacteria, and Rickettsiales. **(F)** The percentage of the expected genome size and chromosome sizes that has been anchored. Ranges represent range in Canu and Falcon assemblies. Colors as in Figure 1. **(G)** The Canu assembly with scaffolds drawn as rectangles corresponding to their lengths, colored according to the chromosome they were anchored into (or unanchored), and at their mean coverage from PacBio reads, the dataset used to determine X-linked sequences by haploid level coverage. The white background highlights sequences in the primary assembly whereas the grey and cyan backgrounds are set behind associated and bacterial sequences, respectively. All sequences to the left of the first vertical dashed line are anchored.

We next iteratively filled and polished the gaps using PBJelly (English et al. 2012) and Quiver, respectively. In the Canu (Falcon) scaffolds, 31 (14) gaps were completely closed and over 972 kb (1.06 Mb) of gap sequence was filled in (Figure 4C, Supplemental Table S8). In the final round of gap filling, we allowed PBJelly to “meta-scaffold” the hybrid scaffolds using connections from long-read alignments. This decreased the total number of sequences in Canu (Falcon) from 857 to 769 (608 to 565) while increasing the NG50 of 6.7 Mb to 8.3 Mb (10.0 Mb to 10.5 Mb) and the assembly size from 311 Mb to 312 Mb (303 Mb to 304 Mb) (Figure 4A, Supplemental Table S6, S7). We used both Quiver and Pilon to correct errors in the gap-filled meta-scaffolds. In the final round, Pilon made only 18-27 changes to the consensus sequences, translating to 1 change per 16.9 Mb and 11 Mb of non-gap sequence for Canu and Falcon, respectively.

### Assembly cleaning

BlobTools (Laetsch and Blaxter 2017) was used to identify contaminating contigs in the final scaffolds (Figure 4C-E, Supplemental Figure S10, S11). *Sciara* male embryo coverage from Illumina, PacBio, and the MinION all gave similar results (Supplemental Figure S10). The vast majority of the final Canu and Falcon scaffolds (≥97.7% of the total sequence length) was identified as Arthopoda, >99% of which was also Dipteran (Figure 4C, 4E, Supplemental Figure S11). Canu and Falcon had 25 and 8 bacterial contigs respectively, with total lengths of 2.0-2.3 Mb (<1% of the total sequence length) and bacterial contig N50s of 1.0-1.3 Mb (Figure 4C, 4D, 4E, 4G; Supplemental Figure S11, Supplemental Table S9). There were no BioNano optical map alignments over the bacterial contigs, and accordingly there were no bacterial contigs attached to or found in any of the final Arthropod-associated scaffolds. Removing bacterial contigs only marginally affected contig size statistics of the *Sciara* assemblies (Figure 4G; Supplemental Tables S6, S7).

No bacterial contigs were labeled as Delftia in the long-read assemblies despite it being the major bacterial representation in short-read assemblies. The majority of the bacterial sequence (87-96%) in the Canu and Falcon scaffolds was labeled as Rickettsiales (Figure 4D-E, Supplemental Figure S11), nearly all of which was associated with *Rickettsia prowazekii* (88.5-90.1%) and *Rickettsia peacockii* (9.9-10.8%). Given that the published genome sizes for these *Rickettsia* species range from ∼1.1-1.3 Mb (Andersson et al. 1998; Felsheim et al. 2009), it is possible that a complete *Rickettsia* genome sequence was co-assembled. The genus *Rickettsia* includes obligate intracellular bacteria that may be the closest extant relatives to the ancestor of the mitochondrial endosymbiont (Andersson et al. 1998). *Rickettsia* is closely related to *Wolbachia* that is found in many strains of *Drosophila melanogaster* (Clark et al. 2005). The *Rickettsia* or *Rickettsia*-like species in our *Sciara* datasets may be an important part of *Sciara* biology. Interestingly, in the Illumina, PacBio, and MinION datasets, the *Rickettsia* genome has nearly the same coverage as the *Sciara* genome (Figure 4D, 4G, Supplemental Figure S10). This indicates that there is approximately one *Rickettsia* genome per haploid *Sciara* genome or two *Rickettsia* for each diploid *Sciara* cell in male embryos on average. The current evidence can only suggest that this correspondence is coincidental. Despite high Rickettsia coverage in embryos, there were no *Rickettsia* optical maps from pupae. This may reflect the DNA plug isolation procedure used and/or a far lower abundance of *Rickettsia* in pupal cells or its restriction to a small subset of cells.

After removing bacterial sequences, each assembly was partitioned into “primary” and “associated” sequences. Primary sequences represent one haplotype of the genome whereas associated sequences consist of short redundant contigs called haplotigs that represent other haplotypes of specific loci (Figure 4G). The Canu (Falcon) assembly contained 744 (557) sequences of which 205 (138) were primary and 539 (419) were associated, giving a primary assembly size of ∼299 Mb (∼295 Mb) with ∼13 Mb (9.4 Mb) of associated sequences (Figure 4A-C, Supplemental Tables S6, S7). The difference of ∼4 Mb between the Canu and Falcon primary assembly sizes is in part owed to Canu having ∼2.2 Mb more gap length than Falcon. Given that the associated sequences are generally short (∼23 kb on average), computing size statistics on the primary assembly has relatively large effects on the mean and median contig sizes (Supplemental Tables S6, S7). For example, the mean scaffold size in Canu (Falcon) increased from ∼416 kb to 1.5 Mb (542 kb to 2.1 Mb).

### Assembly anchoring

We used previously known sequences with associated *in situ* hybridization results from polytene chromosomes to anchor some of the scaffolds into chromosomes (Table 2). The results span all 3 autosomes and the X chromosome. We anchored 7-8 primary autosome-linked contigs from each assembly that sum to 64.9-75.6 Mb, or ∼23-27% of the expected somatic genome size and 28-33% of the expected autosomal sequence length. Given the number of regions determined for each chromosome from polytene banding patterns (Gabrusewycz-Garcia 1964), we expect chromosomes II, III, and IV to be approximately 62-66 Mb, 66-71 Mb, and 88-94 Mb, respectively (Supplementary Table S1E). Therefore, across both assemblies we expect to have anchored 20-46% of II, 8-19% of III, and 37-52% of IV. Since it is possible to transfer anchoring information from one assembly to the other, the overall anchoring percentages for both assemblies are essentially the higher end of each range above. We also anchored between 1-2 Mb of X-linked contigs using repetitive sequences specific to the X (Table 2, e.g. rDNA). In addition to chromosome-specific sequences, “Sccr” (*Sciara* centromere consensus sequence) that hybridized to the centromeres of all *Sciara* chromosomes (Escribá et al. 2011) mapped to 48-105 contigs, the majority of which were not primary sequences (Table 2).

The majority of genomic DNA sequenced from male embryos came from somatic cells that are haploid for the X and diploid for all autosomes. Therefore, X-linked contigs could be defined as primary contigs with haploid level coverage across 80% or more of their lengths. The Canu (Falcon) assembly contained 69 (36) contigs called as X that summed to 71 Mb (62 Mb) with the longest X-linked contig reaching 9.68 Mb (12 Mb) and an X-linked contig N50 of 5.95 Mb (7.3 Mb). In both assemblies, contigs containing the X-hybridized repetitive sequences (Table 2: rDNA, B4) were called as X as expected (Figure 4G, Supplemental Figure S11C). Upon closer inspection, contigs with rDNA not called as X had haploid level coverage regions interrupted by regions of unusually high coverage over collapsed rDNA repeats, and were therefore consistent with being X-linked sequences as well. We also found X-linked contigs that contained the F7 repeat sequence known to be on X, IV, and L (Escribá et al. 2011) (Figure 4G, Supplemental Figure S11C). The X chromosome is estimated to be ∼50 Mb based on DNA-Feulgen cytophotometry or ∼62 Mb based on the number of polytene bands (Rasch 2006; Gabrusewycz-Garcia 1964; Supplementary Table S1 A-E). Therefore, >93% of the X chromosome sequence could be anchored. In total, at least 136.6-138.0 Mb of *Sciara* sequence, or ∼49% of the expected somatic genome size, was anchored into specific chromosomes with 100% of the assembly characterized as either X or autosomal from the coverage analysis.

### Repeats in the *Sciara* genome

To learn more about the repeat content of the *Sciara* genome and to facilitate repeat masking, *de novo* repeat families were created from both the Canu and Falcon assemblies using RepeatModeler (Smit and Hubley 2008). There were close to 2700 repeat families in each library of which 15-19 were classified as SINEs, 160-186 as LINEs, 48-53 as LTR, 130-131 as DNA elements, and 43-50 as other repeat classes (Figure 5A, Supplemental Figure S11D, Supplemental Tables S10, S11). The majority of repeats in each library were unclassified. For repeat masking, the *de novo* repeat libraries were combined with the few previously known repeat sequences (see Methods) as well as repeats from across Arthropods. Using this comprehensive repeat library, RepeatMasker (Smit et al. 2013) classified ∼121-126 MB or 39-41% of the Canu and Falcon assemblies as repeats (Figure 5B, Supplemental Figure S11E, Supplemental Tables S12, S13). Assuming that scaffold gaps also correspond to repeats leaves ∼180 Mb of unique sequence (∼58%) in the Canu assembly (Figure 5B). The majority (76.6-76.9%) of repeats were unclassified and spanned 93.3-96.7 Mb (Figure 5B) whereas SINE, LINE, LTR, RC, and DNA elements each constitute 0.4-3.4% of the assemblies (Figure 5B). DNA elements made up the largest class in terms of total span and Crypton-I was the largest sub-class therein (Figure 5C). However, Helitron elements from the RC class were the largest sub-class in the assembly overall (Figure 5C). Simple repeats made up ∼1% of the assemblies (Supplemental Table S12). Similar results were obtained when repeat masking with only the *de novo* repeat libraries (Figure 5C). However, using only known arthropod repeats found fewer, had a higher composition of LINE elements, and found the RTE sub-class therein to be the most abundant sub-class in the assembly (Figure 5C).

**Figure 5:**
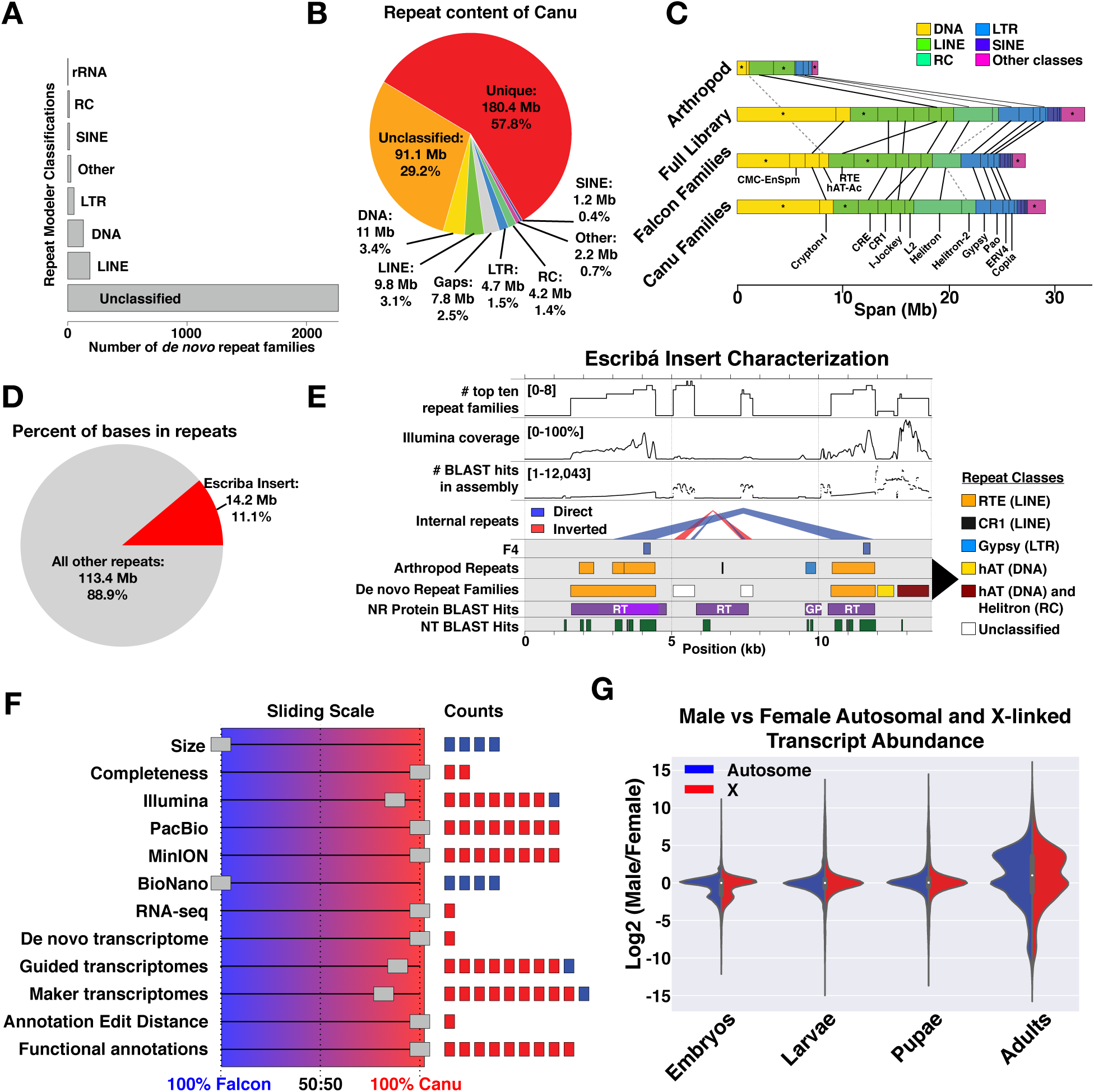
Repeats and genes in the chosen assembly. **(A)** Number of *de novo* repeat families trained on Canu with each classification. **(B)** Pie chart partition of the Canu assembly into the major repeat categories. Note that the “DNA” label used by RepeatMasker refers to DNA transposons. **(C)** The major sub-classes of repeats in each repeat class in the Canu assembly, showing the results when using different repeat libraries for masking. **(D)** Pie chart representing the number of bases masked by the Escribá insert (Escribá et al. 2011) alone compared to all masked bases. **(E)** Characterization of the Escribá insert, highlighting major repeats in the *Sciara* genome. Black arrowhead on right side pointing to repeat classes legend corresponds only to the two repeat family rows. RT = reverse transcriptase. GP = gag-pol. **(F)** The ranking results of the final two assemblies demonstrating how many metrics in each category for which each assembler scored better. **(G)** Split violin plots showing the log2 of the male to female ratio of transcript abundance for the X (red) and autosomes (blue) across multiple life stages.

Previously, Escribá et al. (2011) published a 13.8 kb lambda phage insert sequence that contains two copies of the non-LTR retrotransposon named ScRTE. A corresponding probe (F4) predominantly labeled pericentromeric regions of all *Sciara* chromosomes by FISH (Escribá et al. 2011). We found that the 13.8 kb Escribá insert contains some of the most abundant sequences in the *Sciara* genome, although there was only one full-length copy of the lambda insert in each assembly presumably from the locus that was originally cloned (Supplemental Figure S12). Otherwise, pieces of the insert were scattered across the assembly corresponding to nearly 60,000 alignments spanning ∼14.2 Mb, or ∼11% of bases labeled as repeats (Figure 5D-E). Of the top ten most abundant *de novo* repeat families, eight map to the Escribá insert across most of their lengths at sites that are consistently over-represented in DNA sequencing coverage and BLAST hits from other regions of the genome, and correspond to direct repeats of the ScRTE element, unclassified inverted and direct repeats, as well as hAT and Helitron elements (Figure 5E).

### Transcriptome assembly and gene annotation

We annotated protein-coding genes in the Canu and Falcon genome assemblies with Maker2 (Holt and Yandell 2011) guided by transcriptome assemblies from poly-A enriched RNA-seq datasets from *Sciara* male and female embryos, larvae, pupae, and adults (Figure 3). With the gene sets available from each assembly, we performed a final set of reference-free evaluations to choose a final assembly: Canu or Falcon (Figure 3). The Falcon assembly had slightly longer contig size statistics and a corresponding lead in metrics from optical map alignments (Figure 5F, Supplemental Figure S13). However, the Canu assembly outperformed Falcon in completeness metrics, RNA-seq and *de novo* transcriptome alignments, as well as metrics from Illumina, PacBio, and MinION datasets (Figure 5F, Supplemental Figure S13). Moreover, both the Canu-guided transcriptome assembly and the transcripts in the final Canu annotation received higher evaluation scores than their Falcon counterparts (Figure 5F, Supplemental Tables S14, S15). Finally, the Canu annotation had lower annotation edit distances, more genes with GO terms, Pfam domains, and/or BLAST hits in the UniProt-SwissProt database, more BUSCOs, as well as more hits with proteomes from *Drosophila melanogaster* and *Anopheles gambiae* (Figure 5F, Supplemental Figure S14, Supplemental Table S16). We conclude that the Canu assembly had higher consistency with the genome sequencing datasets and yielded the superior gene set. We therefore chose the Canu assembly as the first draft genome for *Sciara coprophila*.

The final annotation of the Canu assembly had 23,117 protein-coding gene models with 28,870 associated transcripts (Supplemental Table S15A). There are more genes than expected from other fly genomes, which may be a result of gene splitting in the annotation. To increase the quality of the *Sciara* gene set, the annotation was deposited at the i5k-workspace for community-enabled manual curation (https://i5k.nal.usda.gov/). Nevertheless, in its current form, the annotation contains nearly all expected Dipteran genes: 94.2% complete Dipteran BUSCOs were found in the final gene set, 97% when including fragmented BUSCOs (Supplemental Figure S14E, Supplemental Table S15A). The majority of genes in the annotation (87.5%) had only a single associated transcript isoform (Supplemental Figure S14B). The median gene and transcript lengths are ∼2.6 kb and ∼1.3 kb, respectively (Supplemental Table S15A). Genes had a median of 4 exons, ranging from just one (10.8% of genes) to over 100 exons. There are 10,801 genes with both 5’ and 3’ UTRs annotated and 13,335 with one or the other. Exons, introns, 5’ and 3’ UTRs had median lengths of 182 bp, 80 bp, 165 bp and 184 bp, respectively. Of all genes, we were able to attach functional information to ∼65%. Specifically, 8671 (37.5%) have Ontology Terms, 13745 (59.5%) have UniProt/SwissProt hits, 13789 (59.6%) have Pfam descriptions (El-Gebali et al. 2019), 8252 (35.7%) have all three, and 14961 (64.7%) have one or more (Supplemental Figure S14F, Supplemental Table S16). Genes spanned over 54% of the Canu assembly, mostly attributable to introns, and ∼20% of the assembly was both unique (not repetitive) and intergenic (Supplemental Figure S14H).

In the standard Dipteran model, *Drosophila melanogaster*, where males are XY and females are XX, male flies exhibit dosage compensation of X-linked genes. We used the *Sciara* gene annotation and anchoring information to explore dosage compensation in *Sciara.* The majority of cells in *Sciara* embryos, larvae, pupae, and adults are somatic where X ploidy differs between males and females. Males are haploid and females are diploid for the X, respectively. Both sexes are diploid for autosomes. We defined genes as X-linked if they were on contigs anchored into the X chromosome by the coverage analysis described above. We then determined if there was dosage compensation for X-linked genes, or if they consistently had 2-fold lower transcript abundances in male samples. Across each stage of development sequenced, the distributions of log2 fold changes between male and female transcript abundance were the same for autosomal and X-linked genes (Figure 5G, Supplemental Figure S15). There were many examples of both autosomal and X-linked genes that were differentially expressed between males and females, but there was no difference between males and females for the majority of genes in both classes. Therefore, the evidence strongly supports the existence of dosage compensation of most X-linked genes in *S. coprophila*.

### DNA modification signatures in single-molecule data

The mechanism for imprinting in *Sciara* remains unknown. Since imprinting in mammals utilizes DNA methylation (Li et al. 1993), it was if interest to determine whether DNA modifications are present in *Sciara*. The gene annotation contains the proteins involved in cytosine and adenine methylation pathways (reviewed in Armstrong et al 2019; Rausch et al 2020) that are expected to be found in Dipterans, including putative homologs for DNMT2, TET-family proteins, DAMT-1/METTL4, N6AMT1, ALKBH1, jumu, and several proteins with methyl-CpG binding domains (Supplemental Table S17A-C). There was also evidence of DNA modifications in the *Sciara* genome found using anomalies in the raw signal of both single-molecule, long-read sequencing datasets (Flusberg et al. 2010; Clark et al. 2012; Simpson et al. 2017). The high-coverage PacBio dataset was used to call site-specific modifications in the assembly for 5mC, 4mC, and 6mA. The low-coverage MinION dataset was used to find kmers with signal distributions that were shifted compared to expected models, which could result from DNA modifications. These kmers were used to find sub-motifs for comparison to motifs obtained in the PacBio analysis.

Using the PacBio SMRT kinetics data, we estimated that ∼0.13-0.24% of adenine sites in the *Sciara* male embryo genome were potentially modified with up to ∼0.04-0.06% of adenine sites exhibiting the 6-methyl-Adenine (6mA) signature (Figure 6A, Supplemental Table S18A), which is similar to 6mA densities seen for humans (∼0.05%; Xiao et al. 2018), some fungi (∼0.05%; Mondo et al. 2017), *Drosophila embryos* (0.07%; Zhang et al. 2015), and pig (0.05%; Liu et al. 2016). The tens of thousands of modified adenines were distributed ubiquitously throughout the assembly, including in genes and repeats as well as on both autosomal and X-linked sequences (Supplemental Figure S16A-C). Over 50% of the reads aligning to the majority of 6mA sites were estimated to contain 6mA (Figure 6B), suggesting that while the mark may be rare in the genome it is common at those sites. Although adenine modifications were found in many dimer and trimer contexts, AG and GAG were most enriched (Figure 6C, Supplemental Figure S16D, Supplemental Tables S19, S20). GAG sites were modified up to 7-8 times more frequently than the rate for A alone, with 0.9-1.7% GAG sites flagged as modified and 0.3-0.5% flagged as 6mA specifically (Supplemental Table S18B). The frequencies of bases surrounding the 6mA position in enriched 7mers showed a prominent 4 bp GAGG motif (Figure 6D), which did not differ between X and autosomal sequences (Supplemental Figure S17). Other motifs associated with 6mA in the *Sciara* genome included CAG within them (Supplemental Figure S18). AG, GAG, GAGG, and CAG motifs were also previously found associated with 6mA sites in human, rice, and *C. elegans* genomes (Greer et al. 2015; Xiao et al. 2018; Zhou et al. 2018). We found that 6mers defined by the sequence logo from enriched 7mers showed shifted MinION signal distributions whereas other control kmers fully agreed with the expected model (e.g. CGAGGT; Figure 6E-F, Supplemental Figure S19). From the set of all kmers with shifted MinION signals, we found similar motifs to those found in the analysis of 6mA sites identified in the PacBio analysis (Figure 6G, Supplemental Figure S18).

**Figure 6:**
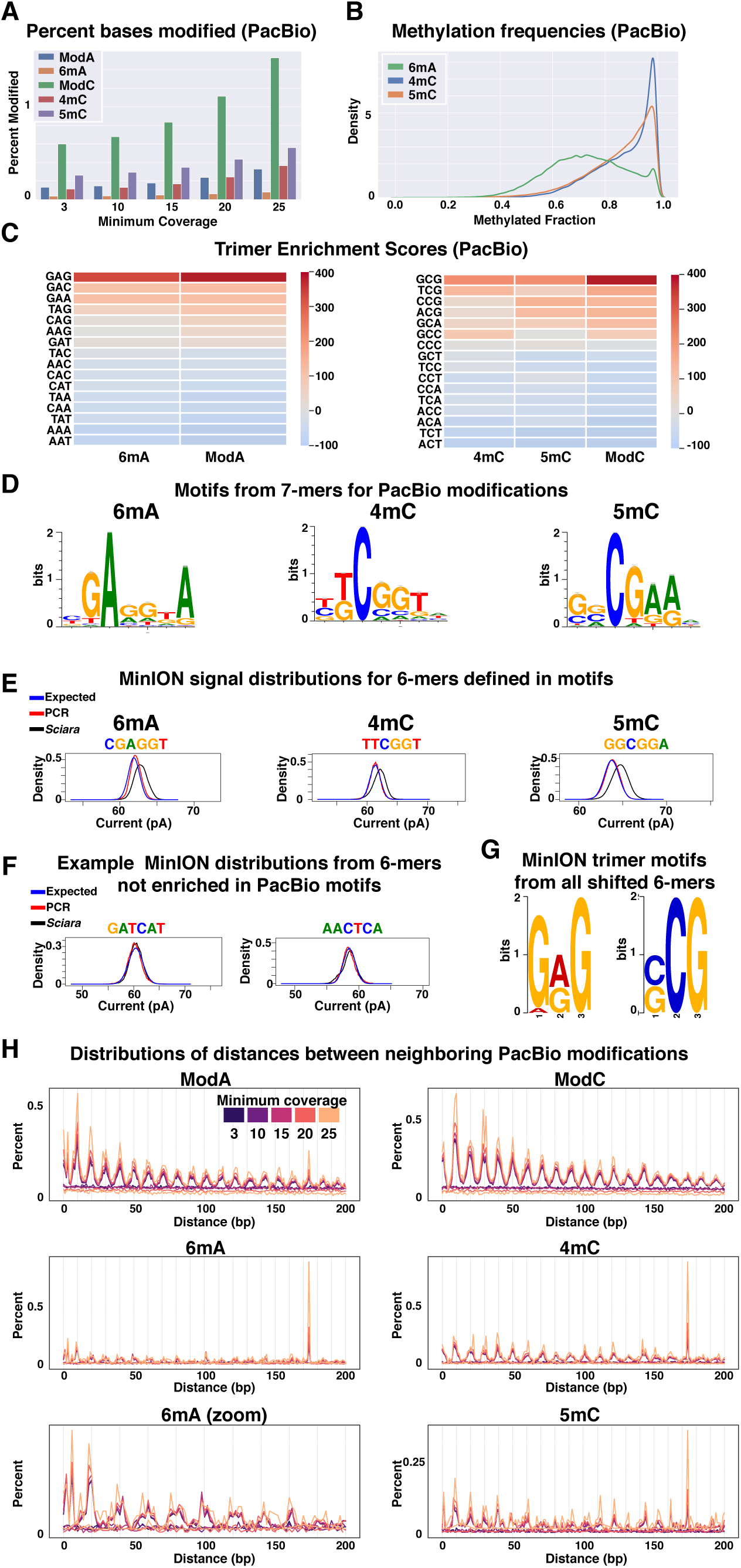
DNA modifications in male embryo genomic DNA of *Sciara coprophila*. **(A)** Percent of adenines or cytosines assigned to a modification class given a minimum coverage level in the PacBio analysis. ModA and ModC are the sets of all adenines or cytosines, respectively, flagged as modified whereas 6mA, 4mC, and 5mC are the subsets of adenines or cytosines therein with those specific classifications. **(B)** Methylation frequencies from the PacBio analysis at sites classified as having the given methylation type. **(C)** Chi-square standardized residuals (enrichment scores) indicating how many standard deviations away each observation is from expectation for trimers with middle adenines or middle cytosines from the PacBio analysis. **(D)** Position weighted motifs from the sets of 7-mers (where the modified base occurs at position 3) enriched for 6mA, 4mC, or 5mC. **(E)** The distribution of ionic current means from the MinION data for 6-mers defined by the PacBio motifs in (D). The blue line shows the expected distribution given the MinION model for each kmer. The red line shows the distribution learned from whole *E. coli* genome PCR data (Simpson et al. 2017) using only canonical nucleotides. The black line shows the distribution learned from native genomic DNA from *Sciara*. Distributions are from template reads. **(F)** As in (E), but showing examples of 6-mers not defined by motifs learned in the PacBio analysis. **(G)** Two of the top three trimer motifs learned from the set of all 6-mers that had shifted MinION signal distributions with respect to the expected models. **(H)** Distributions of distances between neighboring DNA modifications on the

We also used the PacBio SMRT kinetics data to look at cytosine methylation, which has been previously shown to mark heterochromatic regions in *Sciara* chromosomes by immunofluorescence (Eastman et al. 1980; Wei et al. 1981; Greciano et al. 2009). Up to 0.6-1.1% of cytosines were modified with up to 0.11-0.24% and 0.26-0.43% showing 4-methylcytosine (4mC) and 5-methylcytosine (5mC) signatures, respectively (Figure 6A, Supplemental Table S17C). Modified cytosines were present throughout autosomal and X-linked sequences (Supplemental Figure S16A-C). The frequency of methylation at the majority of 4mC and 5mC sites was estimated to be over 80% (Figure 6B), despite being rare in the genome overall. Modified cytosines were found in all dimer and trimers, but the most enriched were CG and GCG (Figure 6C, Supplemental Figure S16D). This is reflected in the sequence logos constructed from enriched 7mers centered on the modified C position (Figure 6D, Supplemental Tables S19, S20), and was the same for autosomes and the X (Supplemental Figure S17). Up to ∼1.3-2.5% of CpG dinucleotides were estimated to be modified with 0.26-0.57% and 0.55-0.96% specifically classified as 4mCpG and 5mCpG, respectively (Supplemental Table S18D). A more sensitive algorithm (Suzuki et al. 2016) estimated as high as 6.4% of CpG dinucleotide sites in the genome as targets for methylation in male *Sciara* embryos (Supplemental Table S18E). GCG sites were modified up to ∼4-5 times more frequently than the rate for C alone and 2 times more than CG, with 2.5-4.9% of GCG sites flagged as modified and 0.5-1.2% and 0.9-1.5% of GCG sites flagged as 4mC and 5mC, respectively (Supplemental Table S18F). Interestingly, GCG trimers are depleted in the genome sequence whereas GTG trimers are enriched (Supplemental Figure S20). This suggests that GCG may be a methylation target in the germline where 5mC deamination and conversion to thymine can deplete GCG trimers over evolutionary time. We found that 6mers defined by the sequence logos from enriched 7mers displayed shifted MinION signal distributions (e.g. for TTCGGT and GGCGGA) whereas control kmers did not (Figure 6E-F, Supplemental Figure S19), and that many motifs similar to those in the PacBio analysis specific to 4mC and 5mC were found when looking for motifs in kmers with shifted MinION signal distributions (e.g. GCG; Figure 6G, Supplemental Figure S18).

The distribution of distances between adjacent DNA modifications, for both methylated C and A, showed an enrichment of shorter distances than expected by chance (Figure 6H). There were spikes of enrichment with a periodicity of 10 bp out to distances of at least 200 bp when looking at both strand-agnostic and strand-specific spacings (Figure 6H, Supplemental Figures S21-22). This periodicity is highly suggestive of turns of the DNA helix. Periodic spacing of 10 bp between methylation sites and target motifs has been observed enriched over nucleosome positions in *Arabidopsis* and mammals (Jia et al. 2007; Chodavarapu et al. 2010; Collings and Anderson 2017). Moreover, 6mA was shown to be phased between nucleosomes in *Chlamydomonas* and *Tetrahymena* (Fu et al. 2015; Wang et al. 2017; Luo et al. 2018). Indeed, ∼175 bp is one of the most enriched distances separating two modifications in our *Sciara* male embryo data (Figure 6H, Supplemental Figures S21-22), a length reminiscent of nucleosomal spacing in general and the exact length found for nucleosome intervals in *Drosophila* (Mavrich et al. 2008).

We searched for relationships between DNA modifications and genomic features. The trends were the same for all modification types (6mA, 4mC, 5mC). With respect to annotated protein-coding genes, DNA modifications were random or slightly depleted, though there were slight depletions in exons and promoters and slight enrichments in introns (Supplemental Figure S16B, Supplemental Table S21A-B). These trends were the same when using gene locations defined by the StringTie transcriptome assembly (Supplemental Table S21C) and were generally true even when splitting the genes into categories of not expressed, lowly expressed, and highly expressed using male embryo RNA-seq data (Supplemental Table S21D). Repeat regions in the genome had more modifications than expected, and conversely the non-repeat regions had fewer than expected (Supplemental Figure S16B, Supplemental Table S21E-F). In the *de novo* repeat library, there were repeat families, including simple repeats, with 2-100 fold more modifications than expected and many families with no modifications indicating that specific classes of repeats are targeted for DNA methylation.

### Candidate germline-limited L sequences

L chromosome sequences are likely to be absent or of very low abundance in our datasets from late male embryos. Nevertheless, an effort was made to identify candidate L-sequences. Similar to identifying X-linked contigs in the assembly based on haploid-level coverage, L candidates were gathered based on very low coverage, which may include junk or redundant contigs. There were 25 contigs summing to ∼230 kb that had at most 3X PacBio coverage across their lengths in contrast to the genomic average of ∼34X. These sequences were comprised of ∼60% repeats compared with ∼40% genome-wide. The most abundant repeats were unclassified, but Gypsy, Pao, and Zator transposons were highly represented. There were 15 mRNA isoforms annotated in these low-coverage contigs corresponding to 13 genes (Supplemental Table S22). Seven genes had putative functional information. Six had best hits to UniProt-SwissProt proteins corresponding mostly Drosophila proteins, including Facilitated trehalose transporter Tret1, Ras-related protein Rab-3, Ubiquitin carboxyl-terminal hydrolase 36, RNA-directed DNA polymerase (jockey reverse transcriptase), Vitellogenin-2, Rho GTPase-activating protein. One was a protein of unknown function, but was classified in the ribosomal L22e protein family from the Pfam domain analysis.

An additional attempt to identify L candidate sequences was made by rescuing unassembled reads. Since sequences of relatively low abundance may not have been assembled, short and long reads that do not map to the Canu *Sciara* genome assembly were used to generate new assemblies using Platanus and Canu, respectively. Using the unmapped long reads, Canu returned 250 contigs summing to 2.55 Mb in length and 2247 unassembled reads summing to 7.45 Mb for a total of 10 Mb. The majority of the sequences were identified as bacterial: 89% of the total contig length and 69% of the total unassembled length for a total of 7.4 Mb of bacterial sequence (Supplemental Figure S23). Similarly, when assembling the unmapped short read pairs, Platanus returned 330 scaffolds summing to 6.8 Mb, and ∼96% was identified as bacterial (Supplemental Figure S23).

We focused on the 698 long-read sequences (1.7 Mb) and the 159 short-read scaffolds (141.5 kb) that were either identified as Arthropod or had no hits and 20-45% GC content (Supplemental Figure S23). Only 12% of the total length of these sequences aligned to the assembly, and just 14.5% and 28.3% of the targeted long-read and short-read sequences were identified as repeats. The most abundant repeats present were unclassified. The repeats were also enriched for simple repeats as well as transposons, such as Helitron (RC), Pao (LTR), RTE (LINE), Jockey (LINE), and Mariner (DNA). The centromere-associated Sccr repeat (Escribá et al 2011) was on 14 contigs. There were also contigs with rDNA and rDNA transposons R1 or R2. A small fraction of short reads contained the peri-centromeric tandem repeat B4 (Escribá et al 2011). Neither B4 nor rDNA has been observed on L chromosomes by in situ hybridization (Escribá et al 2011), suggesting that at least some of these sequences are not from L chromosomes. About 8% of the combined long- and short-read sequence length was covered by hits from 116 proteins, 16% of which were transposon-related and another 65% of which had other functional information. The most convincing alignments matched proteins (Supplemental Figure S23), such as (i) an Integrator complex subunit that is involved in snRNA transcription and processing, (ii) two zinc-finger transcription factors, (iii) a TATA-box binding protein, (iv) proteins involved with chromosome cohesion, recombination, and segregation like Wings apart-like protein (WAPL), Structural maintenance of chromosomes protein 6 (SMC6), and MOB kinase activator-like 1 (mats), and (v) proteins involved in the nervous system. The majority of these were on contigs with no matches to the genome assembly.

## DISCUSSION

### Derivation of the *Sciara* genome and anchoring

We report here the assembled sequence of the male somatic genome of the lower Dipteran fly, *Sciara coprophila*, as well as its gene annotation from transcriptomes covering both sexes and all life stages. To find the assembly approach that worked best for the *Sciara* genome, we used a battery of reference-free metrics to evaluate assemblies generated from different technologies, algorithms, inputs, and parameters. *Sciara* genome sequences assembled with Canu and Falcon from a blend of PacBio and MinION data, and polished with PacBio and Illumina data, performed best and were selected for scaffolding using optical maps from the BioNano Genomics Irys platform. Ultimately, the Canu scaffolds were the final selection for the first draft sequence owing to their higher quality gene annotation. This release of the *Sciara* male somatic genome assembly contains 299 Mb of sequence on 205 primary contigs with 50% of the expected genome size on only 12 scaffolds that range from 8.2-23 Mb long. Annotating the *Sciara* protein-coding genes with guidance from RNA-seq data gave a gene set that contained 97% of Dipteran BUSCOs, suggesting it is essentially complete. We have anchored a significant amount of the *Sciara* genome sequence on the three autosomes using previous *in situ* hybridization data, accounting for 20-46% of chromosome II, 8-19% of chromosome III and 37-52% of chromosome IV. As *Sciara* male somatic cells have only one X chromosome in contrast to two of each autosome, we were able to use coverage levels in addition to *in situ* hybridization to anchor most or all of the X chromosome sequences. In total, ∼137-138 Mb of sequence, or ∼49% of the expected genome size, was anchored into chromosomes. Future research with targeted approaches to study the L chromosome and variations associated with the X’ chromosome will be of interest beyond the current male somatic genome assembly presented here.

In its current state, the *Sciara* genome assembly is already more contiguous than up to 95% of all Arthropod genomes described (http://i5k.github.io/arthropod_genomes_at_ncbi). Its contiguity statistics exceed 42 of the 43 currently available lower Dipteran genome assemblies, over 75% of which have sub-100 kb N50s. The low contiguity of most available Dipteran genome sequences and the lack of anchoring to chromosomes limits their utility. However, the *Sciara* genome assembly presented here may be useful for scaffolding currently available and future *Nematoceran* genomes by synteny. The long contigs in the *Sciara* genome assembly reflect the success of using long read technologies and optical maps, both of which span repeats. The long-read datasets and the resulting assembly will be important and extremely useful for analyzing regions of repetitive DNA, like rDNA, centromeres, telomeres, and transposable elements.

### Comparative phylogenomics

Comparative genomics provides an understanding into the rates and patterns of evolution of genes as well as populations and species (Wiegmann and Richards 2018). The phylogenetic position of *Sciara (Bradysia) coprophila* makes its genome and transcriptome sequences valuable for future comparative genomics studies. *Sciara* is a lower Dipteran fly (Nematocera) whereas *Drosophila* is a higher Dipteran fly (Brachycera) and they diverged from one another ∼200 MYA (Wiegmann et al 2011). The *Sciara* genome size of 280 Mb (362 Mb with the L chromosomes) is larger than the 175 Mb size of the *Drosophila melanogaster* genome (Elllis et al 2014), but similar to the 264 Mb genome of the Nematoceran *Anopholes gambia* (Sharakhova et al 2007). Dipteran phylogenetics has been much studied (Hennig 1973; McAlpine and Wood 1989) but some unresolved questions remain. Previously, morphological criteria suggested that the Brachycera (containing *Drosophila*) and the Nematocera (containing *Sciara*) diverged from a common ancestor. However, more recent molecular data supports a model where the Nematoceran infraorder Bibionomorpha ultimately gave rise to the Brachycera (Wiegmann et al 2011). The *Sciara* genome and transcriptome sequences reported here will be valuable resources to further describe Dipteran phylogenetic relationships, and will further our understanding of the evolution and molecular structure of genes and pathways in Dipterans including Drosophila.

### Evolution of sex determination

The evolution of sex determination is a topic of much current interest. The most common occurrence is male heterogamety where males are XY and females are XX. In contrast, in female heterogamety, females have heteromorphic sex chromosomes (e.g., ZW), and males are homomorphic (e.g., ZZ). Female heterogamety is rare in insects (Blackmon et al 2017), but is exhibited by *Sciara* where males have a single X in their soma and females have two (Gerbi 1986). Female *Sciara* can be either XX or X’X where the X’ chromosome carries a long paracentric inversion that inhibits crossing over with the X. Thus, the heterogametic *Sciara* female determines the sex of her offspring. In *Sciara coprophila*, XX mothers have only sons and X’X mothers have only daughters. Presumably, the ooplasm is conditioned by the *Sciara* mother to determine the sex of the offspring via X chromosome elimination. In agreement with this hypothesis, sex is determined by a temperature-sensitive maternal effect that controls X-chromosome elimination in *Sciara ocellaris* (Nigro et al. 2007). As for the single X in male soma, *Sciara* males are haploid only for the X but diploid for the autosomes, unlike haplodiploid males that are haploid for their entire genome. This is accomplished by X chromosome elimination in the early *Sciara* embryo and was noted by White (1949) to occur in the Nematoceran families of Sciaridae and Cecidomyidae (including the Hessian fly *Mayetiola destructor*). Comparisons of the genomes/transcriptomes of *Sciara* and *M. destructor* might help to elucidate the molecular regulation of X chromosome elimination.

Cytoplasmic sex determination, as suggested above for *Sciara*, occurs if sex is under the control of cytoplasmic elements, such as endosymbionts. *Wolbachia* and *Rickettsia* are related groups of intracellular alpha proteobacteria that can distort the sex ratio of their arthropod hosts (Lawson et al, 2001, Serbus et al 2008). They are transmitted through the egg cytoplasm and alter reproduction in their arthropod hosts in various ways, including cytoplasmic incompatibility, feminization of genetic males, and male killing (Werren and Windsor 2000; Serbus et al 2008). Both can induce parthogenesis (Blackmon et al 2017). The latter is of interest since (i) parthenogenetic *Sciara* embryos have been observed, but their development arrests in embryogenesis (de Saint Phalle and Sullivan 1998), and (ii) although we did not find *Wolbachia* sequences in *Sciara* genomic DNA, we essentially co-assembled an entire *Rickettsia* genome. Moreover, our genomic copy number analyses suggest there are two *Rickettsia* cells per *Sciara* cell on average in 1-2 day old male embryos. Further evidence is needed to ascertain if *Rickettsia* plays a role in *Sciara* sex determination.

### Paternal chromosome imprinting

The first example of a chromosome or a chromosomal locus “remembering” its maternal or paternal origin was noted in *Sciara* and the term “imprinting” was coined (Crouse 1960). Specifically, in *Sciara* male meiosis I, the paternally derived chromosomes move away from the single pole of the naturally occurring monopolar spindle and are discarded in a bud of cytoplasm. This is an example of paternal genome elimination (PGE) that can give rise to haplodiploidy in other systems (Blackmon et al 2017). Thus, only the maternal genome is passed down through sperm in *Sciara*. Although sperm in *Sciara* is haploid for autosomes, it is diploid for the X chromosomes due to non-disjunction of the X in *Sciara* male meiosis II (Gerbi 1986). After fertilization of the haploid egg, diploidy is re-established for the autosomes, but the X chromosome is temporarily triploid. Either one or both copies of the paternally-derived X are eliminated from female or male embryos, respectively, during the 7^th^-9^th^ embryonic cleavage division, representing another example of imprinting in *Sciara* (de Saint Phalle and Sullivan 1996). Nevertheless, the mechanism for imprinting in *Sciara* remains elusive. It is of interest to learn if DNA modifications occur in *Sciara* since different imprints in mammalian genomes are laid down in eggs and sperm through a DNA methylation mechanism, leading to differential gene expression at imprinted loci in the offspring (Li et al 1993).

DNA methylation typically occurs at CpG sites where it is established *de novo* by DNA methyltransferase 3 (DNMT3) and is maintained by DNMT1 (Goll and Bestor 2005, Kato et al 2007). In contrast to vertebrates, DNA methylation in invertebrates is relatively sparse (Bird 1980). DNMT1 is found in all orders of insects except Diptera, which also lack DNMT3 (Bewick et al 2017). In agreement, our gene annotations suggest that *Sciara* also lacks DNMT1 and DNMT3. Some bisulfite sequencing studies revealed that CpG DNA methylation is found in all insect Orders except Dipteran flies (Bewick et al 2017) and failed to find specific patterns for methylated C in *Drosophila* embryos (Zemach et al 2010; Raddatz et al 2013). Other studies have asserted that *Drosophila melanogaster* has DNA methyltransferase activity and CpC methylation (Panikar et al 2015), has low levels of 5-methylcytosine (5mC) (Capuano et al 2014, Takayama et al 2014, Deshmukh et al. 2018), and has more cytosine methylation in stage 5 *Drosophila* embryos than oocytes (Takayama et al 2014). Moreover, 6-methyladenine (6mA) has been recently reported to be in the genomic DNA of *Drosophila* and other eukaryotes (Fu et al. 2015; Greer et al. 2015; Zhang et al. 2015; Liu et al. 2016; Mondo et al. 2017; Wang et al. 2018; Xiao et al. 2018). Typically, the level of 6mA is quite low, such as 0.001% in *Drosophila* but rises to 0.07% in early embryos (Zhang et al 2015). DAMT-1 appears to be the methyltransferase for 6mA in insect cells and DMAD has 6mA demethylating activity in *Drosophila* (Luo et al 2015, Zhang et al 2015). Our gene annotations suggest that *Sciara* has both DAMT-1 and DMAD.

Before it can determined whether or not imprinting in *Sciara* involves DNA modifications, it needs to be determined if the *Sciara* genome harbors DNA modifications at all. Previous immunofluorescence studies have suggested the presence of 5-methylcytosine in *Sciara* chromosomes (Eastman 1980, Greciano 2009). Similarly, our sequencing data support the presence of base modifications in the *Sciara* genome. Overall, up to 0.6-1.1% of cytosines may be modified in the *Sciara* genome, especially at GCG sites, with specifically 0.1-0.2% and 0.3-0.4% identified as 4mC and 5mC, respectively. In addition, 0.13-0.24% of adenine sites in the *Sciara* male embryo genome were potentially modified with up to ∼0.04-0.06% of adenine sites containing 6mA, especially GAG sites. Moreover, the PacBio analysis suggests that these DNA modifications are phased with 10 bp and 175 bp periodicities, suggesting physical interactions between the 10 bp turns of the DNA helix and methylation machinery as well as relationships with nucleosome spacing, both of which have been seen previously using orthogonal methods (Jia et al. 2007; Chodavarapu et al. 2011; Fu et al. 2015; Collings and Anderson 2017; Wang et al. 2017; Luo et al. 2018). Lastly, the distribution of modifications we observed with respect to genes and repeats are concordant with previous observations (i) of methyl-C in *Drosophila* (Takayama et al. 2014) and (ii) that heterochromatic regions of the *Sciara* genome, where most repeats reside, are enriched for 5mC (Eastman et al. 1980; Greciano et al. 2009). Overall, the evidence from single-molecule sequencing lends support to the presence of methylated cytosines and adenines in the autosomes and X chromosome in the male embryo genome of *Sciara*. However, the analyses suggest that the levels of DNA modifications are low. Their abundance in females and other developmental stages and tissues as well as their biological significance remains to be determined in future investigations. Nevertheless, given the evidence from the current study and previous work, base modifications may be a promising avenue for the study of imprinting in *Sciara*.

### Summary

The *Sciara* genome sequence provides a foundation for future studies to delve into the many unique biological properties of *Sciara* (reviewed by Gerbi 1986) that include **(i)** chromosome imprinting; **(ii)** sex determination by the mother; **(iii)** a monopolar spindle in male meiosis I; **(iv)** non-disjunction of the X chromosome in male meiosis II; **(v)** chromosome elimination in early embryogenesis; **(vi)** germ line limited L chromosomes; **(vii)** DNA amplification in late larval salivary gland polytene chromosomes; **(viii)** high resistance to radiation.

## METHODS

### Tissue collection, DNA extraction, and DNA sequencing

*Sciara coprophila* (renamed *Bradysia coprophila)* was used for these studies. *Sciara* (stock: Holo2) matings were performed to produce only male offspring from which embryos aged 2 hours – 2 days (genome sequencing datasets) or pupae (BioNano Irys genome mapping datasets) were collected. For a minority of MinION sequencing data, adult males were used. Genomic DNA (gDNA) was isolated using DNAzol (ThermoFisher) as per the manufacturer’s instructions with some modifications. gDNA was cleaned with AMPure beads (Beckman Coulter). Purity was checked with NanoDrop (ThermoFisher) and concentration was checked with Qubit (ThermoFisher).

For Illumina HiSeq 2000 sequencing, male embryo gDNA was sonicated to a size range of 100-600 bp, prepared using the NEBNext kit (New England Biolabs) following the manufacturer’s directions, run on a 2% NuSieve agarose (Lonza) gel, size-selected near the 500 bp marker, gel purified (Qiagen), and sequenced to obtain 100 bp paired-end reads.

For Pacific Biosciences RSII Single Molecule Real Time sequencing datasets (P5-C3 chemistry), male embryo gDNA was given to the Technology Development Group at the Institute of Genomics and Multiscale Biology at the Icahn School of Medicine at Mount Sinai for library construction and sequencing. Two DNA libraries were prepared and sequenced across 24 SMRTcells as described further in the Supplemental Methods.

MinION data was collected using multiple early iterations of the technology (original MinION and MkI), kits (SQK-MAP002, MAP004, MAP005, MAP006), and pores (R7.3 and R7.3 70 bps 6mer). We prepared 15 libraries from male *Sciara* embryo gDNA (making up >97% of the data) and 2 from male adult gDNA. The manufacturer’s instructions were followed with modifications to increase read lengths (Urban et al. 2015 and Suppl. Methods). Libraries were loaded onto the MinION, sequenced, and basecalled with Metrichor. Reads were extracted from Fast5 files and analyzed using our own custom set of tools (Fast5Tools: github.com/JohnUrban/fast5tools).

For BioNano Genomics (BNG) Irys optical maps, male pupae were flash frozen ground in liquid nitrogen and high molecular weight gDNA was isolated (Suppl. Methods), nicked with BssSI (CACGAG, New England BioLabs), labeled, and repaired according to the IrysPrep protocol (BioNano Genomics).

### Genome assemblies

After optional trimming/filtering with Trimmomatic (Bolger et al 2014) and/or error-correction with BayesHammer (Nikolenko et al 2013), short-read assemblies were generated using ABySS (Simpson et al. 2009), Megahit (Li et al. 2015), Platanus (Kajitani et al. 2014), SGA (Simpson and Durbin 2010), SOAP (Luo et al. 2012), SPAdes (Bankevich et al. 2012), Velvet (Zerbino and Birney 2008). Hybrid assemblies were generated using DBG2OLC (Ye et al. 2016) and PBDagCon (http://bit.ly/pbdagcon) starting with Platanus contigs and long reads. Non-hybrid long-read assemblies were generated with Canu (Koren et al. 2017), Falcon (Chin et al. 2016), Miniasm (Li 2016) with RaCon (Vaser et al. 2017), ABruijn (Lin et al. 2016), and SMARTdenovo (https://github.com/ruanjue/smartdenovo). For all assemblers, we varied filtering, error correction, inputs, and parameters as detailed further in the Suppl. Methods. Long-read assemblies were polished with Quiver (Chin et al. 2013) and Pilon (Walker et al. 2014). BlobTools (Laetsch and Blaxter 2017) was used to identify contaminating contigs.

### Assembly evaluations

Assembly evaluations included subsets of the following: contig size statistics, percent of Illumina reads that mapped using Bowtie2 (Langmead and Salzberg 2012), probabilistic scores from LAP (Ghodsi et al. 2013) and ALE (Clark et al. 2013), number of features from FRC^bam^ (Vezzi et al. 2012), percent error-free bases and/or the mean base score from REAPR (Hunt et al. 2013), completeness of gene content with BUSCO (Simão et al. 2015), the percent of long reads that aligned with BWA (Li and Durbin 2009), the average number of split alignments per long read, structural variations using Sniffles (Sedlazeck et al. 2018), the percent of raw BioNano map alignments using Maligner (Mendelowitz et al. 2015), the resulting optical map alignment M-scores, the number of bases covered by optical maps (span), and the total coverage from aligned optical maps. Evaluations were automated and parallelized on SLURM with a custom package (github.com/JohnUrban/battery).

### Scaffolding

For hybrid scaffolding, optical maps >150 kb were assembled into consensus maps (CMAPs) using BioNano Pipeline Version 2884 and RefAligner Version 2816 (BioNano Genomics). Each selected assembly was used with the BNG CMAPs to create genome-wide hybrid scaffolds using hybridScaffold.pl version 4741 (BioNano Genomics). Quiver and PBJelly (English et al. 2012) were used to polish and gap-fill the scaffolds. PBJelly was used additionally to join more scaffolds with long-read evidence into “meta-scaffolds”, and Quiver and Pilon were used for final polishing.

### Assembly anchoring

Haplotigs were identified using Minimap2 (Li 2018) and purge_haplotigs (Roach et al. 2018). To anchor contigs into chromosomes, sequences that were previously mapped to chromosomes experimentally were mapped to the assemblies using BLAST (Altschul et al. 1990). Differentiating between autosomal and X-linked contigs was performed by requiring haploid coverage levels across at least 80% of a contig to be called as X-linked (else autosomal), using Minimap2 and BEDTools (Quinlan and Hall 2010).

### Transcriptome assemblies

For strand-specific RNA-sequencing libraries, poly-A RNA was prepared from a given sex and stage using TRIzol (Invitrogen/Thermofisher), DNase (Qiagen), RNeasy columns (Qiagen), and Oligo-dT DynaBeads (Life Technologies). RNA integrity was evaluated on 1.1% formaldehyde 1.2% agarose gels. RNA purity and quantity were measured with the NanoDrop (ThermoScientific) and Qubit (ThermoFisher) throughout. Libraries were prepared from poly-A RNA using NEB’s Magnesium Fragmentation Module, SSIII (Invitrogen) first strand synthesis with random primers, NEBNext Second Strand Synthesis module with ACGU nucleotide mix (10 mM each of dATP, dCTP, dGTP, and 20 mM of dUTP), NEBNext End Repair and dA-Tailing (NEB), and ligation (NEB: NEBNext Quick Ligation Reaction Buffer, NEB Adaptor, Quick T4 Ligase). The libraries were size-selected with AMPure beads (Beckman Coulter). Uracil-cutting for strand-specificity (and hairpin adapter cutting) was performed with NEBNext USER enzyme, followed by PCR using NEBNext High-Fidelity 2X PCR Master Mix and NEBNext indexed and universal primers for 12 cycles. A final size-selection of PCR products was performed with AMPure beads. Purity, quantity, and size of the libraries were checked with NanoDrop, Qubit and Fragment Analyzer (Agilent). Traces suggested the mean estimated fragment sizes was around 420 bp, indicating mean insert sizes near 300 bp. Libraries were sequenced to yield 100 bp paired-end reads using the Illumina HiSeq 2000. The strand-specific RNA-seq datasets were combined and assembled with Trinity (Grabherr et al. 2011) or using HiSat2 (Kim et al. 2019) and StringTie (Pertea et al. 2015). Transcriptome assemblies were evaluated with BUSCO (Simão et al. 2015), RSEM-Eval (Li et al. 2014), and TransRate (Smith-Unna et al. 2016).

### Repeat and gene annotation

Species-specific repeat libraries were built using RepeatModeler (Smit and Hubley 2008). These were combined with previously known repeat sequences from *Bradysia coprophila* as well as all Arthropod repeats in the RepeatMasker Combined Database: Dfam_Consensus-20181026 (Hubley et al. 2016), RepBase-20181026 (Bao et al. 2015). To predict protein-coding genes, Maker2 (Holt and Yandell 2011) was used with transcriptome evidence described above, transcript and protein sequences from related species for homology evidence, Augustus (Hoff and Stanke 2019), SNAP (Korf 2004), GeneMark-ES (Ter-Hovhannisyan et al. 2008), and RepeatMasker (Smit et al. 2013) with repeat libraries described above. InterProScan (Quevillon et al. 2005) was used to identify Pfam domains and GO terms from predicted protein sequences, and BLASTp was to find the best matches to curated proteins in the entire UniProtKB/Swiss-Prot database (The UniProt Consortium 2019). Maker2 transcriptomes were evaluated using annotation edit distances, BUSCO, RSEM-Eval, and TransRate.

### DNA modification analyses

PBalign (github.com/PacificBiosciences/pbalign) with BLASR v2 (Chaisson and Tesler 2012) was used to align PacBio reads to the entire unfiltered assembly to avoid forcing incorrect mappings. Pbh5tools (github.com/PacificBiosciences/pbh5tools) was used to merge and sort the mapped reads. ipdSummary from kineticsTools v0.6.0 (github.com/PacificBiosciences/kineticsTools) was used to predict base modifications across the Canu genome assembly (--pvalue 0.01 --minCoverage 3 --methylMinCov 10 --identifyMinCov 5). AgIn (Suzuki et al. 2016) was also used to look at CpG methylation. For all analyses on predicted DNA modifications, we used only primary contigs labeled as Arthopoda. Kmer enrichment scores for dimers and trimers were obtained from the Chi-square standardized residuals found when comparing the distribution of kmers that had a specific modification at a fixed position with the genome-wide distribution of kmers with the target base at that position. We also used this approach to define enriched 7-mers for position weight matrix motifs using WebLogo (Crooks et al. 2004). In addition, the 9 bp sequences centered on the top 500 or 5000 scoring specific modification calls were used with MEME (Bailey and Elkan 1994) to identify motifs using a second order Markov model background file trained on the *Sciara* genome assembly (fasta-get-markov -m 2 -dna). We determined if DNA modifications were enriched/depleted in various genomic regions using binomial models. When separating genes by expression level for this analysis, Salmon (Patro et al. 2017) was used to quantify expression over our Maker2 protein-coding gene annotation using male embryo RNA-seq. BEDtools was used to obtain spacing distances between modified bases as well as between random bases of the same type (e.g. m6A vs random A). Although 10 bp periodicities were obvious by visual examination, we formally determined the periodicities observed in counts of inter-modification distances between 0-200 bp by running a discrete Fourier transform (DFT) analysis using the Fast Fourier Transform (FFT) from Python’s Numpy package.

For the MinION analysis, only datasets generated from the MkI, SQK-MAP006 kit, and R7.3 70 bps 6mer pore model were used, and only reads that aligned to primary contigs annotated as Arthropoda. We compared the signal distributions for each kmer in our *Sciara* dataset to the expected ONT kmer models, and to a MinION dataset generated from whole genome PCR on *E. coli* genomic DNA using the same kit and pore model (BioProject PRJEB13021; Run ERR1309547; www.ebi.ac.uk/ena; Simpson et al. 2017). MinION reads were aligned with BWA. Nanopolish (Simpson et al. 2017) was used to learn updated kmer models from the native *Sciara* and E. coli PCR MinION datasets. MEME was used to identify short motifs in all 6mers that differed from the expected ONT model.

### Further bioinformatics

The Supplemental Methods contains software versions, as well as further details and exact commands for: read processing, genome assembling, polishing, evaluating, scaffolding, gap filling, bacterial filtering, haplotig filtering, anchoring, transcriptome assemblies and evaluations, repeat library construction, repeat-masking, training gene predictors, alternative transcript and protein evidence, Maker2 iterations and evaluations, and the PacBio and MinION DNA modification analyses. Bioinformatics analyses were largely aided by custom scripts located at github.com/JohnUrban/sciara-project-tools, github.com/JohnUrban/fast5tools, github.com/JohnUrban/battery, github.com/JohnUrban/lave, and github.com/JohnUrban/fftDnaMods.

## Supporting information

All supplemental materials including supplemental figures, tables, and methods.

## DATA ACCESS

Raw Illumina (DNA and RNA-seq), PacBio, MinION, and BioNano data generated in this study as well as BioNano CMAPs and PacBio kinetics and DNA modification results have been submitted to the NCBI BioProject database (http://www.ncbi.nlm.nih.gov/bioproject/) under accession number PRJNA123456. This Whole Genome Shotgun project has been deposited at DDBJ/ENA/GenBank under the accession VSDI00000000, and the Canu assembly version selected as the first draft genome release in this paper (Bcop v1.0) is version VSDI01000000. The automated Bcop_v1.0 annotation for the Canu assembly is available at the i5k Workspace (i5k.nal.usda.gov) where manual curation updates will be made.

## ACKNOWLEDGMENTS

We thank Oxford Nanopore for early access to the MinION and for strong continual support, particularly from Michael Micorescu, Sissel Juul, Daniel Turner, Stuart Reid, David Stoddart, Margherita Coccia, Richard Ronan, and Jackie Evans. We thank the members of the nanopore community, and particularly members of the MinION Access and Reference Consortium (MARC), for lively discussions. Thanks to Benjamin Raphael for helpful discussions on genome assembly and the Center for Computational Molecular Biology (CCMB) at Brown University for providing computing resources for our MinION. We thank the Technology Development Group at the Institute of Genomics & Multiscale Biology at the Icahn School of Medicine at Mount Sinai, particularly Gintaras Deikus for help in obtaining PacBio data, and Ali Bashir and Robert Sebra for discussions on PacBio chemistries, and genome assembly. Thanks to Adam Phillippy, Sergey Koren, and Brian Walenz for helpful correspondence and guidance about Canu and genome assembly in general. Thanks to Mark Howison, Stefano Lonardi and Stephen Richards for helpful correspondence and sharing experiences with various assemblers and tools. Thanks to Jared Simpson for providing guidance on Nanopolish. Thanks to Jennifer Urban for help drawing *Sciara* cartoons, to Yutaka Yamamoto for *Sciara* photographs, and to Miiko Sokka, Steven DeLuca, Ethan Greenblatt, and other members of the Spradling and Gerbi laboratories for useful discussions and comments. Thanks to Kevin Urban at Early Signal for valuable insights on FFT analysis. Thanks to the Center for Computation and Visualization (CCV) at Brown University, NSF EPSCoR, and Carnegie Science’s Scientific Computing Committee for High-Performance Computing for computational resources and support. This work was supported by NSF/MCB-1607411 and NIH/GM121455 to SAG; predoctoral traineeships to JMU from NIH/T32-GM 007601, NSF/EPSCoR #1004057, and NSF predoctoral fellowship GRFP-DGE-1058262; NIH/P20 GM103418 to SJB; and funding through the Howard Hughes Medical Institute (ACS, JMU).

## AUTHOR CONTRIBUTIONS

John Urban (JMU) collected all embryos, larvae, pupae, and adult Sciara needed for all experiments. JMU prepared all MinION libraries and performed all MinION sequencing and analyses. JMU wrote the suites of tools for working with MinION data (https://github.com/JohnUrban/fast5tools), automating the battery of assembly evaluations (https://github.com/JohnUrban/battery), genome alignment visualizations (https://github.com/JohnUrban/lave), and all general bioinformatics over the course of this project (https://github.com/JohnUrban/sciara-project-tools). JMU obtained high molecular weight genomic DNA and delivered it to the Technology Development Group at the Institute of Genomics & Multiscale Biology at the Icahn School of Medicine at Mount Sinai, where PacBio sequencing libraries were prepared and sequenced. JMU performed all short- and long-read assemblies, genome polishing, assembly evaluations, repeat modeling and annotation, and gene annotation. JMU did all RNA work and library preparations for all RNA-seq samples representing replicates from both sexes at different stages, and performed all transcriptome assemblies and RNA-seq data analysis. JMU performed DNA modification analyses with PacBio single molecule kinetics data and MinION single molecule ionic current data. CMC made DNA plugs from *Sciara* pupae collected and sent to her by JMU, and performed the BioNano preparations and imaging on Irys platform. RM and NL performed BioNano hybrid scaffolding with selected assemblies sent to them. SJB provided guidance in our acquisition of BioNano data and provided oversight to CMC, RM, and NL. MSF prepared the Illumina DNA library. JEB did all *Sciara* mass matings. ACS provided support and guidance on this work. SAG pioneered and guided the *Sciara* genome effort. JMU conceived the experiments and analyses. JMU and SAG wrote the manuscript.

