## Supplementary material for "Single-molecule sequencing of long DNA molecules allows high contiguity *de novo* genome assembly for the fungus fly, *Sciara coprophila*": All supplemental materials including supplemental figures, tables, and methods.

**Table of Contents:**
**Pages 2-13**

**Glossary:**
**Pages 14-15**

#### Section 1: Supplemental Figures
#### Pages 16-41

**Supplemental Figure S1:**
Comparing evaluations of short read assemblies to long read assemblies
Page 16

**Supplemental Figure S2**:
Assembly ranking correlation matrices
Page 17

**Supplemental Figure S3**:
Filtering out non-Arthropod, contaminating reads using Taxonomy- annotated GC plots
Page 18

**Supplemental Figure S4:**
Length Distributions for Illumina Scaffolds, PacBio Reads and MinION Molecules
Page 19

**Supplemental Figure S5:**
Percent identity of MinION reads compared to a PacBio-only assembly
Page 20

**Supplemental Figure S6:**
Evaluations across Quiver polishing rounds
Page 21

**Supplemental Figure S7:**
Blended assemblies with both PacBio and MinION data tended to receive better ranks than PacBio-alone assemblies
Page 22

**Supplemental Figure S8:**
Metrics comparing assemblies after scaffolding
Page 23

**Supplemental Figure S9:**
Aligning chosen and discarded scaffolds from each assembler (Canu and Falcon)
Page 24

**Supplemental Figure S10:**
BlobTools analysis of the Canu scaffolds
Page 25

**Supplemental Figure S11:**
BlobTools analysis and anchoring Falcon scaffolds
Page 26

**Supplemental Figure S12:**
The single locus that contains the full-length Escribá insert
Page 27

**Supplemental Figure S13:**
Pairwise comparisons of final Canu and Falcon scaffolds
Page 28

**Supplemental Figure S14:**
Pairwise comparisons of final Canu and Falcon annotations
Page 29-30

**Supplemental Figure S15:**
Dosage compensation of X-linked genes in *Sciara coprophila*
Page 32-33

**Supplemental Figure S16:**
Distribution of DNA modifications across *Sciara* genome (PacBio analysis)
Page 33

**Supplemental Figure S17:**
Position weighted 7-mer motifs learned from different filtering and different subsets of the genome sequence (PacBio analysis)
Page 34

**Supplemental Figure S18:**
MEME motifs in the PacBio and MinION analyses
Page 35

**Supplemental Figure S19:**
MinION signal distributions for 6mers defined by motifs learned in the PacBio analysis and negative controls
Page 36

**Supplemental Figure S20:**
The GCG trimer is depleted in the genome and transcriptome compared to expectation
Page 37

**Supplemental Figure S21:**
Distribution of distances between adjacent DNA modifications (PacBio analysis) on the same strand shows enrichment of short distances, a 10 bp periodicity, and a spike of enrichment at mono-nucleosome lengths of ~175 bp
Page 38,40

**Supplemental Figure S22:**
Distribution of distances between adjacent DNA modifications (PacBio analysis) on either strand also shows enrichment of short distances, a 10 bp periodicity, and a spike of enrichment at mono-nucleosome lengths of ~175 bp for 6mA and 5mC
Page 39-40

**Supplemental Figure 23:**
BlobTools analysis of assemblies produced from long and short reads that did not map to the Bcop_v1 (Canu) Sciara genome assembly
Page 41

### Section 2: Supplemental Tables
#### Pages 42-79

**Supplemental Table S1 A-E:**
Expected genome and chromosome sizes
Page 42-43

**Supplemental Table S2:**
RNA-seq samples spanning both sexes and 4 life cycle stages
Page 44

**Supplemental Table S3:**
Short read assembly size statistics
Page 45

**Supplemental Table S4:**
Long read assembly size statistics
Page 46

**Supplemental Table S5:**
Pairwise comparisons of size statistics of hybrid scaffolds from pair of Canu or Falcon assemblies
Page 47

**Supplemental Table S6:**
Size statistics of Canu C3.2 across the work flow
Page 48

**Supplemental Table S7:**
Size statistics of Falcon F9 across the work flow
Page 49

**Supplementary Table 8 A-C:**
Gap size statistics
Page 50

**Supplemental Table S9:**
Bacterial contig statistics in each assembly
Page 51

**Supplemental Table S10:**
*Sciara* (*Bradysia*) *coprophila* repeat family classes from RepeatModeler
Page 52

**Supplemental Table S11:**
Sub-classification of classified Repeat Families found in *Sciara coprophila* genome with RepeatModeler
Page 53-54

**Supplemental Table S12 A-B:**
Repeat Masking on Canu
Page 55

**Supplemental Table S13 A-B:**
Repeat Masking on Falcon
Page 56

**Supplemental Table S14:**
Transcriptome Evaluations
Page 57

**Supplemental Table S15 A-B:**
Maker Annotation Transcript Evaluations on Canu and Falcon
Page 58-60

**Supplemental Table S16:**
Additional characterization and comparisons of the final annotations of Canu and Falcon assemblies
Page 61

**Supplemental Table S17 A-C:**
Putative Sciara homologs for proteins involved in reading, writing, and erasing DNA methylation marks for adenine and cytosine
Page 62-66

**Supplemental Table S18 A-F:**
DNA modification percentages in male embryonic genomic DNA
Page 67-69

**Supplemental Table S19:**
Which dimers are observed with modifications more often than expected?
Page 70

**Supplemental Table S20:**
Which trimers are observed with modifications more often than expected?
Page 71

**Supplemental Table S21 A-F:**
Binomial tests for enrichment or depletion of DNA modifications in various genomic features
Page 72-76

**Supplemental Table 22:**
L candidate contigs and corresponding proteins
Page 77

**Supplemental Table 23**
Highlighted proteins that aligned to putative Sciara contigs produced by reads that did not map to the Bcop_v1 (Canu) Sciara genome assembly
Page 78-79

### Section 3: Detailed experimental methods
### Pages 80-85

**3.1 Embryo Collection**
Page 80

**3.2 DNA extraction**
Page 80

**3.3 Illumina PE Genomic DNA library**
Page 80

**3.4 PacBio Sequencing Details**
Page 80-81

**3.5 MinION sequencing details**
Page 82-84

**3.6 BioNano Irys optical mapping details**
Page 84

**3.7 Strand-specific RNA-seq details**
Page 84-85

### Section 4: Detailed Bioinformatic Methods
### Pages 86-165

#### 4.1 Illumina assemblies
Page 86-100

##### 4.1.1 Inputs
Page 86-88

4.1.1.1 Trimming and quality filtering
4.1.1.2 Short read error-correction
4.1.1.3 Naming of short read datasets used in assemblies
4.1.1.4 Insert size statistics

##### 4.1.2 Assembling the short reads
Page 88-92

4.1.2.1 Abyss
4.1.2.2 Megahit
4.1.2.3 Platanus
4.1.2.4 SGA
4.1.2.5 SOAPdenovo2
4.1.2.6 SPAdes
4.1.2.7 Velvet

##### 4.1.3 Evaluations of the Initial 40 Short Read Assemblies
Page 93-95

4.1.3.1 Size statistics
4.1.3.2 Mapping the reads back to each assembly
4.1.3.3 Percent mapped
4.1.3.4 ALE
4.1.3.5 FRC$^{bam}$
4.1.3.6 LAP
4.1.3.7 REAPR
4.1.3.8 BUSCOv1

##### 4.1.4 Contamination analyses for the selected Platanus short read assembly
Page 96-100

4.1.4.1 Obtaining taxonomy IDs from BLAST hits for each contig.
4.1.4.2 Using BlobTools and custom scripts to label each contig with a taxonomy
4.1.4.3 Obtaining contamination-filtered reads for re-assembly
4.1.4.4 Platanus contamination-filtered re-assemblies
4.1.4.5 Evaluations of contamination-filtered re-assemblies

#### 4.2 Long Read Assemblies using PacBio and Oxford Nanopore Data
Page 101-131

**4.2.1 Long read inputs to assemblers**
Page 102-106

4.2.1.1 Long read set naming
4.2.1.2 Obtaining fastx files from PacBio .bax.h5 files using bash5tools
4.2.1.3 Obtaining fastx files from MinION fast5 files using fast5tools
4.2.1.4 A more detailed description of Fast5Tools and MinION read analyses
4.2.1.5 MarginStats for percent identity of MinION Reads

**4.2.2 Hybrid-assembling long reads with short reads**
Page 107

4.2.2.1 DBG2OLC with Platanus short read contigs and PBDAGCON

**4.2.3 Assembling the long reads alone**
Page 107-114

4.2.3.1 ABruijn
4.2.3.2 Canu
4.2.3.3 Falcon
4.2.3.4 Miniasm and RaCon
4.2.3.5 SMARTdenovo

**4.2.4 Polishing the long-read assemblies**
Page 115-116

4.2.4.1 Iterative polishing with signal-level PacBio data using Quiver
4.2.4.2 Evaluations during Quiver Polishing
4.2.4.3 Iterative polishing with Illumina short reads using Pilon

**4.2.5 Evaluations of the Initial 50 long-read assemblies**
Page 117-121

4.2.5.1 Size statistics
4.2.5.2 BUSCO completeness metrics
4.2.5.3 Illumina metrics
4.2.5.4 PacBio and MinION long read metrics
4.2.5.5 BioNano optical map metrics

4.2.5.6 Selection for BioNano scaffolding
4.2.5.7 Automating the battery of metrics

**4.2.6 BioNano scaffolding**
Page 121-122

4.2.6.1 BioNano CMAP assembly
4.2.6.2 BioNano hybrid scaffolding
4.2.6.3 Obtaining gap intervals for gap sizing
4.2.6.4 Scaffold comparisons and evaluation

**4.2.7 Assembly refining: Gap-filling, polishing, contamination removal, haplotig identification, and anchoring**
Page 122-131

4.2.7.1 Final polishing, gap-filling, and meta-scaffolding
4.2.7.2 Identifying contaminating contigs and scaffolds
4.2.7.3 Identifying primary and associated sequences
4.2.7.4 Anchoring sequences into chromosomes using known, localized, and/or homologous sequences
4.2.7.5 Classifying/anchoring assembled sequences as autosomal or X based on diploid vs haploid coverage levels

#### 4.3 Gene and repeat annotation
Page 132-153

**4.3.1 Transcriptome Assemblies**
Page 132-133

4.3.1.1 De novo Transcriptome assembly using Trinity
4.3.1.2 Genome-guided transcriptome assembly using Stringtie

**4.3.2 Evaluations of transcriptome assemblies**
Page 133-134

4.3.2.1 BUSCO
4.3.2.2 RSEM-Eval
4.3.2.3 TransRate

**4.3.3 Building a comprehensive repeat library for use in Maker2**
Page 135-136

4.3.3.1 *De novo* repeat libraries with RepeatModeler
4.3.3.2 Known *Sciara* repeats
4.3.3.3 Known Arthropod Repeats

**4.3.4 Maker2 gene annotations**
Page 137-152

4.3.4.1 Overview
4.3.4.2 Training GeneMark-ES for the first gene prediction round of Maker
4.3.4.3 Using BUSCO to train Augustus for the first gene prediction round of Maker
4.3.4.4 Using Maker est2genome/protein2genome to train SNAP for the first gene prediction round of Maker
4.3.4.5 Repeat Libraries
4.3.4.6 EST evidence
4.3.4.7 Alternative EST evidence
4.3.4.8 Protein homology evidence
4.3.4.9 Other parameter choices
4.3.4.10 Maker2 Round 1
4.3.4.11 Maker2 Round 2
4.3.4.12 Maker2 Round 3 (keep_preds=1)
4.3.4.13 Maker2 Round 3 with gene-filtered repeat library (keep_preds=1)
4.3.4.14 Maker2 Round 3 combined standard gene sets and finalization
4.3.4.15 Maker2 gene set evaluations

**4.3.5 Dosage compensation analysis**
Page 153

**4.3.6 Characterization of the Lambda Phage Insert Containing ScRTE**
Page 153

#### 4.4 DNA modifications
Page 154-165

##### 4.4.1 PacBio Analysis
Page 154-161

4.4.1.1 PacBio Modification Prediction
4.4.1.2 DNA sequence motifs found near sites of predicted DNA modifications
4.4.1.3 Kmers enriched for DNA modifications
4.4.1.4 MEME motifs in the highest scoring sites
4.4.1.5 Enrichment/Depletion of DNA modifications in various genomic regions
4.4.1.6 Binomial tests for higher or lower modification rates in genomic regions
4.4.1.7 Binomial tests for enrichment/depletion of DNA modifications in genomic regions
4.4.1.8 Spacing between DNA modifications

##### 4.4.2 Orthogonal support for PacBio modification results using the MinION data
Page 162-163

##### 4.4.3 Calculating observed:expected ratios of dimers and trimers
Page 164-165

#### 4.5 Candidate L sequences
Page 165

### Section 5: Software versions and Supplemental References
### Pages 166-175

#### 5.1 Software
Page 166-170

#### 5.2 Supplemental References
Page 171-175

### Glossary and Acronyms:

| | |
|---|---|
| **ALE** | Assembly Likelihood Evaluation. A probabilistic measure of assembly accuracy given read quality, mate pair orientation, insert length, coverage, alignments, and k-mer frequencies. See Clark et al 2013. |
| **Anchoring** | Anchoring is the process of mapping contigs/scaffolds into chromosomes, typically into specific loci. For flies like Drosophila and Sciara, this is done using data from in situ hybridization experiments on polytene chromosomes. |
| **BUSCO** | Benchmark Universal Single Copy Orthologs. Sets of genes expected to be present given evolutionarily related species. See Simao et al 2015. |
| **CMAP** | BioNano **C**onsensus **MAP**s produced by "assembling" raw BioNano optical maps together to produce a consensus optical map representation of the genome.See Lam et al 2012, and other BioNano literature. |
| **Contig** | Contiguous sequence put together by aligning reads together. The ends of contigs represent areas of uncertainty where multiple choices were present, commonly caused by repeats. |
| **Coverage** | This is the sum of read lengths divided by the genome size – i.e. the average number of times any given base in the genome is read. Thus, 10X coverage means that the sum of read lengths was 10 times the genome length, and each base is represented 10 times on average in the dataset. |
| **EGS** | Expected Genome Size – the expected size or length of a haploid (single copy) complement of a genome. The EGS for Sciara was obtained with DNA content measurements by Rasch. |
| **FRC / $FRC^{bam}$** | Feature Response Curve. $FRC^{bam}$ is a program that evaluates assemblies using read-layouts of aligned reads. We used the number of features it flags as a proxy for the number of assembly errors. See Vezzi et al 2012. |
| **Haplotigs** | Diploid genomes can have loci that are represented twice (or more) in an assembly. Each contig representation of the same locus is a haplotig. Common practice is to use only the version on the longest contig as part of the primary assembly. See Roach et al 2018 for more information. |
| **LAP** | Log Average Probability. LAP is a probabilistic measure of assembly quality given a set of reads. The more an assembly resembles the genome sequence, the higher the score. |
| **L50 / LG50** | Ordering contigs from longest to shortest, the number of the longest contigs needed to reach or exceed 50% of the assembly size (L50) or expected genome size (LG50). LG50 is used to directly compare different assemblies. |
| **LINEs** | Long interspersed nuclear elements, a family of transposons. |
| **LTR** | Long Terminal Repeat – LTR transposons are a family of transposons. |
| **N50 / NG50** | The length of the shortest contig (or scaffold or read) such that 50% of the assembly or dataset size (N50) or 50% of the expected genome length (NG50) is on sequences of that length or longer. NG50 is used to directly compare different assemblies. |
| **RC** | Rolling Circle family of transposons. |
| **SINEs** | Short interspersed nuclear elements, a family of transposons. |
| **Scaffolds** | A set of contigs ordered and oriented with respect to each other with unknown sequence in between them (gaps). Gaps are usually represented as a stretch of Ns given the estimated gap length. |
| **Sniffles** | A program to detect structural variants (SVs) with long reads. See Sedlazeck et al 2018. |

| | |
|---|---|
| **Span** | The length of sequence in an assembly covered at least once by reads (or optical maps). Thus, if optical map alignments span 50 Mb of an assembly, then 50 Mb had at least 1 optical map over it. |
| **SV** | Structural variant – a variation between two genomes involving the structure of the genome rather than a point mutation. Examples are short and long deletions, copy number variations, duplications, insertions, and translocations. Mis-assemblies will increase the number of apparent SVs. So, assemblies that look more like the true underlying genome structure will have fewer SVs than assemblies with many errors. |

### Section 1: Supplemental Figures

#### Supplemental Figure S1: Comparing evaluations of short read assemblies to long read assemblies.

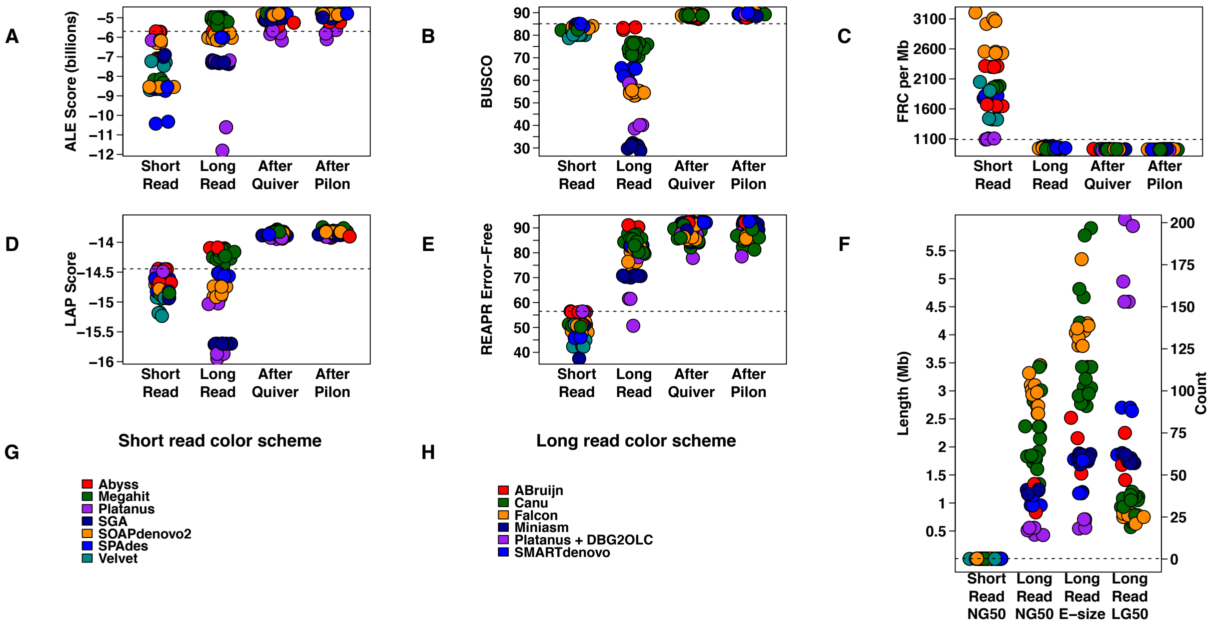

**Supplemental Figure S1: Comparing evaluations of short read assemblies to long read assemblies (prior to any scaffolding).**

**(A)** ALE scores. **(B)** BUSCO: percent complete Arthropod BUSCOs. **(C)** Average number of features per Mb detected by FRC$^{bam}$. **(D)** LAP scores**. (E)** REAPR percent error-free bases. **(F)** Size statistics. NG50 = size of contig such that at least 50% of the expected genome size is on contigs of that size or larger. Expected contig size is as defined in the Genome Assembly Gold-standard Evaluations (GAGE) paper (Salzberg et al. 2012), which is designed to give the expected size of contig containing a randomly selected base in the assembly. The expected genome size was used in the denominator of the equation instead of individual assembly sizes for direct comparisons as done in the GAGE paper. LG50 is the number of contigs (count) it takes to reach at least 50% of the expected genome size when selecting contigs from longest to shortest. Lower LG50s are better. **(G-H)** Legends for A-F. “Short read” indicates assemblies generated using Illumina data. “Long read” indicates assemblies generated using long read data before any polishing. “After Quiver” and “After Pilon” indicate the long read assembly evaluations taken after each of those polishing steps. In (F), the size statistics for long read assemblies are from those after Pilon polishing.

#### Supplemental Figure S2: Assembly ranking correlation matrices

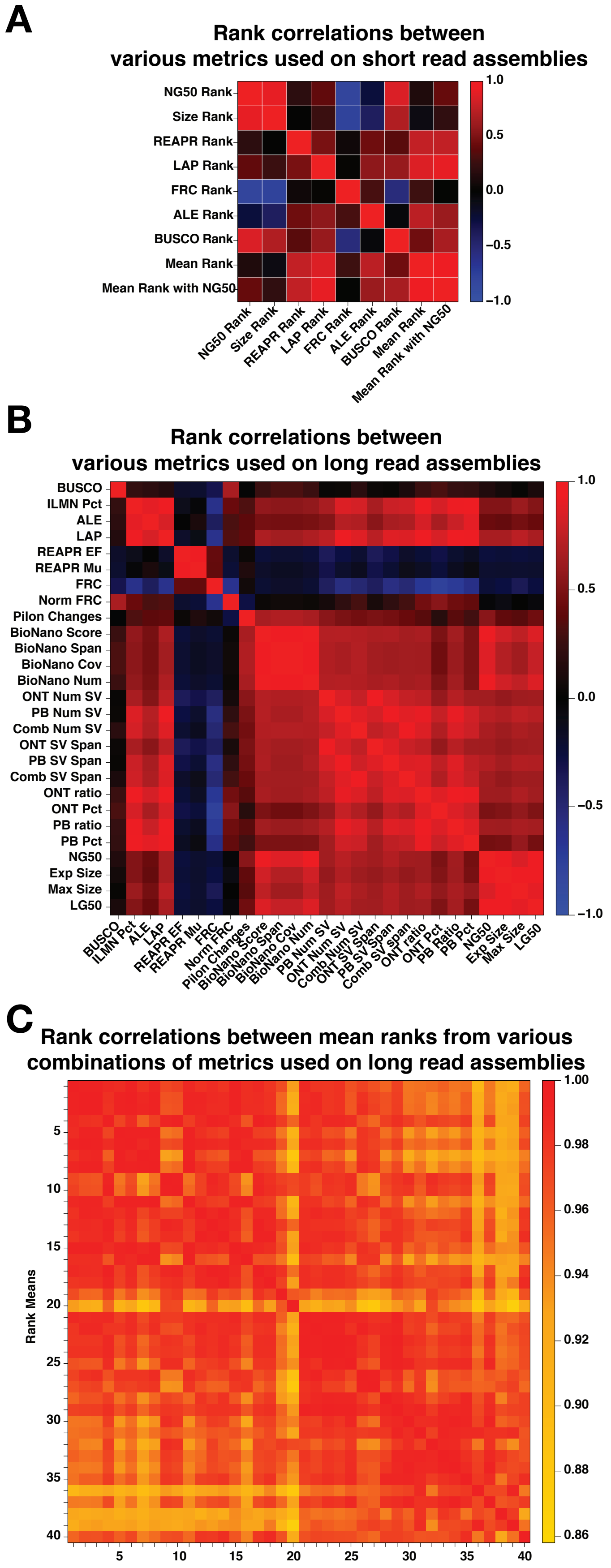

**Supplemental Figure S2: Assembly ranking correlation matrices**

**(A)** Matrix of pairwise correlations between short read assembly rankings for given metrics. Also see Figure 2A.

**(B)** Matrix of pairwise correlations between long read assembly rankings for given metrics. Also see Figure 2E.

**(C)** Matrix of pairwise correlations between long read assembly rankings for mean ranks from different combinations of metrics. Also see Figure 2E.

Note: These matrices represent both Pearson and Spearman correlation coefficients since the correlations were computed on ranks.

#### Supplemental Figure S3: Filtering out non-Arthropod, contaminating reads using Taxonomy- annotated GC plots.

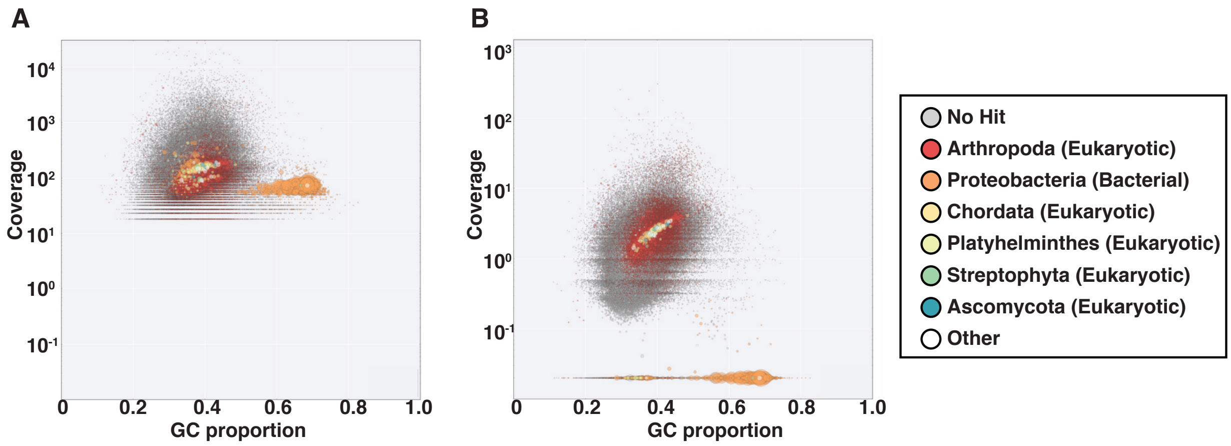

**Supplemental Figure S3: Filtering out non-Arthropod, contaminating reads using Taxonomy- annotated GC plots.**
Taxonomy-Annotated GC (TAGC) plots can be used to visualize the GC proportion and coverage for each contig in an assembly. In the TAGC plots, each circle represents a contig. The size of each circle is proportional to contig size. The colors correspond to phylum-level taxonomy assignments. Notice that the largest contigs in this Platanus assembly are all bacterial (i.e. the largest circles are orange). Contaminating genomes, from the microbiome on and in embryos for example, are expected to be at different copy numbers and have different GC contents than the target organism. Therefore, when contaminating genomes are present, more than one cluster of contigs is typically seen, allowing coverage and GC proportion cutoffs to be chosen for filtering unannotated contigs.
**(A)** TAGC plot of the chosen Platanus assembly with kmer coverage from the reads input into the assembly on the y-axis and GC proportion of contigs on the x-axis. The bacterial cluster has similar coverage and though it has a higher average GC content, the clusters overlap in that dimension as well.
**(B)** TAGC plot with read coverage from pre-amplification stage salivary glands on the y-axis and GC proportion of contigs on x-axis. In this case, using a different sample from a different tissue and prepared by a different person, resulted in differential coverage between the *Sciara* genome and the bacterial cluster.

**Supplemental Figure S4: Length Distributions for Illumina Scaffolds, PacBio Reads and MinION Molecules.**

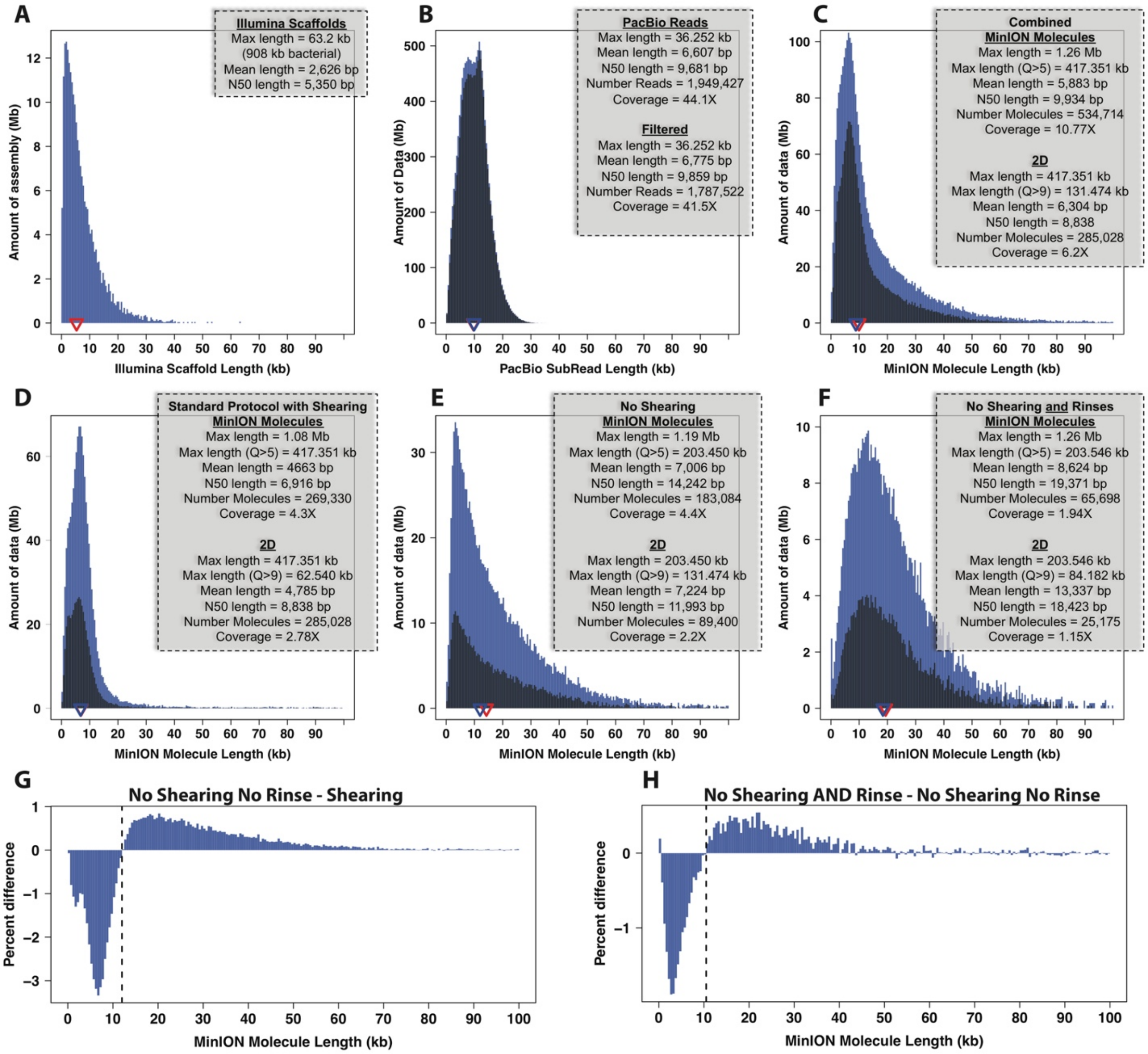

**Supplemental Figure S4: Length Distributions for Illumina Scaffolds, PacBio Reads and MinION Molecules.**

**(A)** Platanus scaffold lengths from Illumina data. **(B)** Sub-read length distribution for all PacBio subreads (blue) or filtered (shaded area). **(C-F)** MinION molecule length (blue) and 2D read only (shaded) distributions from **(C)** combining all libraries, **(D)** only libraries that followed the standard ONT protocol with shearing, targeting 8 kb, **(E)** libraries that skipped shearing and used other long read principles, but did not include rinse steps, and **(F)** libraries that skipped shearing AND included rinse steps. **(G)** Libraries that skipped shearing are depleted for molecules around 10-12 kb and enriched for longer molecules with respect to the libraries from the standard protocol. **(H)** Libraries that skipped shearing AND included rinse steps are additionally depleted for molecules <10-12 kb and additionally enriched for longer molecules, as demonstrated by comparing to the libraries that only included shearing, but no rinse. We described the rinse steps during AMPure cleanups elsewhere (Urban et al. 2015) and in the supplemental methods. Note that a single “MinION molecule length” is defined for each sequenced molecule (that can produce up to 3 reads) as described in the supplemental methods and elsewhere (Urban et al. 2015).

**Supplemental Figure S5: Percent identity of MinION reads compared to a PacBio-only assembly.**

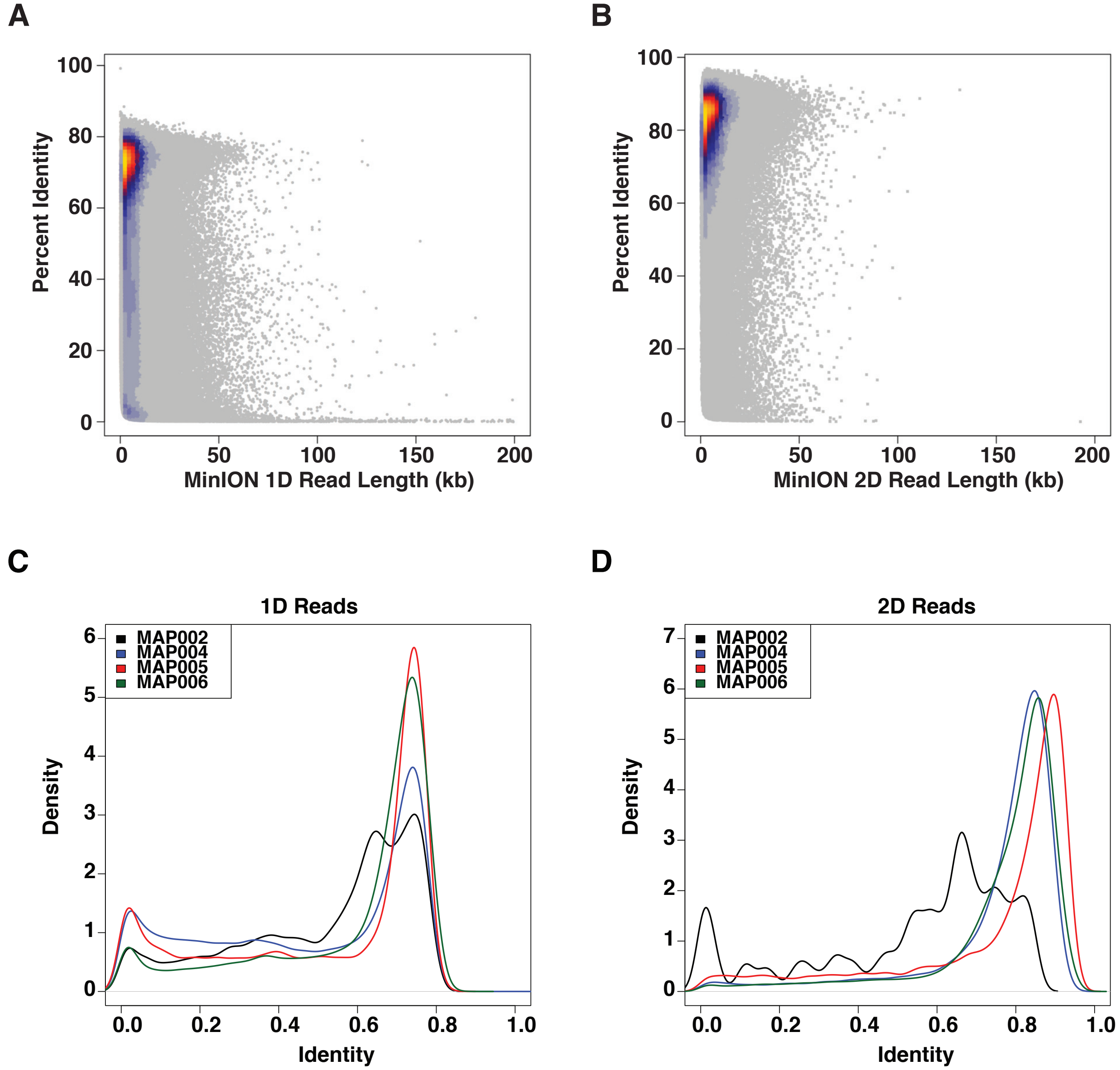

**Supplemental Figure S5: Percent identity of MinION reads compared to a PacBio-only assembly.**
**(A)** 1D read length vs percent identity. **(B)** 2D read length vs percent identity. **(C-D)** Percent identity distributions separated by kit version for **(C)** 1D reads and **(D)** 2D reads. In both (A) and (B), the scatter plot has heat colors representing where the least (grey) and most (yellow) data is located. The limits on the x-axes was set to 200 kb to better visualize the data shorter than that length. The several reads longer than 200 kb appear to be quite low quality. In all, the percent identity is the result of summing the identities in the section of the read that aligned and dividing by the length of the read. Since we did not perform a re-alignment step (Jain et al. 2015), low quality reads with only small sections mapped appear to have a much lower percent identity than is likely the case.

#### Supplemental Figure S6: Evaluations across Quiver polishing rounds.

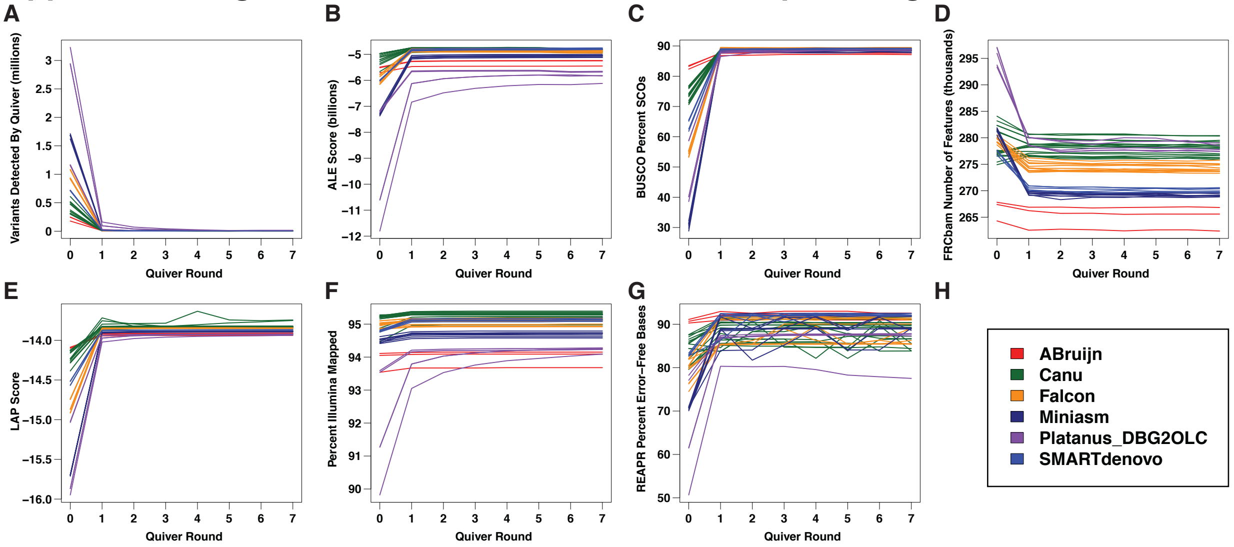

**Supplemental Figure S6: Evaluations across Quiver polishing rounds.**
**(A)** Number of variants detected by Quiver. **(B)** ALE scores. **(C)** BUSCO (Benchmark Universal Single Copy Ortholog): percent complete Single Copy Orthologs (SCOs). **(D)** Number of features detected by FRC$^{bam}$. **(E)** LAP scores. **(F)** Percent of Illumina dataset that maps to the assembly. **(G)** REAPR percent error-free bases. **(H)** The legend for all plots. Note that “round 0” represents the scores of the assemblies being input into the first Quiver round. The largest improvement in all metrics is seen after the first round (at Quiver Round = 1).

**Supplemental Figure S7: Blended assemblies with both PacBio and MinION data tended to receive better ranks than PacBio-alone assemblies.**

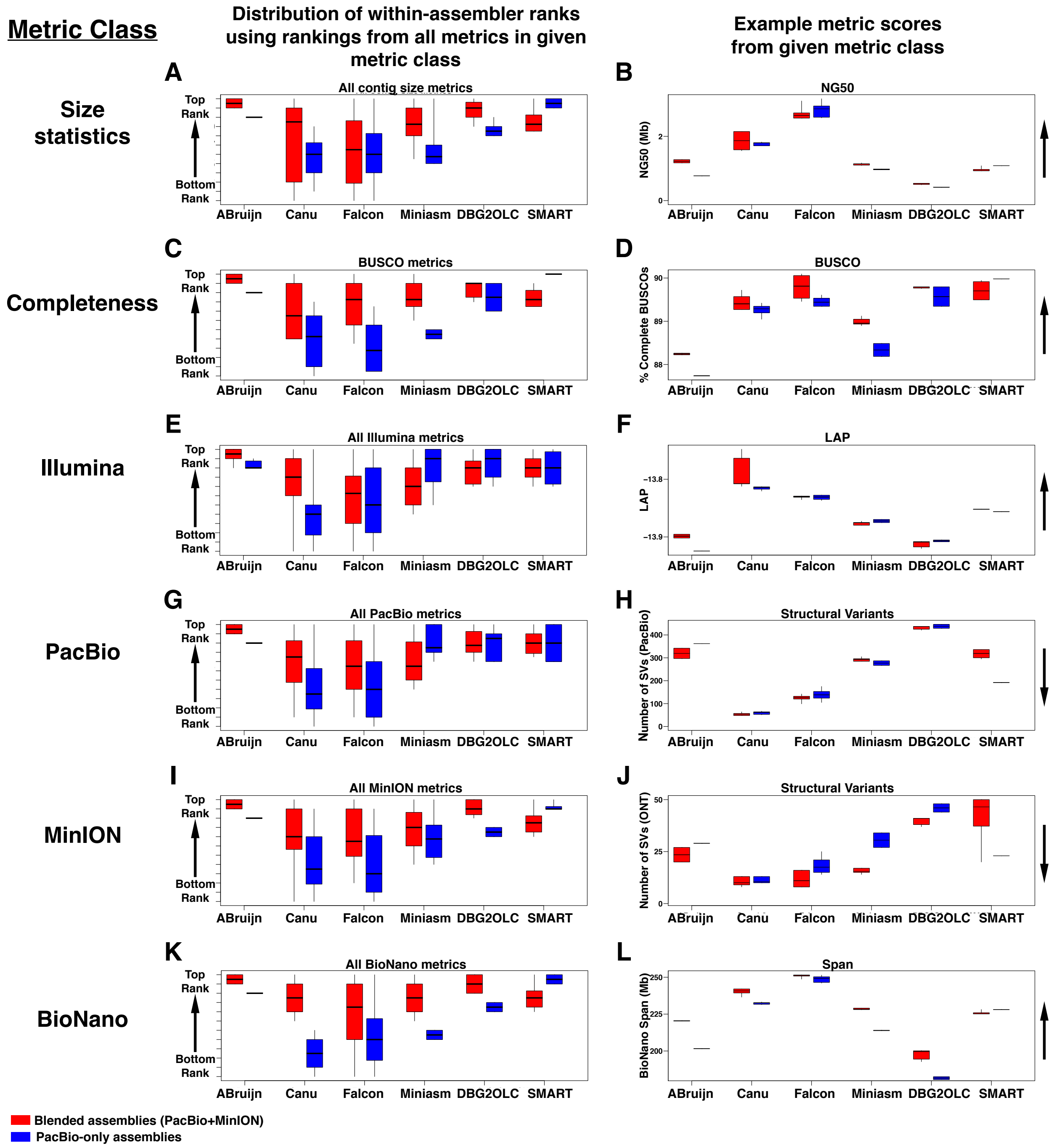

**Supplemental Figure S7: Blended assemblies with both PacBio and MinION data tended to receive better ranks than PacBio-alone assemblies.**
Box and whisker plots where the boxes show the 25th-75th percentile, the black line is the median, and the whiskers span the range (min to max). **(A, C, E, G, I, K)** Assemblies from a given assembler were ranked using all individual metrics from given metric class (to the left of plots). The ranks from individual metrics were combined and partitioned into those from blended versus PacBio-only assemblies. Better ranks are higher. **(B, D, F, H, J, L)** The actual scores (not ranks) from an example metric from the given metric class is plotted. The arrows to the right of plots show the direction that the given score gets better.

#### Supplemental Figure S8: Metrics comparing assemblies after scaffolding

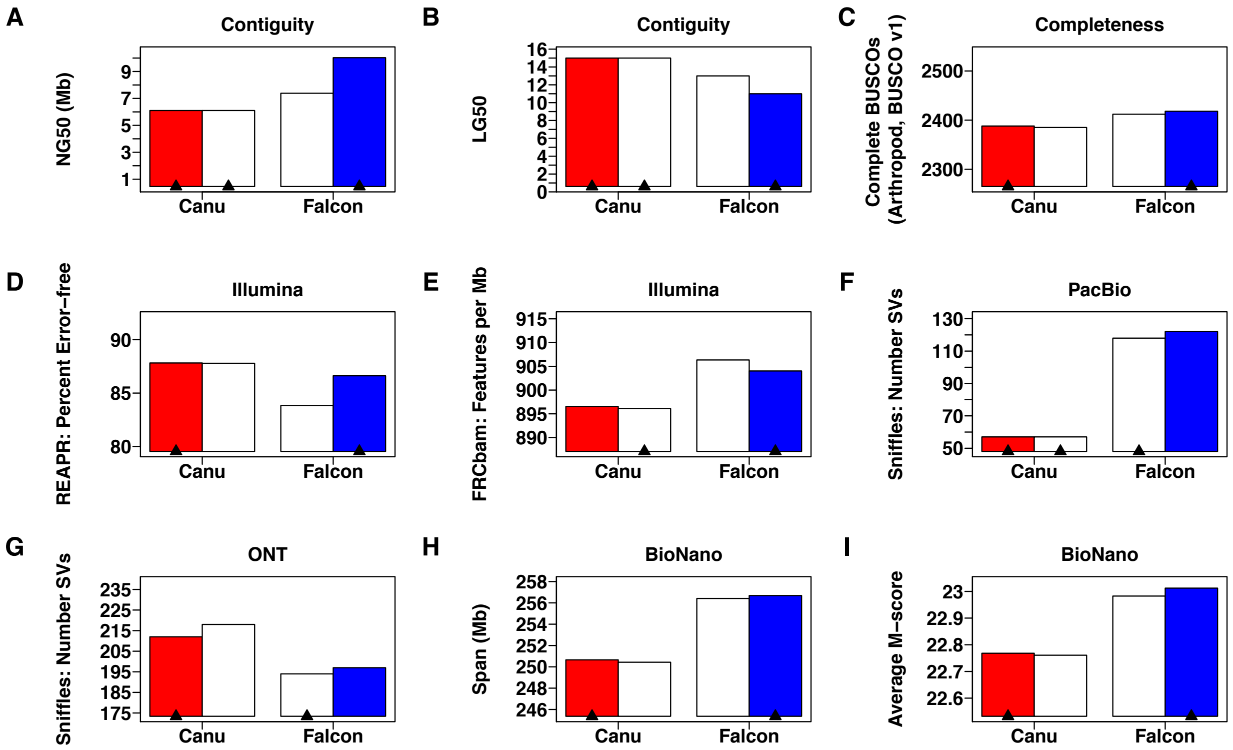

**Supplemental Figure S8: Metrics comparing assemblies after scaffolding**
We chose two Canu and two Falcon assemblies for BioNano scaffolding. However, we found them to be extremely similar, so used only the one version of each that tended to do slightly better in these metrics. Also see Supplemental Figure S9 and Supplemental Table S5. In each panel, the red bar is the chosen Canu assembly and the blue bar is the chosen Falcon assembly. The white bars are the assemblies that we did not continue working on. For each assembler, the black arrowheads indicate which assembly in the pair did better in that metric (both in a pair have arrowheads in the case of a tie). **(A-B)** Show contiguity metrics: NG50 and LG50. Higher NG50s and lower LG50s are considered better. **(C)** Shows BUSCO as a completeness metric where higher numbers are better. **(D-E)** Show Illumina-based metrics from REAPR and FRC$^{bam}$, where higher and lower numbers, respectively, are better. **(F-G)** Show long read metrics for PacBio and MinION (ONT) respectively. Both show the number of structural variants (SVs) as detected by Sniffles given those datasets. Lower numbers are better. **(H-I)** Show BioNano optical map metrics. Span is the number of bases in the assembly covered by at least one optical map (higher is better). M-score is a score based on the alignments from Maligner that is typically negative, but is multiplied by -1 here for positive values such that higher is better.

#### Supplemental Figure S9: Aligning chosen and discarded scaffolds from each assembler (Canu and Falcon)

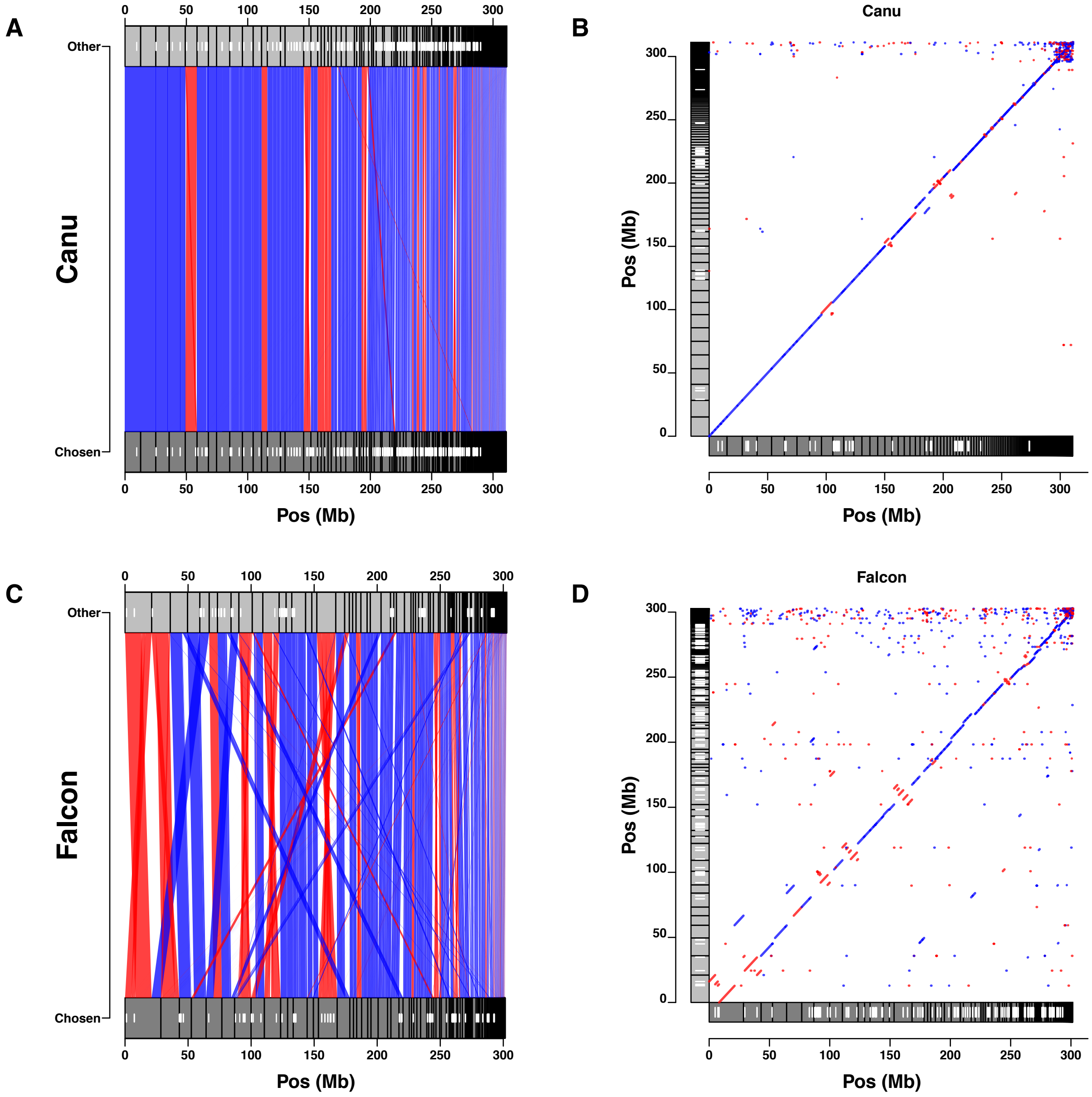

**Supplemental Figure S9: Aligning chosen and discarded scaffolds from each assembler (Canu and Falcon)**
We took two Canu and two Falcon assemblies into BioNano scaffolding. For a given assembler we aligned the two assembly versions with Minimap2 and visualized the alignments as pairwise plots (A, C) and dot plots (B, D) using our own tools made in R (github.com/JohnUrban/lave). In both types of plots, blue represents that the positive strand from both sequences aligned whereas red indicates that the alignment is in the reverse direction (negative strand with respect to each other). The polygons at the top and bottom of the pairwise plots or the x- and y- axes of the dot plots represent the scaffolds, and the white smaller polygons within them represent gap locations. In both, the chosen assembly is the bottom set of polygons (i.e. x-axis in dot plots). **(A-D)** Pairwise alignment plot and dot plot for the two sets of scaffolds derived from Canu assemblies (A-B) or Falcon assemblies (C-D). For more on analyzing the two versions of scaffolds from each assembler, see Supplemental Figure S8 and Supplemental Table S5.

In the figure labels, “Pos” = Position.

#### Supplemental Figure S10: BlobTools analysis of the Canu scaffolds

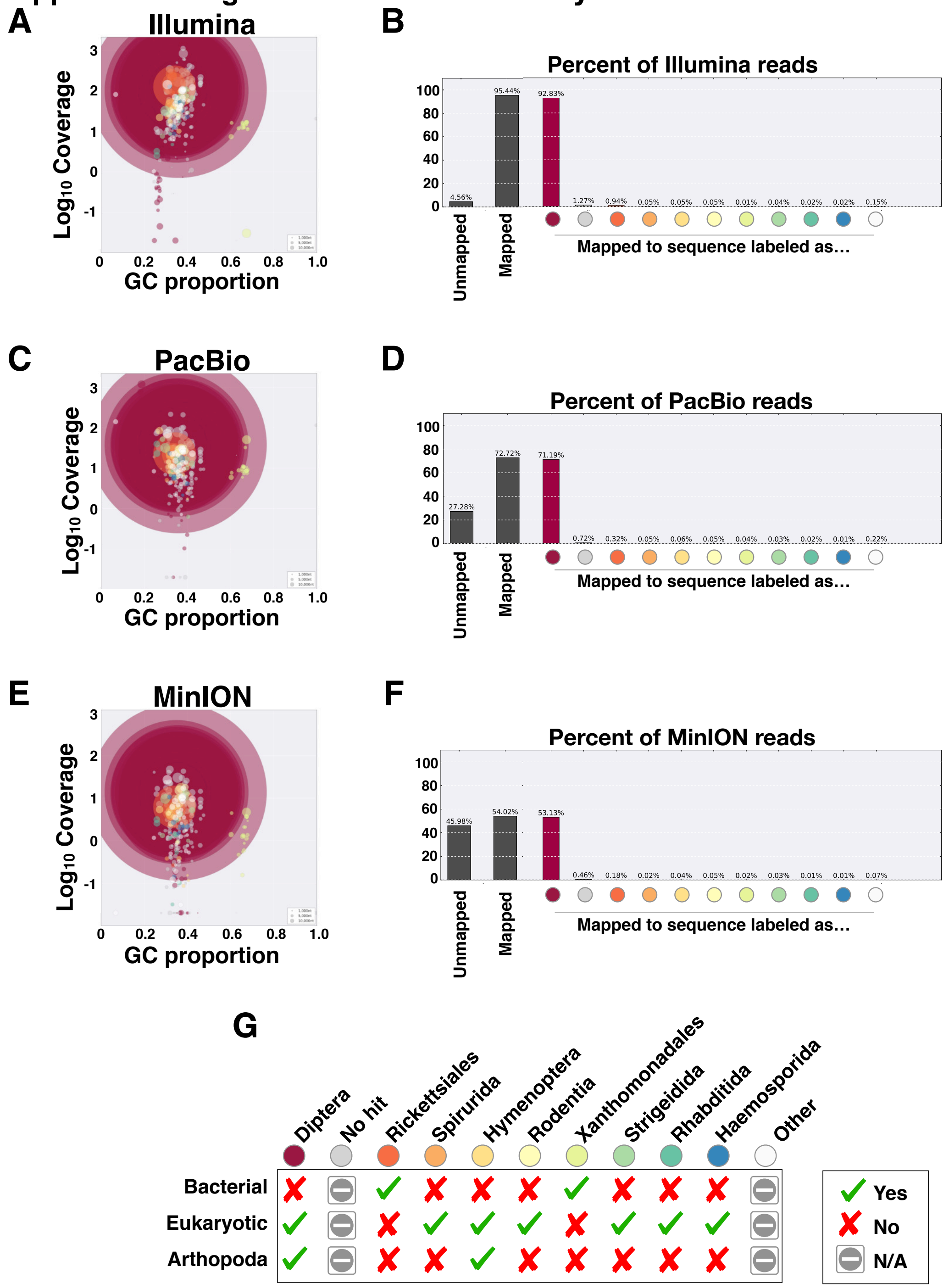

**Supplemental Figure S10: BlobTools analysis of the Canu scaffolds**
Note that essentially identical results were obtained for the Falcon assembly. **(A, C, E)** These plots show contigs/scaffolds as circles sized relative to the contig size, colored according to taxon information, and positioned according to GC content and read coverage from **(A)** Illumina, **(B)** PacBio, or **(C)** MinION. The coloring scheme is as detailed in (G). The largest contigs (and majority of assembled sequence) are labeled as Dipteran (Arthropoda). **(B, D, F)** Percent of reads that mapped (or did not) to any contig and percent that mapped to contigs labeled as the given taxonomy for the **(B)** Illumina, **(D)** PacBio, and **(F)** MinION datasets. Color scheme of circles on x-axis is as in (G). **(G)** Circle color legend for A-F with some additional taxon information.

#### Supplemental Figure S11: BlobTools analysis and anchoring Falcon scaffolds

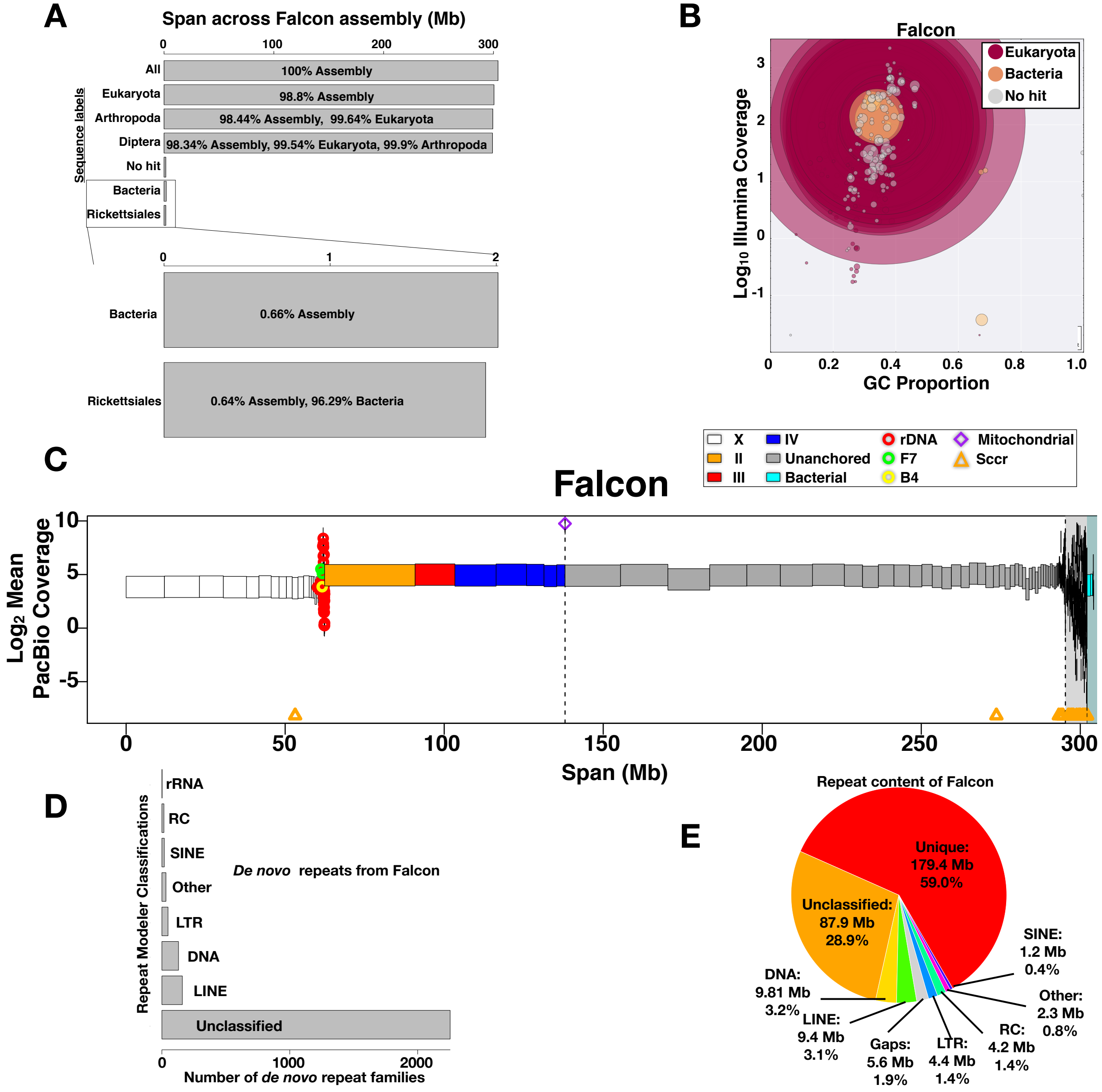

**Supplemental Figure S11: BlobTools analysis and anchoring Falcon scaffolds**

**(A)** The proportions of the Falcon assembly taxonomically labeled as Eukaryotic, Arthropoda, Diptera, Bacteria, and Rickettsiales.

**(B)** Log10 Illumina coverage versus GC content over the Falcon assembly.

**(C)** The Falcon assembly with scaffolds drawn as rectangles corresponding to their lengths, colored according to the chromosome they were anchored into (or unanchored), and positioned at their mean coverage from PacBio reads, the dataset used to determine X-linked sequences by haploid level coverage. The white background highlights sequences in the primary assembly whereas the grey and cyan backgrounds are set behind associated and bacterial sequences, respectively. All sequences to the left of the first vertical dashed line are anchored.

**(D)** Repeat modeler family classifications when run on Falcon assembly.

**(E)** Repeat content of the Falcon assembly from repeat masking. Note that the DNA, LINE, LTR, RC, and SINE are also classes of transposon elements ("DNA" stands for DNA elements).

#### Supplemental Figure S12: The single locus that contains the full-length Escribá insert

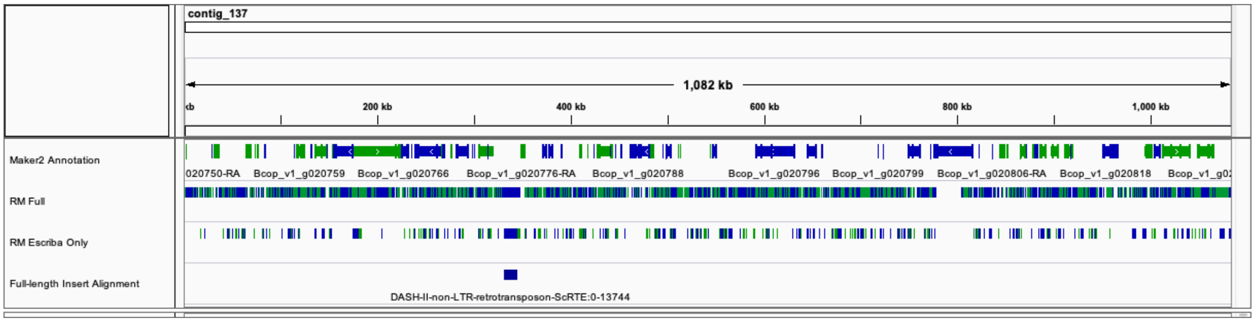

**Supplemental Figure S12: The single locus that contains the full-length Escribá insert**
This figure shows the ~1.1 Mb contig (contig_137) that harbors the full-length 13.8 kb insert sequence from Escribá et al (2011). In all rows, green shows positive strand alignments and blue shows negative strand alignments. The top row shows the Maker2 annotated regions. “RM Full” shows all regions marked as repeats by repeat masker when using the full comprehensive repeat library. “RM Escribá only” shows regions marked as repeats by repeat masker when using only the 13.8 kb Escribá insert sequence (Escribá et al. 2011). “Full-length Insert Alignment” shows the precise site that the insert corresponds to on this contig. The site is intergenic.

#### Supplemental Figure S13: Pairwise comparisons of final Canu and Falcon scaffolds

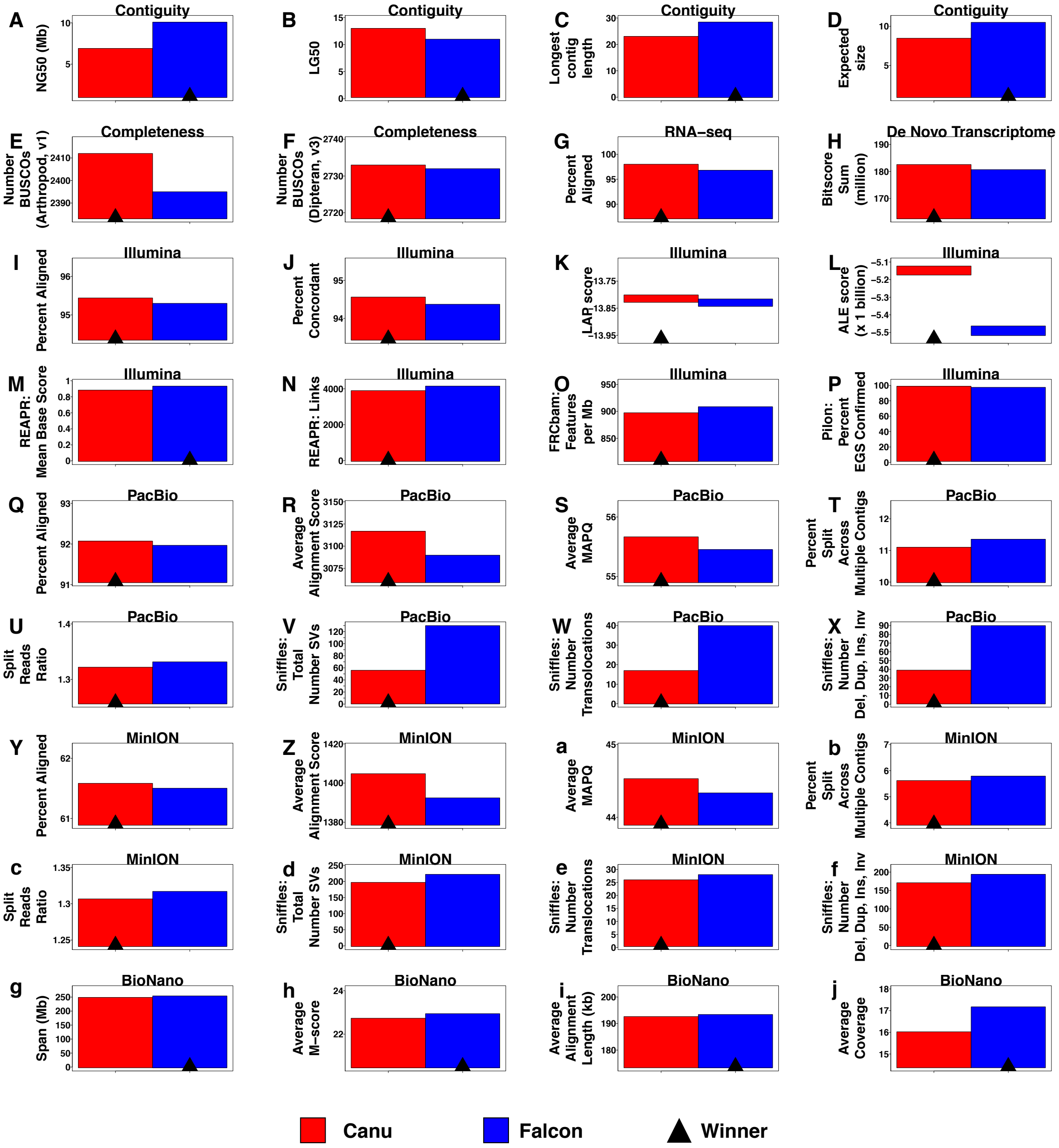

**Supplemental Figure S13: Pairwise comparisons of final Canu and Falcon scaffolds.** This figure shows actual scores, summarized in Figure 5F, for each of the evaluations of the assemblies for contiguity, completeness, and genome sequencing metrics. Evaluations of the transcriptomes and annotations guided by these assemblies are reported in Supplemental Tables S14, S15, and S16. Overall, the Canu scaffolds and annotation consistently received better scores.

Note that the black arrowhead shows the winner in each category, and that in some categories higher scores are better, and other categories lower scores are better. For example, a better assembly will have fewer putative mis-assembled regions as proxied by Sniffles structural variants, split long read alignments, and FRC[bam] features.

**Supplemental Figure S14: Pairwise comparisons of final Canu and Falcon annotations.**

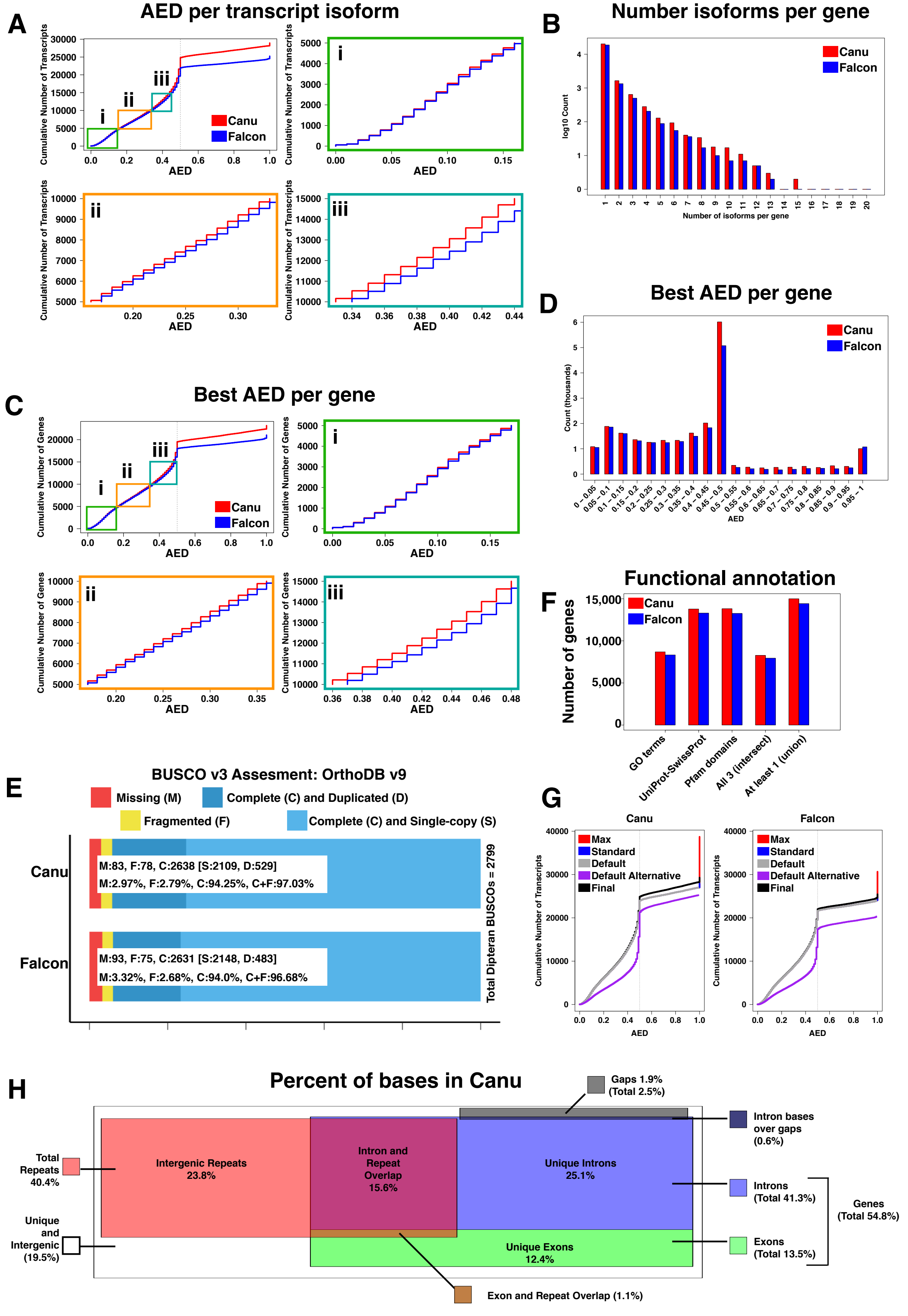

**Supplemental Figure S14: Pairwise comparisons of final Canu and Falcon annotations.**

**(A)** Shows the cumulative number of transcripts (y-axis) with respect to annotation edit distance (AED; x-axis). The (i) green, (ii) orange, and (iii) cyan boxes show subsequent subsets of 5000 transcripts up close. Canu had more transcripts with lower AEDs. **(B)** Canu typically has slightly more isoforms per gene than Falcon. **(C)** Shows the cumulative number of genes with respect to the best (lowest) AED used for each gene. Still the Canu annotation tends to have more genes with lower AEDs. **(D)** Shows the number of genes in each AED category. **(E)** Shows the BUSCO assessment results on the annotation transcript sets using all Dipteran BUSCOs (n=2799). Canu's annotation has a slightly higher % of BUSCOs found than Falcon's (97.03% vs 96.68%). **(F)** Visualization of the functional annotation results. The Canu annotation has more genes with GO terms, BLASTp hits to the entire UniProt-SwissProt protein dastabase, and/or with recognizable Pfam protein domains than the Falcon annotation. **(G)** Shows the cumulative number of transcripts with respect to AED for the different subsets (default, standard, and max) in the annotation that used the original repeat library. It also shows the default transcript set (purple) for the annotation that used the alternative/filtered repeat library for masking. Notice that the AED distribution is worse for that annotation. Therefore, only genes that had no overlapping representation in the original annotation were taken from that annotation (and they tended to be repetitive genes such as histones and olfactory receptors). See supplemental methods for more information. **(H)** A Venn-like diagram of boxes representing percentages of the Canu assembly annotated as genes (exons, introns), repeats, gaps, and overlaps amongst features.

**Supplemental Figure S15: Dosage compensation of X-linked genes in *Sciara coprophila***

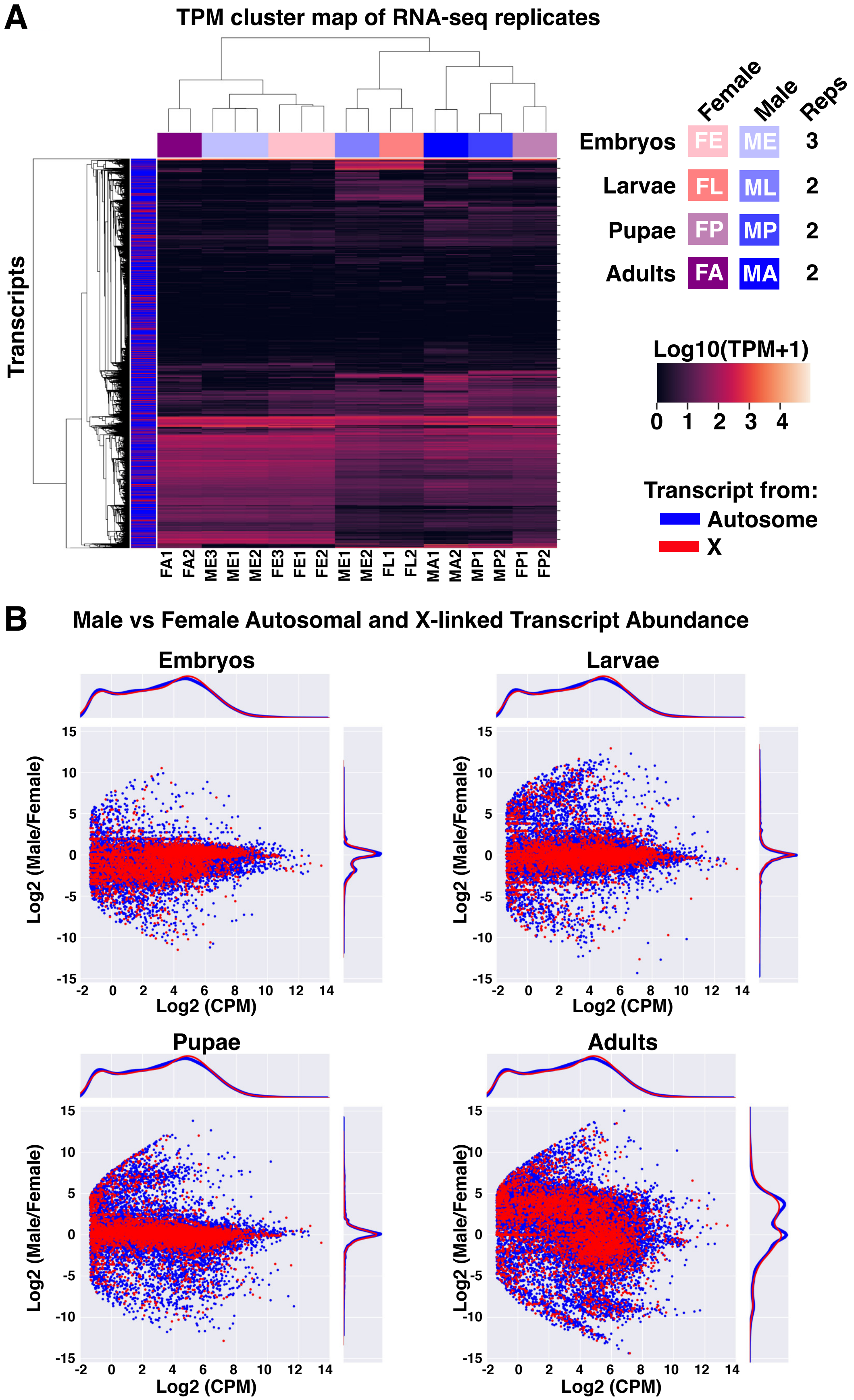

**Supplemental Figure S15: Dosage compensation of X-linked genes in *Sciara coprophila***

**(A)** Clustered heatmap of transcript abundances represented as log10 Transcripts Per Million (TPM) demonstrates that the replicates were highly reproducible, that autosomal and X-linked genes span all expression levels in both sexes, and that transcript abundances of X-linked genes appear similar across sexes in the same stage.

**(B)** Scatter plots of the Log2 average Count Per Million vs the Log2 ratio of male to female transcript abundances. Dots are colored blue for autosomal genes and red for X-linked genes as in (A).

#### Supplemental Figure S16: Distribution of DNA modifications across *Sciara* genome (PacBio analysis)

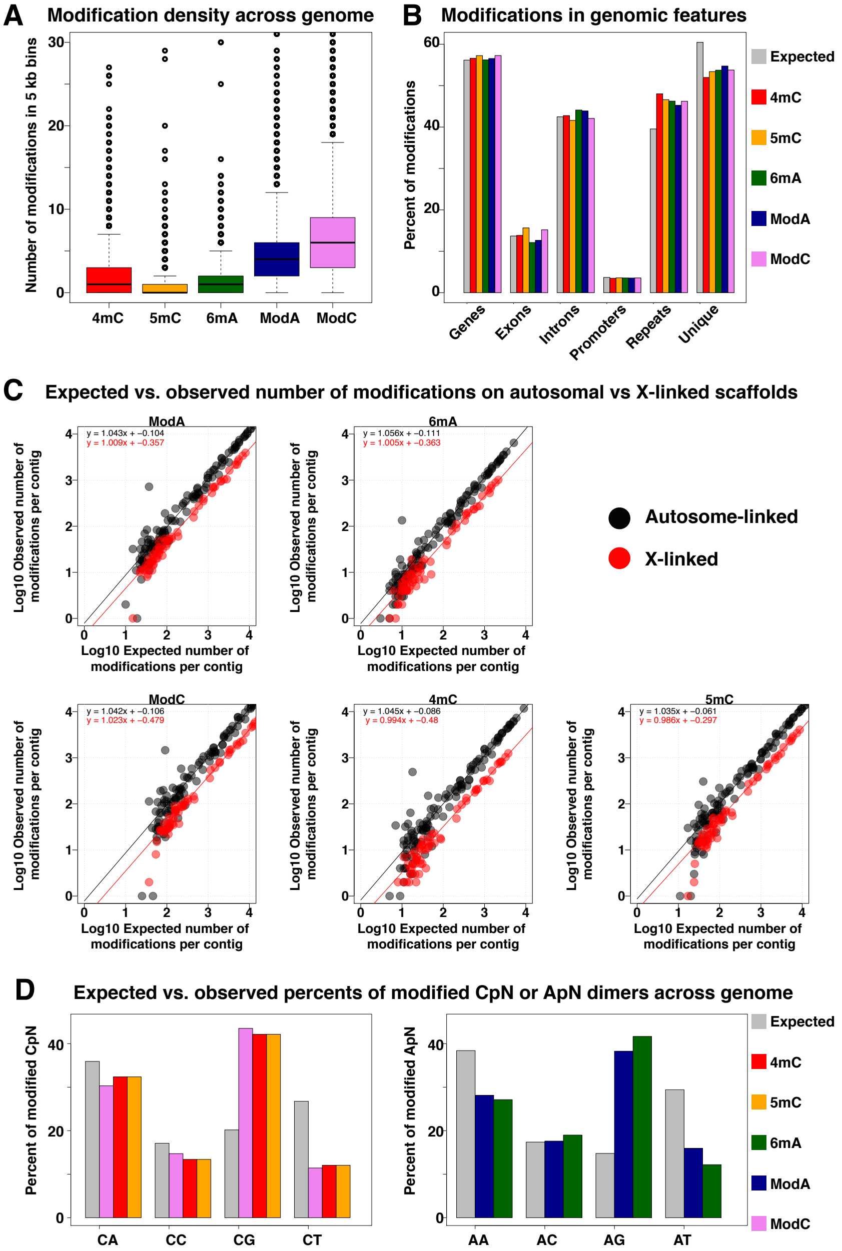

**Supplemental Figure S16: Distribution of DNA modifications across *Sciara* genome (PacBio analysis)**

**(A)** Boxplots of the number of modifications in 5 kb bins. Note that ModA and ModC are all adenines and cytosines, respectively, flagged as putative modified bases and 6mA or 4mC/5mC are subsets therein. Boxes span from the 25$^{th}$-75$^{th}$ percentiles with the median denoted as a black line. Whiskers extend up/down from the box to datapoints no more than an additional 1.5x the interquartile range (IQR). Outliers are shown as circles. **(B)** Expected vs Observed proportion of DNA modifications in genomic features. See supplemental tables S20 A-D as well. **(C)** Expected vs observed counts on contigs/scaffolds in the primary assembly labeled as Arthropoda (phylum). X-linked sequences have fewer modifications than autosomal sequences, but this is likely due to the loss of sensitivity from having half the strand coverage. **(D)** For given C or A modification type, this shows the percent of modified CpN or modified ApN that each dimer makes up for that category. The expected percents are obtained from the percent of all CpN or ApN genome-wide. Also see Supplemental Table S18.

**Supplemental Figure S17: Position weighted 7-mer motifs learned from different filtering and different subsets of the genome sequence (PacBio analysis)**

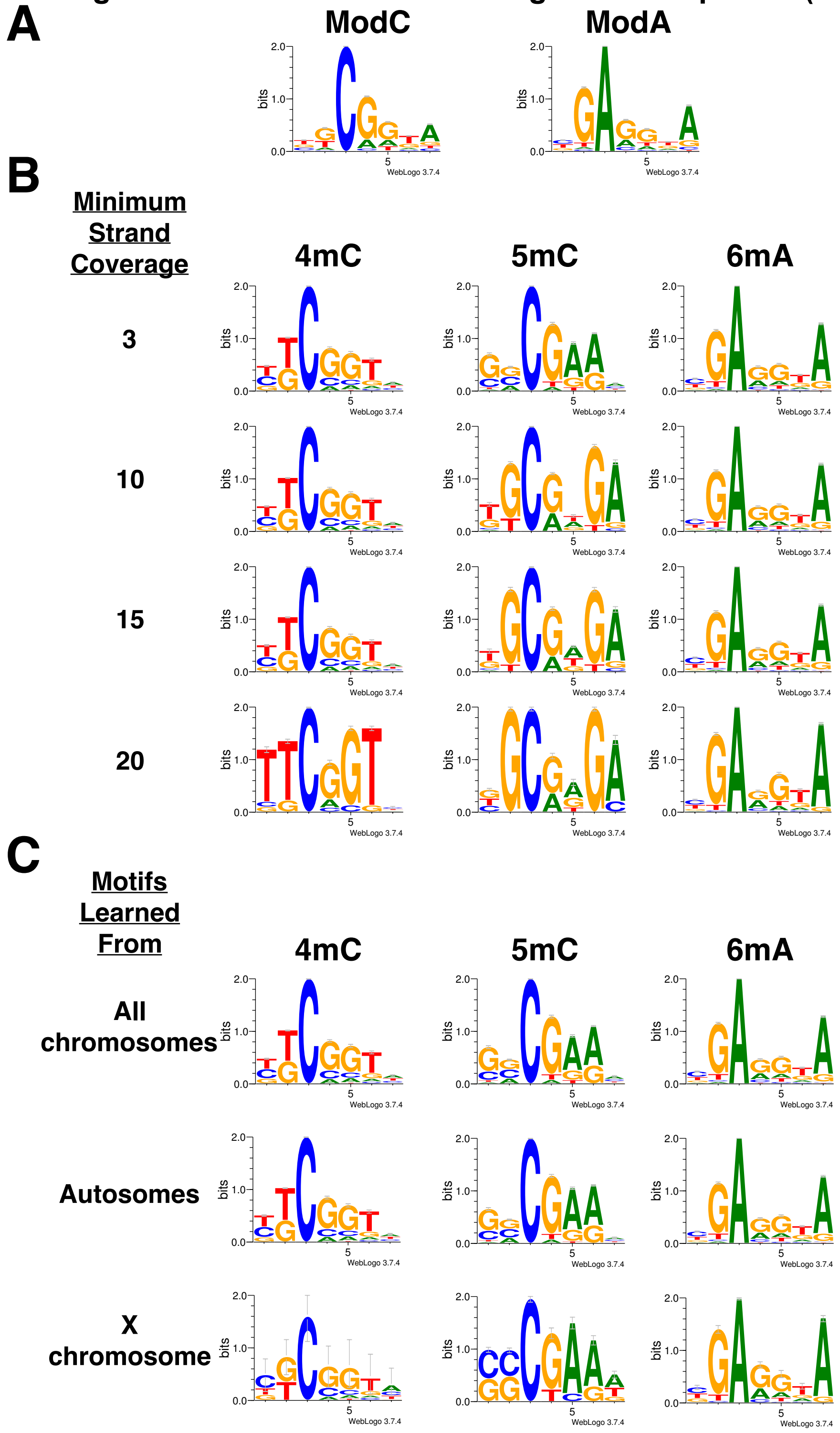

**Supplemental Figure S17: Position weighted 7-mer motifs learned from different filtering and different subsets of the genome sequence (PacBio analysis)**
**(A-C)** contain position weighted motifs from the sets of 7-mers (where the modified base occurs at position 3) enriched for 6mA, 4mC, or 5mC in the PacBio analysis. **(A)** Motifs from enriched 7-mers found in the analysis of all adenine or cytosine sites found as putatively modified. **(B)** Motifs learned from putative 4mC, 5mC, and 6mA sites with increasingly stringent filtering. **(C)** Either all sites were considered or the sites were partitioned into those on contigs we classified as chromosome X or autosomal. The motifs are the same in all cases.

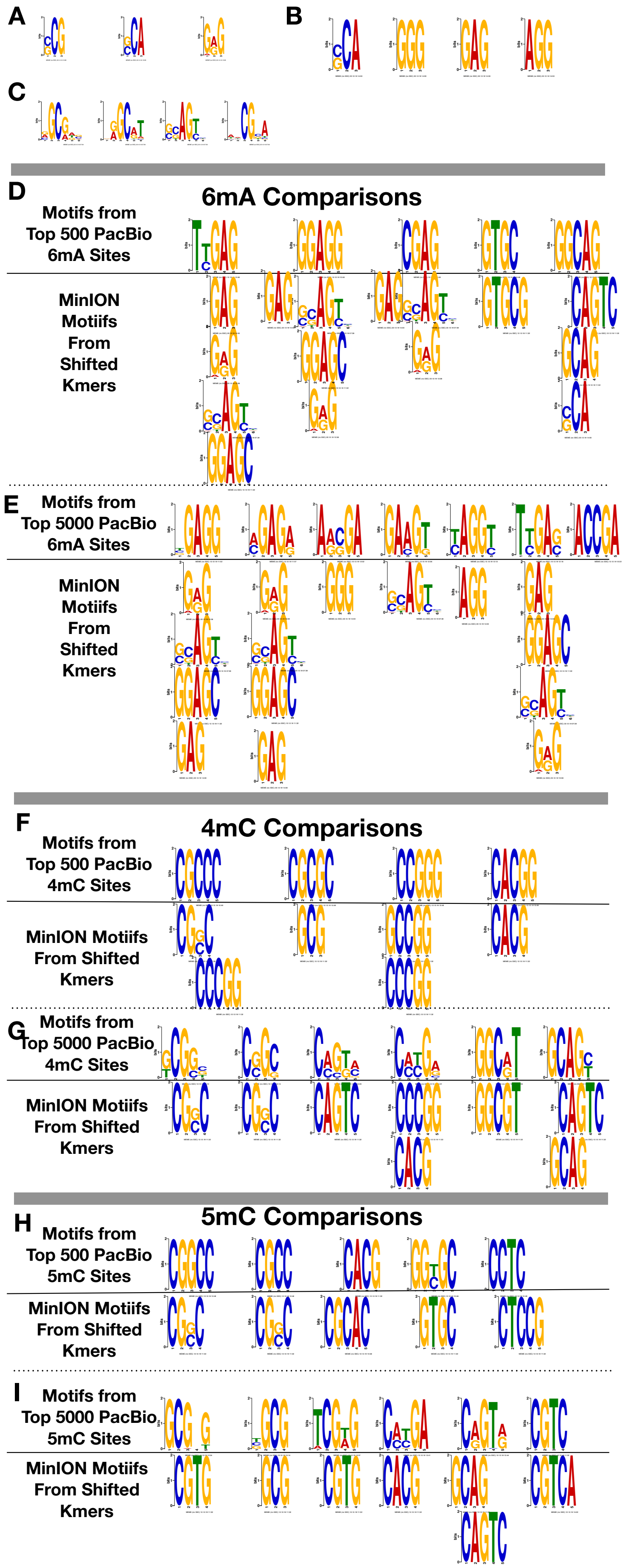

**Supplemental Figure S18: MEME motifs in the PacBio and MinION analyses**

**(A)** The top 3 trimer motifs in MEME analysis of kmers with shifted MinION signal.

**(B)** The top 4 trimer motifs in MEME analysis of kmers with shifted MinION signal when removing kmers with the CG dinucleotide from consideration.

**(C)** The top 4 6mer motifs in MEME analysis of kmers with shifted MinION signal.

**(D)** MEME motifs from top 500 PacBio 6mA sites compared to similar 2-6 bp motifs found in kmers with shifted MinION signal.

**(E)** MEME motifs from top 5000 PacBio 6mA sites compared to similar 2-6 bp motifs found in kmers with shifted MinION signal.

**(F)** MEME motifs from top 500 PacBio 4mC sites compared to similar 2-6 bp motifs found in kmers with shifted MinION signal.

**(G)** MEME motifs from top 5000 PacBio 4mC sites compared to similar 2-6 bp motifs found in kmers with shifted MinION signal.

**(H)** MEME motifs from top 500 PacBio 5mC sites compared to similar 2-6 bp motifs found in kmers with shifted MinION signal.

**(I)** MEME motifs from top 5000 PacBio 5mC sites compared to similar 2-6 bp motifs found in kmers with shifted MinION signal.

#### Supplemental Figure S19: MinION signal distributions for 6mers defined by motifs learned in the PacBio analysis and negative controls

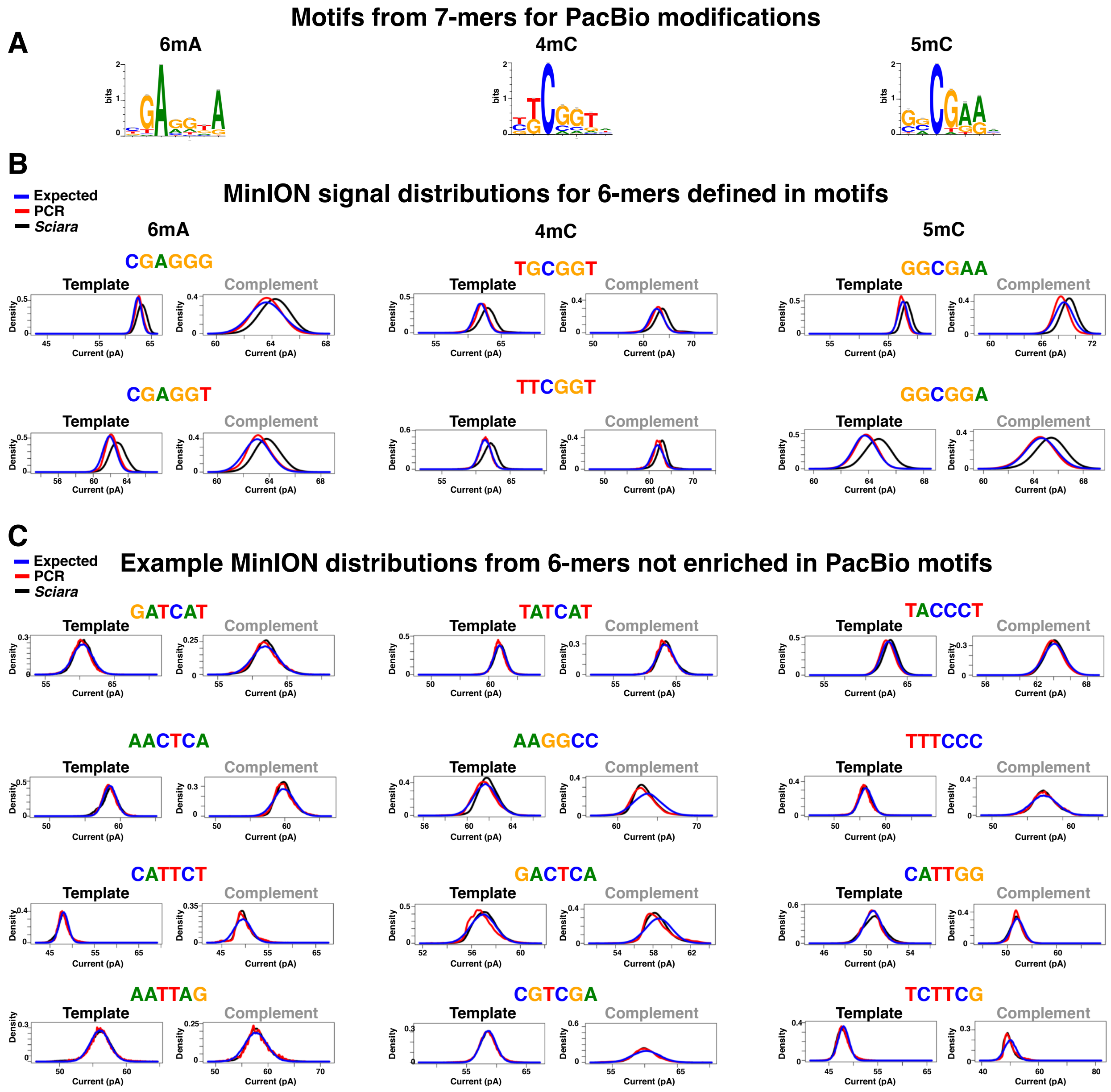

**Supplemental Figure S19: MinION signal distributions for 6mers defined by motifs learned in the PacBio analysis and negative controls**

**(A)** The motifs learned from PacBio analysis as shown elsewhere.

**(B)** MinION signal distributions for 6mers defined by motifs found for each DNA modification type in the PacBio analysis. For each 6mer, the expected distribution given the model provided by Oxford Nanopore Technologies (ONT) is shown in blue; the distribution learned from whole genome PCR of E. coli (using only canonical nucleotides) is shown in red; the distribution learned from native genomic DNA from male *Sciara* embryos is shown in black. Each 6mer shows the the expected and learned distributions for both the template and complement reads (R7.3).The distributions for *Sciara* are slightly offset from the model expectation whereas the learned model from PCR data is not.

**(C)** MinION signal distributions for 6mers not found enriched in the PacBio analysis. The distributions for *Sciara* match the distributions learned by PCR and expected from the model provided by ONT.

**Supplemental Figure S20: The GCG trimer is depleted in the genome and transcriptome compared to expectation.**

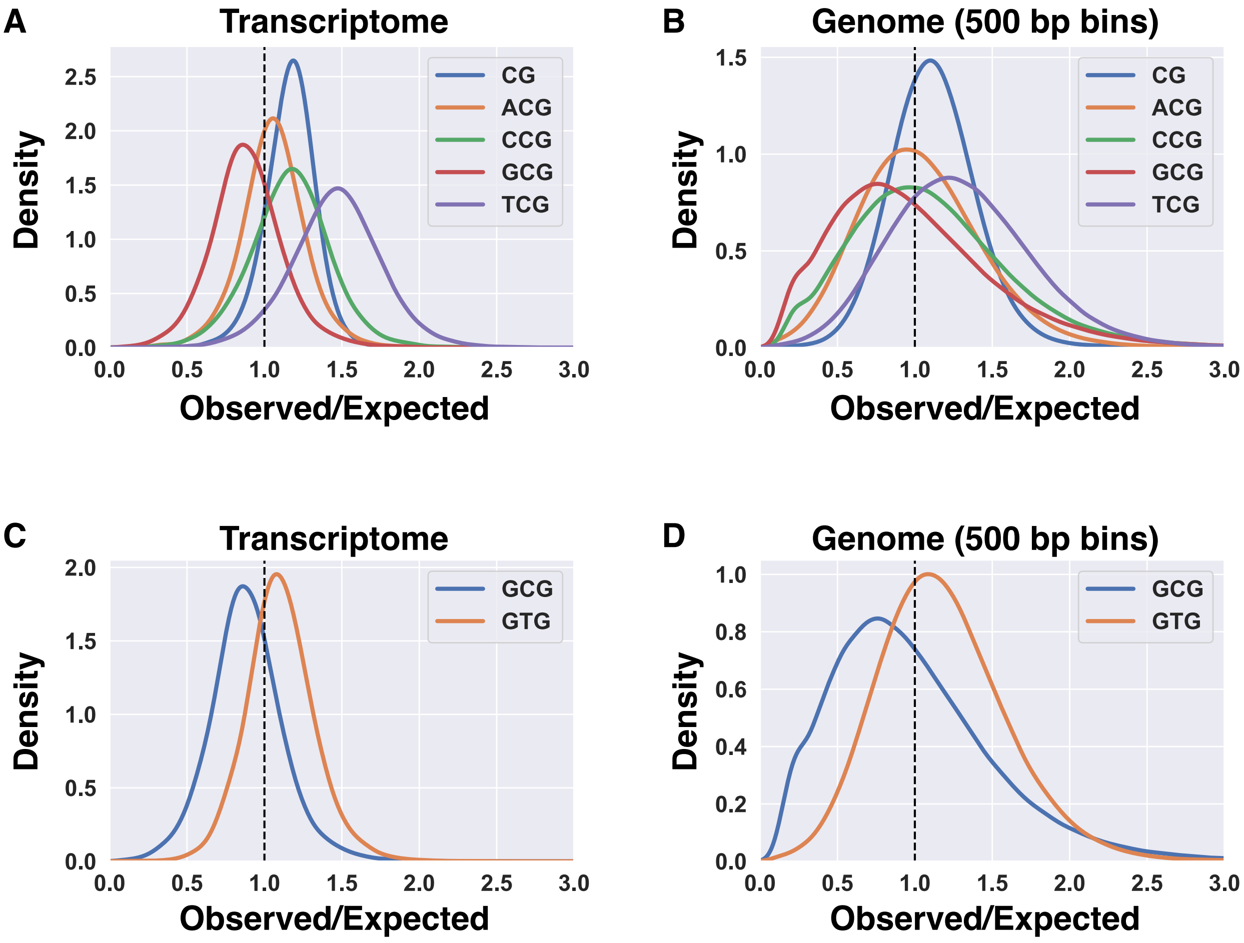

**Supplemental Figure S20: the GCG trimer is depleted in the genome and transcriptome compared to expectation.**
Methylated cytosines can spontaneously convert to thymine by deamination and thus germline CpG methylation predicts a depletion of CpG dimers in the genome over evolutionary time. In contrast, we found that there are more CpG dimers in the genome sequence than expected given the GC content. This may that the germline methylation and conversion rate(s) may be too low for this analysis or may be offset by other mutagenic forces in the germline genome (e.g. non-CG cytosine methylation). Alternatively, it may imply that CG is not a methylation target in the germline or at least not in all contexts. In support of the latter possibility, we found that GCG trimers were more enriched for DNA methylation than C or CG, which would predict a depletion of GCG trimers specifically. Interestingly, that is what we see. GCG trimers are depleted in the **(A)** transcriptome and **(B)** genome sequences with respect to expected levels (see supplemental methods). GCG depletion in the germline by way of C→T conversion over evolutionary timescales would also predict that GTG is enriched over expectation, which we also see in the **(C)** transcriptome and **(D)** genome sequences.

**Supplemental Figure S21: Distribution of distances between adjacent DNA modifications (PacBio analysis) on the same strand shows enrichment of short distances, a 10 bp periodicity, and a spike of enrichment at mono-nucleosome lengths of ~175 bp.**

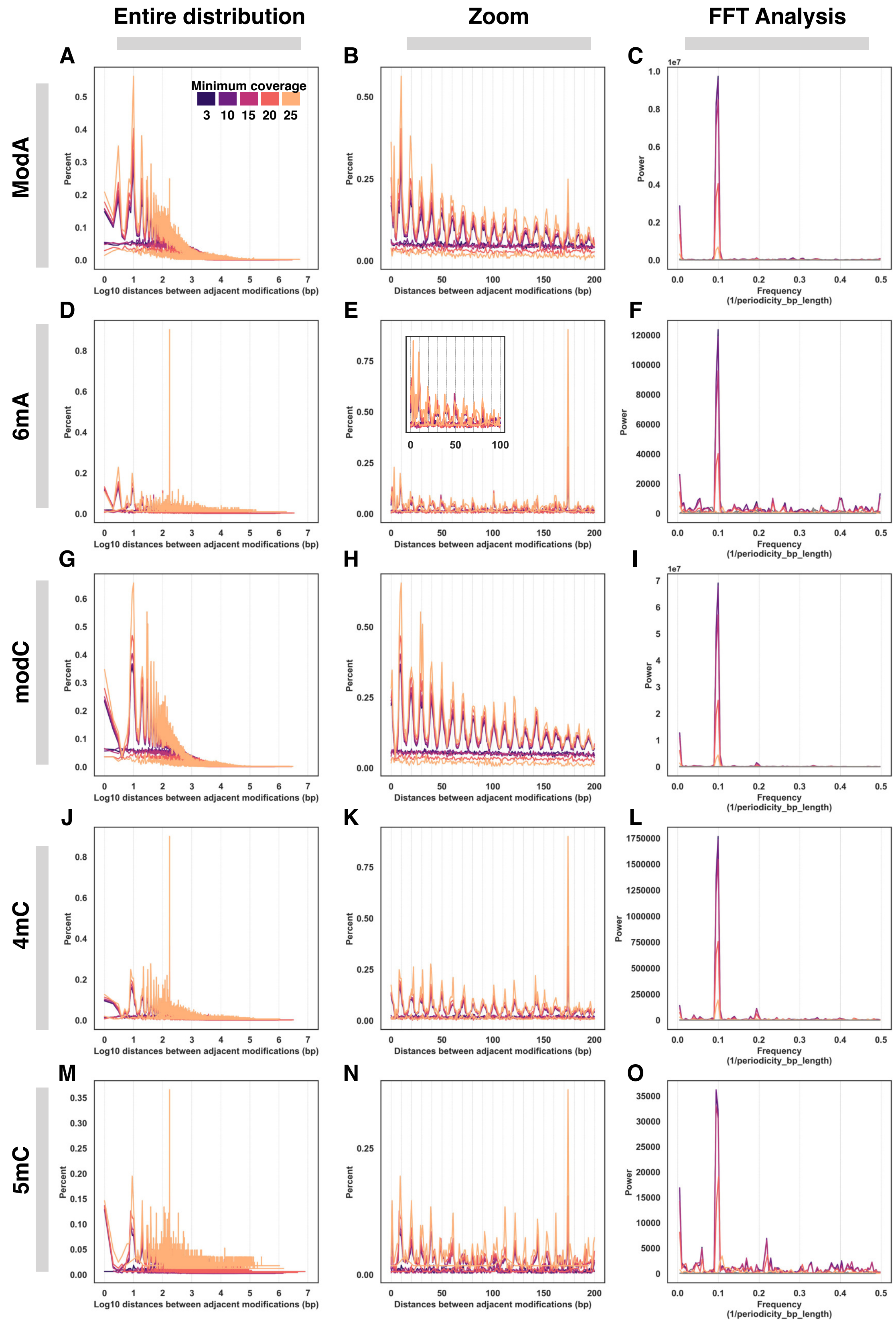

**Supplemental Figure S22: Distribution of distances between adjacent DNA modifications (PacBio analysis) on either strand also shows enrichment of short distances, a 10 bp periodicity, and a spike of enrichment at mono-nucleosome lengths of ~175 bp for 6mA and 5mC.**

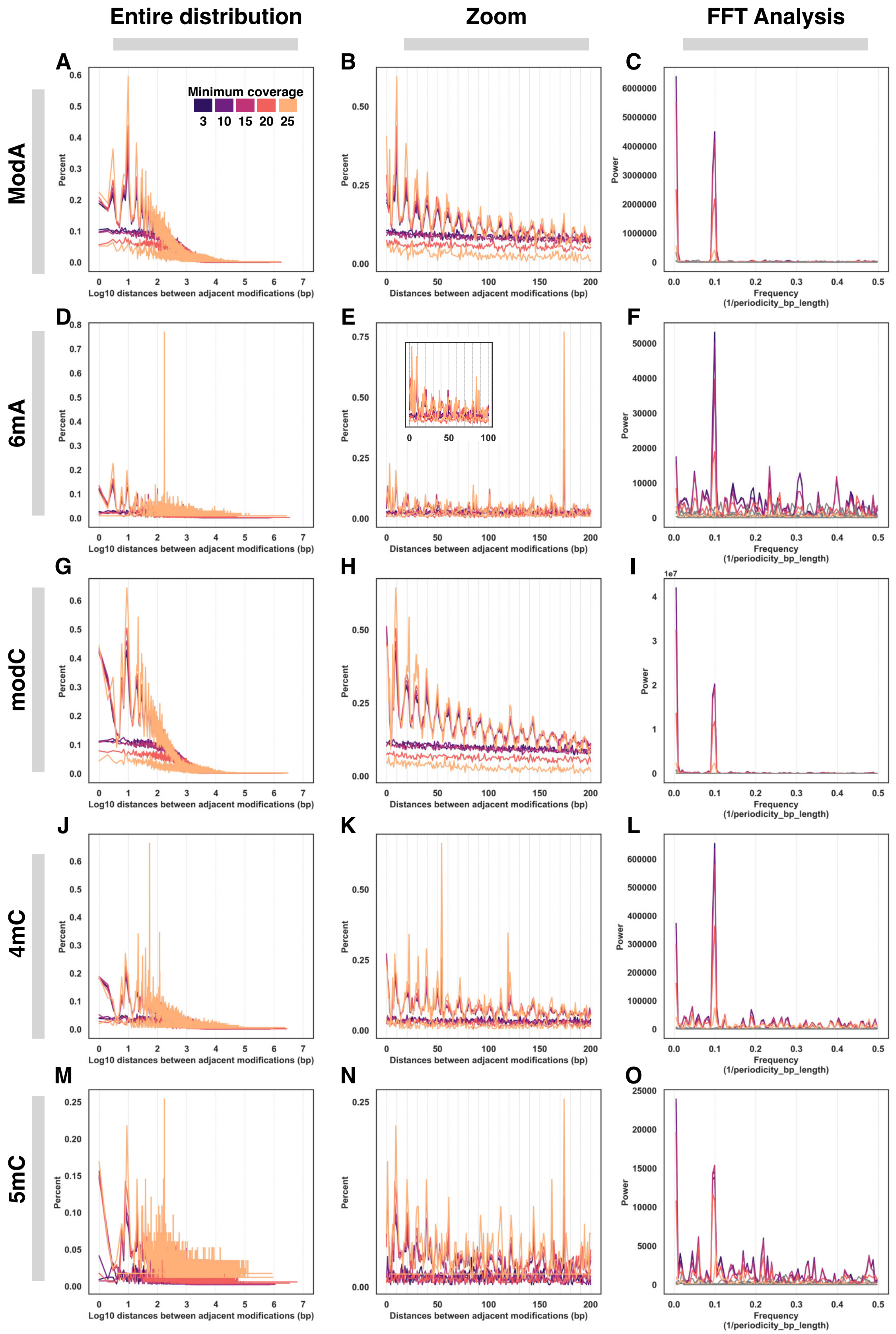

**Legend for both Supplemental Figure S21 and S22:**
For each row that features a given type of modification identified in the PacBio analysis, the left column shows the entire distribution of log10 distances between adjacent modifications normalized as the percent of all distances, the middle column shows a zoom in of distances between 0-200 bp normalized as the percent of all distances, and the right column shows the fast fourier transform analysis of the distances to identify the frequency with the highest power that is converted to periodicity by 1/frequency (1/0.1 = 10 bp). The inset in the E panels shows a further zoom in to distances between 0-100 to illustrate that the wavelength pattern is there, but looks extremely compressed due to the high spike at ~175 bp.

modA is the set of adenine sites flagged as modified by kineticsTools and 6mA is the subset therein identified specifically as 6mA. modC is the set of cytosine sites flagged as modified by kineticsTools whereas 4mC and 5mC are the subsets therein specifically identified as those modifications. Sites with modification scores (phred-transformed P-value that a kinetic deviation exists) <20 were filtered out and each plot shows the results of further filtering for sites with minimum coverages of 3X, 10X, 15X, 20X, or 25X. Moreover, ModA and ModC can be considered the unfiltered set of all putative modified sites (given the score and coverage cutoffs) whereas 6mA, 4mC, and 5mC can be considered the further filtered subsets of those sites that met classification criteria for those modifications. Finally, we also looked at the distances between neighboring modifications on the same strand or neighboring modifications on either strand. The results and conclusions were highly robust in all scenarios, with slightly stronger and cleaner effects in the strand-specific analysis.

In the left and middle columns that show distributions of distances, there are pairs of tests and controls with matching colors based on the minimum coverage used. The tests are the actual distribution of distances between DNA modifications and are the set of wave-like lines above the flatter ones (controls). The controls are the distances between the same number of “modifications” as the test obtained by randomly selecting adenines for modA/6mA or randomly selected cytosines for modC/4mC/5mC. The control lines are the set of relatively flat lines under the taller waves (tests). Control line colors match the colors for the paired test. For the FFT analysis, test lines are the same colors, but control lines were made grey as they all are essentially flat lines at 0.

**Supplemental Figure S23: BlobTools analysis of assemblies produced from long and short reads that did not map to the Bcop_v1 (Canu) *Sciara* genome assembly.**

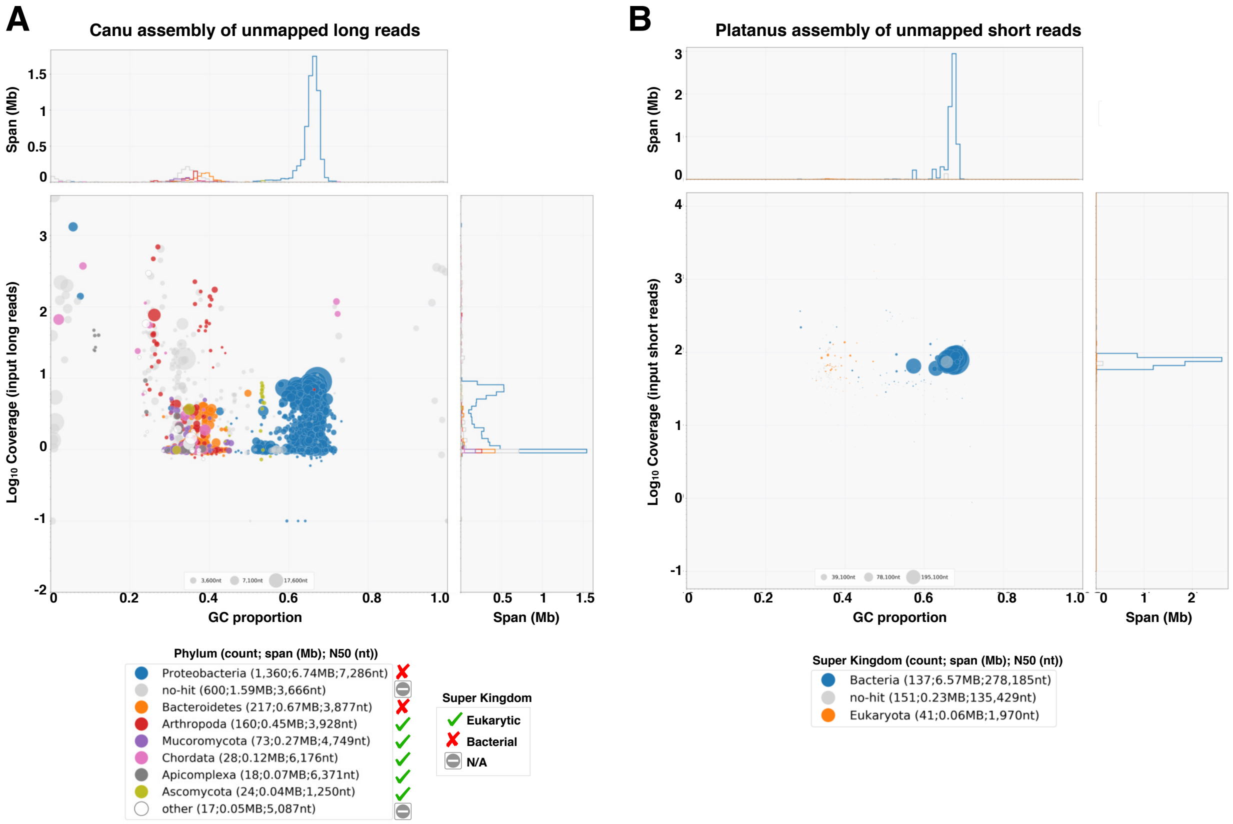

**Supplemental Figure S23: BlobTools analysis of assemblies produced from long and short reads that did not map to the Bcop_v1 (Canu) *Sciara* genome assembly.** (A) BlobPlot of the assembly of unmappable long reads. (B) BlobPlot of the assembly of unmappable short reads. The majority in both cases comes from bacterial species. Arthropod sequences tend to fall between 20-45% GC content whereas bacterial contigs tend to be >50% GC content in these plots.

### Section 2: Supplemental Tables

#### Supplemental Table S1 A-E: Expected genome and chromosome sizes

##### Supplementary Table S1A

| DNA content measurements (Rasch 2006) | | | | | | | |
|---|---|---|---|---|---|---|---|
| Sex | Stage / Tissue | Cell Type | Chromosomes* | N | Mean (pg) | Std Error (pg) | Total Length (Mb)** |
| Female | 4th instar larvae | Hemocytes | 2X+2A | 65 | 0.571 | 0.004 | 558.438 |
| Male | 4th instar larvae | Hemocytes | 1X+2A | 77 | 0.523 | 0.007 | 511.494 |
| Female | Pharate pupae | Hemocytes | 2X+2A | 74 | 0.591 | 0.004 | 577.998 |
| Male | Pharate pupae | Hemocytes | 1X+2A | 38 | 0.540 | 0.003 | 528.12 |
| Female | Adult | Hemocytes | 2X+2A | 74 | 0.554 | 0.003 | 541.812 |
| Female | Adult | Hemocytes | 2X+2A | 62 | 0.557 | 0.004 | 544.746 |
| Male | Adult | Hemocytes | 1X+2A | 100 | 0.503 | 0.002 | 491.934 |
| Male | Adult 44A testis | Sperm | 2X+1A+2L | 162 | 0.491 | 0.016 | 480.198 |
| Male | Adult 44A duct | Sperm | 2X+1A+2L | 60 | 0.494 | 0.017 | 483.132 |
| Male | Adult 43 E testis | Sperm | 2X+1A+2L | 100 | 0.519 | 0.018 | 507.582 |
| Male | Adult 43 E duct | Sperm | 2X+1A+2L | 50 | 0.521 | 0.013 | 509.538 |
| Male | Adult 45 testis | Sperm | 2X+1A+2L | 65 | 0.535 | 0.014 | 523.23 |
| Male | Adult 45 duct | Sperm | 2X+1A+2L | 50 | 0.484 | 0.019 | 473.352 |

*Assumes two L chromosomes are present in the sperm. Though this is most common, it can range from 0-4.
** The Mb length of all DNA in the nucleus given pg mean and Dolezel conversion (Dolezel et al. 2003).

##### Supplementary Table S1B

| Chromosome X | | | | |
|---|---|---|---|---|
| Stage | Female (pg) | Male (pg) | Chromosome X (pg) (Difference) | Chromosome X (Mb) |
| Larvae | 0.571 | 0.523 | 0.048 | 46.944 |
| Pupae | 0.591 | 0.540 | 0.051 | 49.878 |
| Adult* | 0.555 | 0.503 | 0.052 | 50.856 |
| Mean | 0.572 | 0.522 | 0.05 | 48.9 |

*Since two female adult values are in Table A, the female adult value was obtained by: (74 ∗ 0.554 + 62 ∗ 0.557)/(74 + 62) = 0.555

##### Supplementary Table S1C

| Expected length of the haploid somatic genome: Chromosomes X, II, III, and IV | | | | | | | |
|---|---|---|---|---|---|---|---|
| Stage | Sex | Total DNA content (pg) | X DNA content (pg) | Total – X (pg) | Diploid length of autosomes: II, III, and IV (Mb) | Haploid length of autosomes: II, III, and IV (Mb) | Haploid length of X + autosomes (Mb) |
| Larvae | Female | 0.571 | 0.1 | 0.471 | 460.638 | 230.319 | 279.219 |
| Larvae | Male | 0.523 | 0.05 | 0.473 | 462.594 | 231.297 | 280.197 |
| Pupae | Female | 0.591 | 0.1 | 0.491 | 480.198 | 240.099 | 288.999 |
| Pupae | Male | 0.540 | 0.05 | 0.490 | 479.22 | 239.61 | 288.51 |
| Adult | Female | 0.555 | 0.1 | 0.455 | 444.99 | 222.495 | 271.395 |
| Adult | Male | 0.503 | 0.05 | 0.453 | 443.034 | 221.517 | 270.417 |
| **Mean** | **-** | **-** | **-** | **0.472** | **461.779** | **230.889** | **279.789** |

*Haploid estimates range from ~270-290 Mb

**Supplementary Table S1D**

| DNA content contribution from L chromosomes in sperm and expected germline genome size | | | | | | | |
|---|---|---|---|---|---|---|---|
| **DNA content in sperm*** | **Length of DNA in Sperm: 2X+1A+2L** (Mb)** | **Length of haploid autosomes (Mb)** | **Length of X (Mb)** | **Length of 2X + 1 set of autosomes (Mb)** | **Length of contribution from L chromosomes: Total – (2X+A) (Mb)** | **Length of L** | **Haploid germline size (1A + 1X + 1L)** |
| 0.5053531 | 494.235 | 230.889 | 48.9 | 328.689 | 165.546 | 82.773 | 362.562 |

* The weighted average for DNA content in sperm was the sum(N*mean)/sum(N) for the N and M values in Table A.
**Assumes two L chromosomes are present in the sperm. Though this is most common, it can range from 0-4. If one assumed the most majority of sperm had 1, 2, 3, or 4 Ls, the possible L chromosome lengths are between 41.4-165.5 Mb and the haploid genome size is between 321.2-445.3 Mb.

**Supplementary Table S1E**

| Expected X and Autosome Lengths given polytene zones (Gabrusewycz-Garica 1964) | | | | | |
|---|---|---|---|---|---|
| **Chromosome** | **Number of polytene zones** | **Expected proportion of somatic genome: (X, II, III, IV)** | **Expected Length (Mb) given haploid somatic genome size (279.789)** | **Expected proportion of autosomes: (II, III, IV)** | **Expected Length (Mb) given sum of haploid autosome sizes (230.889)** |
| X | 14 | 0.222222222 | 62.17533333 | 0 | (48.9) |
| II | 14 | 0.222222222 | 62.17533333 | 0.285714286 | 65.96828571 |
| III | 15 | 0.238095238 | 66.61642857 | 0.306122449 | 70.68030612 |
| IV | 20 | 0.317460317 | 88.82190476 | 0.408163265 | 94.24040816 |
| Sum | 63 | 1 | 279.789 | 1 | 279.789 |

**Supplementary Table S1 A-E: Expected genome and chromosome sizes**

**(A)** We wanted to estimate the approximate length of the haploid genome in the germline (ChrII, ChrIII, ChrIV, ChrX, ChrL) as well as the somatic haploid genome length (ChrII, ChrIII, ChrIV, ChrX). To do so, we used started with the values in Table 2 from Rasch (2006) (Rasch 2006), adapted here.
**(B)** Since male somatic cells have a single X and female somatic cells have two, the pg weight and bp length of a single chromosome X can be inferred by subtracting the male value from the female value.
**(C)** The diploid autosome DNA content and bp length can be inferred by subtracting the value of 1X for males or 2X for females. One can then deduce the haploid autosome DNA content and length by dividing by 2, and can subsequently deduce the expected somatic haploid genome (chrII, chrIII, chrIV, chrX) DNA content and length by adding the value of a single X. Thus, the genome assemblies we generate should be ~280 Mb reflecting the expected haploid somatic genome length (chrII, chrIII, chrIV, chrX).
**(D)** The expected length of the haploid germline genome can be deduced by adding the weight of a single L chromosome. The sperm data from sperm in Table S1A represents one set of autosomes, two X chromosomes, and a variable number of L chromosomes (most commonly two). So the L content can be found by subtracting 2X+A from the total. We took the weighted average DNA content in sperm. As discussed by Rasch (Rasch 2006) and demonstrated by Rieffel and Crouse (Rieffel and Crouse 1966), there can be between 0 and 4 copies of the L chromosome in a sperm nucleus, though the majority have 2 copies (78% according to Rasch). The length in the table assumes two L per sperm. Effects from other possibilities are noted too.
**(E)** Assuming the number of polytene zones correlates with chromosome length in base pairs, one can estimate the relative chromosome sizes compared to each other, and use those relative proportions with the estimated summed length of all four to estimate chromosome lengths. This results in an X chromosome length that longer than DNA content estimates (49 Mb vs 62 Mb). We can also fix the X length as estimated above and use the relative proportions of only chromosomes II, III, and IV with their estimated summed length from DNA content. Overall, we expect X to be 48-62 Mb, II to be 62-66 Mb, III to be 66-71 Mb, and IV to be 88-94 Mb.

#### Supplemental Table S2: RNA-seq samples spanning both sexes and 4 life cycle stages

| Sample | Replicate # | NEB Set # | Barcode Index | inPrimerSeq* | ReadSeq* | Number of paired-reads |
|---|---|---|---|---|---|---|
| **Female embryos** | 1 | 1 | index 1 | CGTGAT | ATCACG | 5,904,586 |
| **Female embryos** | 2 | 1 | index 2 | ACATCG | CGATGT | 6,136,865 |
| **Female embryos** | 3 | 1 | index 3 | GCCTAA | TTAGGC | 4,941,561 |
| **Male embryos** | 1 | 1 | index 4 | TGGTCA | TGACCA | 5,608,743 |
| **Male embryos** | 2 | 1 | index 5 | CACTGT | ACAGTG | 4,891,639 |
| **Male embryos** | 3 | 1 | index 6 | ATTGGC | GCCAAT | 6,942,968 |
| **Female Larvae** | 1 | 1 | index 7 | GATCTG | CAGATC | 5,465,807 |
| **Female Larvae** | 2 | 1 | index 8 | TCAAGT | ACTTGA | 5,776,190 |
| **Male Larvae** | 1 | 1 | index 9 | CTGATC | GATCAG | 6,453,818 |
| **Male Larvae** | 2 | 1 | index 10 | AAGCTA | TAGCTT | 5,694,394 |
| **Female Pupae** | 1 | 1 | index 11 | GTAGCC | GGCTAC | 5,177,694 |
| **Female Pupae** | 2 | 1 | index 12 | TACAAG | CTTGTA | 5,532,750 |
| **Male Pupae** | 1 | 2 | index 13 | TTGACT | AGTCAA | 15,280,707 |
| **Male Pupae** | 2 | 2 | index 14 | GGAACT | AGTTCC | 14,022,001 |
| **Female adults** | 1 | 2 | index 15 | TGACAT | ATGTCA | 5,485,023 |
| **Female adults** | 2 | 2 | index 16 | GGACGG | CCGTCC | 13,704,291 |
| **Male adults** | 1 | 2 | index 18* | GCGGAC | GTCCGC | 11,741,449 |
| **Male adults** | 2 | 2 | index 19 | TTTCAC | GTGAAA | 10,582,159 |

#### Supplemental Table S3: Short read assembly size statistics

| Assembly | N | ASM_SIZE | MAX | MIN | MEAN | MEDIAN | N50 | L50 | E | NG50 280M | LG50 280M | E 280M | Matrix Row | Nickname |
|---|---|---|---|---|---|---|---|---|---|---|---|---|---|---|
| **abyss.k55** | 303676 | 277836422 | 1054107 | 55 | 915 | 109 | 5938 | 12383 | 18271 | 5880 | 12566 | 18130 | 1 | a1 |
| **abyss.k55.bh** | 302988 | 278198827 | 1054115 | 55 | 918 | 109 | 5993 | 12280 | 18542 | 5943 | 12431 | 18422 | 2 | a2 |
| **abyss.k55.q5** | 303987 | 276373215 | 1025867 | 55 | 909 | 109 | 5796 | 12678 | 16597 | 5695 | 12993 | 16382 | 3 | a3 |
| **abyss.k55.q5.bh** | 303005 | 276740830 | 1053964 | 55 | 913 | 109 | 5851 | 12559 | 17462 | 5764 | 12840 | 17258 | 4 | a4 |
| **abyss.k77** | 186647 | 273406983 | 1060849 | 77 | 1465 | 169 | 5026 | 14870 | 15634 | 4875 | 15536 | 15265 | 5 | a5 |
| **abyss.k77.bh** | 186091 | 275264727 | 1054512 | 77 | 1479 | 168 | 5120 | 14670 | 16138 | 5024 | 15137 | 15865 | 6 | a6 |
| **abyss.k77.q5** | 188860 | 273874388 | 707552 | 77 | 1450 | 171 | 4971 | 15031 | 13590 | 4833 | 15655 | 13292 | 7 | a7 |
| **abyss.k77.q5.bh** | 186022 | 273581412 | 1054487 | 77 | 1471 | 170 | 5039 | 14825 | 15353 | 4893 | 15471 | 15001 | 8 | a8 |
| **megahit** | 203144 | 314764763 | 805548 | 200 | 1549 | 540 | 4309 | 17696 | 13631 | 5123 | 13994 | 15323 | 9 | m1 |
| **megahit.bh** | 200747 | 312783966 | 687499 | 200 | 1558 | 539 | 4340 | 17552 | 13114 | 5104 | 14065 | 14650 | 10 | m2 |
| **megahit.q5** | 194936 | 308417993 | 682464 | 200 | 1582 | 541 | 4379 | 17352 | 12459 | 5053 | 14330 | 13723 | 11 | m3 |
| **megahit.q5.bh** | 194640 | 308016525 | 682470 | 200 | 1582 | 542 | 4379 | 17394 | 12473 | 5034 | 14408 | 13721 | 12 | m4 |
| **platanus** | 88584 | 235992359 | 801720 | 81 | 2664 | 1354 | 5454 | 11586 | 17358 | 4287 | 16133 | 14630 | 13 | p1 |
| **platanus.bh** | 88132 | 237450965 | 936960 | 96 | 2694 | 1359 | 5536 | 11484 | 18261 | 4418 | 15787 | 15486 | 14 | p2 |
| **platanus.q5** | 88177 | 240633956 | 908679 | 94 | 2729 | 1373 | 5660 | 11477 | 17730 | 4616 | 15333 | 15238 | 15 | p3 |
| **platanus.q5.bh** | 87570 | 241582189 | 908703 | 81 | 2759 | 1382 | 5755 | 11387 | 17501 | 4718 | 15079 | 15100 | 16 | p4 |
| **sga** | 160775 | 266601738 | 907684 | 200 | 1658 | 628 | 3880 | 18376 | 12870 | 3642 | 20157 | 12254 | 17 | sg1 |
| **sga.q5** | 167119 | 264933802 | 907675 | 200 | 1585 | 607 | 3702 | 19463 | 12648 | 3452 | 21571 | 11967 | 18 | sg2 |
| **soapdenovo2.k55** | 339967 | 315872819 | 907788 | 100 | 929 | 115 | 6525 | 12023 | 20769 | 7729 | 9497 | 23430 | 19 | so1 |
| **soapdenovo2.k55.bh** | 357836 | 318047022 | 909706 | 100 | 889 | 111 | 6327 | 12399 | 19605 | 7609 | 9658 | 22269 | 20 | so2 |
| **soapdenovo2.k55.q5** | 315548 | 311171324 | 907784 | 100 | 986 | 126 | 6596 | 11897 | 19175 | 7630 | 9700 | 21310 | 21 | so3 |
| **soapdenovo2.k55.q5.bh** | 302550 | 310634015 | 907798 | 100 | 1027 | 145 | 6589 | 11855 | 19173 | 7625 | 9695 | 21271 | 22 | so4 |
| **soapdenovo2.k77** | 244672 | 282948474 | 907817 | 100 | 1156 | 155 | 5747 | 13036 | 17684 | 5828 | 12781 | 17870 | 23 | so5 |
| **soapdenovo2.k77.bh** | 261683 | 285615548 | 908360 | 100 | 1091 | 155 | 5710 | 13165 | 17883 | 5869 | 12681 | 18242 | 24 | so6 |
| **soapdenovo2.k77.q5** | 237972 | 280534633 | 1025525 | 100 | 1179 | 155 | 5621 | 13222 | 17567 | 5635 | 13175 | 17600 | 25 | so7 |
| **soapdenovo2.k77.q5.bh** | 237446 | 281739587 | 907721 | 100 | 1187 | 155 | 5776 | 12954 | 17037 | 5825 | 12804 | 17143 | 26 | so8 |
| **spades.bh.k213355** | 380982 | 347635213 | 1026350 | 1 | 912 | 146 | 4229 | 19112 | 15992 | 5882 | 12309 | 19855 | 27 | sp1 |
| **spades.bh.k21335577** | 286478 | 348582982 | 1228455 | 18 | 1217 | 252 | 5552 | 14564 | 22018 | 7768 | 9340 | 27411 | 28 | sp2 |
| **spades.bh.k21335577auto** | 279669 | 345817875 | 1228455 | 18 | 1237 | 249 | 5630 | 14317 | 22188 | 7768 | 9340 | 27404 | 29 | sp3 |
| **spades.q5.bh.k213355** | 382055 | 344292232 | 1220242 | 1 | 901 | 152 | 4214 | 18998 | 16637 | 5803 | 12479 | 20457 | 30 | sp4 |
| **spades.q5.bh.k21335577** | 278353 | 343736955 | 1227608 | 18 | 1235 | 260 | 5618 | 14336 | 20503 | 7675 | 9477 | 25170 | 31 | sp5 |
| **spades.q5.bh.k21335577auto** | 271253 | 340884856 | 1227608 | 18 | 1257 | 257 | 5703 | 14085 | 20668 | 7675 | 9478 | 25163 | 32 | sp6 |
| **velvet.k55** | 160994 | 233337453 | 1109576 | 109 | 1449 | 350 | 4202 | 15121 | 18120 | 3266 | 21424 | 15100 | 33 | v1 |
| **velvet.k55.bh** | 186320 | 235611858 | 1110712 | 109 | 1265 | 337 | 3515 | 18288 | 15265 | 2780 | 25390 | 12845 | 34 | v2 |
| **velvet.k55.q5** | 159581 | 231243264 | 1108829 | 109 | 1449 | 352 | 4189 | 15075 | 17438 | 3217 | 21718 | 14402 | 35 | v3 |
| **velvet.k55.q5.bh** | 159511 | 233215384 | 1108932 | 109 | 1462 | 358 | 4211 | 15078 | 17083 | 3280 | 21374 | 14229 | 36 | v4 |
| **velvet.k77** | 125780 | 229591350 | 1109932 | 153 | 1825 | 823 | 4075 | 15985 | 16289 | 3158 | 23010 | 13356 | 37 | v5 |
| **velvet.k77.bh** | 125522 | 232398455 | 1110079 | 153 | 1851 | 851 | 4104 | 16073 | 17076 | 3237 | 22606 | 14173 | 38 | v6 |
| **velvet.k77.q5** | 124497 | 226321787 | 882517 | 153 | 1818 | 824 | 4045 | 15879 | 14163 | 3066 | 23502 | 11448 | 39 | v7 |
| **velvet.k77.q5.bh** | 124026 | 230127881 | 1111298 | 153 | 1855 | 850 | 4113 | 15846 | 17911 | 3203 | 22718 | 14721 | 40 | v8 |
| **platanus.blobfilt1** | 87119 | 227977694 | 75265 | 81 | 2617 | 1368 | 5314 | 11982 | 7242 | 3979 | 17647 | 5897 | NA | p5 |
| **platanus.blobfilt1.bh** | 87696 | 228227902 | 75308 | 96 | 2602 | 1355 | 5299 | 12005 | 7233 | 3985 | 17647 | 5895 | NA | p6 |
| **platanus.blobfilt1.q5** | 86482 | 232676523 | 60214 | 95 | 2690 | 1393 | 5519 | 11815 | 7472 | 4283 | 16683 | 6209 | NA | p7 |
| **platanus.blobfilt1.q5.bh** | 87509 | 229834980 | 63160 | 99 | 2626 | 1366 | 5350 | 12030 | 7269 | 4073 | 17403 | 5967 | NA | p8 |

**Supplemental Table S3: Short read assembly size statistics**
The naming scheme is "*assembler.parameters.input*" where parameters for some assemblers were kmer sizes of 55 or 77 (k55, k77), or up to k=55 or k=77 for SPAdes (e.g. k21335577 used kmer sizes 21, 33, 55, and 77), and where the input was quality-filtered (q5), error-corrected (bh), both (q5.bh), or the raw input reads otherwise. For SPAdes, "auto" means "—cov-cutoff auto" was set. For Platanus, "blobfilt1" indicates that these were "re-assemblies" with contaminating reads removed after blobtools analysis (Laetsch and Blaxter 2017). The assemblies are presented in the same order that their ranks are visualized in the ranking matrix in Figure 2B. The "Matrix Row" and "Nickname" columns additionally help connect the table with that figure.

#### Supplemental Table S4: Long read assembly size statistics

| Assembly | N | ASM_SIZE | MAX | MIN | MEAN | MEDIAN | N50 | L50 | E | NG50 280M | LG50 280M | E 280M | Matrix Row | Nickname |
|---|---|---|---|---|---|---|---|---|---|---|---|---|---|---|
| abruijn.pball.ontmol | 544 | 285817077 | 10540070 | 53798 | 525399 | 227013 | 1222757 | 58 | 2112258 | 1281449 | 56 | 2156141 | 1 | A1 |
| abruijn.pball | 732 | 281500939 | 5690677 | 50217 | 384564 | 161202 | 826831 | 76 | 1516868 | 832247 | 75 | 1525000 | 2 | A2 |
| abruijn.pbfilt.ont2d | 541 | 284906646 | 10747290 | 48902 | 526630 | 210027 | 1294301 | 49 | 2475897 | 1338369 | 47 | 2519284 | 3 | A3 |
| canu10.minrl1000.dip3x.pbfilt | 1316 | 303769988 | 21228270 | 1732 | 230828 | 21211 | 2554706 | 23 | 5321262 | 3425454 | 19 | 5772999 | 4 | C0.1 |
| canu10.minrl500.dip3x.pball | 1236 | 303631862 | 26730757 | 1213 | 245657 | 21871 | 2570070 | 27 | 5442641 | 3008968 | 22 | 5901998 | 5 | C0.2 |
| canu11.corcov80minrl500.dip3x.pball.ontmol | 1537 | 296991896 | 20910109 | 3770 | 193228 | 32147 | 1188978 | 42 | 3975708 | 1337456 | 35 | 4216975 | 6 | C1.1 |
| canu12.corcov80minrl500.dip3x.pball.ontmol | 1225 | 299099912 | 17969101 | 4139 | 244163 | 34799 | 2399381 | 26 | 4373842 | 2819977 | 23 | 4672199 | 7 | C2.1 |
| canu12.minrl500.dip3x.pball.ontmol | 1213 | 297711070 | 19547680 | 4524 | 245434 | 39809 | 1958195 | 30 | 4530411 | 2149314 | 26 | 4816976 | 8 | C2.2 |
| canu.corcov500minrl500.aspb-e025.pball.ontmol | 1036 | 302365066 | 12318386 | 793 | 291858 | 28771 | 1908694 | 37 | 3173265 | 2367475 | 31 | 3426730 | 9 | C3.1 |
| canu.corcov500minrl500.aspb-e02.pball.ont2d | 1044 | 302258594 | 12317152 | 671 | 289520 | 28311 | 1908286 | 37 | 3173951 | 2363127 | 31 | 3426264 | 10 | C3.2 |
| canu.corcov500minrl500.aspb-e02.pbfilt.ont2d | 1044 | 302239832 | 12315397 | 671 | 289502 | 28307 | 1908501 | 37 | 3173821 | 2361875 | 31 | 3425912 | 11 | C3.3 |
| canu.corcov500minrl500.pball.ont2d | 1104 | 303257430 | 12695061 | 792 | 274690 | 31720 | 1506847 | 45 | 2690343 | 1724831 | 38 | 2913809 | 12 | C3.4 |
| canu.corcov500minrl500.pball.ontmol | 1107 | 303232626 | 12696256 | 792 | 273923 | 31352 | 1516021 | 44 | 2721930 | 1777033 | 37 | 2947779 | 13 | C3.5 |
| canu.corcov500minrl500.pball | 1231 | 300698388 | 11589796 | 587 | 244272 | 27608 | 1764129 | 42 | 2623585 | 1918491 | 36 | 2817528 | 14 | C3.6 |
| canu.corcov500minrl500.pbfilt | 1224 | 300534892 | 11589685 | 793 | 245535 | 29497 | 1654609 | 43 | 2722880 | 1823318 | 37 | 2922573 | 15 | C3.7 |
| canu.corcov500.pball | 1240 | 300804020 | 11589802 | 1133 | 242584 | 28527 | 1674691 | 43 | 2580825 | 1848220 | 37 | 2772581 | 16 | C3.8 |
| canu.corcov500.pbfilt.ont2d | 1139 | 301454870 | 10743861 | 1023 | 264666 | 30368 | 1391244 | 47 | 2535311 | 1603630 | 40 | 2729578 | 17 | C3.9 |
| canu.corcov500.pbfilt | 1232 | 300545669 | 11589555 | 1133 | 243949 | 29275 | 1676452 | 41 | 2993594 | 1823280 | 35 | 3213256 | 18 | C3.10 |
| canu.default.pball | 1192 | 298757505 | 12235276 | 1189 | 250635 | 30880 | 1649665 | 42 | 2860691 | 1832171 | 37 | 3052331 | 19 | C3.11 |
| canu.minrl500.pball | 1226 | 299103658 | 12235409 | 792 | 243967 | 28570 | 1636496 | 42 | 2868504 | 1832154 | 37 | 3064214 | 20 | C3.12 |
| falcon.default.pball | 702 | 294632888 | 11528295 | 406 | 419705 | 32855 | 2516799 | 29 | 3755975 | 2730335 | 26 | 3952264 | 21 | F1 |
| falcon.default.pbfilt | 704 | 295192605 | 12753606 | 283 | 419308 | 30728 | 2785086 | 29 | 3850555 | 2973441 | 27 | 4059483 | 22 | F2 |
| falcon.seed25.pball.ont2d | 728 | 297366589 | 11078332 | 519 | 408471 | 22849 | 2546290 | 30 | 3585374 | 2596709 | 27 | 3807751 | 23 | F3 |
| falcon.seed25.pball.ontmol | 737 | 296668450 | 12400955 | 406 | 402535 | 25656 | 2519867 | 28 | 3809884 | 2730274 | 25 | 4036687 | 24 | F4 |
| falcon.seed25.pball | 671 | 295939241 | 12785397 | 148 | 441042 | 25122 | 3072189 | 28 | 3978720 | 3456700 | 25 | 4205212 | 25 | F5 |
| falcon.seed25.pbfilt.ont2d | 724 | 297400499 | 11078438 | 519 | 410774 | 22977 | 2546297 | 30 | 3585986 | 2596645 | 27 | 3808836 | 26 | F6 |
| falcon.seed25.pbfilt | 712 | 295735287 | 9828760 | 145 | 415359 | 21976 | 2943872 | 28 | 3599286 | 3316978 | 26 | 3801557 | 27 | F7 |
| falcon.seed25.relaxed.pball.ont2d | 709 | 296742085 | 12794951 | 117 | 418536 | 19083 | 2626834 | 28 | 3927881 | 3100919 | 25 | 4162742 | 28 | F8 |
| falcon.seed25.relaxed.pball.ontmol | 713 | 296559753 | 20165452 | 119 | 415932 | 18659 | 2878051 | 24 | 5050163 | 3454019 | 21 | 5348840 | 29 | F9 |
| falcon.seed25.relaxed.pbfilt.ont2d | 705 | 296734699 | 12795016 | 117 | 420900 | 19089 | 2626878 | 28 | 3928187 | 3100874 | 25 | 4162962 | 30 | F10 |
| falcon.seed30.pball | 703 | 296128944 | 15921957 | 148 | 421236 | 19231 | 2883831 | 30 | 3890853 | 3014947 | 27 | 4114979 | 31 | F11 |
| falcon.seed30.pbfilt | 713 | 296392938 | 12458309 | 86 | 415698 | 19624 | 2564544 | 29 | 3845400 | 2929617 | 26 | 4070533 | 32 | F12 |
| miniasm.racon.pball.ont2d | 682 | 290016397 | 7561531 | 4803 | 425244 | 150330 | 1161069 | 64 | 1732387 | 1221342 | 60 | 1794359 | 33 | M1 |
| miniasm.racon.pball.ontall | 669 | 289847742 | 7563691 | 2427 | 433255 | 154789 | 1179233 | 62 | 1736678 | 1237602 | 58 | 1797758 | 34 | M2 |
| miniasm.racon.pball.ontmol | 691 | 289860660 | 7565108 | 2576 | 419480 | 146701 | 1095353 | 67 | 1682131 | 1161141 | 63 | 1741370 | 35 | M3 |
| miniasm.racon.pball | 913 | 289699308 | 6296002 | 1616 | 317305 | 103582 | 1008157 | 66 | 1632180 | 1100088 | 61 | 1688720 | 36 | M4 |
| miniasm.racon.pbfilt.ont2d | 687 | 290025924 | 7561922 | 4803 | 422163 | 146787 | 1161098 | 61 | 1807223 | 1234830 | 57 | 1871934 | 37 | M5 |
| miniasm.racon.pbfilt.ontall | 671 | 289995708 | 7560375 | 4803 | 432184 | 150177 | 1161104 | 62 | 1794981 | 1234832 | 58 | 1859060 | 38 | M6 |
| miniasm.racon.pbfilt.ontmol | 689 | 290105515 | 7560362 | 4803 | 421053 | 145499 | 1161100 | 61 | 1816611 | 1234831 | 57 | 1882174 | 39 | M7 |
| miniasm.racon.pbfilt | 920 | 289863434 | 8266389 | 4948 | 315069 | 104506 | 962801 | 67 | 1724677 | 1100085 | 62 | 1785432 | 40 | M8 |
| platanus.dbg2olc.pball.ont2d | 1309 | 304011299 | 2365216 | 1000 | 232247 | 104763 | 519314 | 176 | 652377 | 557486 | 153 | 708321 | 41 | PD1 |
| platanus.dbg2olc.pball.ontmol | 1385 | 306594506 | 2113652 | 999 | 221368 | 102361 | 473926 | 192 | 605128 | 513068 | 165 | 662603 | 42 | PD2 |
| platanus.dbg2olc.pball | 1491 | 302754463 | 1940333 | 1479 | 203055 | 113097 | 396110 | 229 | 502846 | 428139 | 202 | 543711 | 43 | PD3 |
| platanus.dbg2olc.pbfilt.ont2d | 1309 | 304009150 | 2364200 | 1000 | 232245 | 104763 | 519111 | 176 | 652333 | 557461 | 153 | 708268 | 44 | PD4 |
| platanus.dbg2olc.pbfilt | 1479 | 302538025 | 2204874 | 2336 | 204556 | 113907 | 405052 | 225 | 515659 | 433416 | 198 | 557166 | 45 | PD5 |
| smartdenovo.pball.ont2d | 713 | 292871890 | 3556032 | 13027 | 410760 | 199388 | 928894 | 97 | 1121622 | 956687 | 90 | 1173184 | 46 | S1 |
| smartdenovo.pball.ontmol | 698 | 292525170 | 3556036 | 15626 | 419091 | 208056 | 932578 | 94 | 1143652 | 959911 | 88 | 1194811 | 47 | S2 |
| smartdenovo.pball | 638 | 291784268 | 7505804 | 7277 | 457342 | 177734 | 1098245 | 67 | 1689097 | 1146154 | 62 | 1760185 | 48 | S3 |
| smartdenovo.pbfilt.ont2d | 713 | 292871890 | 3556032 | 13027 | 410760 | 199388 | 928894 | 97 | 1121622 | 956687 | 90 | 1173184 | 49 | S4 |
| smartdenovo.pbfilt | 633 | 291543073 | 7505612 | 11967 | 460574 | 184935 | 1083964 | 66 | 1708535 | 1140563 | 61 | 1778969 | 50 | S5 |

**Supplemental Table S4: Long read assembly size statistics**
The naming scheme is approximately "assembler.[parameters].pacbio-input.[minion-input]" where items inside square brackets are optionally present. The assemblies are presented in the same order that their ranks are visualized in Figure 2E. The "Matrix Row" and "Nickname" columns additionally help connect the table with Figure 2E. The assemblies chosen for scaffolding are in red and green, the latter being the assemblies chosen to move forward with after scaffolding. The Canu assembly in green (C3.2) is the assembly that went on to be chosen as the first released draft genome sequence.

**Supplemental Table S5: Pairwise comparisons of size statistics of hybrid scaffolds from pair of Canu or Falcon assemblies**

| | Canu | | Falcon | |
|---|---|---|---|---|
| **Version** | 1 (C3.2) | 2 (C3.1) | 1 (F8) | 2 (F9) |
| **Scaffold Set** | 1 | 2 | 3 | 4 |
| **PacBio** | All | All | All | All |
| **MinION** | 2D | Molecule | 2D | Molecule |
| **Number of sequences** | 857 | 847 | 603 | 608 |
| **Total length** | 311,020,781 | 311,186,377 | 302,764,727 | 303,252,712 |
| **Max sequence length** | 15,214,119 | 15,220,704 | 21,064,549 | 28,442,430 |
| **Min sequence length** | 671 | 793 | 117 | 119 |
| **Mean sequence length** | 362,918.1 | 367,398.3 | 502,097.4 | 498,770.9 |
| **Median sequence length** | 21,642 | 21,891 | 16,349 | 15,738 |
| **Sequence length N50** | 5,861,178 | 5,860,508 | 7,077,677 | 9,560,626 |
| **Sequence length L50** | 16 | 16 | 12 | 12 |
| **Expected sequence length** | 6,349,017 | 6,378,079 | 8,115,036 | 9,440,067 |
| **Sequence length NG50** | 6,689,144 | 6,689,144 | 7,509,349 | 10,026,878 |
| **Sequence length LG50** | 14 | 14 | 12 | 11 |
| **Normalized expected sequence length** | 7,052,416 | 7,088,469 | 8,774,810 | 10,224,021 |

#### Supplemental Table S6: Size statistics of Canu C3.2 across the work flow

### Canu

| | Assembly | Hybrid Scaffolds | Meta-scaffolds | *Meta-scaffolds: Bacterial removed | **Primary assembly | Anchored |
|---|---|---|---|---|---|---|
| Number of sequences | 1,044 | 857 | 769 | 744 | 205 | 92 |
| Total length | 302,258,594 | 311,020,781 | 312,049,034 | 309,775,056 | 298,965,442 | 136,508,149 |
| Max sequence length | 12,317,152 | 15,214,119 | 23,039,227 | 23,039,227 | 23,039,227 | 15,234,923 |
| Min sequence length | 671 | 671 | 671 | 671 | 4264 | 8,393 |
| Mean sequence length | 289,519.7 | 362,918.1 | 405,785.5 | 416,364.3 | 1,458,368 | 1,483,784.2 |
| Median sequence length | 28,310.5 | 21,642 | 21,093 | 21,428 | 94,665 | 76,618 |
| Sequence length N50 | 1,908,286 | 5,861,178 | 6,689,181 | 6,790,317 | 6,790,317 | 6,666,656 |
| Sequence length L50 | 37 | 16 | 15 | 14 | 14 | 7 |
| Expected sequence length | 3,173,951 | 6,349,017 | 7,930,712 | 7,982,431 | 8,269,986 | 7,447,096 |
| Sequence length NG50 | 2,363,127 | 6,689,144 | 8,288,951 | 8,288,951 | 8,288,951 | 8,393 |
| Sequence length LG50 | 31 | 14 | 12 | 12 | 12 | 92 |
| Normalized expected sequence length | 3,426,264 | 7,052,416 | 8,838,469 | 8,831,279 | 8,830,144 | 3,630,676 |

*All sequences here deposited to NCBI. One short contig (< 1kb) was flagged for removal. Thus, one can expect 743 sequences.

**The primary assembly contains contigs such that bacterial, ‘remove tigs’, and ‘redundant contigs’ are removed and those retained are marked as KEEP by purge_haplotigs. When keeping all of the non-bacterial KEEP scaffolds from purge_haplotigs (Roach et al. 2018) there are 265 instead of 205. The difference is that purge_haplotigs retains 54 of the contigs originally marked for removal (max size of 27.4 kb, mean size of 12.5 kb, sum of ~673 kb) and retained 6 contigs our minimap2-alone approach marked as “redundant” (max size of 27.5 kb, mean size of 20.4 kb, and sum of ~122 kb).

#### Supplemental Table S7: Size statistics of Falcon F9 across the work flow

### Falcon

| | Assembly | Hybrid Scaffolds | Meta-scaffolds | Meta-scaffolds: Bacterial removed | *Primary assembly | Anchored |
|---|---|---|---|---|---|---|
| Number of sequences | 713 | 608 | 565 | 557 | 138 | 84 |
| Total length | 296,559,753 | 303,252,712 | 304,155,157 | 302,144,604 | 294,751,939 | 137,975,970 |
| Max sequence length | 20,165,452 | 28,442,430 | 28,458,505 | 28458505 | 28,458,505 | 28,458,505 |
| Min sequence length | 119 | 119 | 889 | 889 | 1,079 | 2,103 |
| Mean sequence length | 415,932.3 | 498,770.9 | 538,327.7 | 542,449.9 | 2,135,883.6 | 1,642,571.1 |
| Median sequence length | 18,659 | 15,738 | 17,238 | 17,226 | 187,754 | 29,360 |
| Sequence length N50 | 2,878,051 | 9,560,626 | 10,085,069 | 10,085,069 | 10,085,069 | 11,007,303 |
| Sequence length L50 | 24 | 12 | 11 | 11 | 11 | 5 |
| Expected sequence length | 5,050,162 | 9,440,067 | 10,090,295 | 10,152,132 | 10,405,732 | 12,395,557 |
| Sequence length NG50 | 3,454,019 | 10,026,878 | 10,522,735 | 10,522,735 | 10,522,735 | 2,103 |
| Sequence length LG50 | 21 | 11 | 10 | 10 | 10 | 84 |
| Normalized expected sequence length | 5,348,840 | 10,224,021 | 10,960,769 | 10,955,043 | 10,953,963 | 6,108,175 |

* The primary assembly contains contigs such that bacterial, ‘remove tigs’, and ‘redundant contigs’ are removed and those retained are marked as KEEP by purge_haplotigs. When keeping all of the non-bacterial KEEP scaffolds from purge_haplotigs there are 162 instead of 138. The difference is that purge_haplotigs retains 4 of the contigs originally marked for removal, which were associated-bubble contigs from Falcon (max size of 13.7 kb, mean size of 8.4 kb, sum of ~33.5 kb), and retained 20 contigs that our minimap2-alone approach marked as “redundant” (max size of 20.2 kb, mean size of 9.4 kb, and sum of ~187 kb).

**Supplementary Table 8 A-C: Gap size statistics**

**A. Gap size statistics before and after gap-filling**:

| | Canu (C3.2) | | Falcon (F9) | |
|---|---|---|---|---|
| | **Before** | **After** | **Before** | **After** |
| **Number of gaps** | 187 | 156 | 105 | 91 |
| **Total gap length** | 8,761,718 | 7,789,542 | 6,692,707 | 5,628,294 |
| **Max gap size** | 677,224 | 674,379 | 965,004 | 953,978 |
| **Min gap size** | 500 | 25 | 500 | 25 |
| **Mean gap size** | 46,854.1 | 499,323 | 63,740.1 | 61,849.4 |
| **Median gap size** | 20,800 | 21,425 | 30,519 | 24,983 |
| **Gap N50** | 102,377 | 119,644 | 144,312 | 124,939 |
| **Gap L50** | 22 | 19 | 11 | 8 |
| **Expected gap size** | 167,075 | 176,849 | 288,100 | 314,580 |

**B. Gap filling progress by stage for Canu (C3.2)**:

**CANU**

| **Stage** | **Initial** | **Q1** | **PBJ1** | **Q2** | **PBJ2** | **PBJScaff** | **Q3** | **Pilon** |
|---|---|---|---|---|---|---|---|---|
| **Number** | 187 | 186 | 166 | 165 | 162 | 157 | 156 | 156 |
| **Total Length** | 8,761,718 | 8,761,218 | 7,952,958 | 7,952,624 | 7,819,479 | 7,789,567 | 7,789,542 | 7,789,542 |
| **N Diff** | - | 1 | 20 | 1 | 3 | 5 | 1 | 0 |
| **Len Diff** | - | 500 | 808,260 | 334 | 133,145 | 29,912 | 25 | 0 |

31 gaps closed totally. 972,176 bp filled overall.

**C. Gap filling progress by stage for Falcon (F9)**:

**FALCON**

| **Stage** | **Initial** | **Q1** | **PBJ1** | **Q2** | **PBJ2** | **PBJScaff** | **Q3** | **Pilon** |
|---|---|---|---|---|---|---|---|---|
| **Number** | 105 | 104 | 97 | 95 | 93 | 92 | 91 | 91 |
| **Total Length** | 6,692,707 | 6,692,207 | 5,778,078 | 5,777,035 | 5,669,089 | 5,628,485 | 5,628,294 | 5,628,294 |
| **N Diff** | - | 1 | 7 | 2 | 1 | 1 | 1 | 0 |
| **Len Diff** | - | 500 | 914,129 | 1,043 | 107,946 | 40,604 | 191 | 0 |

14 gaps closed totally. 1,064,413 bp filled overall.

#### Supplemental Table S9: Bacterial contig statistics in each assembly

| Bacterial contig statistics | Canu | Falcon |
|---|---|---|
| Number of contigs | 25 | 8 |
| Total length | 2,273,978 | 2,010,553 |
| Max contig size | 1,348,403 | 1,048,146 |
| Min contig size | 4,017 | 7,239 |
| Mean contig size | 90,959.12 | 251,319.125 |
| Median contig size | 13,445 | 71,061.5 |
| Contig N50 | 1,348,403 | 1,048,146 |
| Contig L50 | 1 | 1 |
| Expected contig size | 885,299 | 797,499 |

**Supplemental Table S10: *Sciara* (*Bradysia*) *coprophila* repeat family classes from RepeatModeler**

| Class | From Canu | From Falcon |
|---|---|---|
| **Unclassified** | 2267 | 2254 |
| SINEs | 15 | 19 |
| LINEs | 186 | 160 |
| LTR | 53 | 48 |
| DNA elements | 131 | 130 |
| rRNA | 2 | 2 |
| Satellites | 4 | 5 |
| Simple Repeats | 22 | 25 |
| RC | 14 | 16 |
| Buffer | 1 | 1 |
| snRNA | 0 | 1 |
| **TOTAL** | **2695** | **2661** |

**Supplemental Table S11: Sub-classification of classified Repeat Families found in *Sciara coprophila* genome with RepeatModeler**

| Class | Sub-class | Canu | Falcon |
|---|---|---|---|
| **LINE** | CRE | 36 | 30 |
| **LINE** | I-Jockey | 31 | 23 |
| **LINE** | CR1 | 26 | 25 |
| **DNA** | hAT-Ac | 25 | 22 |
| **Simple repeat** | - | 22 | 25 |
| **LINE** | RTE-BovB | 22 | 12 |
| **LINE** | L2 | 20 | 18 |
| **LTR** | Gypsy | 19 | 21 |
| **LINE** | R1-LOA | 19 | 10 |
| **DNA** | Zator | 16 | 17 |
| **LINE** | Penelope | 15 | 16 |
| **DNA** | - | 15 | 15 |
| **DNA** | Crypton-I | 15 | 12 |
| **LTR** | Pao | 14 | 11 |
| **RC** | Helitron | 13 | 16 |
| **LTR** | Copia | 12 | 13 |
| **DNA** | CMC-EnSpm | 9 | 9 |
| **DNA** | hAT | 7 | 3 |
| **DNA** | MULE-MuDR | 6 | 10 |
| **DNA** | P | 6 | 1 |
| **Satellite** | - | 4 | 5 |
| **SINE** | tRNA-V | 4 | 5 |
| **DNA** | hAT-Tag1 | 4 | 3 |
| **LINE** | R1 | 4 | 3 |
| **SINE?** | - | 4 | 2 |
| **LINE** | RTE-RTE | 3 | 6 |
| **LINE** | R2 | 3 | 5 |
| **DNA** | Maverick | 3 | 4 |
| **DNA** | hAT-Tip100 | 3 | 3 |
| **LTR** | - | 3 | 2 |
| **SINE** | tRNA | 2 | 6 |
| **DNA** | hAT-Blackjack | 2 | 5 |
| **DNA** | hAT-hATx | 2 | 3 |
| **DNA** | CMC-Chapaev | 2 | 2 |
| **rRNA** | - | 2 | 2 |
| **DNA** | Kolobok-Hydra | 2 | 1 |
| **LTR** | ERV4 | 2 | 1 |
| **SINE** | tRNA-RTE | 2 | 1 |
| **LTR** | ERV1 | 2 | 0 |
| **DNA** | PIF-Harbinger | 1 | 3 |

| | | | |
|---|---|---|---|
| **LINE** | - | 1 | 3 |
| **DNA** | MULE | 1 | 2 |
| **DNA** | Sola-2 | 1 | 2 |
| **DNA** | TcMar-Tc4 | 1 | 2 |
| **LINE** | I | 1 | 2 |
| **LINE** | L1-Tx1 | 1 | 2 |
| **SINE** | 5S-Deu | 1 | 2 |
| **buffer** | - | 1 | 1 |
| **DNA** | Crypton-H | 1 | 1 |
| **DNA** | Crypton-V | 1 | 1 |
| **DNA** | Ginger-2 | 1 | 1 |
| **DNA** | PiggyBac-X | 1 | 1 |
| **DNA** | TcMar | 1 | 1 |
| **LINE** | CR1-Zenon | 1 | 1 |
| **LINE** | RTE-ORTE | 1 | 1 |
| **DNA** | Dada | 1 | 0 |
| **DNA** | hAT-hATm | 1 | 0 |
| **DNA** | hAT-hobo | 1 | 0 |
| **DNA** | Kolobok-T2 | 1 | 0 |
| **DNA** | P-Fungi | 1 | 0 |
| **LINE** | Dualen | 1 | 0 |
| **LINE** | RTE | 1 | 0 |
| **LTR** | ERVK | 1 | 0 |
| **RC** | Helitron-2 | 1 | 0 |
| **SINE** | tRNA-Deu-L2 | 1 | 0 |
| **SINE** | tRNA-V-RTE | 1 | 0 |
| **DNA** | Novosib | 0 | 2 |
| **LINE** | L1 | 0 | 2 |
| **SINE** | tRNA-Meta | 0 | 2 |
| **DNA** | hAT-hAT5 | 0 | 1 |
| **DNA** | MULE-NOF | 0 | 1 |
| **DNA** | Sola-1 | 0 | 1 |
| **DNA** | TcMar-Tc2 | 0 | 1 |
| **LINE** | Dong-R4 | 0 | 1 |
| **SINE** | tRNA-Deu-RTE | 0 | 1 |
| **snRNA** | - | 0 | 1 |
| **TOTAL CLASSIFIED** | - | 428 | 407 |

*The Canu and Falcon columns represent the number of repeat families with the given classification found in the final Canu and Falcon assemblies respectively.

#### Supplemental Table S12 A-B: Repeat Masking on Canu

**Supplemental Table S12A: Summary of Repeat Masking on Canu**

| Canu: | All Scaffolds | Primary Scaffolds |
|---|---|---|
| **Sequences:** | 769 | 205 |
| **Total Length:** | 312,049,034 | 298,965,442 |
| **Percent GC:** | 35.56 | 35.56 |
| **Bases masked:** | 126,126,531 | 118,515,350 |
| **Percent of assembly masked:** | 40.42 | 39.64 |

Note: Tables describe masking of both primary and associated contigs. The associated sequences that were filtered out during assembly pruning were enriched for repeats containing 7,611,181 masked bases of 13,083,592 bases total (58.17% masked).

**Supplemental Table S12B: RepeatMasker classifications from Canu**

| Class | Sub-class | Number of elements | Length occupied (bp) | Percent of assembly |
|---|---|---|---|---|
| **SINEs:** | - | 6,607 | 1,244,312 | 0.4 |
| | ALUs | 0 | 0 | 0 |
| | MIRs | 0 | 0 | 0 |
| **LINEs:** | - | 21,636 | 9,813,413 | 3.14 |
| | LINE1 | 340 | 55,161 | 0.02 |
| | LINE2 | 2,394 | 1,168,917 | 0.37 |
| | L3/CR1 | 2,581 | 1,433,910 | 0.46 |
| **LTR elements:** | - | 11,150 | 4,668,386 | 1.5 |
| | ERVL | 0 | 0 | 0 |
| | ERVL-MaLRs | 0 | 0 | 0 |
| | ERV_classI | 47 | 3,734 | 0.001 |
| | ERV_classII | 572 | 85,008 | 0.03 |
| **DNA elements:** | - | 5,0957 | 11,011,885 | 3.53 |
| | hAT-Charlie | 15 | 849 | 0.0003 |
| | TcMar-Tigger | 12 | 1,339 | 0.0004 |
| **Unclassified:** | - | 523,519 | 96,659,741 | 30.98 |
| **Total interspersed repeats:** | - | | 123,397,737 | 39.54 |
| **Small RNA:** | - | 1,347 | 451,713 | 0.14 |
| **Satellites:** | - | 1,722 | 227,960 | 0.07 |
| **Simple repeats:** | - | 50,235 | 3,051,062 | 0.98 |
| **Low complexity:** | - | 9,985 | 460,867 | 0.15 |

*Most repeats fragmented by insertions or deletions have been counted as one element.
RepeatMasker Combined Database: Dfam_Consensus-20181026, RepBase-20181026.
Run with rmblastn version 2.2.27+.
The query was compared to classified sequences in the Full Repeat Library constructed for *Sciara coprophila*.
Note: 76.6% of masked bases were unclassified.

#### Supplemental Table S13 A-B: Repeat Masking on Falcon

##### Supplemental Table 13A: Summary of Repeat Masking on Falcon

| Falcon: | All scaffolds | Primary scaffolds |
|---|---|---|
| **Sequences:** | 565 | 138 |
| **Total Length:** | 304,155,157 | 294,751,939 |
| **Percent GC:** | 35.54 | 35.56 |
| **Bases masked:** | 121,347,624 | 116,828,462 |
| **Percent of assembly masked:** | 39.9 | 39.64 |

Note: Tables describe masking of both primary and associated contigs. The associated sequences that were filtered out during assembly pruning were enriched for repeats containing 4,519,162 masked bases of 9,403,218 bases total (48.06%).

##### Supplemental Table 13B: RepeatMasker classifications from Falcon

| Class | Sub-class | Number of elements | Length occupied (bp) | Percent of assembly |
|---|---|---|---|---|
| **SINEs:** | - | 6,565 | 1,239,700 | 0.41 |
| | ALUs | 0 | 0 | 0 |
| | MIRs | 0 | 0 | 0 |
| **LINEs:** | - | 21,297 | 9,424,389 | 3.1 |
| | LINE1 | 338 | 55,260 | 0.02 |
| | LINE2 | 2,389 | 1,160,831 | 0.38 |
| | L3/CR1 | 2,556 | 1,401,689 | 0.46 |
| **LTR elements:** | - | 10,976 | 4,379,978 | 1.44 |
| | ERVL | 0 | 0 | 0 |
| | ERVL-MaLRs | 0 | 0 | 0 |
| | ERV_classI | 45 | 3,474 | 0.001 |
| | ERV_classII | 563 | 84,866 | 0.03 |
| **DNA elements:** | - | 50,340 | 10,123,256 | 3.33 |
| | hAT-Charlie | 13 | 717 | 0.0002 |
| | TcMar-Tigger | 12 | 1,325 | 0.0004 |
| **Unclassified:** | - | 520,383 | 93,317,154 | 30.68 |
| **Total interspersed repeats:** | - | - | 118,484,477 | 38.96 |
| **Small RNA:** | - | 1,372 | 666,998 | 0.22 |
| **Satellites:** | - | 1,733 | 228,935 | 0.08 |
| **Simple repeats:** | - | 49,811 | 2,930,470 | 0.96 |
| **Low complexity:** | - | 10,008 | 460,501 | 0.15 |

*Most repeats fragmented by insertions or deletions have been counted as one element.
RepeatMasker Combined Database: Dfam_Consensus-20181026, RepBase-20181026.
Run with rmblastn version 2.2.27+.
The query was compared to classified sequences in the Full Repeat Library constructed for *Sciara coprophila*.
Note: 76.9% of masked bases were unclassified.

#### Supplemental Table S14: Transcriptome Evaluations

| Assembler | StringTie | | | | | | Trinity |
|---|---|---|---|---|---|---|---|
| Reference genome | Canu | | | Falcon | | | N/A |
| Approach | 1 | 2 | **3** | 1 | 2 | **3** | ***De novo*** |
| Number Genes | 19248 | 22520 | **22875** | 19308 | 22574 | **22958** | **75585** |
| Median NTPG* | 1 | 1 | **1** | 1 | 1 | **1** | **1** |
| Max NTPG | 95 | 54 | **137** | 96 | 59 | **148** | **22** |
| Number Transcripts | 36469 | 34167 | **44987** | 36478 | 34307 | **45099** | **93211** |
| Median TL | 2081 | 1721 | **2118** | 2081 | 1715 | **2118** | **493** |
| Max TL | 64293 | 62815 | **64287** | 59877 | 59879 | **59897** | **33554** |
| BUSCO: C | 2679 | 2693 | **2705** | 2677 | 2693 | **2703** | **2694** |
| BUSCO: F | 45 | 41 | **40** | 44 | 38 | **38** | **53** |
| BUSCO: M | 75 | 65 | **54** | 78 | 68 | **58** | **52** |
| TransRate pRef w/ CRBB (UniProt) | 0.24944 | 0.25016 | **0.25234** | 0.24952 | 0.25089 | **0.2521** | **0.26451** |
| TransRate Reference Coverage (UniProt/ Arthopod) | 0.36455 | 0.36275 | **0.36898** | 0.36434 | 0.3622 | **0.36819** | **0.37041** |
| TransRate pRef w/ CRBB (all) | 0.0883 | 0.08576 | **0.0919** | 0.08781 | 0.08584 | **0.09188** | **0.09598** |
| TransRate Reference Coverage (all) | 0.07644 | 0.07122 | **0.07982** | 0.07609 | 0.07164 | **0.07986** | **0.07322** |
| TransRate Score | 0.05376 | 0.08375 | **0.0368** | 0.05247 | 0.08278 | **0.03599** | **0.08182** |
| TransRate Optimal Score | 0.2522 | 0.28717 | **0.21469** | 0.25012 | 0.28662 | **0.21232** | **0.29233** |
| RSEM-Eval Score ($x10^9$) | -5.83677 | -5.95996 | **-5.73757** | -5.84391 | -5.94631 | **-5.75274** | **-7.02515** |

*NTPG Number Transcripts Per Gene; TL = transcript length

**Supplemental Table S14: Transcriptome Evaluations**
This table contains results summarized in Figure 5F “Reference-guided transcriptomes” category in the bold columns under Canu and Falcon (Approach 3, emboldened). There were three reference-guided transcriptome assembly approaches: (1) independent assemblies of each RNA-seq sample, followed by assembly merging, (2) read merging, followed by assembly, and (3) assembly merging the independent assemblies with the merged read assembly. The order from least to most sensitive for both assemblies was approach 1, then 2, then 3. This is unsurprising since approach 2 effectively has higher read depth for the assembly step, and since approach 3 merges outputs of approaches 1 and 2. For example, approach 3 had the highest number of complete BUSCOs detected and the fewest missing. Similarly, in the TransRate analysis, approach 3 had the highest proportion of reference proteins with Conditional Reciprocal Best BLAST (CRBB) hits and highest coverage across the reference proteins. Approach 3 also received the best RSEM-Eval scores, but did not fare as well as the other approaches with TransRate scores. Nevertheless, since approach 3 seemed to fare best in most metrics, we chose to include it as part of the EST evidence in the Maker2 gene annotation steps below.

#### Supplemental Table S15 A-B: Maker Annotation Transcript Evaluations on Canu and Falcon

##### Supplemental Table S15A: Canu Maker Annotation Transcript Evaluations

| | Canu Annotation Rounds | | | | | Alternative Round 3 | | | Final Canu Annotation |
|---|---|---|---|---|---|---|---|---|---|
| **Round** | 1 | 2 | 3 | | | Alternative Round 3 | | | 3* |
| **Repeat Library** | Original | Original | Original | | | Alternative/Filtered | | | Both* |
| **KeepPred =** | 0 | 0 | 1 | | | 1 | | | 1 |
| **Set** | NA | NA | Default | Standard | Max | Default | Standard | Max | Standard* |
| **# genes** | 15156 | 18250 | 21388 | 22330 | 33044 | 18845 | 21566 | 35492 | 23117 |
| **# transcripts** | 22128 | 22866 | 27003 | 27954 | 38713 | 25250 | 28007 | 42120 | 28870 |
| **Median NTPG** | 1 | 1 | 1 | 1 | 1 | 1 | 1 | 1 | 1 |
| **Max NTPG** | 19 | 19 | 19 | 19 | 19 | 16 | 16 | 16 | 19 |
| **Median NEPG** | 5 | 4 | 4 | 4 | 4 | 4 | 4 | 4 | 4 |
| **Max NEPG** | 75 | 131 | 132 | 132 | 132 | 129 | 129 | 129 | 132 |
| **Min GL** | 18 | 69 | 69 | 69 | 69 | 69 | 69 | 69 | 69 |
| **Median GL** | 2524 | 2677.5 | 2682 | 2660.5 | 2720 | 2912 | 2809 | 2698.5 | 2630 |
| **Max GL** | 406075 | 734151 | 545840 | 545840 | 545840 | 916260 | 916260 | 916260 | 545840 |
| **Median TL** | 1647 | 1553 | 1398 | 1383 | 900 | 1371 | 1332 | 761 | 1330.5 |
| **Max TL** | 31218 | 122902 | 542627 | 542627 | 542627 | 161972 | 161972 | 161972 | 542627 |
| **Median EL** | 212 | 192 | 185 | 185 | 158 | 173 | 173 | 147 | 182 |
| **Max EL** | 13073 | 122229 | 541964 | 541964 | 541964 | 161911 | 161911 | 161911 | 541964 |
| **Median IL** | 71 | 75 | 79 | 80 | 137 | 88 | 88 | 177 | 80 |
| **Max IL** | 218817 | 302637 | 218817 | 218817 | 273521 | 542594 | 542594 | 542594 | 218817 |
| **Median 5L** | 157 | 159 | 165 | 165 | 165 | 169 | 169 | 169 | 165 |
| **Median 3L** | 172 | 184 | 184 | 184 | 184 | 192 | 192 | 192 | 184 |
| **NG 5’ and 3’** | 12583 | 10131 | 10762 | 10762 | 10766 | 8100 | 8100 | 8103 | 10801 |
| **NG 5’ OR 3’** | 14315 | 12840 | 13193 | 13198 | 13213 | 10159 | 10175 | 10195 | 13335 |
| **BUSCO: C** | 2511 | 2601 | 2611 | 2638 | 2641 | 2265 | 2531 | 2562 | 2638 |
| **BUSCO: F** | 77 | 74 | 77 | 77 | 77 | 79 | 108 | 112 | 78 |
| **BUSCO: M** | 211 | 124 | 111 | 84 | 81 | 455 | 160 | 125 | 83 |
| **BUSCO: % Found** | 92.46 | 95.57 | 96.03 | 96.0 | 97.11 | 83.74 | 94.28 | 95.53 | 97.03 |
| **BUSCO: C (proteins)** | 2504 | 2585 | 2593 | 2620 | 2623 | 2248 | 2514 | 2544 | 2620 |
| **BUSCO: F (proteins)** | 81 | 74 | 78 | 78 | 78 | 81 | 111 | 115 | 79 |
| **BUSCO: M (proteins)** | 214 | 140 | 128 | 101 | 98 | 470 | 174 | 140 | 100 |
| **BUSCO: % Found (proteins)** | 92.35 | 95.0 | 95.43 | 96.39 | 96.50 | 83.21 | 93.78 | 95.0 | 96.43 |
| **TransRate pRef (UniProt)** | 0.23549 | 0.24017 | 0.24154 | 0.24396 | 0.24718 | 0.22034 | 0.24057 | 0.24573 | 0.2446 |
| **TransRate RefCov (UniProt)** | 0.34797 | 0.34658 | 0.34779 | 0.34993 | 0.35085 | 0.32425 | 0.34383 | 0.346 | 0.3503 |
| **TransRate pRef (all)** | 0.07887 | 0.07725 | 0.08012 | 0.08125 | 0.0851 | 0.07107 | 0.07846 | 0.08412 | 0.0816 |
| **TransRate RefCov (all)** | 0.06624 | 0.06359 | 0.0649 | 0.06543 | 0.06587 | 0.05922 | 0.06315 | 0.06425 | 0.06555 |
| **TransRate Score** | 0.13249 | 0.12217 | 0.092 | 0.08706 | 0.02364 | 0.06299 | 0.06603 | 0.03388 | 0.03659 |
| **TransRate Optimal Score** | 0.39399 | 0.38755 | 0.41209 | 0.4118 | 0.28501 | 0.34404 | 0.35499 | 0.36033 | 0.28293 |
| **RSEM-Eval Score ($x10^9$)** | -14.1476 | -12.8598 | -12.5512 | -12.3797 | -12.3468 | -16.4623 | -14.9077 | -14.5409 | -12.318 |

*See caption for S15A,B tables for a key to help interpretation of abbreviated metric names.
Note: The final Canu annotation here is Bcop_v1 that was deposited to the i5K workspace.

#### Supplemental Table 15B: Falcon Maker Annotation Transcript Evaluations

| | Falcon Annotation Rounds | | | | | Alternative Round 3 | | | Final Falcon Annotation |
|---|---|---|---|---|---|---|---|---|---|
| **Round** | 1 | 2 | 3 | | | Alternative Round 3 | | | 3* |
| **Repeat Library** | Original | Original | Original | | | Alternative/Filtered | | | Both* |
| **KeepPred =** | 0 | 0 | 1 | | | 1 | | | 1 |
| **Set** | NA | NA | Default | Standard | Max | Default | Standard | Max | Standard* |
| **# genes** | 15084 | 18060 | 19856 | 20596 | 26435 | 16452 | 19179 | 27473 | 21010 |
| **# transcripts** | 22028 | 22523 | 24036 | 24786 | 30653 | 20327 | 23088 | 31489 | 25225 |
| **Median NTPG** | 1 | 1 | 1 | 1 | 1 | 1 | 1 | 1 | 1 |
| **Max NTPG** | 19 | 15 | 14 | 14 | 14 | 16 | 16 | 16 | 14 |
| **Median NEPG** | 5 | 4 | 5 | 5 | 4 | 5 | 5 | 5 | 4 |
| **Max NEPG** | 63 | 124 | 159 | 159 | 159 | 242 | 242 | 242 | 159 |
| **Min GL** | 18 | 100 | 100 | 100 | 100 | 72 | 72 | 72 | 72 |
| **Median GL** | 2542 | 2770 | 2988 | 2982 | 3462 | 3441.5 | 3502 | 3981 | 2962.5 |
| **Max GL** | 406075 | 748256 | 548033 | 548033 | 548033 | 844972 | 844972 | 844972 | 548033 |
| **Median TL** | 1653 | 1560 | 1491 | 1475 | 1202 | 1506 | 1471 | 1089 | 1447 |
| **Max TL** | 31218 | 146821 | 543194 | 543194 | 543194 | 99085 | 99085 | 99085 | 543194 |
| **Median EL** | 212 | 190 | 181 | 180 | 159 | 172 | 170 | 145 | 179 |
| **Max EL** | 13073 | 146569 | 542688 | 542688 | 542688 | 98842 | 98842 | 98842 | 542688 |
| **Median IL** | 71 | 76 | 83 | 85 | 146 | 90 | 98 | 198 | 84 |
| **Max IL** | 223782 | 302656 | 255207 | 255207 | 255207 | 510038 | 510038 | 510038 | 255207 |
| **Median 5L** | 157 | 159 | 160 | 160 | 160 | 164 | 164 | 163 | 160 |
| **Median 3L** | 172 | 185 | 185 | 185 | 185 | 197 | 197 | 197 | 185 |
| **NG 5' and 3'** | 12549 | 10104 | 10083 | 10083 | 10084 | 7179 | 7179 | 7182 | 10119 |
| **NG 5' OR 3'** | 14262 | 12827 | 12895 | 12899 | 12912 | 9429 | 9438 | 9465 | 12971 |
| **BUSCO: C** | 2506 | 2584 | 2599 | 2630 | 2633 | 2176 | 2503 | 2543 | 2631 |
| **BUSCO: F** | 79 | 80 | 75 | 75 | 75 | 83 | 122 | 124 | 75 |
| **BUSCO: M** | 214 | 135 | 125 | 94 | 91 | 540 | 174 | 132 | 93 |
| **BUSCO: % Found** | 92.35 | 95.18 | 95.53 | 96.64 | 96.75 | 80.71 | 93.78 | 95.28 | 96.68 |
| **BUSCO: C (proteins)** | 2497 | 2565 | 2580 | 2611 | 2614 | 2161 | 2485 | 2526 | 2612 |
| **BUSCO: F (proteins)** | 85 | 83 | 79 | 79 | 79 | 87 | 127 | 128 | 79 |
| **BUSCO: M (proteins)** | 217 | 151 | 140 | 109 | 106 | 551 | 187 | 145 | 108 |
| **BUSCO: % Found (proteins)** | 92.25 | 94.61 | 95.0 | 96.11 | 96.21 | 80.3 | 93.32 | 94.82 | 96.14 |
| **TransRate pRef (UniProt)** | 0.23493 | 0.23864 | 0.24122 | 0.24371 | 0.24637 | 0.21236 | 0.23751 | 0.24307 | 0.24396 |
| **TransRate RefCov (UniProt)** | 0.34764 | 0.34576 | 0.34671 | 0.34864 | 0.34931 | 0.31321 | 0.33969 | 0.34237 | 0.34875 |
| **TransRate pRef (all)** | 0.07854 | 0.07728 | 0.07853 | 0.07959 | 0.08252 | 0.06678 | 0.07543 | 0.08035 | 0.07979 |
| **TransRate RefCov (all)** | 0.06596 | 0.06359 | 0.06359 | 0.06411 | 0.06451 | 0.05674 | 0.06166 | 0.06275 | 0.06421 |
| **TransRate Score** | 0.13294 | 0.12285 | 0.10809 | 0.10314 | 0.06423 | 0.08193 | 0.08601 | 0.04975 | 0.09869 |
| **TransRate Optimal Score** | 0.39056 | 0.39052 | 0.39057 | 0.39166 | 0.39179 | 0.32995 | 0.33836 | 0.34169 | 0.39171 |
| **RSEM-Eval Score (x$10^9$)** | -14.2099 | -13.0089 | -12.9233 | -12.7318 | -12.6894 | -17.1623 | -15.3031 | -14.9171 | -12.6757 |

*See caption for S15A,B tables for a key to help interpretation of abbreviated metric names.

**Supp Table 15 A and B: Canu and Falcon Maker2 Annotation Transcript Evaluations**

These tables contain results summarized in Figure 5F "Maker transcriptomes" category (Final Canu and Falcon annotation columns in A and B, respectively) as well as additional statistics. We evaluated the transcript outputs of the Maker2 annotation process. For both genome assemblies, the gene annotation improved across subsequent rounds. For example, there were more complete BUSCOs and fewer missing BUSCOs in each subsequent round of Maker. Similarly, a Transrate completeness metric was highest after the third round. RSEM-eval scores improved across Maker rounds, and for the Canu assembly, the TransRate Optimal score was highest after the third round. We also saw that the annotation edit distance (AED) scores from Maker2 reflected an improvement in round 3 over round 2 (not shown).

The transcript models learned from the Canu assembly contained more complete BUSCOs, fewer missing BUSCOs, more reference proteins with CRBB hits and higher reference protein coverage, higher TransRate optimal scores, and higher RSEM-Eval scores than those learned from Falcon. We also saw that in the final annotation sets from the Canu and Falcon assemblies there were more Canu models with the lowest annotation edit distances than Falcon models (Supplemental Figure S14). Moreover, the Canu annotation had more functionally-annotated genes (Supplemental Figure S14, Supplemental Table S16). Overall, this suggested Canu had a superior annotation.

Interestingly, the final set of Maker annotations for both assemblies had higher Transrate scores and higher Transrate optimal scores than the input transcriptome assemblies. In contrast, they had lower RSEM-Eval scores and lower completeness scores from both BUSCO and Transrate than the input transcriptomes. The final transcript models from Maker contained 94.2% and 94.0% complete Dipteran BUSCOs for Canu and Falcon, respectively (more when considering fragmented BUSCOs). For comparison, ~93.7% complete Dipteran BUSCOs were detected in both genome assemblies directly and ~96.6% were detected in StringTie transcriptome assemblies.

**Key**:
NTPG = Number Transcripts Per Gene;
NEPG = Number Exons Per Gene;
GL = Gene Length;
TL = Transcript Length;
EL = Exon Length;
IL = Intron Length;
5L = 5' UTR Length;
3L = 3' UTR Length;
NG 5' and 3' = Number of Genes that have both 5' and 3' UTRs annotated;
NG 5' or 3' = Number of Genes that have 5' and/or 3' UTRs annotated.
BUSCO C, F, M = Complete, Fragmented, Missing.
BUSCO: % Found = 100 * (C+F)/2799 = percent of Dipteran BUSCOs found as complete or fragmented.
TransRate pRef (reference used) = proportion of reference with CRBB.
TransRate RefCov (reference used) = Reference Coverage.
TransRate references used = UniProt (Arthropod proteins from UniProt/SwissProt); All (All proteins gathered for use in Maker2 annotation).

#### Supplemental Table S16: Additional characterization and comparisons of the final annotations of Canu and Falcon assemblies.

| | Final Canu Annotation | Final Falcon Annotation |
|---|---|---|
| Number of Genes with Ontology Term | 8671 | 8324 |
| Number of Genes with UniProt hit(s) | 13745 | 13283 |
| Number of Genes with Pfam domain | 13789 | 13248 |
| Number of Genes with All 3 above (intersect) | 8252 | 7933 |
| Number of Genes with $\geq$ 1 of 3 above (union) | 14961 | 14406 |
| Number of Genes with Drosophila hit(s) | 9739 | 9547 |
| Number of Genes with Anopheles hit(s) | 9057 | 8939 |
| Percent Drosophila Proteome with *Sciara* hit(s) | 87.3184 | 87.0166 |
| Percent Anopheles Proteome with *Sciara* hit(s) | 89.3487 | 89.3152 |

These are the results summarized in Figure 5F "Functional Annotations" category. The Maker2 annotation for the Canu assembly had more genes with GO terms, more genes with UniProt hits, and more genes with recognizable Pfam domains than found in the Maker2 Falcon annotation. The Canu annotation was also judged to be more complete by BUSCO (Supplemental Table S15), as well as through the number of hits back and forth with the proteomes of two other flies: *Drosophila melanogaster* and *Anopheles gambiae*. As with results reported elsewhere (e.g. Supp. Table S15), this suggests the Canu scaffolds had the superior annotation.

### Supplemental Table S17A: Putative Sciara homologs for proteins involved in reading, writing, and erasing DNA methylation marks for adenine and cytosine

| | | | | Analysis of putative Sciara homolog | | | | | | | | |
|---|---|---|---|---|---|---|---|---|---|---|---|---|
| Protein | Protein length (AA) | Possible Function | Putative Sciara homolog | Sciara protein length (AA) | Pfam domain | Inter Pro Annotation | GO term(s) | UniProt-SwissProt Functional Annotation: "Similar to:" | Drosophila melanogaster Top Hit | Homo sapiens Top Hit | Other top hit(s) [species] | Other Sciara proteins of interest |
| Dnmt2 | 329-369 | cytosine methylation | Bcop_v1_g020988-RA | 336 | C-5 cytosine-specific DNA methylase | C-5 cytosine methyltransferase | 0008168 | TRDMT1 tRNA (cytosine(38)-C(5))-methyltransferase (Homo sapiens OX=9606) | methyltransferase 2, isoform C | Chain A, DNA CYTOSINE METHYLTRANSFERA SE DNMT2 | . | . |
| TET1/2/3 and DMAD (most like DMAD and TET2) | 2136, 2002, 1811, 2860 | cytosine demethylation; adenine demethylation | Bcop_v1_g016381-RA | 1653 | Oxygenase domain of the 2OGFeDO superfamily | 2OGFeDO, oxygenase domain | . | TET2 Methylcytosine dioxygenase TET2 (Homo sapiens OX=9606) | Ten-Eleven translocation (TET) family protein, isoform D (also called: DNA N6-methyl adenine demethylase (DMAD) and Tet2) | Chain A, Methylcytosine Dioxygenase Tet2 | . | . |
| TET1/2/3 and DMAD (runner up) | 2136, 2002, 1811, 2860 | cytosine demethylation; adenine demethylation | Bcop_v1_g016384-RA/RB | 337-469 | CXXC zinc finger domain | Zinc finger, CXXC-type | 0003677, 0008270 | Tet DNA N6-methyl adenine demethylase (Drosophila melanogaster OX=7227) | Ten-Eleven translocation (TET) family protein, isoform A | Chain C, Methylcytosine dioxygenase (TET3) | TET1 splice variant VP_DE4 [H. sapiens], zf-CpG_bind_C domain-containing protein [Caenorhabditis elegans], methylcytosine dioxygenase TET3 isoform X2 [Mus musculus] | . |
| TDG | 410 | cytosine demethylation | Bcop_v1_g006678-RA | 1148 | Uracil DNA glycosylase superfamily | Uracil-DNA glycosylase-like | . | TDG G/T mismatch-specific thymine DNA glycosylase (Homo sapiens OX=9606) | G/T mismatch-specific thymine DNA glycosylase (AAD33588.1) | G/T mismatch-specific thymine DNA glycosylase (AAC50540.1) | . | . |
| MBD2 | 302-411 | methylated cytosine binding | Bcop_v1_g000093-RA | 185 | C-terminal domain of methyl-CpG binding protein 2 and 3; p55-binding region of Methyl-CpG-binding domain proteins MBD | Methyl-CpG-binding domain protein 2/3, p55-binding region; Methyl-CpG binding protein 2/3, C-terminal domain | . | Mbd2 Methyl-CpG-binding domain protein 2 (Mus musculus OX=10090) | Methyl-CpG binding domain protein-like, isoform B | methyl-CpG-binding domain protein 2 isoform 1 | . | Bcop_v1_g003486-RA, Bcop_v1_g003486-RB |
| MBD5 | 1494-1727 | methylated cytosine binding | Bcop_v1_g017396-RD/RE/RH/RI | 1394-1557 | PWWP domain | PWWP domain | . | Mbd5 Methyl-CpG-binding domain protein 5 (Mus musculus OX=10090) | six-banded, isoform H (Methyl-CpG binding domain, PWWP domain) | methyl-CpG-binding domain protein 5 isoform X1 (and PWWP domain) | . | . |
| MBD-R2 [D. melanogaster] | 1081-1169 | methylated cytosine binding | Bcop_v1_g019755-RA | 1193 | Jumonji domain-containing protein 2A Tudor domain; Methyl-CpG binding domain; THAP domain | Methyl-CpG DNA binding; THAP-type zinc finger | 0003677, 0005634; 0003676 | PHF20L1 PHD finger protein 20-like protein 1 (Homo sapiens OX=9606) | MBD-R2 | PHD finger protein 20-like protein 1 isoform 4 | . | . |
| CFP1 | 660 | Unmethylated CpG binding and recruitment of methylases | Bcop_v1_g003348-RA | 477 | PHD-finger;CpG binding protein zinc finger C terminal domain | Zinc finger, PHD-finger;CpG binding protein, C-terminal | . | Cfp1 CXXC-type zinc finger protein 1 (Drosophila melanogaster OX=7227) | CXXC finger protein 1, isoform B | CXXC-type zinc finger protein 1 isoform 1 | . | . |
| SETDB1 | 397-1292 | methylated cytosine binding | Bcop_v1_g019039-RA | 1325 | Histone methyltransferase Tudor domain; Methyl-CpG binding domain; SET domain; Histone methyltransferase Tudor domain 1 | Methyl-CpG DNA binding; SET domain | 0003677, 0005634; 0005515 | egg Histone-lysine N-methyltransferase eggless (Drosophila pseudoobscura pseudoobscura OX=46245) | eggless | histone-lysine N-methyltransferase SETDB1 isoform 4 | . | Bcop_v1_g019996-RA |
| SETDB2* | 707-719 | methylated cytosine binding | [Bcop_v1_g019996-RA]* | 1152 | Pre-SET motif; Histone methyltransferase Tudor domain; Methyl-CpG binding domain; SET domain; Histone methyltransferase Tudor domain 1 | Methyl-CpG DNA binding; Pre-SET domain; SET domain | 0003677, 0005634; 0005634, 0008270, 0018024, 0034968; 0005515 | egg Histone-lysine N-methyltransferase eggless (Drosophila pseudoobscura pseudoobscura OX=46245) | eggless | histone-lysine N-methyltransferase SETDB1 isoform 4 | . | Bcop_v1_g019039-RA |
| BAZ2B | 2132-2168 | methylated cytosine binding | Bcop_v1_g011218-RA/RB | 2486-2499 | DDT domain; Bromodomain; Methyl-CpG binding domain; PHD-finger; Williams-Beuren syndrome DDT (WSD), D-TOX E motif | Methyl-CpG DNA binding; Zinc finger, PHD-finger; DDT domain; Bromodomain; WHIM2 domain | 0003677, 0005634; 0005515 | BAZ2B Bromodomain adjacent to zinc finger domain protein 2B (Homo sapiens OX=9606) | toutatis, isoform F | bromodomain adjacent to zinc finger domain protein 2B isoform c (BAZ2B) | . | Bcop_v1_g015386-RA, Bcop_v1_g004566-RB |
| DAMT-1 and METTL4 | 365, 419 | adenine methylation | Bcop_v1_g005400-RA (DAMT-1/METTL4) | 367 | MT-A70 | MT-A70-like | . | METTL4 Methyltransferase-like protein 4 (Homo sapiens OX=9606) | CG14906 | N(6)-adenine-specific DNA methyltransferase METTL4 isoform X2 | DNA N6-methyl methyltransferase [Caenorhabditis elegans] | . |
| N6AMT1 | 186-214 | adenine methylation | Bcop_v1_g003680-RA | 233 | Methyltransferase small domain | Methyltransferase small domain | 0008168 | N6AMT1 Methyltransferase N6AMT1 (Homo sapiens OX=9606) | HemK methyltransferase 2 (N6AMT1) | Chain A, Methyltransferase N6AMT1 | . | Bcop_v1_g003124-RA/RB |
| ALKBH1 | 389 | adenine demethylation (6mA) | Bcop_v1_g006515-RA | 639 | 2OG-Fe(II) oxygenase superfamily | Alpha-ketoglutarate-dependent dioxygenase AlkB-like | . | Alkbh1 Nucleic acid dioxygenase ALKBH1 (Mus musculus OX=10090) | AlkB, isoform B | ALKBH1 | ALKBH1-like hits [Insecta] | . |
| ALKBH2 | 157-261 | adenine demethylation (1mA, 3mA) | Bcop_v1_g001386-RA | 230 | 2OG-Fe(II) oxygenase superfamily | Alpha-ketoglutarate-dependent dioxygenase AlkB-like | . | Alkbh2 DNA oxidative demethylase ALKBH2 (Mus musculus OX=10090) | *No significant hits. | DNA oxidative demethylase ALKBH2 isoform 1 | ALKBH2-like hits [Insecta] | . |
| ALKBH6 | 161-266 | adenine demethylation | Bcop_v1_g018977-RA | 241 | 2OG-Fe(II) oxygenase superfamily | Alpha-ketoglutarate-dependent dioxygenase AlkB-like | . | alkbh6 Alpha-ketoglutarate-dependent dioxygenase alkB homolog 6 (Danio rerio OX=7955) | CG6144, isoform B | alpha-ketoglutarate-dependent dioxygenase alkB homolog 6 isoform 3 | . | Bcop_v1_g000722-RA |
| jumu/jumeau | 719 | methylated adenine binding | Bcop_v1_g008508-RA/Bcop_v1_g008508-RC | 540-583 | Forkhead domain | Forkhead domain | 0003700, 0006355, 0043565 | foxn4 Forkhead box protein N4 (Danio rerio OX=7955) | jumeau | forkhead box protein N1 | . | Bcop_v1_g008508-RD |
| ALKBH8 | 614-664 | RNA methylation | Bcop_v1_g000722-RA | 602 | 2OG-Fe(II) oxygenase superfamily; Methyltransferase domain | Alpha-ketoglutarate-dependent dioxygenase AlkB-like;Methyltransferase type 11 | 0008168 | alkbh8 Alkylated DNA repair protein alkB homolog 8 (Xenopus tropicalis OX=8364) | CG17807 | alkylated DNA repair protein alkB homolog 8 isoform 1 | Fe2OG dioxygenase domain-containing protein [Caenorhabditis elegans] | Bcop_v1_g000775-RA |
| METTL3 | 221-580 | adenine methylation (RNA) | Bcop_v1_g006197-RA | 599 | MT-A70 | MT-A70-like | . | Mettl3 N6-adenosine-methyltransferase MT-A70-like protein (Drosophila melanogaster OX=7227) | methyltransferase like 3 | N6-adenosine-methyltransferase catalytic subunit (METTL3) | . | . |
| METTL14 | 456 | adenine methylation (RNA) | Bcop_v1_g016925-RA | 392 | MT-A70 | MT-A70-like | . | Mettl14 N6-adenosine-methyltransferase non-catalytic subunit (Drosophila melanogaster OX=7227) | methyltransferase like 14 | Chain B, N6-adenosine methyltransferase subunit METTL14 | . | . |

[Bcop_v1_g019996-RA]* may be another isoform of SETDB1 or SETDB2. Sequences for Sciara homologs reported below and available in the Bcop_v1 annotation at i5k.

#### Supplemental Table S17B: Other methylation-related proteins

| Protein | Length | Putative Function | Best Sciara Hit | Putative homolog for: |
|---|---|---|---|---|
| MECP2 | 486-498 | metylated cytosine binding | Bcop_v1_g019755-RA | MBD-R2 |
| MBD3 | 259-291 | methylated cytosine binding | Bcop_v1_g000093-RA | MBD2 |
| MBD4 | 262-580 | methylated cytosine binding, cytosine demethylation | Bcop_v1_g019755-RA | MBD-R2 |
| MBD6 | 1003 | methylated cytosine binding | Bcop_v1_g017396-RD/RE | MBD5 |
| MB3L1 | 194 | methylated cytosine binding | Bcop_v1_g000093-RA | MBD2 |
| NMAD-1 | 208-291 | adenine demethylation | Bcop_v1_g000722-RA | ALKBH8 |
| ALKBH3 | 614-664 | adenine/cytosine demethylation (1mA, 3mC) | Bcop_v1_g001386-RA | ALKBH2 |
| BAZ2A | 1873-1905 | methylated cytosine binding | Bcop_v1_g011218-RA/RB | BAZ2B |

These proteins had significant hits to Sciara proteins, but the Sciara proteins were best reciprocal hits with and putative homologs for the proteins named in column 5. The significant hits were likely due to similar methyl-binding domains and other shared domains.

More information on the methylation-related proteins in Tables S17A and S17B can be found in recent reviews (Armstrong et al. 2019; Long et al. 2013; Rausch et al. 2020; Kohli and Zhang 2013; Iyer et al. 2016; Zhu et al. 2018).

#### Supplemental Table S17C: FASTA sequences of Sciara proteins that were putative homologs

```
>Bcop_v1_g005400-RA, DAMT-1-like, METTL4-like
MSVVYSEKDCVLINHKEYIDHIYRTTIVNQGDSGSDLKTIYTLREDVFNFAPKNRNDTDDTKRHKINAKKSSILISQNPELHELNQTIAAKSKEFLEDCKRRNVIFNESSHHTTDSSLSVFKEVCSEET
VNVPLNGKNTLDNAIVTKISGQTYLIPPKCKFTNTCVSAIDKFADEFYDFIVIDPPWWNKYIRRVRTWKREEGYDMMYNDDILKIPLEDLIHPNTIVAIWCTNSPTHSEFVKKIALPKWNLKLLATYYW
VKITKSGDPICEFNAPLKKQPYEQLFLATHIESERLLKDDINLIFSISSVVHSHKPPLLEVFKDYLPDQPKCLEIFARYLLPHFSSIGREVLKLQNTILFEEKEQETVR
>Bcop_v1_g006197-RA, METTL3-like
MSDGTWDQIQEIKIKRNSLREKLEKRKKERQDILLGSTNLAAASSFIKSESSGSEDKKPVLSNIKHEIDADPEVERQLLQILSDNALTLPISSKQLAQRMNLLRFKPVNQDILYYFLQKLVAQSHISIN
NINSSTESGYEVTLVDHSRVQSLFNDIVEETEATVKSEDILKRKLDDDSDEGSGSKFSKNLSNTDKKVSLTKDTSNNNSDIMSLLSMPSTREKHNKQVGEEILELLSKPTAKERSLAEKFKSQGGAQVM
EFCPHGTRIECLRAQQATEELQSVKKADDQTVLIKLEESAEPVVVDSMKCNKLHFKKIIQSHTDESLGDCSFLNTCFHMDSCKYVHYEVDTLLTSEGCKADNNKNNRCIALPKLKTIDPAAKLDPPQWI
QCDLRFLDMTVLGKFAVIMADPPWDIHMELPYGTMSDDEMRQLGVPALQDDGLIFLWVTGRAMELGRECLKLWGYERFDELIWVKTNQLQRIIRTGRTGHWLNHGKEHCLVGTKGNPQNLNRGLDCDVI
VAEVRATSHKPDEIYGIIERLSPGTRKIELFGRPHNVQPNWITLGNQLDGIRLVDPELIAQFQKRYPDGNCMAPSQVAAKPPA
>Bcop_v1_g003348-RA, CFP1-like
MFKKSKRSKEEIAKEFDLPERKSKIATILKQGEQYCLCRSSDCSRFMIGCDGCEEWYHGDCIFVSEKEAKHIKHYYCQRCKENDPSLQTVFRAVPVAAVPSNPSDDRLPKKPKEKVRESDAKEGSKSKG
RCGNCDGCSNKYNCGRCDSCRSTGKKQRCDKRICINQRVKKEKPATSRTKKRRRSPSPEITNPELEGPRQCYGPGCRMTARPQSKYCSDLCGKKLASSRIYQVLPQRIQEWNLSPCIAEEENKVQLDNI
RAKKMTVRATLAQLDERQKELDLLVERAKRCVLDPNASETNEVEDEMSMYCVTCGFEIHSRTAIRHMEKCFNKYESQASFGSIFKTRIEGSSMFCDFYNPASKTYCKRLRVLCPEHCKKPKISDTDVCG
CPLVKNTFELTGEFCCAPKKKCFRHFFWEKIRRAEIDCERVRQWLKMDELMEQERQVRSSMASRAGVLHLMLHSTYNHELMEQLCRSGRQ
>Bcop_v1_g003680-RA, N6MAT1-like
MERIRFDQKGFEEYFNEVFQFIYRPDDDSKMLVDAVERLESLIFDHIKPAVCLEIGSGSGYVINSISKNFGSKIYKYYATDINPIAIETTLASGRANGVSNIHVVMTSLVDDIKKNLTHQVDLLVTNPP
FEPSPVEDVGRKGAICAWSAGKNGRAIIDRIIIELPDIMSDHGVALMCCVKENDVNDIIRRMGEQDFYSNIVIEREGTTSSNYNRIYQQYVIAFSKTNYWAEPY
>Bcop_v1_g006515-RA, ALKBH1-like
MFTDSFKYYKSKCPPPSFESVVNFSENDDNLFTRRTTIKAGEEFKHLGLTDPTHWHIYQFSSGLIFIKNPFTPHGQRYWIARCLRDYPKSPHANNLRSKSISPAAIDDWWTEMVSSRTKSDKESLRSAL
RWCTLGYHHQWDTKVYSDEMKHPFPVDLGTLSAYIARTLDFGEYSAEAAIVNFYPIGTTLAGHTDHSERNLDAPLFSFSFGQSAIFLIGGTTKDIKPTAMFLHSGDIVIMSKESRLCYHAVPRVFKSDD
SVWNNTIEIGDDDDRNEMPKKRKIENEAGYMVKDLWNDVDDRFYWEPFNEYVENCRINVNMFTDSFKYYKSKCPPPSFESVLFTRRTTIKTGEEFKHLGLTDPTHWHIYQFSSGLIFIKNPFTPHGQRY
WIARCLRDYPKSPHANNLRSKSISPAAIDDWWTEMVSSRTKSDKESLRSALRWCTLGYHHQWDTKVYSEEMKHPFPVDLGTLSAYIARTLDFGEYSAEAAIVNFYPIGTTLAGHTDHSERNFDAPLFSF
SFGQSAIFLIGGTTKDIKPTAMFLHSGDIVIMSKESRLCYHAVPRVVKSDDSVWNNTIEIGDDDDRNEMPKKRKIDNEAGYMVKDLWNDVDDRFYWEPFNEYVENCRINVNVRQVLRKGEHSL
>Bcop_v1_g006678-RA, TDG-like (TDG G/T mismatch-specific thymine DNA glycosylase)
MELSNNAMTSPMAATNVPYSPAPLQSDLSYSPDHSKLTAPSPASSQNATERPNDLSPKPNPDDQLVSNDHQQHSVRNFINHSNETENETDKSQVKFIEESQLKSTQPSPMDFQSNNQDDDNKQHELKAY
NPINKEASPTHTNHIEHDIKFDKNNEDKNVGEPYTPQYLPPPNNYNNAYVNKPNTTFKPYRMSNIPSNMKHEMSDDGYETSATGDGSHLTQNGSHMMGNQVLMPKCEISPPDDPYNFVDDDMQGSMSPH
LMPSSMQPHMRVPYSQSSLAGGGAGGNIVNNVPKLASMHGNMHAMSTVHSPVLSSPPIDGSTPKKRGRKKKQRDDTGQILNGVVKQIPAKERKKHDRFNGLSEEEVSKRTLPDHLANNLDIIIIGINPG
LFAAYKGHHYAGPGNHFWKCLYLSGLTHEQMNAEEDYKLLDFGIGFTNMVERATKGSADLTRKEIKEGCRLLQDKLRRFRPKIAVFNGKLIFEVFSGKKDFNFGRQPECIEGTNTYMWVMPSSSARCAQ
LPRAADKVPFYTALKKFRDYLNGLIPHLDESECVFSDPKFKACYEIEPKADNISIPSIQSATDITDLSNNHKMLDVPGKKKRGRPKKVRGLDGEEPKVTPRRTPSVVTLDSDGMPKKKRGRPKKVKTDA
INGMSNGMEAISNSIDAMCNTSIGMCSSVNGQNRMYPSPMQSPNPNNTFCGMNSMNPSAYGSMNSMALNPNRPNSYNHQQSPQNLHSQPFSHSDLSSEISAAISSEHLGESPVPTSPSLGPPDFEPPNN
CMPTDDNNTNECGFASPAPSTISEHTGPPLSQLHSNNNSTSYSPYRTTPSNNFIDPRSNDPNVNTTNESYALQAQQRQQQQNGLSNPMLRQTIPTNVDQHSPRHQQQQQQQQQTQQQQQQHSTTGDVY
KLPDVSAKSLSGLESLVDQIPSLNDSESPSSNNSLAGHSLNVSNLTGASVPISDSRTADVSSLSEQYQNSCIYSGYPSNLNSMPGPTSVNPSQIPQFSNSNYSPPNSSSPSNATNAYGSLSSSFSVSSL
TSSYPSSAMNTYHSNLMAASTHHLGGTGAPMYMEPHIPMAVNPLYHPYSQHASYTGYSPIAPPPTIHMPSPNYPYGYTNSTYGQPAHPTTHPAAGYLSNHHSMFDRIKPDIGYGGF
>Bcop_v1_g019755-RA, MBD-R2-like
MAPRRCTIKGCPSVSGRQQHRGVTFHTFPLNPIARQFWLTNCNLPDTKNITKSILVCSRHFRRADFQPLKNNKYLLKQGAVPTIFPWGTLPYVEPTPHSVIESGEPMVIADDKPAEPLPLVNQQLVDEI
VKIKSEMKATIGVKSPGKRSASVDEPGTSEPKKKAIRVSLDSKKLSNEPIATASFDPFSKFSPGARIEAQDFNGAWHTAKVVEVDGDEKEVLINFEKNSKSKAPATATEWIPMNSSRLRQVVKKPIASY
VPGEKCLARWTDSRKFPATIQKVLENNTYEVLFDDGFVKVVRAGHISKVKVPEPKTPKSDKSGRKSNPHLPIPKFDLAKLNLLPVPKDGEWCCNWVNEIPIGAEGYLDGPDGHKRPTVLVEDWRLPAGW
TKHLYQRSSVSGKWDVVLVGPTKKRFRSKNDVKVYLEEKGEVYNPDIYDFSIHKRRAKDIGLFVFTDEYKEELKNKQLLAASTAATAASSITPLLLNTSDDAFQGFSSKESSFLSSTLPSTPLSSIGSL
NASAIPSTPQSIPLLQPHPPPPPLPPTDPGYVYVGALKVQLIDNLFHCPEATCNKNFRKENHLQIHVKHYHLDLAKLLGVCPNMSDLAYQRTVGMPIDEAVPKNQIPNSQFFEKVYQSDLSSKLQRKSM
SPSTQVKAEPRSPNFSEKAANISAHSDEVDKPVAEPIAQSQEIANILHQSLSNEKNVEQIIHAQECIGSPKKPEAMPLDADVAIVKKSETPTARTKATKRRPIKPLNRQKARRNIKKAFMKSLQEAKAS
IVKDFGDANEGRVPLSSVSELSPTKSSRLKFLQPVDIGSSKTIAHDRQHVPEMNAPHPKYINENGEVIKIVRMRQEEIINCLCSYVEEDGLMIQCELCLCWQHGICNGIEKESQVPEKYICYICRNPQR
GRESMKYAHDQDWLLAGKLPVANYHPANSKHTERFDMLKQSHTLTGNLIELKRFMHSLNVKINIAENKDHPKMYLWSKKWEKCPTTAELTGIEMLAQSDKSKFVTNEALNELFSNTEVKKDAVKEEELN
TPMEVAADTSDPNEDKTATVPVKEEEAALTEEKPNVFGDDKMIVDQSILAGLLNSSGGVNVEQTQETVAVKIPQPEAAINPSECQLLLLDHIQKQQNLVMGRLQTIEAQIIALECRDDKMDIMEQPESC
SKTKQIISLLLNDLQKMRKIAAINSMENKLEI
>Bcop_v1_g019996-RA, SETDB1-like, SETDB2-like
MEHCEQTSEIMDIEKNAKLSKIIEDNSASNHSNRSDGWSDAITGGHHDVVSLLSDNEGEVMNSNGSADDNAESMGYGLNNEATWTLVEPTIVQRDSNDSDSGIVMPDNSVVNFREQSAERTQNSQKQEA
FNINCTDLQPNCESVTSNILNSLASNEFQHDAGSADDKTTRKKCINSECSSDKKKTTFYDTPIWSLNYFNVPRNYSRRQSVCQRCFDVSINDYERLGAALVNQQPLMLERLPIRPEVVEIIDSEEEDKG
GVGDTEALPSNTLTLLEENVDVVLQETVNRFGIEQQMRWANQILEHRVDVNSAISDRIDEEIRSMDQLRDSMYKNLYSRSRYAIEELPPFDLNTNKRLHMFGPNYPPCGKVVYPPVDKNSFYYAVRTKL
LSTWMPCKVTEYGDGPVDGKPMMFTVQFLRRNDDKTVPGNHLAYGTAPVVRLNVGTRVIALTMEKSVNVKCLNFHPGIVGETLQEYNRWRYMIFFDDGCVRYVTPDNVRLICSPSPNVWEDVSHDAVAA
AFIKSYLEQYKTERPMVQVKPGQKMLTQWRSEWKSARVANVDASLVHMHFEDVKRWEWIYRGSTRLGPLFKDNHQRQRRFAIAAKGNESFVELVSSDDDDESKNCRETTSPTVKSEDSSVPNTFTIPSQ
VDPERKQENRAIAKKGTGTMPSHPTVQHMNKATIYIDEDSPAKGKVVHYTVKQRILPRKFVSHICGTACLYPVKYNLSSYSPLVKPLLSGWDRKSVMNTRTSRRIVYQTPCGRIMRNMQELHRYLRDTK
CTSLDVDNFDFGGMTSCLAEYLVDNSFIQTKDLSNGCEAMTVSSVNFYDNTTPPPCEYSAQRIPTDGVPLNLDPDFLCGCDCDDDCWDKTKCQCWKLTLEGAKLISPYIPIDRMGYKYKRLLDPIPTGV
YECNAQCKCSKMCLNRVAQHPLQLKLQVFKTKSRGWGLRCLNDVPEGSFVCVYAGDLLTDKSANIAGGEYGDEYFAELDFIEVIESLKEGYEPEAMDVDSGRNSVHVSSETSIDVSKSLKVEPVFNTDD
DEIQVLNEPSLDENASSVCLLRSNSTLRFYDKNEMVFTMDAKKHGNIGRYINHSCDPNLFVQNVFVDTHDLRFPWVAFFAKRYIAAGTELTWDYNYVVGSVPGKRLFCECGAQNCRRRIL
>Bcop_v1_g008508-RA, jumu/jumeau-like
MNQFEEISTGFHDIFLNKFADSSGPILDFYVSDSMQDMLDIDIRSEVATVVGGVSEFTSLMNDLPSLDLESNSNDGDILWSSGQKWPTTNDIYADVGACINPNSVHTIAPVTQSSILMSPKAQLNLNTH
NYHSNNSPLPSPKEKKSHLTFSPNTIKVPVVQENKKQSLLCQPQRTSVEAIKKDLTDDLRDEKVVKLTHSMTVNGQTNGHNTIKLAAGIGGLSFANSQMYAKLKQNNSKIVANGNITHLKRESSPLNNS
SMIGNNGSLNGHSNGQKMFARSISVQNGKSHISTQHILPSNSLSHHNGVNKQMKPKITGQEFPKPAYSYSCLIAMALKNSRAGSLPVSEIYSFMCEHFPYFKTAPNGWKNSVRHNLSLNKCFEKIEKPV
TNGGQRKGCLWAMNPSKITKMDEEIQKWSRKDPMAIRKAMVFPDNLELLEKGEMKHGSTGDSEGENEVGDEDTETNTDSEQDVTELDDTIQNTTEPESCDEDDRPQIDYDIEVTDLYDDIDLNDANQKD
LLSPEDIHPAKRARLDINYSIGPAGTFQSLNGSQQPQQNRRKMTLVNRIVIRWDHNRKRGSVVCVEQ
>Bcop_v1_g008508-RC,  jumu/jumeau-like
MNQFEEISTGFHDIFLNKFADSSGPILDFYVSDSMQDMLDIDIRSEVATVVGGVSEFTSLMNDLPSLDLESNSNDGDILWSSGQKWPTTNDIYADVGACINPNSVHTIAPVTQSSILMSPKAQLNLNTH
NYHSNNSPLPSPKEKKSHLTFSPNTIKVPVVQENKKQSLLCQPQRTSVEAIKKDLTDDLRDEKVVKLTHSMTVNGQTNGHNTIKLAAGIGGLSFANSQMYAKLKQNNSKIVANGNITHLKRESSPLNNS
SMIGNNGSLNGHSNGQKMFARSISVQNGKSHISTQHILPSNSLSHHNGVNKQMKPKITGQEFPKPAYSYSCLIAMALKNSRAGSLPVSEIYSFMCEHFPYFKTAPNGWKNSVRHNLSLNKCFEKIEKPV
TNGGQRKGCLWAMNPSKITKMDEEIQKWSRKDPMAIRKAMVFPDNLELLEKGEMKHGSTGDSEGENEVGDEDTETNTDSEQDVTELDDTIQNTTEPESCDEDDRPQIDYDIEVTDLYDDIDLNDANQKD
LLSPEDIHPAKRARLDINYSIGPAGTFQSLNGSQQPQQNRRKMTLVNRIV
>Bcop_v1_g000093-RA, MBD2-like
MNVTIERKSDCAALPKGWQREEIMRKSGLSAGKVDVFYYRQTASIFKQPVTVYKTQDSKSKADLKHGTLDKPNQLFWAKRLEGLRACDVDGVELGPVELPKGLKPVGPNVDDGTLLQSVATSLHVTSQA
VTGQQASKAVLQMNPGVFLNPDQPLMHSVNINEDDIRRQEDRVQVARRKLQEALKC
>Bcop_v1_g000722-RA, ALKBH8-like
MTSKSIERKAGRKQKRCQHIITSETGIECNDAPKQYLVVCNAGLSTGLKNDVLLKEAAIFGTVLQIILLPGKSYSFMKCKTIDDAISIYNGMNARSTLGQNGSVLYLLYCEDVPDSVNPWNHQVPPGLI
LIDNFITELEEESLLNRVSDDFTGHDSSLGNSGSMKHRQVYHYGYEFKYDTNNVDATQPLAMQIPDECNRLWPRLRSSMPNLSFQVPDQITVNKYEPGQGIPPHCDTHSAFCDPIFSLSLASGTVMEFR
RPSDGRHISVWLPRRSLLVMSKESRYGWTHGITPRKTDVVPVNEGLTIVERKLRVSFTFRRLSFDKCRCEFTSLCDTYQKSQQETSENIEKLVLPAELELRNVHKVYDQIADHFSETRHSPWPKVKEFI
ESFPAGSIVVDVGCGNGKYLPLNDSIVKIGCDRSHGLLNVCNQRGFNIFQCDCLHLPIKSNSVDGCISIAVIHHLVTEERRQQAISEMSRILVTGGRCLIYVWAKNQAKGTKSSYLRQNKKNNKVDVVA
DDASSCNAKFCDGLPIHTNRTQFSHSDMLVPWKRKDETGDEQKTFLRYYHVFEEAELQRVCERVSDIAVIETYYDQGNWCVIFEKK
>Bcop_v1_g001386-RA, ALKBH2-like
MMSVIEQLNEIGNKPIETKNLKNYRRIVAEHLDVETAILFDKRLSYEIYKGLDDAVEYYEENSLKVKVFGKWHPVPRKMASLIPFAAYCDEGITYTFSGLTMPTRNWFPLLSQLRDIVTKLTGFDYNFV
LVNRYENGNQHIGEHRDGEKEIDPSSPIVSLSFGQHRDFVFRHSNSKARPDLTPIKMELPNGSILVMNPPTNQFWLHSLPIRKKALHPRINLTFRKIFRSE
>Bcop_v1_g016925-RA, METTL14-like
MGDVLKNLRIRSQKRKKLLAQTLGVSKVEDLKHVLGTAEDVQSTKTQRDDDEPTSSTKKFQSESMFYRDSSTFLKGTQSSNPHNDYCQHFVDTGQRPQNFIRDVGLADRFEEYPKLRELIRLKDELIEQ
TATPPMYLQADLKTLDLKSLGTKFDVILIEPPLEEYARAGAGVAVSSSTRTFWSWDEILSLDIGEVAAHRSFVFLWCGSSEGLDMGRNCLRKWGFRRCEDICWIRTNIENPGHSKILEPKAVFQRTKEH
CLMGIKGTVRRSTDFDFIHANVDIDLIISEEEQLGSFEKPIELFHIIEHFCLGRRRLHIFGRDTSIRPGWLTVGPELTNSNFNGELYASAFADNTTTGCTDRIEALRPKSPPTNGKMMRGRGRGGYVRG
RGRGR
>Bcop_v1_g011218-RB, BAZ2B-like
MDKENGDNNEKGKGGGHSGGGSGQDPSSLLDAASLFAYWGRDPTAAQAAAAASSQLFGSPFGAAAAGLGLLPGSGGSSNDRYSMNNHQHHQNTMAVAASQAASLAGLHPASWWSMAQLAAQDYFARLQA
SGLSPFQHPELAAFPGMGMGGNVGGSSSGTGRQGAGEGKSSTKGRKEKKSNNSNNSMGGSSGNASTTVTSSAGTSSAYKQSSSQSYKSAANQYSPATLHKELLAMQAAAAASGSGHSSSRKSSQSSHKQ
HQSSAHHTSSSGHGSMGNHSSGRDSHHGSNSGKESSSKNSSSALATSLASSNATLTMNALNSLSQFGNLGSLSSQHSMQAAMNALAGNSSSKSSKDYIPSGILSAALERNDPSSLLGVRLPPDTEIIKY
TSSIVGPKIPGTTNRGRKKTISLDSNPLLAGIPGLTAGTGVSPAKRARLEAEYGNMVQSAHQSDRDRVEVIKLPSTINSNGAYNLSKSNAQAKDTSFDSSNEWATGLNLSGKASTTSSTDDGDAPLNLC
MKSDGKSGDAGTGTTGSNSLQSLSHITAALGSGSGNDRMLHYKEGRPRNLGRGVSKPKKNTVASLLAQSRAVGLKPMLNTQQLLSQGADLEKIRQAIAEAHLAMEVSTDSESLADASGMSDSEGEEMVN
LAELRAPLDKGWKRETIIRGLTKNMQIRGDVCYYAPGSQMKLKHIGQIQTILEQQSSNFTTDNFSFSARPIIGSFLQPYSSATDAECIRMTDSEVAKRLEELKLYTKQSLNVEQRIEIARQQQALRDAK
KMAKDDMSRTKEKPVSYQRGKQNRDVEKVERLEAQRKERELKNQQALEERERKRQEALLQKEQIYLQDLNKQKELLYTVELERERRRQHMTLIRQLENRRKYEERERKKHQMVLDKLISREKKLMVKKR
DAEILAELRKPQEDSEIPNQKDLPELQRLPGLKLTGQALSDLLMVFEFLHNFGETLGFDMDSLPTLQSLHQALCSDGTATDAEDELISVMTHLLVCAIEDPGIPNPNRHTTLLGQSLRTADITNSNISE
ILRIYLYAVATGEVRQTSGLTFDRDRERRVADHHSTEQFDISTSKNQQYYEQLHENTTWKLSECLKENPFVSLNSTVKSQILAHLCNDLLLNKAVSKQIEGSLESMAQYKREKYLLDNRIRKYKHLHTR
KIRMEQFEKQQMLAKQSESIPKDGFVEPTRQGPVGDNLVNENNNETPDKLNESVSNSNLIRTASNLTNASSIDNLTDVSDLANHKDDSLHSTDVALTKNGDEMSPVKSVDDIKANKSLDDFSLIDTPRS
LNNGTSTPDMNSLLNKKIINRELNDVSSTADLNLDEDLSDLESEGTILEEDEDNRMTAEEVLKKLEKIMKAGQQNKMLLEQSCNQLRATCYGQDRFWRRYWYLPKAGGIFVEGLESAQPDILKYHSLDD
EVLSTSSHSVSVPVGENTEKKKRGRKRKHQTDEEESSSKNVTEDAQRVNNETKPTNEVKPVTELKPGDEVKATNDIKPTSDMHQNDDGMMDIEDSIPRAILVQKACTIDDVEMNSKISGVSSSVVDHVS
QTATNSTSNPICNQEEGSSNSGASSDVVVVADDTQIAKLENGAVKEEPATETDANPDPSKAGSEVAVKQEIKDEIKNEGDDDTDGVVKSETLMEKWFSIANRELQLTSTECTPTMACQASFSNVTCDNT
IQYQGNRWEIGNNAQYFNVPVEGLTSTLTFNRDSFLTLSGLDEDMMSQALNSTLKPKDDTLQDVTMVDDEIAKSTINKIDVKQEEMLVPFNLPSFINMSLGNISAYIQCDNPSPLQMTPDEQKLLEEVK
INGLPKRLERNLVPKDLRYGWWKIDDIEVIKEMIQCLHVRGVRERELRQHLLTALSENLDLTTQCHVAHVRAPPPSKGYIDPEPMNAWNPQIARRVELNLLDQVETLEDKIASASMQMKGWQVPNKDDV
```

ENGMDMIGGISLIRERILGLEAAIERRYLKPPLGTSTADAHIAAIATQNQNQANPPTSTSQAASSSTTPTSSSATSATTNSDQQANDTDQDMDREESPEPEVLPKALQSWREAVARSHTTSQLAMALYV
LESCVAWDKSIMKAPTQNGSNTQRSLQNCQFCSSGENEDKLLLCDGCDKGYHTYCFKPKMVNIPEGDWYCYECVNKATGDRKCIVCGGHRQQPVGKMIYCEICPRAYHHDCYIPPMIKVPRGKWYCHGC
ISKAPPPKKRGPKKGHKEKGEKSEKGEKSDKTEKADKADKSSNSTLTATPTLNSSGQEDGALSPTHSVASTSFEEHSTSVDSKPPSTTTAQTAPIISATTSADIVDEEMPIADDVSDSAAPSSPPHTSQ
TVASVQSPASSPPPSSPPPPPPPQAVQPTTAPITTAAAAMEELNHTAALLNEAKEKLKLEKKERHAAKKLIKELAACKTMLEAMELHEDSWPFLLPVNTKQFPTYKKIIKSPMDLSTIKKRIQDLQYKS
RDDFIADVRLIFDNCEIFNEDDSPVGKAGHGMRKFFEVRWAELTDKHS
>Bcop_v1_g011218-RA, BAZ2B-like
MDKENGDNNEKGKGGGHSGGGSGQDPSSLLDAASLFAYWGRDPTAAQAAAAASSQLFGSPFGAAAAGLGLLPGSGGSSNDRYSMNNHQHHQNTMAVAASQAASLAGLHPASWWSMAQLAAQDYFARLQA
SGLSPFQHPELAAFPGMGMGGNVGGSSSGTGRQGAGEGKSSTKGRKEKKSNNSNNSMGGSSGNASTTVTSSAGTSSAYKQSSSQSYKSAANQYSPATLHKELLAMQAAAAASGSGHSSSRKSSQSSHKQ
HQSSAHHTSSSGHGSMGNHSSGRDSHHGSNSGKESSSKNSSSALATSLASSNATLTMNALNSLSQFGNLGSLSSQHSMQAAMNALAGNSSSKSSKDYIPSGILSAALERNDPSSLLGVRLPPDTEIIKY
TSSIVGPKIPGTTNRGRKKTISLDSNPLLAGIPGLTAGTGVSPAKRARLEAEYGNMVQSAHQSDRDRVEVIKLPSTINSNGAYNLSKSNAQAKDTSFDSSNEWATGLNLSGKASTTSSTDDGDAPLNLC
MKSDGKSGDAGTGTTGSNSLQSLSHITAALGSGSGNDRMLHYKEGRPRNLGRGVSKPKKNTVASLLAQSRAVGLKPMLNTQQLLSQGADLEKIRQAIAEAHLAMEVSTDSESLADASGMSDSEGEEMVN
LAELRAPLDKGWKRETIIRGLTKNMQIRGDVCYYAPGSQMKLKHIGQIQTILEQQSSNFTTDNFSFSARPIIGSFLQPYSSATDAECIRMTDSEVAKRLEELKLYTKQSLNVEQRIEIARQQQALRDAK
KMAKDDMSRTKEKPVSYQRGKQNRDVEKVERLEAQRKERELKNQQALEERERKRQEALLQKEQIYLQDLNKQKELLYTVELERERRRQHMTLIRQLENRRKYEERERKKHQMVLDKLISREKKLMVKKR
DAEILAELRKPQEDSEIPNQKDLPELQRLPGLKLTGQALSDLLMVFEFLHNFGETLGFDMDSLPTLQSLHQALCSDGTATDAEDELISVMTHLLVCAIEDPGIPNPNRHTTLLGQSLRTADITNSNISE
ILRIYLYAVATGEVRQTSGLTFDRDRERRVADHHSTEQFDISTSKNQQYYEQLHENTTWKLSECLKENPFVSLNSTVKSQILAHLCNDLLLNKAVSKQIEGSLESMAQYKREKYLLDNRIRKYKHLHTR
KIRMEQFEKQQMLAKQSESIPKDGFVEPTRQGPVGDNLVNENNNETPDKLNESVSNSNLIRTASNLTNASSIDNLTDVSDLANHKDDSLHSTDVALTKNGDEMSPVKSVDDIKANKSLDDFSLIDTPRS
LNNGTSTPDMNSLLNKKIINRELNDVSSTADLNLDEDLSDLESEGTILEEDEDNRMTAEEVLKKLEKIMKAGQQNKMLLEQSCNQLRATCYGQDRFWRRYWYLPKAGGIFVEGLESAQPDILKYHSLDD
EVLSTSSHSVSVPVGENTEKKKRGRKRKHQTDEEESSSKNVTEDAQRVNNETKPTNEVKPVTELKPGDEVKATNDIKPTSDMHQNDDGMMDIEDSIPRAILVQKACTIDDVEMNSKISGVSSSVVDHVS
QTATNSTSNPICNQEEGSSNSGASSDVVVVADDTQIAKLENGAVKEEPATETDANPDPSKAGSEVAVKQEIKDEIKNEGDDDTDGVVKSETLMEKWFSIANRELQLTSTECTPTMACQASFSNVTCDNT
IQYQGNRWEIGNNAQYFNVPVEGLTSTLTFNRDSFLTLSGLDEDMMSQALNSTLKPKDDTLQDVTMVDDEIAKSTINKIDVKQEEMLVPFNLPSFINMSLGNISAYIQCDNPSPLQMTPDEQKLLEEVK
INGLPKRLERNLVPKDLRYGWWKIDDIEVIKEMIQCLHVRGVRERELRQHLLTALSENLDLTTQCHVAHVRAPPPSKGYIDPEPMNAWNPQIARRVELNLLDQVETLEDKIASASMQMKGWQVPNKDDV
ENGMDMIGGISLIRERILGLEAAIERRYLKPPLGTSTADAHIAAIATQNQNQANPPTSTSQAASSSTTPTSSSATSATTNSDQQANDTDQDMDREESPEPEVLPKALQSWREAVARSHTTSQLAMALYV
LESCVAWDKSIMKANCQFCSSGENEDKLLLCDGCDKGYHTYCFKPKMVNIPEGDWYCYECVNKATGDRKCIVCGGHRQQPVGKMIYCEICPRAYHHDCYIPPMIKVPRGKWYCHGCISKAPPPKKRGPK
KGHKEKGEKSEKGEKSDKTEKADKADKSSNSTLTATPTLNSSGQEDGALSPTHSVASTSFEEHSTSVDSKPPSTTTAQTAPIISATTSADIVDEEMPIADDVSDSAAPSSPPHTSQTVASVQSPASSPP
PSSPPPPPPPQAVQPTTAPITTAAAAMEELNHTAALLNEAKEKLKLEKKERHAAKKLIKELAACKTMLEAMELHEDSWPFLLPVNTKQFPTYKKIIKSPMDLSTIKKRIQDLQYKSRDDFIADVRLIFD
NCEIFNEDDSPVGKAGHGMRKFFEVRWAELTDKHS
>Bcop_v1_g017396-RE, MBD5-like
MAAASSGTLQIQGSVPSISQSHLHQQQQQSFQQRLSYVQRSGSQLNVVSGCGPVTVSAIQQRGPNVAPGWRRQVNNGEIIYISPSGAVLRNLGQIKEYLLSAGTCKCGLPCPLRPDYFFEFNSQVPNVK
LQIPLEDAGTIRTTSCLHQARLLGSNNLLPKDTSPSKKRKLDGAWSPASSISSSDSTRSSQTTALCNYVGSPIPNQMVIANPRVQPTIIQKRQVVGIPAQQHLPQQQSQTVVYQNQNVGENKVANVTVS
RTPPWRKNSIRPNFKEDPTGYLDHQTAILHSSILNVHSPDIAENSNQSTISNSNVSYQSEPKLSNQLNVSTPNNDSVQPHTIGSEQSLSVMQQHQHNQNVFIQQQQQQSGVESAGGKTQPHVIIGSNGQ
PIRILQNGQIIYSNPTMQSVKVQDTQNNASTGIVSSGPRIISDEIIQPVQTTNVHQQVFQQQNAPRMMAPQQIIPQGQMEPVQIQHNRQVLQQQPQQQASSGQTQIITVQQHVPTSQPFAMNQSVRQPH
PQSIVQQQAVNRSHQDHRIIMPSSNQTDSNQSMELSIGGQTVTHLPNGMVQVQQNCDIAHMKQQQQIQLQQQIQQQNMLFRQQQELQQNPRLKDQYQTCNSTQDTIRQQHQVSTRTVATSQESPVSSST
TYRNKIIEKRSSSKGPAQVGAISTSNESPPATSSCSSPADSPESLQARTVTSAAALCPPKIQTGGIVRSQTTGKNTITSVLAGKAMTSTTTTNHGFGKQSGNANEKDTAPNTFDRQIVHVSSSQMVQIP
SSVHQAKISTAGSNQNRPIASMIQNCVPVSAALTNLQLQPICRGEPGSQIIMTSSGQILVMPPQSKSSNNQMIISGGNGSSLVMNNSPQGVVINQQGQQILSGDMIHGINNQVLDGQSANILQNSGNQV
VIQGQNLMSGNGNRVIGSNNSNFIVNSTNNMQPMIINNSNIISHNGNVLQQGQNVIHGNQGNLISGTKVISSNGNILNGSNIISNQSNVLASNNQIIGNSNSTNLISPNGTVVLPNGYVLQPHQFTTVD
GQVVNVINQDNGTHQFIQSTQQRIILSPDSKRRAKKRKSSSGTPPTPHSLSPQQSPTIQQHSPTIVQQNQSGAMLQITPQYQQQSFQISPGMSGITLVQNKPQTNSGPQQQILLQNGQTIIQPLNIIGQ
QLLVPAGLMVAPDTTLLQIQNVGPCGSILTPQGMMIRASSPQNKNFLSPNSGGQQFIVSGNGQISPIGQMYSTPMGLVVPQTNSNNSGPTYVQQNTTILQQQTTMINSANGNGGSSNVDADIRSQNQQS
MSQSQQQQHHHHRTVSVSPPDTTTHSPRSPERPPSQRSNGSDTNMVQCVSSSEPDAVSPVADSARSPSNNSDYERSGGISYQQNTFKPSDAKIRRIQTPTQLSNSHVITQGNQSKMRPDSHVWGAARGH
PAWPGKIVSPPETYKGVATSSDSTWVQWFGGRPNVELVSINSLKSLSEGLEAHHKAQKDTRKSRKLNSQLECAIQEAMAELDRTVTATSSTPTTPSNRNKARTLKPTSTQQRTKPVKIAPAPPNSTTHL
PAKYLRQSK
>Bcop_v1_g017396-RD, MBD5-like
MAAASSGTLQIQGSVPSISQSHLHQQQQQSFQQRLSYVQRSGSQLNVVSGCGPVTVSAIQQRGPNVAPGWRRQVNNGEIIYISPSGAVLRNLGQIKEYLLSAGTCKCGLPCPLRPDYFFEFNSQVPNVK
LQIPLEDAGTIRTTSCLHQARLLGSNNLLPKDTSPSKKRKLDGAWSPASSISSSDSTRSSQTTALCNYVGSPIPNQMVIANPRVQPTIIQKRQVVGIPAQQHLPQQQSQTVVYQNQNVGENKVANVTVS
RTPPWRKNSIRPNFKEDPTGYLDHQTAILHSSILNVHSPDIAENSNQSTISNSNVSYQSEPKLSNQLNVSTPNNDSVQPHTIGSEQSLSVMQQHQHNQNVFIQQQQQQSGVESAGGKTQPHVIIGSNGQ
PIRILQNGQIIYSNPTMQSVKVQDTQNNASTGIVSSGPRIISDEIIQPVQTTNVHQQVFQQQNAPRMMAPQQIIPQGQMEPVQIQHNRQVLQQQPQQQASSGQTQIITVQQHVPTSQPFAMNQSVRQPH
PQSIVQQQAVNRSHQDHRIIMPSSNQTDSNQSMELSIGGQTVTHLPNGMVQVQQNCDIAHMKQQQQIQLQQQIQQQNMLFRQQQELQQNPRLKDQYQTCNSTQDTIRQQHQVSTRTVATSQESPVSSST
TYRNKIIEKRSSSKGPAQVGAISTSNESPPATSSCSSPADSPESLQARTVTSAAALCPPKIQTGGIVRSQTTGKNTITSVLAGKAMTSTTTTNHGFGKQSGNANEKDTAPNTFDRQIVHVSSSQMVQIP
SSVHQAKISTAGSNQNRPIASMIQNCVPVSAALTNLQLQPICRGEPGSQIIMTSSGQILVMPPQSKSSNNQMIISGGNGSSLVMNNSPQGVVINQQGQQILSGDMIHGINNQVLDGQSANILQNSGNQV
VIQGQNLMSGNGNRVIGSNNSNFIVNSTNNMQPMIINNSNIISHNGNVLQQGQNVIHGNQGNLISGTKVISSNGNILNGSNIISNQSNVLASNNQIIGNSNSTNLISPNGTVVLPNGYVLQPHQFTTVD
GQVVNVINQDNGTHQFIQSTQQRIILSPDSKRRAKKRKSSSGTPPTPHSLSPQQSPTIQQHSPTIVQQNQSGAMLQITPQYQQQSFQISPGMSGITLVQNKPQTNSGPQQQILLQNGQTIIQPLNIIGQ
QLLVPAGLMVAPDTTLLQIQNVGPCGSILTPQGMMIRASSPQNKNFLSPNSGGQQFIVSGNGQISPIGQMYSTPMGLVVPQTNSNNSGPTYVQQNTTILQQQTTMINSANGNGGSSNVDADIRSQNQQS
MSQSQQQQHHHHRTVSVSPPDTTTHSPRSPERPPSQRSNGSDTNMVQCVSSSEPDAVSPVADSARSPSNNSDYERSGGISYQQNTFKPSDAKIRRIQTPTQLSNSHVITQGNQSKMRPDSHVWGAARGH
PAWPGKIVSPPETYKGVATSSDSTWVQWFGGRPNVELVSINSLKSLSEGLEAHHKAQKDTRKSRKLNSQLECAIQEAMAELDRTVTATSSTPTTPSNRNKARTLKPTSTQQRTKPVKIAPAPPNSTTHL
PAKYLRQSK
>Bcop_v1_g017396-RI, MBD5-like
MADTSPSKKRKLDGAWSPASSISSSDSTRSSQTTALCNYVGSPIPNQMVIANPRVQPTIIQKRQVVGIPAQQHLPQQQSQTVVYQNQNVGENKVANVTVSRTPPWRKNSIVNPTGYLDHQTAILHSSIL
NVHSPDIAENSNQSTISNSNVSYQSEPKLSNQLNVSTPNNDSVQPHTIGSEQSLSVMQQHQHNQNVFIQQQQQQSGVESAGGKTQPHVIIGSNGQPIRILQNGQIIYSNPTMQSVKVQDTQNNASTGIV
SSGPRIISDEIIQPVQTTNVHQQVFQQQNAPRMMAPQQIIPQGQMEPVQIQHNRQVLQQQPQQQASSGQTQIITVQQHVPTSQPFAMNQSVRQPHPQSIVQQQAVNRSHQDHRIIMPSSNQTDSNQSME
LSIGGQTVTHLPNGMVQVQQNCDIAHMKQQQQIQLQQQIQQQNMLFRQQQELQQNPRLKDQYQTCNSTQDTIRQQHQVSTRTVATSQESPVSSSTTYRNKIIEKRSSSKGPAQVGAISTSNESPPATSS
CSSPADSPESLQARTVTSAAALCPPKIQTGGIVRSQTTGKNTITSVLAGKAMTSTTTTNHGFGKQSGNANEKDTAPNTFDRQIVHVSSSQMVQIPSSVHQAKISTAGSNQNRPIASMIQNCVPVSAALT
NLQLQPICRGEPGSQIIMTSSGQILVMPPQSKSSNNQMIISGGNGSSLVMNNSPQGVVINQQGQQILSGDMIHGINNQVLDGQSANILQNSGNQVVIQGQNLMSGNGNRVIGSNNSNFIVNSTNNMQPM
IINNSNIISHNGNVLQQGQNVIHGNQGNLISGTKVISSNGNILNGSNIISNQSNVLASNNQIIGNSNSTNLISPNGTVVLPNGYVLQPHQFTTVDGQVVNVINQDNGTHQFIQSTQQRIILSPDSKRRA
KKRKSSSGTPPTPHSLSPQQSPTIQQHSPTIVQQNQSGAMLQITPQYQQQSFQISPGMSGITLVQNKPQTNSGPQQQILLQNGQTIIQPLNIIGQQLLVPAGLMVAPDTTLLQIQNVGPCGSILTPQGM
MIRASSPQNKNFLSPNSGGQQFIVSGNGQISPIGQMYSTPMGLVVPQTNSNNSGPTYVQQNTTILQQQTTMINSANGNGGSSNVDADIRSQNQQSMSQSQQQQHHHHRTVSVSPPDTTTHSPRSPERPP
SQRSNGSDTNMVQCVSSSEPDAVSPVADSARSPSNNSDYERSGGISYQQNTFKPSDAKIRRIQTPTQLSNSHVITQGNQSKMRPDSHESPSTPTLCSTATFCIGELVWGAARGHPAWPGKIVSPPETYK
GVATSSDSTWVQWFGGRPNVELVSINSLKSLSEGLEAHHKAQKDTRKSRKLNSQLECAIQEAMAELDRTVTATSSTPTTPSNRNKARTLKPTSTQQRTKPVKIAPAPPNSTTHLPAKYLRQSK
>Bcop_v1_g017396-RH, MBD5-like
MADTSPSKKRKLDGAWSPASSISSSDSTRSSQTTALCNYVGSPIPNQMVIANPRVQPTIIQKRQVVGIPAQQHLPQQQSQTVVYQNQNVGENKVANVTVSRTPPWRKNSIVNPTGYLDHQTAILHSSIL
NVHSPDIAENSNQSTISNSNVSYQSEPKLSNQLNVSTPNNDSVQPHTIGSEQSLSVMQQHQHNQNVFIQQQQQQSGVESAGGKTQPHVIIGSNGQPIRILQNGQIIYSNPTMQSVKVQDTQNNASTGIV
SSGPRIISDEIIQPVQTTNVHQQVFQQQNAPRMMAPQQIIPQGQMEPVQIQHNRQVLQQQPQQQASSGQTQIITVQQHVPTSQPFAMNQSVRQPHPQSIVQQQAVNRSHQDHRIIMPSSNQTDSNQSME
LSIGGQTVTHLPNGMVQVQQNCDIAHMKQQQQIQLQQQIQQQNMLFRQQQELQQNPRLKDQYQTCNSTQDTIRQQHQVSTRTVATSQESPVSSSTTYRNKIIEKRSSSKGPAQVGAISTSNESPPATSS
CSSPADSPESLQARTVTSAAALCPPKIQTGGIVRSQTTGKNTITSVLAGKAMTSTTTTNHGFGKQSGNANEKDTAPNTFDRQIVHVSSSQMVQIPSSVHQAKISTAGSNQNRPIASMIQNCVPVSAALT
NLQLQPICRGEPGSQIIMTSSGQILVMPPQSKSSNNQMIISGGNGSSLVMNNSPQGVVINQQGQQILSGDMIHGINNQVLDGQSANILQNSGNQVVIQGQNLMSGNGNRVIGSNNSNFIVNSTNNMQPM
IINNSNIISHNGNVLQQGQNVIHGNQGNLISGTKVISSNGNILNGSNIISNQSNVLASNNQIIGNSNSTNLISPNGTVVLPNGYVLQPHQFTTVDGQVVNVINQDNGTHQFIQSTQQRIILSPDSKRRA
KKRKSSSGTPPTPHSLSPQQSPTIQQHSPTIVQQNQSGAMLQITPQYQQQSFQISPGMSGITLVQNKPQTNSGPQQQILLQNGQTIIQPLNIIGQQLLVPAGLMVAPDTTLLQIQNVGPCGSILTPQGM
MIRASSPQNKNFLSPNSGGQQFIVSGNGQISPIGQMYSTPMGLVVPQTNSNNSGPTYVQQNTTILQQQTTMINSANGNGGSSNVDADIRSQNQQSMSQSQQQQHHHHRTVSVSPPDTTTHSPRSPERPP
SQRSNGSDTNMVQCVSSSEPDAVSPVADSARSPSNNSDYERSGGISYQQNTFKPSDAKIRRIQTPTQLSNSHVITQGNQSKMRPDSHVWGAARGHPAWPGKIVSPPETYKGVATSSDSTWVQWFGGRPN
VELVSINSLKSLSEGLEAHHKAQKDTRKSRKLNSQLECAIQEAMAELDRTVTATSSTPTTPSNRNKARTLKPTSTQQRTKPVKIAPAPPNSTTHLPAKYLRQSK
>Bcop_v1_g016381-RA, TET-family, TET2-like, DMAD-like
MKEQPQQPMAPMAFYPTWQADPSQGWQNQFIQQIPQNTPAITSLNSIEFQPQGYSYQPNGYVQTGLGFEPNYGRTYATPVQRYEFQTNQMSPINQVSLQSVSFAPAVTYTSQVSTGSTPMLLSSQNDLN
KTVSGNDMPGYPRVNSVPPRSLNCNGYSGDYSGNQNHNLTNNVPQQQQHNQHQQQQQQQMQQHHQQQHNNSNQMHPPQVPQSPNRSNMSTPVSQHMMVPTGPGSNHMSQIPQSPNSLQNNGNQQQIQMQ
SPNHHHSQLQTASRYPSHPGTPHSMQQSPNKNPQMLHSHSGMHTPNPQHTAHQHNMTAMSPNHNMNHVQEWSWNNHTVPAGNNEIFNQSDRVNLNTRLKTMILSKNDQKDHDHQNGDGTASAANQQNTA
QTGHFLSYSHHLRDINPTTNDSRCLQPCSTQVAEPIGGGGESIWKPFGYSKPNDFQYSDQKLENSVRKTDYGTDETKEKNAKGKATAAQKSQNVDGVKVKIEPEKVKPKRQRKKPAAKQTQVNDGPNKS
SNNESCHNANSRQSYSTYGIDHIHPPHEQPMQQQPIPKPYYLPKENETVPTPYDPAIKIKQEPMSFSNYNPTTNHPVENFDLAQVKMEGYERNYQNFINYADYCQSQNPTANRGQQQQPTQNETDYPCF
NPNYNYPSFSTANYQTNYPNYNQYTGQQVQQQPMNIEPVPQATKQEPYLQEQHVKTENESENKIVLTNYEKEIPVYTYPNPSKLPAGDTNQKAEPQKFPYLGEGGPATLSESTGFSCCRQGGTNQPSDE
HIRDGTCVGRQTKDELLLDDDAEKSQDEAGERNAVPSPDRTEKGLKPEVPDCDCFSSDKNPPEPGSYYTHLGSAASLEDLRKDFEQRVGITGRQLRIEKIVYTGKEGKTGQGCPLAKWIIRRVDLEEKI
LCLVKRRQGHRCQASFIVVCIVAWDGLPKYEADNAYKLLVHKLNKFGLPTTRRCATNENRTCACQGLDPETCGASFSFGCSWSMYYNGCKYARSKTVRKFRLSVKSEEPEIEEHMNVLATILSPLYVNV
APKAYENQCKYEQEAPDCRLGLKQGKPFSGVTTCIDFCAHSHRDLHNMQDGCTVQVGLLRQRGPGVAAAAEDEQLHVLPLYTMDTTDEFGSAEAQTEKNKTGAVQVLEKFPCEVRVRATPLQPCRRHGK
KRNGKEVAEDPPPPPTLPAEIQPAKKESSKKSAKNKSKQPPPSSPSPRAQTPQASNTSPSTNNTSAFSTPINQNLMNTNSSLINMASMIDNFTDAQLQSNHISSTVLDSPYSYDYTTGSYIDNRNFYNQ
WPPGSDYGYPDRCEQVKSRGSEENPDSTTLSTANSAFSPKITDPNKFSLLDDSTKDQGFAKPKPPEYPHNYGYHSNKPAYSPMYPPYTPYEHYNNMDYYSNDKLRYNFYNPHHHQPYPQYNMYHSNPAL
SAPSPSPQPNWNMYPSPAPMSMHPMQPQMQQPSPKMPEHITPTSEPIGEITEVNDNLECFQDSQMGGVGIALGHGSVLIECAKHEMHATTALRKPNRLNPTRITLIFYQHRNLNRHKHGIEEWEEKMRL
KKLGITPQDVEEANKEKAIKLEEDIKAEMRDEPPAGEKFELDYEKLNRDKEEVSLRAPTLTTTSWTTLFPMHPCVVTGPYQEGNNPTATQSVAEGSVQQQQQQQQ
>Bcop_v1_g016384-RA, TET-family, TET2-like, DMAD-like
MSVQTTEDGTDNSLPSFSTFSSELDPSWDYYDRDRSSTDVITSQSNNYTRPWEMETKDKGFEPFSKLPSFQSQFHAYSETNLVPEPSLPQITPVPVPVSPASPGNGSLTQLTPINSIQSGLTQLQSPSF
HTLTAVNTRTYPLVPAPIQARDIPAINQQYLEERHIQLYQPISTFPPQNMVTVIKNEPNYEIKNGLHHTNFMNPLIDNGFDIKVEKSESPVVRCDARKKERRKNRASSLESSAESESSAMDMGDGNPGQ
VAAISSTASFKSPMGSMGMNEDSEGNGEKQTKKKRKRCGECIGCQKKDNCGDCAPCRNDKSHQICKQRRCEKLTDKKILLSLCFCILDTLDTTLQQQQKKVTQCLIHQLYTPIGTTPKLESYFDFEKGY
KVEIFWTVDIYCSIGKLICIHQMLALVTLVQSQQSRVLSEILSTKIKLGREAHIILLLRIHFSVMLCKNNLELELENLKMFH
>Bcop_v1_g016384-RB, TET-family, TET2-like, DMAD-like
MSVQTTEDGTDNSLPSFSTFSSELDPSWDYYDRDRSSTDVITSQSNNYTRPWEMETKDKGFEPFSKLPSFQSQFHAYSETNLVPEPSLPQITPVPVPVSPASPGNGSLTQLTPINSIQSGLTQLQSPSF
HTLTAVNTRTYPLVPAPIQARDIPAINQQYLEERHIQLYQPISTFPPQNMVTVIKNEPNYEIKNGLHHTNFMNPLIDNGFDIKVEKSESPVVRCDARKKERRKNRASSLESSAESESSAMDMGDGNPGQ
VAAISSTASFKSPMGSMGMNEDSEGNGEKQTKKKRKRCGECIGCQKKDNCGDCAPCRNDKSHQICKQRRCEKLTDKKSC

>Bcop_v1_g020988-RA, DNMT2-like
MDDFRVLELFSGIGGMHYALKSSQIPGTVVAAADINTIANDVYRHNHPDIKVLNNNIQKFTPEFIRTLNVNTILMSPPCQPFTRVGNQMDVEDRRADPFVHICQILPELKCIDWILMENVKGFDTSKAR
EMFVESLRQSEFHYQEFILSPTQLGIPNSRHRYYCLARRSKAFTFASERILTSLPSADTMVPDVQQISQFISSVDANGLPLPEKVLQKYFSVLDICYPDSINSMCFTKAYTHYAEGTGSVFCPLDRNAV
DKCCSQLELIFDDKTTVDLLKSLQLRYFTPAEVSRLMCFPENDFSFPEKISNKSRYRLLGNSINVLVVSELIKLLYED
>Bcop_v1_g019039-RA, SETDB1-like
MDKTPDCNLDRIKDEPAECKNEPLSRCDGQANEFVDNTSIMVSVPLSASEISEIDSIVLNDSVSKEVENDKPADVPPVEPMEVNEQDKSDIPDANLTEDIDLPSTATQSTTVATCDKSIADPSSTQPNI
EKGDDTNVDENDVVMLLSDSEDEDTARKSRLVDTAEMIEKFSTQGVTLEVVKPANDKLRAASVRTENVENKTNDNDTSVIMLDDSPREVEAPTNDRDSAKRNAIEVDEEPPPQTANETDDPAQNALQMI
RQLVDGCYPDDKESTAATTATTTTTTTTASKSTDSPAPNESQKTQSTFRLVGMDKLLETNSVKSGEKRECINFDCKSKKSVEFYKAPLLALSHFNVPRKVNRAQFICQNCFDVSVNDYERMGVALVNQQ
PLLLEEVPVRPEVVEILDSEEEDNGGGSSKYVDDIKPLSLDTLTLLEDHFEDVLKEAFNRINIAQQMAWTDQILNHKIDKNETKTEELELQVKSMQKLADSMYDKLYKNTNFVIEELPPFDLNENKQLH
MYGPNYPPQGELIYPPVDMNSLYYAVRAKLLTTWIPCKVTERVDASVDGKKMFKVRFLRTKGVLNKTVPGNHLAYGTAPSVRLTVGTRVIALFTMEPHDNLRSTFFPGIVAEPLQQYNKWRYLIFFDDG
YAQYVTPDNLRLICSPSPNVWEDVYPDAAEFIKSYLEQYKTQRPMVQVKRGQRMITEWNSEWTHARVHDIDGSLVQMYFENAKRSEWIYRGSTRLGPLYKDRQLQKHSVSSGKRNEPFVEYMTIDDDGK
NKTSTAAKPVQAQKPVAAKPSHTNTPSQTPNPPSHTPNTSSQVAPQSGEEKRSVARKSVTAPQRPQPAVQHMNSATIYIEEDNRPKGKVVYYTAKKHMPPRKFVPHKCGVGCLYEVKHNLSTYSPLAKP
LLSGWERKILRNKTKKSVEYRTPCGRNLRNMAELHRYLRQTKCALNVDNFDFDFMVHCLAEYVIDTCVVQKQDLSDGCEPMVVPCVNSYDNTMPPPCVYSAKRIPNEGFGNPNTPIDQVGYEFKRLPNP
VPTGIYECNTRCKCSKTCLNRVAQHPLQLKLQVFKTVSRGWGLRCLNDVPKGSFICIYAGDLLTEQNANIAGDNYGDEYFAELDYIEVVENLKEGYEPGVMDDDFDDDEFDPDKTKDDSDDDEYVATVS
QGPRQTRYNTRHSVDASNKKKTTETPKLKVDPASGSNSDEEDAQRQLISFMPNADSVEFENNTTNRYRSIRKFYGKNESVFIMDAKKLGNIGRYFNHSCSPNLFVQNVFVDTHDLRFPWVAFFALSHIR
AGSELTWNYNYDVGSVPGKVMYCECGAENCRGRLL
>Bcop_v1_g018977-RA, ALKBH6-like
MDISDFTVQSCPNTVIYIPNFITPAEEQKILTSVYNAPKPKWDQLTHRRLLNYGGVPHKNGMIAEEMPGWLQNYVDKVNNLGIFDTKKANHVLVNEYLPKQGIMPHLDGPLFYPIITTISCGSHTVLEF
LEANDNNPEFSETLMSANDGSSDQMDGKLSRKIMCKLFIEPRSLLILRDDMYHKYLHAICEIDEDVISENVANLQHCSNSYKLGDKIKRETRVSLTIRHVPKTSKMKLKLGN

#### Supplemental Table S18 A-F: DNA modification percentages in male embryonic genomic DNA.

Note in tables A and B below, the effective genome size is doubled since coverage is calculated per strand and counts of how many bases and how many modified bases are from both strands.

**S18A: Statistics on all putative modified adenines given minimum strand coverage levels**

| Min. Strand Coverage | #A = Number of adenines ≥ Min. Coverage | #A as % of all adenines in the genome | #modA = Number of modified adenines ≥ Min. Coverage | #modA as % of #A | #6mA = Number of 6-methyl adenines ≥ Min. Coverage | #6mA as % of #A |
|---|---|---|---|---|---|---|
| 3 | 184940136 | 98.9739 | 247015 | 0.1336 | 66414 | 0.0359 |
| 5 | 182482744 | 97.6587 | 245841 | 0.1347 | 66414 | 0.0364 |
| 10 | 158406693 | 84.774 | 233673 | 0.1475 | 63472 | 0.0401 |
| 15 | 110866645 | 59.3322 | 198921 | 0.1794 | 53535 | 0.0483 |
| 20 | 50775937 | 27.1736 | 120845 | 0.238 | 31325 | 0.0617 |
| 25 | 12788726 | 6.8441 | 41950 | 0.328 | 10218 | 0.0799 |
| 50 | 52445 | 0.0281 | 1090 | 2.0784 | 210 | 0.4004 |

*modA is any adenine that was flagged as “modified_base” by kineticsTools and 6mA is the subset of modA that was predicted to be 6mA by kineticsTools.

**S18B: Statistics on all putative modified adenines inside GAG motifs given minimum strand coverage levels**

| Min. Strand Coverage | #GAG = Number of GAG ≥ Min. Coverage | #GAG as % of all GAG in the genome | #modGAG = Number of modified GAG ≥ Min. Coverage | #modGAG as % of #A | #6mA-GAG = Number of 6mA-GAG ≥ Min. Coverage | #6mA-GAG as % of #GAG |
|---|---|---|---|---|---|---|
| 3 | 5365180 | 98.9799 | 47082 | 0.8775 | 16464 | 0.3069 |
| 5 | 5296391 | 97.7108 | 47022 | 0.8878 | 16464 | 0.3109 |
| 10 | 4614029 | 85.1222 | 45850 | 0.9937 | 15875 | 0.3441 |
| 15 | 3229440 | 59.5785 | 40036 | 1.2397 | 13314 | 0.4123 |
| 20 | 1446315 | 26.6824 | 24534 | 1.6963 | 7447 | 0.5149 |
| 25 | 347929 | 6.4188 | 8261 | 2.3743 | 2310 | 0.6639 |
| 50 | 1218 | 0.0225 | 101 | 8.2923 | 11 | 0.9031 |

*modGAG is an GAG flagged as containing a modified A and 6mA-GAG is the subset of modGAG where the modification was predicted to be 6mA by kineticsTools.

**S18C: Statistics on all putative modified cytosines given minimum strand coverage levels**

| Min. Strand Cover-age | #C = Number of cytosines ≥ Min. Coverage | #C as % of all cytosines in the genome | #modC = Number of modified cytosines ≥ Min. Coverage | #modC as % of #C | #4mC = Number of 4-methyl cyto-sines ≥ Min. Cover-age | #4mC as % of #C | #5mC = Number of 5-methyl cytosines ≥ Min. Coverage | #5mC as % of #C |
|---|---|---|---|---|---|---|---|---|
| 3 | 102025314 | 98.9127 | 610766 | 0.5986 | 115955 | 0.1137 | 267966 | 0.2626 |
| 5 | 100469957 | 97.4048 | 609640 | 0.6068 | 115955 | 0.1154 | 267757 | 0.2665 |
| 10 | 86136561 | 83.5087 | 583489 | 0.6774 | 112556 | 0.1307 | 253146 | 0.2939 |
| 15 | 58293210 | 56.5148 | 486588 | 0.8347 | 98439 | 0.1689 | 202877 | 0.348 |
| 20 | 25049775 | 24.2856 | 278927 | 1.1135 | 61097 | 0.2439 | 108946 | 0.4349 |
| 25 | 5881088 | 5.7017 | 89906 | 1.5287 | 21491 | 0.3654 | 32944 | 0.5602 |
| 50 | 22901 | 0.0222 | 2347 | 10.2485 | 706 | 3.0828 | 579 | 2.5283 |

*modC is any cytosine that was flagged as "modified_base" by kineticsTools whereas 4mC and 5mC are subsets of modC predicted to be those specific modifications by kineticsTools.

**S18D: Statistics on all putative modified cytosines in CpG dimers given minimum strand coverage levels**

| Min. Strand Cover-age | #CG = Number GCG ≥ Min. Coverage | #CG as % of all CG in the genome | #modCG = Number of modified CG ≥ Min. Coverage | #modCG as % of #CG | #4mCG = Number of 4-methyl CG ≥ Min. Cover-age | #4mCG as % of #CG | #5mCG = Number of 5-methyl CG ≥ Min. Coverage | #5mCG as % of #CG |
|---|---|---|---|---|---|---|---|---|
| 3 | 20574879 | 98.8276 | 265810 | 1.2919 | 53713 | 0.2611 | 112978 | 0.5491 |
| 5 | 20242476 | 97.231 | 265491 | 1.3116 | 53713 | 0.2653 | 112919 | 0.5578 |
| 10 | 17288660 | 83.0429 | 255990 | 1.4807 | 52335 | 0.3027 | 107824 | 0.6237 |
| 15 | 11610868 | 55.7706 | 215423 | 1.8554 | 46055 | 0.3967 | 87656 | 0.7549 |
| 20 | 4973418 | 23.8889 | 123759 | 2.4884 | 28576 | 0.5746 | 47657 | 0.9582 |
| 25 | 1170657 | 5.623 | 39692 | 3.3906 | 9986 | 0.853 | 14609 | 1.2479 |
| 50 | 4195 | 0.0201 | 777 | 18.5221 | 309 | 7.3659 | 139 | 3.3135 |

*modCG is any CG flagged as containing a modified C and 4mCG of 5mCG are the subsets of modCG where the modification was predicted to be 4mC or 5mC by kineticsTools.

**S18E: CpG methylation according to the more sensitive AgIn Algorithm:**

| *CpG Methylation State* | Count | Percent |
|---|---|---|
| *0* | 10022667 | 93.6 |
| *1* | 689756 | 6.4 |

Blocks containing 6.4% of interrogated CpGs were not considered to be unmethylated. That indicates an upper limit of CpG methylation at 6.4%. This may be an overestimate since the algorithm calls "blocks" of CpGs as either methylated or unmethylated. Each block contains data from both strands of 40 or more consecutive CpGs and the methylation signals may come from only a subset therein. Nevertheless, the less sensitive approach that looked at individual Cs strand-specifically estimated 0.5-1% CpGs containing 5mC or 1.3-2.5% with modification signals in general. Thus, the algorithms together suggest 0.5-6.4%.

**S18F: Statistics on all putative modified cytosines in GCG motif given minimum strand coverage level**

| Min. Strand Cover-age | #GCG = Number GCG ≥ Min. Coverage | #GCG as % of all GCG in the genome | #modGCG = Number of modified GCG ≥ Min. Coverage | #modGCG as % of #GCG | #4mC-GCG = Number of 4-methyl GCG ≥ Min. Cover-age | #4mC-GCG as % of #GCG | #5mC-GCG = Number of 5-methyl GCG ≥ Min. Coverage | #5mC-GCG as % of #GCG |
|---|---|---|---|---|---|---|---|---|
| 3 | 3156626 | 98.9159 | 80487 | 2.5498 | 16971 | 0.5376 | 28215 | 0.8938 |
| 5 | 3113898 | 97.5769 | 80450 | 2.5836 | 16971 | 0.545 | 28213 | 0.906 |
| 10 | 2701392 | 84.6507 | 78284 | 2.8979 | 16680 | 0.6175 | 27123 | 1.004 |
| 15 | 1869990 | 58.5979 | 67593 | 3.6146 | 14997 | 0.802 | 22651 | 1.2113 |
| 20 | 819836 | 25.6903 | 39841 | 4.8596 | 9480 | 1.1563 | 12595 | 1.5363 |
| 25 | 193764 | 6.0718 | 12745 | 6.5776 | 3340 | 1.7237 | 3764 | 1.9426 |
| 50 | 945 | 0.0296 | 387 | 40.9524 | 103 | 10.8995 | 68 | 7.1958 |

*modGCG is any GCG flagged as containing a modified C and 4mC-GCG of 5mC-GCG are the subsets of modGCG where the modification was predicted to be 4mC or 5mC by kineticsTools.

**Supplemental Table S19: Which dimers are observed with modifications more often than expected?**

| Modification | K | Position | Kmer | Observed | Expected | Chi-Square | P value | Std Residual |
|---|---|---|---|---|---|---|---|---|
| m4C | 2 | 1 | CA | 29905 | 41657.1 | 57124.7 | 0 | -57.6 |
| m4C | 2 | 1 | CC | 22323 | 19846.1 | 57124.7 | 0 | 17.6 |
| **m4C** | **2** | **1** | **CG** | **53713** | **23404.2** | **57124.7** | **0** | **198.1** |
| m4C | 2 | 1 | CT | 10014 | 31047.7 | 57124.7 | 0 | -119.4 |
| m5C | 2 | 1 | CA | 86775 | 96267.5 | 88914.7 | 0 | -30.6 |
| m5C | 2 | 1 | CC | 35919 | 45863.2 | 88914.7 | 0 | -46.4 |
| **m5C** | **2** | **1** | **CG** | **112978** | **54085.8** | **88914.7** | **0** | **253.2** |
| m5C | 2 | 1 | CT | 32294 | 71749.5 | 88914.7 | 0 | -147.3 |
| m6A | 2 | 1 | AA | 18023 | 25507.8 | 41495 | 0 | -46.9 |
| m6A | 2 | 1 | AC | 12618 | 11545.7 | 41495 | 0 | 10 |
| **m6A** | **2** | **1** | **AG** | **27676** | **9816.2** | **41495** | **0** | **180.3** |
| m6A | 2 | 1 | AT | 8097 | 19544.3 | 41495 | 0 | -81.9 |
| modA | 2 | 1 | AA | 69516 | 94871.6 | 114285.5 | 0 | -82.3 |
| modA | 2 | 1 | AC | 43516 | 42942.3 | 114285.5 | 0 | 2.8 |
| **modA** | **2** | **1** | **AG** | **94554** | **36509.7** | **114285.5** | **0** | **303.8** |
| modA | 2 | 1 | AT | 39429 | 72691.5 | 114285.5 | 0 | -123.4 |
| modC | 2 | 1 | CA | 185193 | 219419.3 | 225872.7 | 0 | -73.1 |
| modC | 2 | 1 | CC | 89933 | 104534.5 | 225872.7 | 0 | -45.2 |
| **modC** | **2** | **1** | **CG** | **265810** | **123276** | **225872.7** | **0** | **406** |
| modC | 2 | 1 | CT | 69830 | 163536.3 | 225872.7 | 0 | -231.7 |

*modA is any adenine that was flagged as “modified_base” by kineticsTools and m6A is the subset of modA that was predicted to be 6mA by kineticsTools.
*modC is any cytosine that was flagged as “modified_base” by kineticsTools whereas m4C and m5C are subsets of modC predicted to specifically be 4mC and 5mC by kineticsTools.

#### Supplemental Table S20: Which trimers are observed with modifications more often than expected?

| Modification | K | Positon | Kmer | Observed | Expected | Chi-Square | P value | Std Residual |
|---|---|---|---|---|---|---|---|---|
| m4C | 3 | 2 | ACA | 4986 | 13691.2 | 104230.7 | 0 | -74.4 |
| m4C | 3 | 2 | ACC | 3470 | 6221.9 | 104230.7 | 0 | -34.9 |
| m4C | 3 | 2 | ACG | 9484 | 6881.9 | 104230.7 | 0 | 31.4 |
| m4C | 3 | 2 | ACT | 1146 | 9723 | 104230.7 | 0 | -87 |
| m4C | 3 | 2 | CCA | 4156 | 7440.3 | 104230.7 | 0 | -38.1 |
| m4C | 3 | 2 | CCC | 3582 | 3172.8 | 104230.7 | 0 | 7.3 |
| m4C | 3 | 2 | CCG | 6623 | 4181.2 | 104230.7 | 0 | 37.8 |
| m4C | 3 | 2 | CCT | 1669 | 5051.7 | 104230.7 | 0 | -47.6 |
| m4C | 3 | 2 | GCA | 11349 | 7265.5 | 104230.7 | 0 | 47.9 |
| m4C | 3 | 2 | GCC | 9415 | 3911.2 | 104230.7 | 0 | 88 |
| **m4C** | **3** | **2** | **GCG** | **16971** | **3587.5** | **104230.7** | **0** | **223.4** |
| m4C | 3 | 2 | GCT | 4966 | 6028.6 | 104230.7 | 0 | -13.7 |
| m4C | 3 | 2 | TCA | 9414 | 13260.2 | 104230.7 | 0 | -33.4 |
| m4C | 3 | 2 | TCC | 5856 | 6540.2 | 104230.7 | 0 | -8.5 |
| m4C | 3 | 2 | TCG | 20635 | 8753.6 | 104230.7 | 0 | 127 |
| m4C | 3 | 2 | TCT | 2233 | 10244.3 | 104230.7 | 0 | -79.2 |
| m5C | 3 | 2 | ACA | 24931 | 31639.6 | 126488.4 | 0 | -37.7 |
| m5C | 3 | 2 | ACC | 8954 | 14378.4 | 126488.4 | 0 | -45.2 |
| **m5C** | **3** | **2** | **ACG** | **31508** | **15903.7** | **126488.4** | **0** | **123.7** |
| m5C | 3 | 2 | ACT | 5767 | 22469.4 | 126488.4 | 0 | -111.4 |
| m5C | 3 | 2 | CCA | 12446 | 17194.1 | 126488.4 | 0 | -36.2 |
| m5C | 3 | 2 | CCC | 8422 | 7332.2 | 126488.4 | 0 | 12.7 |
| **m5C** | **3** | **2** | **CCG** | **23316** | **9662.6** | **126488.4** | **0** | **138.9** |
| m5C | 3 | 2 | CCT | 10670 | 11674.3 | 126488.4 | 0 | -9.3 |
| m5C | 3 | 2 | GCA | 26080 | 16790.2 | 126488.4 | 0 | 71.7 |
| m5C | 3 | 2 | GCC | 9145 | 9038.6 | 126488.4 | 0 | 1.1 |
| **m5C** | **3** | **2** | **GCG** | **28215** | **8290.5** | **126488.4** | **0** | **218.8** |
| m5C | 3 | 2 | GCT | 7396 | 13931.8 | 126488.4 | 0 | -55.4 |
| m5C | 3 | 2 | TCA | 23318 | 30643.6 | 126488.4 | 0 | -41.8 |
| m5C | 3 | 2 | TCC | 9398 | 15114 | 126488.4 | 0 | -46.5 |
| m5C | 3 | 2 | TCG | 29939 | 20229 | 126488.4 | 0 | 68.3 |
| m5C | 3 | 2 | TCT | 8461 | 23674 | 126488.4 | 0 | -98.9 |
| m6A | 3 | 2 | AAA | 2026 | 10597 | 160668.1 | 0 | -83.3 |
| m6A | 3 | 2 | AAC | 2064 | 3995.7 | 160668.1 | 0 | -30.6 |
| m6A | 3 | 2 | AAG | 3372 | 3383.1 | 160668.1 | 0 | -0.2 |
| m6A | 3 | 2 | AAT | 711 | 7532 | 160668.1 | 0 | -78.6 |
| m6A | 3 | 2 | CAA | 1150 | 4745.7 | 160668.1 | 0 | -52.2 |
| m6A | 3 | 2 | CAC | 1055 | 2450.6 | 160668.1 | 0 | -28.2 |
| m6A | 3 | 2 | CAG | 2649 | 2185.5 | 160668.1 | 0 | 9.9 |
| m6A | 3 | 2 | CAT | 1733 | 3788.8 | 160668.1 | 0 | -33.4 |
| **m6A** | **3** | **2** | **GAA** | **12857** | **4940.6** | **160668.1** | **0** | **112.6** |
| **m6A** | **3** | **2** | **GAC** | **7147** | **2136.1** | **160668.1** | **0** | **108.4** |
| **m6A** | **3** | **2** | **GAG** | **16464** | **1926.6** | **160668.1** | **0** | **331.2** |
| m6A | 3 | 2 | GAT | 3828 | 3263.4 | 160668.1 | 0 | 9.9 |
| m6A | 3 | 2 | TAA | 1990 | 5224.5 | 160668.1 | 0 | -44.7 |
| m6A | 3 | 2 | TAC | 2352 | 2963.3 | 160668.1 | 0 | -11.2 |
| m6A | 3 | 2 | TAG | 5191 | 2321.1 | 160668.1 | 0 | 59.6 |
| m6A | 3 | 2 | TAT | 1825 | 4960.1 | 160668.1 | 0 | -44.5 |

*modA is any adenine that was flagged as “modified_base” by kineticsTools and m6A is the subset of modA that was predicted to be 6mA by kineticsTools.
*modC is any cytosine that was flagged as “modified_base” by kineticsTools whereas m4C and m5C are subsets of modC predicted to specifically be 4mC and 5mC by kineticsTools.

#### Supplemental Table S21 A-F: Binomial tests for enrichment or depletion of DNA modifications in various genomic features.

##### S21A: Do gene regions/features as defined by the Maker2 Annotation have higher or lower modification rates than the genome-wide rate?

| MAKER2 GENE FEATURES | MODIFICATION | PERCENT MODIFIED ACROSS GENOME | PERCENT MODIFIED ACROSS FEATURE | BINOMIAL P-VALUE FOR ENRICHMENT | BINOMIAL P-VALUE FOR DEPLETION |
|---|---|---|---|---|---|
| GENES | 6mA | 0.0355 | 0.0352 | 0.98128157 | 0.018956188 |
| | modA | 0.1322 | 0.1316 | 0.964203427 | 0.036007625 |
| | 4mC | 0.1124 | 0.1108 | 0.999929591 | 7.15E-05 |
| | 5mC | 0.2598 | 0.2589 | 0.910102733 | 0.090312522 |
| | modC | 0.5921 | 0.5902 | 0.970909305 | 0.029203296 |
| EXONS | 6mA | 0.0355 | 0.0329 | 1 | 7.54E-13 |
| | modA | 0.1322 | 0.1278 | 0.999999999 | 6.76E-10 |
| | 4mC | 0.1124 | 0.0984 | 1 | 1.02E-66 |
| | 5mC | 0.2598 | 0.2572 | 0.978871468 | 0.021377372 |
| | modC | 0.5921 | 0.5689 | 1 | 4.71E-35 |
| INTRONS | 6mA | 0.0355 | 0.0358 | 0.072865683 | 0.927941142 |
| | modA | 0.1322 | 0.1327 | 0.106287986 | 0.894268744 |
| | 4mC | 0.1124 | 0.1154 | 1.87E-09 | 0.999999998 |
| | 5mC | 0.2598 | 0.2595 | 0.627989008 | 0.373145715 |
| | modC | 0.5921 | 0.5983 | 5.98E-08 | 0.999999941 |
| PROMOTERS | 6mA | 0.0355 | 0.0332 | 0.999509543 | 0.000527151 |
| | modA | 0.1322 | 0.1228 | 1 | 1.76E-12 |
| | 4mC | 0.1124 | 0.1052 | 0.999990523 | 1.02E-05 |
| | 5mC | 0.2598 | 0.2504 | 0.999866666 | 0.000138694 |
| | modC | 0.5921 | 0.5681 | 1 | 3.02E-10 |

**S21B: Are gene regions/features as defined by the Maker2 Annotation enriched with or depleted for DNA modifications with respect to the prevalence of those features in the genome?**

| MAKER2 GENE FEATURES | MODIFICATION | PERCENT OF MODIFICATIONS EXPECTED TO BE IN REGION AT RANDOM | OBSERVED PERCENT OF MODIFICATIONS IN REGION | BINOMIAL P-VALUE FOR ENRICHMENT | BINOMIAL P-VALUE FOR DEPLETION |
|---|---|---|---|---|---|
| **GENES** | 6mA | 56.1611602 | 56.2185684 | 0.38433787 | 0.61864674 |
| | modA | 56.1611602 | 56.5528409 | 4.38E-05 | 0.9999569 |
| | 4mC | 56.1611602 | 56.6038549 | 0.00119808 | 0.99882524 |
| | 5mC | 56.1611602 | 57.2524126 | 2.26E-30 | 1 |
| | modC | 56.1611602 | 57.2653029 | 3.71E-68 | 1 |
| **EXONS** | 6mA | 13.6878976 | 12.1088927 | 1 | 1.18E-33 |
| | modA | 13.6878976 | 12.6514584 | 1 | 3.93E-52 |
| | 4mC | 13.6878976 | 13.8381269 | 0.06906262 | 0.93206208 |
| | 5mC | 13.6878976 | 15.6490002 | 6.89E-185 | 1 |
| | modC | 13.6878976 | 15.1847025 | 7.82E-247 | 1 |
| **INTRONS** | 6mA | 42.4732626 | 44.1096757 | 8.38E-18 | 1 |
| | modA | 42.4732626 | 43.9013825 | 6.39E-47 | 1 |
| | 4mC | 42.4732626 | 42.7657281 | 0.02214394 | 0.97816755 |
| | 5mC | 42.4732626 | 41.6034124 | 1 | 3.91E-20 |
| | modC | 42.4732626 | 42.0806004 | 1 | 2.66E-10 |
| **PROMOTERS** | 6mA | 3.68304918 | 3.5655133 | 0.94778728 | 0.05447281 |
| | modA | 3.68304918 | 3.54189017 | 0.99991225 | 9.16E-05 |
| | 4mC | 3.68304918 | 3.48583502 | 0.99984234 | 0.00016757 |
| | 5mC | 3.68304918 | 3.59038087 | 0.99477972 | 0.00537773 |
| | modC | 3.68304918 | 3.57370908 | 0.99999746 | 2.63E-06 |

**S21C: Do gene regions/features as defined by StringTie have higher DNA modification rates with respect to the genome-wide average?**

| STRINGTIE GENE FEATURES | MODIFICATION | PERCENT MODIFIED ACROSS GENOME | PERCENT MODIFIED ACROSS FEATURE | BINOMIAL P-VALUE FOR ENRICHMENT | BINOMIAL P-VALUE FOR DEPLETION |
|---|---|---|---|---|---|
| **GENES** | 6mA | 0.0355 | 0.0351 | 0.99619467 | 0.003862711 |
| | modA | 0.1322 | 0.1314 | 0.991948166 | 8.11E-03 |
| | 4mC | 0.1124 | 0.111 | 0.999449561 | 5.58E-04 |
| | 5mC | 0.2598 | 0.2589 | 0.920825033 | 0.079543767 |
| | modC | 0.5921 | 0.5915 | 0.734783388 | 0.265759097 |
| **EXONS** | 6mA | 0.0355 | 0.0337 | 0.999999995 | 5.03E-09 |
| | modA | 0.1322 | 0.1291 | 0.999999449 | 5.65E-07 |
| | 4mC | 0.1124 | 0.1023 | 1 | 2.42E-43 |
| | 5mC | 0.2598 | 0.2599 | 0.448100039 | 0.553623203 |
| | modC | 0.5921 | 0.5812 | 1 | 5.51E-11 |
| **INTRONS** | 6mA | 0.0355 | 0.0356 | 0.316477725 | 0.685634126 |
| | modA | 0.1322 | 0.1323 | 0.400622005 | 0.600569585 |
| | 4mC | 0.1124 | 0.1153 | 2.46E-08 | 0.999999976 |
| | 5mC | 0.2598 | 0.2584 | 0.965035978 | 0.035199744 |
| | modC | 0.5921 | 0.5966 | 0.000100661 | 0.999900138 |
| **PROMOTERS** | 6mA | 0.0355 | 0.0341 | 0.978301218 | 0.022776113 |
| | modA | 0.1322 | 0.1238 | 1 | 3.30E-10 |
| | 4mC | 0.1124 | 0.108 | 0.995606095 | 0.004597575 |
| | 5mC | 0.2598 | 0.247 | 0.999999654 | 3.64E-07 |
| | modC | 0.5921 | 0.568 | 1 | 3.02E-10 |

**S21D: Are there higher or lower DNA modification rates in genes or promoter regions when partitioned by gene expression?**

| MAKER2 GENES AND PROMOTERS | MODIFICATION | PERCENT MODIFIED ACROSS GENOME | PERCENT MODIFIED ACROSS FEATURE | BINOMIAL P-VALUE FOR ENRICHMENT | BINOMIAL P-VALUE FOR DEPLETION |
|---|---|---|---|---|---|
| **GENES: HIGHLY EXPRESSED** | 6mA | 0.0355 | 0.0346 | 0.999093866 | 0.000932321 |
| | modA | 0.1322 | 0.1302 | 0.9997053 | 0.00029946 |
| | 4mC | 0.1124 | 0.1085 | 0.999999992 | 8.78E-09 |
| | 5mC | 0.2598 | 0.2572 | 0.993766357 | 0.006306024 |
| | modC | 0.5921 | 0.587 | 0.999332119 | 0.000674214 |
| **GENES: LOWLY EXPRESSED** | 6mA | 0.0355 | 0.036 | 0.051184353 | 0.949577622 |
| | modA | 0.1322 | 0.1336 | 0.0021631 | 0.997862533 |
| | 4mC | 0.1124 | 0.1145 | 0.000372393 | 0.999635009 |
| | 5mC | 0.2598 | 0.2613 | 0.059873187 | 0.940559983 |
| | modC | 0.5921 | 0.6003 | 6.68E-09 | 0.999999993 |
| **GENES: NOT EXPRESSED** | 6mA | 0.0355 | 0.0339 | 0.99990464 | 0.000100206 |
| | modA | 0.1322 | 0.1289 | 0.999953104 | 4.82E-05 |
| | 4mC | 0.1124 | 0.1059 | 1 | 2.13E-10 |
| | 5mC | 0.2598 | 0.2562 | 0.98919457 | 0.010982399 |
| | modC | 0.5921 | 0.5703 | 1 | 2.70E-20 |
| **PROMOTERS: HIGHLY EXPRESSED** | 6mA | 0.0355 | 0.0341 | 0.888360539 | 0.118420887 |
| | modA | 0.1322 | 0.1211 | 0.99999906 | 1.03E-06 |
| | 4mC | 0.1124 | 0.1066 | 0.977702806 | 0.023739974 |
| | 5mC | 0.2598 | 0.2423 | 0.999968501 | 3.39E-05 |
| | modC | 0.5921 | 0.5625 | 0.999996261 | 3.95E-06 |
| **PROMOTERS: LOWLY EXPRESSED** | 6mA | 0.0355 | 0.0335 | 0.952665437 | 0.050871358 |
| | modA | 0.1322 | 0.1234 | 0.999932439 | 7.27E-05 |
| | 4mC | 0.1124 | 0.1073 | 0.959218121 | 0.043182619 |
| | 5mC | 0.2598 | 0.2562 | 0.792770969 | 0.212254363 |
| | modC | 0.5921 | 0.5767 | 0.988893271 | 0.011450768 |
| **PROMOTERS: NOT EXPRESSED** | 6mA | 0.0355 | 0.0322 | 0.997137664 | 0.00319297 |
| | modA | 0.1322 | 0.1242 | 0.999649671 | 0.000374566 |
| | 4mC | 0.1124 | 0.1007 | 0.999966509 | 3.76E-05 |
| | 5mC | 0.2598 | 0.2532 | 0.927537438 | 0.074943257 |
| | modC | 0.5921 | 0.5649 | 0.999967849 | 3.38E-05 |

**S21E: Do regions of the genome annotated as repetitive or the unique regions have higher or lower modification rates than the genome-wide average?**

| REGIONS DEFINED BY REPEAT MASKER | MODIFICATION | PERCENT MODIFIED ACROSS GENOME | PERCENT MODIFIED ACROSS FEATURE | BINOMIAL P-VALUE FOR ENRICHMENT | BINOMIAL P-VALUE FOR DEPLETION |
|---|---|---|---|---|---|
| REPEATS | 6mA | 0.0355 | 0.0411 | 1.14E-137 | 1 |
| | modA | 0.1322 | 0.1496 | 0 | 1 |
| | 4mC | 0.1124 | 0.1294 | 1.34E-231 | 1 |
| | 5mC | 0.2598 | 0.2902 | 0 | 1 |
| | modC | 0.5921 | 0.6561 | 0 | 1 |
| UNIQUE | 6mA | 0.0355 | 0.0318 | 1 | 1.47E-100 |
| | modA | 0.1322 | 0.1206 | 1 | 1.62E-259 |
| | 4mC | 0.1124 | 0.1003 | 1 | 4.29E-181 |
| | 5mC | 0.2598 | 0.238 | 1 | 4.87E-249 |
| | modC | 0.5921 | 0.5463 | 1 | 0 |

* Note: the unique regions here are regions not labeled as repeats by RepeatMasker. In other words, the assembly was partitioned into repeat and unique regions.

**S20F: Is there enrichment or depletion of DNA modifications in the regions of the genome annotated as repetitive or the unique regions?**

| REGIONS DEFINED BY REPEAT MASKER | MODIFICATION | PERCENT OF MODIFICATIONS EXPECTED TO BE IN REGION AT RANDOM | OBSERVED PERCENT OF MODIFICATIONS IN REGION | BINOMIAL P-VALUE FOR ENRICHMENT | BINOMIAL P-VALUE FOR DEPLETION |
|---|---|---|---|---|---|
| REPEATS | 6mA | 39.5417134 | 46.2613304 | 1.26E-270 | 1 |
| | modA | 39.5417134 | 45.263648 | 0 | 1 |
| | 4mC | 39.5417134 | 48.0281144 | 0 | 1 |
| | 5mC | 39.5417134 | 46.6130031 | 0 | 1 |
| | modC | 39.5417134 | 46.2325997 | 0 | 1 |
| UNIQUE | 6mA | 60.4582866 | 53.7386696 | 1 | 1.26E-270 |
| | modA | 60.4582866 | 54.736352 | 1 | 0 |
| | 4mC | 60.4582866 | 51.9718856 | 1 | 0 |
| | 5mC | 60.4582866 | 53.3869969 | 1 | 0 |
| | modC | 60.4582866 | 53.7674003 | 1 | 0 |

### Supplemental Table S22: L candidate contigs and corresponding proteins

| Contig | Length | Proportion of bases with given PacBio coverage: 0 | 1 | 2 | 3 | 4+ | N transcripts | Genes | Similar to | Pfam | InterPro | GO |
|---|---|---|---|---|---|---|---|---|---|---|---|---|
| genome | 312049034 | 0.026 | 0.002 | 0.002 | 0.002 | 0.968 | . | . | . | . | . | . |
| contig_165 | 8269 | 0.016 | 0.113 | 0.355 | 0.333 | 0.183 | 0 | . | . | . | . | . |
| contig_708 | 8333 | 0.056 | 0.192 | 0.289 | 0.284 | 0.179 | 0 | . | . | . | . | . |
| contig_90 | 10696 | 0.149 | 0.316 | 0.249 | 0.114 | 0.172 | 0 | . | . | . | . | . |
| contig_624 | 8517 | 0.226 | 0.41 | 0.105 | 0.088 | 0.171 | 1 | Bcop_v1_g023096-RA | | RhoGAP domain | Rho GTPase-activating protein domain | GO: 0007165 |
| contig_493 | 8910 | 0.041 | 0.243 | 0.265 | 0.305 | 0.146 | 0 | . | . | . | . | . |
| contig_430 | 9615 | 0.023 | 0.273 | 0.337 | 0.277 | 0.091 | 0 | . | . | . | . | . |
| contig_628 | 5217 | 0.085 | 0.199 | 0.488 | 0.141 | 0.087 | 1 | Bcop_v1_g023110-RA | Protein of unknown function | . | . | . |
| contig_372 | 10512 | 0.024 | 0.518 | 0.232 | 0.14 | 0.086 | 2 | Bcop_v1_g023080-RA | Tret1 Facilitated trehalose transporter Tret1 | Sugar (and other) transporter | Major facilitator, sugar transporter-like | GO: 0016021, 0022857, 0055085 |
| | | | | | | | | Bcop_v1_g023081-RA | Protein of unknown function | . | . | . |
| contig_465 | 10768 | 0.089 | 0.241 | 0.437 | 0.154 | 0.08 | 1 | Bcop_v1_g023078-RA | Protein of unknown function | . | . | . |
| contig_312 | 9760 | 0.001 | 0.068 | 0.467 | 0.412 | 0.052 | 0 | . | . | . | . | . |
| contig_314 | 12874 | 0.531 | 0.234 | 0.163 | 0.023 | 0.049 | 3 | Bcop_v1_g023058-RA | Protein of unknown function | . | . | . |
| | | | | | | | | Bcop_v1_g023059-RA | pol RNA-directed DNA polymerase from mobile element jockey | Reverse transcriptase (RNA-dependent DNA polymerase) | Reverse transcriptase domain | - |
| | | | | | | | | Bcop_v1_g023060-RA | Protein of unknown function | - | - | - |
| contig_334 | 13321 | 0.337 | 0.304 | 0.201 | 0.113 | 0.046 | 0 | . | . | . | . | . |
| contig_294 | 8595 | 0.018 | 0.905 | 0.027 | 0.006 | 0.044 | 0 | . | . | . | . | . |
| contig_274 | 11515 | 0.071 | 0.301 | 0.456 | 0.131 | 0.041 | 0 | . | . | . | . | . |
| contig_329 | 16066 | 0.24 | 0.517 | 0.175 | 0.059 | 0.009 | 3 | Bcop_v1_g022985-RA | Protein of unknown function | Ribosomal L22e protein family | Ribosomal protein L22e | GO: 0003735, 0005622, 0005840, 0006412 |
| | | | | | | | | Bcop_v1_g022985-RB | Protein of unknown function | . | . | . |
| | | | | | | | | Bcop_v1_g022986-RA | Usp36 Ubiquitin carboxyl-terminal hydrolase 36 | Ubiquitin carboxyl-terminal hydrolase | Peptidase C19, ubiquitin carboxyl-terminal hydrolase | GO: 0016579, 0036459 |
| contig_547 | 8225 | 0.371 | 0.417 | 0.183 | 0.029 | 0 | 2 | Bcop_v1_g023099-RA | Yp2 Vitellogenin-2 | Lipase | Lipase/vitellogenin | - |
| | | | | | | | | Bcop_v1_g023099-RB | Yp2 Vitellogenin-2 | Lipase | Lipase/vitellogenin | - |
| contig_749 | 7682 | 0.742 | 0.209 | 0.049 | 0 | 0 | 1 | Protein of unknown function | . | . | . | . |
| contig_196 | 12644 | 0.894 | 0.044 | 0.063 | 0 | 0 | 0 | . | . | . | . | . |
| contig_230 | 4409 | 0.011 | 0.701 | 0.288 | 0 | 0 | 0 | . | . | . | . | . |
| contig_462 | 8003 | 0.365 | 0.635 | 0 | 0 | 0 | 1 | Bcop_v1_g023100-RA | Rab3 Ras-related protein Rab-3 | Ras family | Small GTPase | GO: 0003924, 0005525 |
| contig_491 | 10239 | 0.326 | 0.433 | 0.241 | 0 | 0 | 0 | . | . | . | . | . |
| contig_53 | 4458 | 0.909 | 0.091 | 0 | 0 | 0 | 0 | . | . | . | . | . |
| contig_598 | 4301 | 0.002 | 0.053 | 0.945 | 0 | 0 | 0 | . | . | . | . | . |
| contig_665 | 5862 | 1 | 0 | 0 | 0 | 0 | 0 | . | . | . | . | . |
| contig_721 | 11426 | 0.689 | 0.24 | 0.07 | 0 | 0 | 0 | . | . | . | . | . |

#### Supplemental Table S23: Highlighted proteins that aligned to putative Sciara contigs produced by reads that did not map to the Bcop_v1 (Canu) *Sciara* genome assembly.

| Protein Name | Similar to UniProt-SwissProt Protein: | Pfam | InterPro | GO |
|---|---|---|---|---|
| Bcop_v1_g003039-RB | y Protein yellow (Drosophila yakuba OX=7245) | Major royal jelly protein; Protein of unknown function (DUF1091) | Major royal jelly protein/protein yellow; Protein of unknown function DUF1091 | . |
| Bcop_v1_g003673-RA | jockey\pol RNA-directed DNA polymerase from mobile element jockey (Drosophila funebris OX=7221) | Pao retrotransposon peptidase; Reverse transcriptase (RNA-dependent DNA polymerase); Family of unknown function (DUF5641); Protein of unknown function (DUF1759) | Retrotransposon, Pao; Protein of unknown function DUF1759; Reverse transcriptase domain | . |
| Bcop_v1_g004089-RA | CSN1b COP9 signalosome complex subunit 1b (Drosophila melanogaster OX=7227) | 26S proteasome subunit RPN7; PCI domain | 26S proteasome regulatory subunit Rpn7/COP9 signalosome complex subunit 1; Proteasome component (PCI) domain | . |
| Bcop_v1_g006620-RA | zfh2 Zinc finger protein 2 (Drosophila melanogaster OX=7227) | Zinc finger, C2H2 type; Homeodomain | Homeobox domain; Zinc finger C2H2-type | GO: 0003677; 0003676 |
| Bcop_v1_g007612-RA | AKR1E2 1,5-anhydro-D-fructose reductase (Sus scrofa OX=9823) | Aldo/keto reductase family | NADP-dependent oxidoreductase domain | . |
| Bcop_v1_g007613-RA | HEXO2 Beta-hexosaminidase 2 (Arabidopsis thaliana OX=3702) | Glycosyl hydrolase family 20, catalytic domain; beta-acetyl hexosaminidase like | Beta-hexosaminidase, eukaryotic type, N-terminal; Glycoside hydrolase family 20, catalytic domain | GO: 0004553, 0005975 |
| Bcop_v1_g008161-RB | tbpl1 TATA box-binding protein-like protein 1 (Xenopus tropicalis OX=8364) | Transcription factor TFIID (or TATA-binding protein, TBP) | TATA-box binding protein | GO: 0003677, 0006352 |
| Bcop_v1_g010931-RA | wapl Protein wings apart-like (Drosophila melanogaster OX=7227) | Wings apart-like protein regulation of heterochromatin | Wings apart-like protein, C-terminal | . |
| Bcop_v1_g017684-RA | DIP2 Disco-interacting protein 2 (Drosophila melanogaster OX=7227) | AMP-binding enzyme | AMP-dependent synthetase/ligase | GO: 0003824 |
| Bcop_v1_g017710-RA | Shank3 SH3 and multiple ankyrin repeat domains protein 3 (Mus musculus OX=10090) | Ankyrin repeats (3 copies); PDZ domain; N-terminal or F0 domain of Talin-head FERM; Variant SH3 domain | Talin, N-terminal F0 domain; Ankyrin repeat-containing domain; SH3 domain | GO: 0005515 |
| Bcop_v1_g017897-RA | Oaz Zinc finger protein 423 homolog (Drosophila melanogaster OX=7227) | Zinc finger, C2H2 type | Zinc finger C2H2-type | GO: 0003676 |
| Bcop_v1_g020302-RA | IntS8 Integrator complex subunit 8 (Drosophila melanogaster OX=7227) | . | . | . |
| Bcop_v1_g020596-RA | CYP9E2 Cytochrome P450 9e2 (Blattella germanica OX=6973) | Cytochrome P450 | Cytochrome P450 | GO: 0005506, 0016705, 0020037, 0055114 |
| Bcop_v1_g020722-RD | Annulin (Schistocerca americana OX=7009) | Transglutaminase family, C-terminal ig like domain; Transglutaminase-like superfamily; Transglutaminase family | Transglutaminase-like; Transglutaminase, N-terminal; Transglutaminase, C-terminal | GO: 0003810, 0018149; 0018149 |

Short and long read sequences that did not map to the Bcop_v1 assembly were assembled with Canu or Platanus.
Assembled sequences were filtered for those that were identified as Arthropod or that had no BLASTn hits to the entire NT database.
That subset was aligned to the Bcop_v1 protein database using BLASTx (-culling_limit 1 -evalue 1e-6) to find the most significant protein hits over each locus.
The table highlights proteins that either aligned across at least 90% of their lengths or had at least 3 significant BLASTx hits in a row along the length of the protein and contig.
Similar results were obtained when using BLASTx directly against the UniProt-SwissProt protein database.

#### Direct UniProt-SwissProt hits:

```
sp|Q6AZW2|A1A1A_DANRE Alcohol dehydrogenase [NADP(+)] A OS=Danio rerio GN=akr1a1a PE=2 SV=2
sp|Q9H172|ABCG4_HUMAN ATP-binding cassette sub-family G member 4 OS=Homo sapiens GN=ABCG4 PE=1 SV=2
sp|Q96PC3|AP1S3_HUMAN AP-1 complex subunit sigma-3 OS=Homo sapiens GN=AP1S3 PE=1 SV=1
sp|P52183|ANNU_SCHAM Annulin OS=Schistocerca americana PE=2 SV=1
sp|P56377|AP1S2_HUMAN AP-1 complex subunit sigma-2 OS=Homo sapiens GN=AP1S2 PE=1 SV=1
sp|Q9GKY1|B3A4_RABIT Anion exchange protein 4 OS=Oryctolagus cuniculus GN=SLC4A9 PE=1 SV=2
sp|A9ULZ2|BIR7B_XENLA Baculoviral IAP repeat-containing protein 7-B OS=Xenopus laevis GN=birc7-b PE=2 SV=2
sp|Q9VZA4|BND7A_DROME Band 7 protein CG42540 OS=Drosophila melanogaster GN=CG42540 PE=2 SV=2
sp|Q9D6J3|CCD94_MOUSE Coiled-coil domain-containing protein 94 OS=Mus musculus GN=Ccdc94 PE=1 SV=1
sp|O02751|CFDP2_BOVIN Craniofacial development protein 2 OS=Bos taurus GN=CFDP2 PE=1 SV=2
sp|Q9V4I0|CP9B1_DROME Cytochrome P450 9b1 OS=Drosophila melanogaster GN=Cyp9b1 PE=2 SV=1
sp|Q964T2|CP9E2_BLAGE Cytochrome P450 9e2 OS=Blattella germanica GN=CYP9E2 PE=2 SV=1
sp|Q9V4I1|CP9B2_DROME Cytochrome P450 9b2 OS=Drosophila melanogaster GN=Cyp9b2 PE=2 SV=1
sp|Q9VG82|CP9F2_DROME Probable cytochrome P450 9f2 OS=Drosophila melanogaster GN=Cyp9f2 PE=2 SV=1
sp|Q54QX3|CSN1_DICDI COP9 signalosome complex subunit 1 OS=Dictyostelium discoideum GN=csn1 PE=1 SV=1
sp|Q7K4Y6|DAT_DROME Sodium-dependent dopamine transporter OS=Drosophila melanogaster GN=DAT PE=1 SV=1
sp|Q9W0S9|DIP2_DROME Disco-interacting protein 2 OS=Drosophila melanogaster GN=DIP2 PE=1 SV=2
sp|Q9VS29|DSCL_DROME Down syndrome cell adhesion molecule-like protein Dscam2 OS=Drosophila melanogaster GN=Dscam2 PE=2 SV=3
sp|Q9YMJ7|ENV_NPVLD Envelope fusion protein OS=Lymantria dispar multicapsid nuclear polyhedrosis virus GN=LdOrf-130 PE=3 SV=1
sp|P07511|GLYC_RABIT Serine hydroxymethyltransferase, cytosolic OS=Oryctolagus cuniculus GN=SHMT1 PE=1 SV=2
sp|P35623|GLYC_SHEEP Serine hydroxymethyltransferase, cytosolic OS=Ovis aries GN=SHMT1 PE=1 SV=3
sp|P07686|HEXB_HUMAN Beta-hexosaminidase subunit beta OS=Homo sapiens GN=HEXB PE=1 SV=3
sp|O93353|HXA3_CHICK Homeobox protein Hox-A3 OS=Gallus gallus GN=HOXA3 PE=2 SV=2
sp|Q7PRB8|INT3_ANOGA Integrator complex subunit 3 homolog OS=Anopheles gambiae GN=AGAP002539 PE=3 SV=5
sp|O70546|KDM6A_MOUSE Lysine-specific demethylase 6A OS=Mus musculus GN=Kdm6a PE=1 SV=2
sp|O15550|KDM6A_HUMAN Lysine-specific demethylase 6A OS=Homo sapiens GN=KDM6A PE=1 SV=2
sp|Q867Z4|LOLA4_DROME Longitudinals lacking protein, isoforms F/I/K/T OS=Drosophila melanogaster GN=lola PE=1 SV=1
sp|O00370|LORF2_HUMAN LINE-1 retrotransposable element ORF2 protein OS=Homo sapiens PE=1 SV=1
sp|Q921Y0|MOB1A_MOUSE MOB kinase activator 1A OS=Mus musculus GN=Mob1a PE=2 SV=3
sp|Q54XJ0|MOB1A_DICDI MOB kinase activator-like 1 homolog A OS=Dictyostelium discoideum GN=mobA PE=3 SV=1
sp|Q54BM4|MOB1C_DICDI MOB kinase activator-like 1 homolog C OS=Dictyostelium discoideum GN=mobC PE=3 SV=1
sp|Q95RA8|MOB1_DROME MOB kinase activator-like 1 OS=Drosophila melanogaster GN=mats PE=1 SV=1
sp|P16519|NEC2_HUMAN Neuroendocrine convertase 2 OS=Homo sapiens GN=PCSK2 PE=2 SV=2
sp|P21661|NEC2_MOUSE Neuroendocrine convertase 2 OS=Mus musculus GN=Pcsk2 PE=1 SV=1
sp|O61613|NMT_DROME Glycylpeptide N-tetradecanoyltransferase OS=Drosophila melanogaster GN=Nmt PE=2 SV=2
sp|P20241|NRG_DROME Neuroglian OS=Drosophila melanogaster GN=Nrg PE=1 SV=2
sp|Q24741|OSKA_DROVI Maternal effect protein oskar OS=Drosophila virilis GN=osk PE=2 SV=1
sp|Q6PB75|PAPD7_MOUSE Non-canonical poly(A) RNA polymerase PAPD7 OS=Mus musculus GN=Papd7 PE=2 SV=2
sp|Q6L8Q7|PDE12_HUMAN 2',5'-phosphodiesterase 12 OS=Homo sapiens GN=PDE12 PE=1 SV=2
sp|Q08DF7|PDE12_BOVIN 2',5'-phosphodiesterase 12 OS=Bos taurus GN=PDE12 PE=1 SV=1
sp|P09102|PDIA1_CHICK Protein disulfide-isomerase OS=Gallus gallus GN=P4HB PE=1 SV=3
sp|Q03279|PO21_BRACO Retrovirus-related Pol polyprotein from type-1 retrotransposable element R2 (Fragment) OS=Bradysia
coprophila PE=3 SV=1
sp|P10394|POL4_DROME Retrovirus-related Pol polyprotein from transposon 412 OS=Drosophila melanogaster GN=POL PE=3 SV=1
```

sp|P20611|PPAL_RAT Lysosomal acid phosphatase OS=Rattus norvegicus GN=Acp2 PE=1 SV=1
sp|P39847|PPSC_BACSU Plipastatin synthase subunit C OS=Bacillus subtilis (strain 168) GN=ppsC PE=1 SV=2
sp|Q9W0G1|PTP61_DROME Tyrosine-protein phosphatase non-receptor type 61F OS=Drosophila melanogaster GN=Ptp61F PE=1 SV=1
sp|Q3ZBF3|RM38_BOVIN 39S ribosomal protein L38, mitochondrial OS=Bos taurus GN=MRPL38 PE=1 SV=2
sp|P21329|RTJK_DROFU RNA-directed DNA polymerase from mobile element jockey OS=Drosophila funebris GN=jockey\pol PE=1 SV=1
sp|Q5DTL9|S4A10_MOUSE Sodium-driven chloride bicarbonate exchanger OS=Mus musculus GN=Slc4a10 PE=1 SV=2
sp|Q2Y0W8|S4A8_HUMAN Electroneutral sodium bicarbonate exchanger 1 OS=Homo sapiens GN=SLC4A8 PE=1 SV=1
sp|Q95SX7|RTBS_DROME Probable RNA-directed DNA polymerase from transposon BS OS=Drosophila melanogaster GN=RTase PE=2 SV=1
sp|O13134|S4A4_AMBTI Electrogenic sodium bicarbonate cotransporter 1 OS=Ambystoma tigrinum GN=SLC4A4 PE=2 SV=1
sp|P51143|SC6A2_BOVIN Sodium-dependent noradrenaline transporter OS=Bos taurus GN=SLC6A2 PE=2 SV=1
sp|P51905|SC6A4_DROME Sodium-dependent serotonin transporter OS=Drosophila melanogaster GN=SerT PE=2 SV=1
sp|Q6U841|S4A10_HUMAN Sodium-driven chloride bicarbonate exchanger OS=Homo sapiens GN=SLC4A10 PE=2 SV=1
sp|Q924W5|SMC6_MOUSE Structural maintenance of chromosomes protein 6 OS=Mus musculus GN=Smc6 PE=1 SV=1
sp|Q6P4S6|SIK3_MOUSE Serine/threonine-protein kinase SIK3 OS=Mus musculus GN=Sik3 PE=1 SV=3
sp|O43290|SNUT1_HUMAN U4/U6.U5 tri-snRNP-associated protein 1 OS=Homo sapiens GN=SART1 PE=1 SV=1
sp|P28370|SMCA1_HUMAN Probable global transcription activator SNF2L1 OS=Homo sapiens GN=SMARCA1 PE=1 SV=2
sp|Q9Z315|SNUT1_MOUSE U4/U6.U5 tri-snRNP-associated protein 1 OS=Mus musculus GN=Sart1 PE=1 SV=1
sp|Q8JG30|ST1B1_CHICK Sulfotransferase family cytosolic 1B member 1 OS=Gallus gallus GN=SULT1B1 PE=2 SV=1
sp|Q05344|SSRP1_DROME FACT complex subunit Ssrp1 OS=Drosophila melanogaster GN=Ssrp PE=1 SV=2
sp|Q5U385|TBPL1_DANRE TATA box-binding protein-like protein 1 OS=Danio rerio GN=tbpl1 PE=2 SV=1
sp|Q9VCE9|UBS3A_DROME Protein UBASH3A homolog OS=Drosophila melanogaster GN=CG13604 PE=2 SV=1
sp|Q05B21|VGLU1_XENTR Vesicular glutamate transporter 1 OS=Xenopus tropicalis GN=slc17a7 PE=2 SV=1
sp|Q9W517|WAPL_DROME Protein wings apart-like OS=Drosophila melanogaster GN=wapl PE=1 SV=1
sp|Q9TVM2|XPO1_DROME Exportin-1 OS=Drosophila melanogaster GN=emb PE=1 SV=1
sp|Q0KIA2|Y9801_DROME PP2C-like domain-containing protein CG9801 OS=Drosophila melanogaster GN=CG9801 PE=2 SV=1
sp|P28167|ZFH2_DROME Zinc finger protein 2 OS=Drosophila melanogaster GN=zfh2 PE=1 SV=2
sp|Q10126|YSM6_CAEEL Putative uncharacterized transposon-derived protein F52C9.6 OS=Caenorhabditis elegans GN=F52C9.6 PE=5 SV=1
sp|A1Z9R4|ZN423_DROME Zinc finger protein 423 homolog OS=Drosophila melanogaster GN=Oaz PE=2 SV=2

### Section 3: Detailed Experimental Methods

#### 3.1 Embryo Collection

*Sciara* is 'monogenic': the females produce either only male or only female offspring. Male-producing females have 2 copies of the X chromosome while female-producing females have an X and an X'. The two types of females can be differentiated when they are adult flies based on the wavy wing phenotypic marker that is associated with the X': female-producers have wavy wings, whereas male-producers have normal straight wings. However, the different females cannot be distinguished as embryos, our primary target life stage for sequencing. Therefore, we mated males strictly with male-producing females for male-only embryo collection to avoid sequencing the X' in some female embryos that can complicate the assembly. Mass matings consisted of 6 X'X female and 4 XO male flies per vial. One day after combining the flies for mating, females were separated from males. Embryo laying was induced by squishing the thorax with forceps and plating on 2.2% bactoagar (2.2g per 100 mL). After 2-4 hours, adults were removed from the plate and embryos were transferred to an antibiotic/antimycotic plate, where they were incubated at 20-21˚C for up to 2 days, unless early (2-4 hour embryos) were being collected. Prior to DNA extraction, embryos were washed in TE (10 mM Tris-HCl pH 8, 1 mM EDTA) serially 10 times to remove external contamination.

#### 3.2 DNA Extraction

Genomic DNA (gDNA) was isolated from *Sciara* using DNAzol (ThermoFisher) and following the manufacturer's instructions with some modifications. Either a glass dounce homogenizer or a blue pestle was used each with 10 strokes for the homogenization step. Prior to precipitating the DNA, 5 ul RNase A Solution (Qiagen, 4 mg/mL) was added to the DNA sample and incubated at 37˚C for 10 minutes, followed by 5 ul Puregene Proteinase K (Qiagen, 20 mg/mL), incubated at 37˚C for 10 minutes. After adding 100% ethanol, the tube was slowly inverted 50 times, incubated at room temperature for 2 minutes followed by ice for 2 minutes, then centrifuged at 18000g for 10 minutes. The supernatant was removed. The pellet was washed twice with 75% ethanol, very briefly air-dried, and re-suspended in TE. The gDNA was cleaned with AMPure beads (Beckman Coulter) until NanoDrop reported purity levels of A60/280 > 1.8 and A260/230 of ~2.0. It was re-suspended in Tris-HCl (pH 8.0) before beginning library preparations.

#### 3.3 Illumina PE Genomic DNA library

gDNA from mixed stage embryos (2 hour - 2 day old) was sonicated to a size range of 100-600 bp, and the Illumina library were prepared using the NEBNext kit (New England Biolabs) following the manufacturer's directions. The library was run on a 2% NuSieve agarose (Lonza) gel, size-selected near the 500 bp marker, gel purified (Qiagen), and sequenced on the Illumina HiSeq 2000 platform to obtain 100 bp paired-end reads.

#### 3.4 PacBio sequencing details

For Pacific Biosciences RSII Single Molecule Real Time sequencing datasets (P5-C3 chemistry), male embryo gDNA was brought to the Technology Development Group at the Institute of Genomics & Multiscale Biology at the Icahn School of Medicine at Mount Sinai for library construction and sequencing. There, two DNA libraries were prepared and sequenced according to the manufacturer's instructions, reflecting the P5-C3 sequencing enzyme and chemistry, respectively. For each library, 6 µg of extracted, high-quality, genomic DNA isolated from *Sciara coprophila* was diluted in Qiagen elution buffer to 150 µL. The 150 µL aliquots were individually pipetted into the top chambers of Covaris G-tube spin columns and sheared gently for 60 seconds at 4500 rpm using an Eppendorf 5424 benchtop centrifuge and repeated at 60 seconds at 4500 rpm to further shear the DNA and place the aliquot back into the upper chamber, resulting in a ~20,000 bp DNA shear, verified using a DNA 12000 Agilent Bioanalyzer gel chip. The sheared DNA was then re-purified using a 0.45X AMPure XP purification step (0.45X AMPure beads added, by volume, to each DNA sample dissolved in 200 µL EB, vortexed for 10 minutes at 2,000 rpm, followed by two washes with 70% alcohol and finally diluted in EB).

After purification, 2.7 µg of purified and sheared sample was taken into the DNA damage and end-repair steps. Briefly, the DNA fragments were repaired using DNA Damage Repair solution (1X DNA Damage Repair Buffer, 1X NAD+, 1 mM ATP high, 0.1 mM dNTP, and 1X DNA Damage Repair Mix) with a volume of 21.1 µL and incubated at 37˚C for 20 minutes. DNA ends were repaired next by adding 1X End Repair Mix to the solution, which was incubated at 25˚C for 5 minutes followed by the second 0.45X Ampure XP purification step. Next, 0.75 µM of Blunt Adapter was added to the DNA followed by 1X template Prep Buffer, 0.05 mM ATP low and 0.75 U/µL T4 ligase to ligate (final volume of 47.5 µL) the SMRTbell adapters to the DNA fragments. This solution was incubated at 25˚C overnight followed by a 65˚C 10-minute ligase denaturation step. After ligation, the library was treated with an exonuclease cocktail to remove un-ligated DNA fragments using a solution of 1.81 U/µL Exo III 18 and 0.18 U/µL Exo VII, then incubated at 37˚C for 1 hour. Two additional 0.45X Ampure XP purifications steps were performed to remove <2000 bp molecular weight DNA and organic contaminant.

Upon completion of library construction, the library was validated as ~20 kb using another Agilent DNA 12000 gel chip. All libraries were sufficient for additional size selection to remove any library molecules < 7,000 bp. This step was conducted using Sage Science Blue Pippin 0.75% agarose cassettes to select library in the range of 7,000-50,000 bp. 16% of the input library eluted from the agarose cassette and was available for sequencing. This yield was sufficient to proceed to primer annealing and DNA sequencing on the PacBio RSII machine. Size-selection was confirmed by Bio-Analysis and the mass was quantified using Qubit. Primer was then annealed to the size-selected SMRTbell with the full-length libraries (80˚C for 2 minute 30 seconds followed by decreasing the temperature by 0.1˚ to 25C˚). The polymerase-template complex was then bound to the P5 enzyme using a ratio of 10:1 polymerase to SMRTbell at 0.5 nM for 4 hours at 30˚C and then held at 4˚C until ready for magbead loading, prior to sequencing. The magnetic bead-loading step was conducted at 4˚C for 60-minutes per manufacturer's guidelines. The magbead-loaded, polymerase-bound, SMRTbell libraries were placed onto the RSII machine at a sequencing concentration of 75-100 pM across 24 SMRTcells and configured for 180-minute continuous sequencing runs on each SMRTcell. Sequencing was conducted to ample coverage across 24 SMRTcells.

#### 3.5 MinION sequencing details

We followed similar protocols as reported in our earlier bioRxiv preprint (Urban et al. 2015). There we described our first attempts at sequencing *Sciara* genomic DNA using modifications to the standard protocol that required no additional reagents or equipment and reported that the majority of our data came from reads > 10 kb with many exceeding 50 kb and a high quality 2D read that exceeded 100 kb. In addition to those early datasets, our MinION data were collected over the course of 18 months and span multiple kits (MAP002-MAP006), flow cell/pore (R7.3 – R7.3 70bps 6mer), software, and MinION upgrades (original, MkI). As the ONT protocols changed, we needed to adapt our modifications from that preprint. However, the principles and results of our protocols have stayed the same (e.g. see Supplemental Figures S4 and S5).

The principles we apply to our modified protocols are the following:

(1) Start out with more DNA than required for the standard protocol that assumes 8 kb molecules to target similar molarities,

(2) Skip the Covaris shearing step to keep DNA long,

(3) Always perform a DNA repair step to repair damaged bases and single-stranded nicks,

(4) Use wide-bored tips and very gentle pipetting throughout the protocol,

(5) Use 0.4x ratio of AMPure beads in all clean-up steps,

(6) Add a rinse step before elution of AMPure beads to deplete DNA < 10-12 kb in all clean-up steps, and

(7) Elute DNA off the AMPure beads while adding heat into the system (37-50˚C) for extended periods of time (10-20 minutes) in all clean-up steps.

(8) It is also important to minimize the amount of handling needed, which has been facilitated by updates to the standard protocol by combining End-Repair and dA-tailing into one step.

**Reagents, MinION versions, pore models, and protocols for each library:**

In total, we prepared 17 libraries for this project. The libraries are named/numbered 01-09, 11-16, 20-21. In general, library preparations have the general workflow of optional shearing, DNA repair, End repair, dA-tailing, adapter ligation, and hairpin enrichment in some cases with various clean up steps in between.

Library 01 used SQK-MAP002 reagents. Libraries 02-08 used SQK-MAP004 reagents. Libraries 09, 11, 12, and 13 used SQK-MAP005 reagents. All aforementioned libraries used the original MinION and the R7.3 pore model. Libraries 14, 15, 16, 20, and 21 used SQK-MAP006 reagents, the MinION MkI, and the R7.3 70 bps 6mer model. Libraries 14, 20, and 21 were constructed following Oxford Nanopore's standard protocol. Libraries 02-09, 11-13, and 15-16 were constructed using modifications from above as described below.

**Optional Shearing:**

Libraries 14, 20, and 21 followed the standard protocol, and shearing to ~8 kb was performed in Covaris g-TUBEs as instructed.

For libraries prepared with modified protocols (01-09, 11-13, 15-16), shearing was always skipped and we started out with 2.2-17 µg, using the lower end for early libraries (2.2-5 µg in 01-13) and the higher amounts more recent libraries (15-17 µg in 15-16).

**DNA Repair:**

All libraries except 01 were subject to DNA repair steps. PreCR (NEB) was performed on libraries 02-09 and 11-13. FFPE Repair (NEB) was performed on libraries 14-16 and 20-21.

**End repair and dA-tailing:**

For most early libraries prior to MAP006 (excluding 01 and 03), End-Repair and dA-tailing were done in separate steps using the NEBNext End-Repair Module (NEB) and the NEBNext dA-Tailing Module (NEB). For libraries 01 and 03, these steps were combined using the NEBNext Ultra End Repair/dA-tailing module (NEB). For all libraries using MAP006 MinION reagents, end repair and dA-tailing were performed as a single step using the NEBNext Ultra II End Repair/dA-tailing module (NEB).

**Adapter ligation:**

Ligation was always carried out using MinION kit-specific adapters and volumes with Blunt/TA Ligase Master Mix (NEB). We only needed to manually add the tether and motor proteins for library 01. In all others, the tether and motor proteins were pre-combined with other reagents such as the adapters and elution buffer. The first library did not have a hairpin enrichment step, but all other libraries did concurrent with MinION kits and protocols.

**Hairpin enrichment:**

Seven libraries (02-08) used His-beads (Dynabeads His-tag Isolation and Pulldown; Life Technologies) with MAP004 reagents and four libraries (09, 11-13) used His-beads with MAP005 reagents according to Oxford Nanopore's instructions. The five MAP006 libraries (14-16, 20-21) used MyOne C1 streptavidin beads (Dynabeads/Thermofisher) for hairpin enrichment following Oxford Nanopore's protocol.

**AMPure clean up notes:**

Libraries 07, 08, 09, and 15 included rinses in all clean-up steps (Urban et al. 2015) whereas the other libraries did not. Clean up steps for all modified protocols used a 0.4x AMPure ratio. For all AMPure beads steps for our modified protocols, DNA was eluted off the beads by incubating at 37˚C for 20 minutes in libraries 01-09 and 11-13, and at 50˚C for 10 minutes in libraries 15-16. For standard protocols, DNA was eluted off beads following Oxford Nanopore's protocol. Wide-bored tips and gentle pipetting were used throughout.

**Sequencing notes:**

Sequencing was conducted following standard procedures. For early libraries, smaller amounts of DNA were loaded more frequently (e.g. 4 times) throughout the run. For more recent libraries, half of the library was loaded at the beginning and the second half was added 24 hours in.

**Performance notes:**

Libraries 04, 05, 09, and 11 did not perform well due to flow cells with very limited numbers of pores available for sequencing and consequentially gave little data, but we included them anyway.

Library 11 was attempted on two different flow cells that both started with very few available pores. Moreover, though fragmentation was not performed, the DNA was lower molecular weight than expected as viewed on an agarose gel.

Libraries 06, 07, 08, 09, 12, 13 and 14 were all prepared from the same DNA source to be able to directly compare size distributions after different protocols. Similarly, libraries 15 and 16 were prepared from the same source to make direct comparisons of the effects of including rinse steps in the AMPure clean-ups. When comparing modified protocols without rinse steps to the standard protocols, our modified protocols were enriched for reads >10 kb. Modified protocols that also included rinse steps were additionally enriched over all other protocols for reads >10 kb.

The standard protocol for three libraries gave a combined molecule N50 of 6.9 kb and 2D read N50 of 8.8 kb in agreement with the target of 8 kb molecules. Our modified protocols produced molecule N50s and 2D N50s of 19.3 kb and 18.4 kb respectively, over double that from the standard protocol. MarginStats (Jain et al. 2015) was used to calculate the percent identities of MinION reads aligned to a PacBio genome assembly (details in bioinformatics section below). We obtained ultra-long (>100 kb) MinION reads map to PacBio-only genome assemblies with up to 91% identity across the entire length. For example, we obtained 2D reads of lengths 131.5 kb (Q=10.3), 111.2 kb (Q=9.9), 105 kb (Q=9.3), and 100.7 kb (Q=10.4) that aligned across their full lengths with identities of 91.1%, 88.7%, 63.2%, and 84.2%. The 102.9 kb 2D read that we previously reported as high quality (Q=8.74) (Urban et al. 2015) aligned in full at 84.2% identity, representing a high quality, ultra-long 2D read derived from early MinION sequencing reagents

(MAP004). For all nanopore reads, 85.7% of 2D reads and 57.9% of 1D reads aligned to a high quality PacBio-only assembly from Canu (Koren et al. 2017). The median percent identities for aligned 1D and 2D reads was 68% and 82.1%, respectively, with both having examples with >95% identity. Percent identities for both 2D and 1D reads were correlated with mean quality scores (Spearman's rho for 2D = 0.76 and for 1D = 0.70). There were some exceptionally long 1D reads that exceed 200 kb, but those were strictly of low quality and low percent-identity.

#### 3.6 BioNano Irys optical mapping details

To isolate ultra high molecular weight DNA, male pupae were flash frozen and ground in liquid nitrogen. The powder was resuspended in Nuclear Isolation Buffer (NIB)(10 mM Tris pH 9.4, 60 mM NaCl, 10 mM EDTA, 0.15 mM spermidine, 0.15 mM spermine 0.5% Triton-X 100, 1% beta-mercaptoethanol) on ice and filtered through 100 µM mesh cell strainer. Cellular debris was removed by sedimentation for 15 sec at 1K rpm, 4˚C. Nuclei were recovered from the supernatant by sedimentation for 3 minutes at 1800xg, 4˚C. Nuclei washed 3x with NIB, were resuspended in Cell Suspension Buffer (Chef Mammalian Genomic DNA plug kit, Bio Rad) and embedded in low melt agarose (BioRad), final concentration 0.8%. Nuclei were lysed in Alternate Lysis Buffer (BioNano Genomics) containing 1.6 mg Proteinase K (Qiagen) at 50˚C for 24 hr, and RNA degraded by addition of RNAse A (80 µg, Qiagen). Plugs were liquefied using Gelase (Epicentre) and the resulting high molecular weight DNA was membrane dialyzed against 10mM Tris HCL, 1mM EDTA, pH 8.0). DNA was harvested with wide bore tips and stored at 4˚C.

High Molecular Weight (HMW) DNA was nicked, labeled and repaired according to the IrysPrep protocol (BioNano Genomics). In brief, HMW DNA was digested with the single-stranded nicking endonuclease BssSI (CACGAG, NewEngland BioLabs). Fluorescently labeled nucleotides were incorporated by nick translation. The backbone of the labeled DNA was stained with YOYO-1. Individual labeled DNA molecules were imaged on the Irys platform (BioNano Genomics). Images of 438,139 molecules >100 kb (N50 = 173.8 kb) were collected: 217,194 of these were >150 kb (N50 = 214.1 kb).

#### 3.7 Strand-specific RNA-seq details

Total RNA from male and female embryos, larvae, pupae, and adult flies was extracted using TRIzol (Invitrogen/Thermofisher). RNA quantity and purity were measured with the NanoDrop (ThermoScientific) and Qubit (ThermoFisher). Total RNA was treated with DNase (Qiagen) and subject to RNeasy column clean up (Qiagen). The cleaned total RNA quantity and purity were checked using the NanoDrop and Qubit. RNA integrity was evaluated on 1.1% formaldehyde 1.2% agarose gels. Poly-A RNA was enriched using Oligo-dT DynaBeads (LifeTechnologies). Qubit was used to measure the quantity of poly-A RNA. Poly-A RNA was fragmented with NEB's Magnesium Fragmentation Module for 3 minutes at 94˚C, which was selected after optimizing for conditions for 200-500 bp fragments as determined by the Bioanalyzer. Fragmentation reactions were cleaned up with RNeasy columns. First strand synthesis was performed with SSIII (Invitrogen). Briefly, 1 µl Random Primer (3 µg/µl), 1 µ 10 mM dNTP mix, and 10 µl fragmented RNA were incubated at 65˚C for 5-10 minutes and transferred to ice for 5 minutes. A mix of 4 µl 5X First Strand Synthesis (FSS) Buffer, 1 µl 0.1 M DTT, 1 µl Murine RNase Inhibitor, 1 µl 0.5 µg/µl Actinomycin D, and 1 µl SSIII (200 units) was added to the mixture of RNA, dNTPs, and Random Primers. This was incubated in the thermocycler at 25˚C for 5 minutes to anneal the random primers, 50˚C for 60 minutes to extend from the random primers, and 70˚C for 15 minutes to inactivate SSIII. The reaction was cleaned up with AMPure beads using a ratio of 2.0.

For Second Strand Synthesis (SSS), a mixture of 10 mM each of dATP, dCTP, dGTP, and 20 mM of dUTP (instead of dTTP) was made. For a single reaction, 64 µl of cleaned FSS cDNA:RNA in Ultra Pure Water, was combined with 4 µl ACGU mix, 8 µl of NEB dNTP-free SSS Reaction buffer and 4 µl NEB SSS Enzyme mix from the SSS module. The reaction was incubated for 1 hour at 16˚C, then cleaned with AMPure beads using a 1.0x ratio to begin eliminating DNA smaller than 200 bp, and quantified with the Qubit. There was typically 100-200 ng at this step.

The double-stranded cDNA was then subject to End Repair (NEBNext), and cleaned with AMPure beads using a 0.9x ratio to deplete DNA < 300 bp. The End-Repaired cDNA was then dA-tailed (NEBNext), and cleaned with AMPure beads using a 0.9x ratio. For adapter ligation, 38 µl of DNA was combined with 10 µl 5X NEBNext Quick Ligation Reaction Buffer, 1 µl NEB Adaptor, and 2 µl Quick T4 Ligase. The reaction was incubated at room temperature for 15 minutes, then cleaned with a 0.9x ratio of AMPure beads.

The library was then size-selected with AMPure beads before proceeding to the PCR step. To obtain adapter-ligated fragments in the 300-600 bp range, the DNA was first incubated with 0.6x AMPure beads by adding 60 µl AMPure beads to 100 µl DNA in UPW. The beads were pelleted on a magnet and the supernatant containing DNA smaller than approximately 600 bp was transferred to a new tube (DNA longer than 600 bp stayed on the beads). To make the final ratio 0.9x to select for DNA > 300 bp on the beads, 30 µl more AMPure beads was added to the 160 µl supernatant. From there the AMPure clean-up proceeded as normal.

USER enzyme digestion to cut the DNA at uracils (in the second strand and in the hairpin adapters) and PCR were then performed as follows: 20 µl of cleaned DNA was combined with 25 µl NEBNext High-Fidelity 2X PCR Master Mix and 3 µl NEBNext USER enzyme. This reaction was incubated for 15 minutes at 37˚C to ensure uracil cutting occurs before addition of primers. Then 1 µl indexed primer (NEBNext) and 1 µl universal primer were added. The reaction was put in the thermocycler for 37˚C for 15 minutes to ensure USER digestion went to completion, followed by 98˚C for 30 seconds and 12 cycles of: 98˚C for 10 seconds, 65˚C for 30 seconds, 72˚C for 30 seconds. The PCR products at this stage are approximately 122 bp longer than the target insert size. We adjusted AMPure ratios accordingly for a final clean up and size-selection.

The PCR reactions were cleaned with 0.85x AMPure beads to deplete DNA smaller than 350 bp. The DNA was eluted in 100 µl UPW and AMPure size selection was initiated by incubating with 55 µl AMPure beads (0.55x) to precipitate DNA longer than 650 bp onto the beads. The beads were pelleted on a magnet and the 155 ul of DNA shorter than 650 bp in the supernatant was transferred to a new tube where another 30 µl of AMPure beads was added for a final ratio of 0.85x. The AMPure procedure then continued as normal to obtain DNA > 350 bp. The estimated insert sizes at this step was 230-530 bp. DNA samples were quantified with Qubit and purity was measure by NanoDrop. There was typically 600 ng at the end of this protocol. Bioanalyzer traces suggested the mean estimated fragment sizes was around 420 bp putting the mean insert sizes near 300 bp.

### Section 4: Detailed Bioinformatics Methods

Note: for specific software versions used, see Section 5 below.

#### 4.1 Illumina Assemblies

We obtained ~103x coverage of paired-end Illumina HiSeq 2000 data from a ~430 bp insert genomic DNA library. This data was assembled using several popular genome assemblers: ABySS (Simpson et al. 2009), Megahit (Li et al. 2015), Platanus (Kajitani et al. 2014), SGA (Simpson and Durbin 2010), SOAPdenovo2 (Luo et al. 2012), SPAdes (Bankevich et al. 2012), and Velvet (Zerbino and Birney 2008). We also attempted to use MaSuRCA (Zimin et al. 2013). However, after several attempts that failed due to exceeding allotted time limits, with the final attempt given >8 days, we decided to not pursue it further. For all other assemblers, we tried assembling the raw data as well as a read set that was quality-filtered using Trimmomatic (Bolger et al. 2014). Moreover, SPAdes and SGA both error-correct the input reads followed by assembling the error-corrected reads. Therefore, we also tried the error-corrected raw reads and error-corrected quality-filtered reads produced by BayesHammer (Nikolenko et al. 2013) in the SPAdes pipeline with the other assemblers. Finally, some assemblers, such as Megahit, Platanus, and SPAdes, employ pipelines that iterate over multiple k-mer sizes whereas others, such as Velvet, SOAPdenovo2, and Abyss, require a value of K to be selected. Therefore, we tried K=55 and K=77 for each set of reads for the latter set of assemblers.

##### 4.1.1 Inputs

For most assemblers, we tried using all reads, all reads after BayesHammer (Nikolenko et al. 2013) from SPAdes (Bankevich et al. 2012) correction, filtered reads using Trimmomatic (Bolger et al. 2014), and filtered reads that were subsequently corrected with BayesHammer. Note that the BayesHammer corrected reads were produced as part of running SPAdes.

###### 4.1.1.1 Trimming and quality filtering

For trimming, we used Trimmomatic version 0.32:

```
IN="MSF0007_TGACCA_L004_R1_001.fastq.gz
MSF0007_TGACCA_L004_R2_001.fastq.gz"
OUTPRE=maleSciaraTrimmed
OUT="${OUTPRE}_forward_paired.fq.gz ${OUTPRE}_forward_unpaired.fq.gz
${OUTPRE}_reverse_paired.fq.gz ${OUTPRE}_reverse_unpaired.fq.gz"
PRIMERS=neb_primers.fa
MINQUAL=5
MINLEN=100

java -jar $TRIMMOMATIC_BASE/Trimmomatic-0.32/trimmomatic-0.32.jar PE -
trimlog trim.log $IN $OUT ILLUMINACLIP:${PRIMERS}:2:30:10
LEADING:$MINQUAL TRAILING:$MINQUAL SLIDINGWINDOW:4:$MINQUAL
MINLEN:$MINLEN
```

Trimming produced three files:
One file each for the forward and reverse reads that remained paired and one file containing the orphaned reads. In addition to the paired reads, the orphaned reads were used with most assemblers.

###### 4.1.1.2 Short read error-correction

SPAdes runs BayesHammer error-correction as part of its pipeline. The error-corrected reads are stored in three files: forward reads (R1), reverse reads (R2), and orphaned reads (SE). We used these error-corrected reads with other assemblers as described.

###### 4.1.1.3 Naming of short read datasets used in assemblies

Where relevant below:

R1 = Left read/read_1 from paired-end (PE) reads
R2 = Right read/read_2 from PE reads
SE = reads that were orphaned during quality filtering or error-correction
q5 = Trimmomatic quality-filtered
bh = BayesHammer error-corrected
cf = Contamination filtered (cf)
gz = gzipped extension
fq = altenrative suffix for fastq

<u>Short Read Set #1</u>
R1.fastq(.gz) and R2.fastq(.gz)
= the set of all PE reads - no quality filtering and no error-correction

<u>Short Read Set #2</u>
R1.q5.fastq(.gz), R2.q5.fastq(.gz), and SE.q5.fastq(.gz)
= the quality-filtered reads from Trimmomatic

<u>Short Read Set #3</u>
R1.bh.fastq, R2.bh.fastq, and SE.bh.fastq
= the BayesHammer-corrected reads (using all reads and no quality filtering)

<u>Short Read Set #4</u>
R1.q5.bh.fastq, R2.q5.bh.fastq, and SE.q5.bh.fastq
= the BayesHammer-corrected quality-filtered reads (quality-filtering and bayeshammer orphans all in same SE file)

<u>Short Read Set #5</u>
R1.cf1.fastq and R2.cf1.fastq
= the set of all reads after contamination filtering

<u>Short Read Set #6</u>
R1.cf1.q5.fastq, R2. cf1.q5. fastq, and SE. cf1.q5.fastq
= the set of all reads after contamination filtering that were subsequently quality-filtered

<u>Short Read Set #7</u>
R1.cf1.bh.fastq, R2. cf1.bh.fastq, and SE.cf1.bh.fastq
= the set of all reads after contamination filtering that were subsequently error-corrected

<u>Short Read Set #8</u>
R1.cf1.q5.bh.fastq, R2.cf1.q5.bh.fastq, and SE.cf1.q5.bh.fastq

= the set of all reads after contamination filtering that were subsequently quality-filtered, then error-corrected

###### 4.1.1.4 Insert size statistics

Where relevant below, mean (432), min (83) and deviation of insert sizes were obtained using Picard-Tools CollectInsertSizeMetrics (http://broadinstitute.github.io/picard) after mapping the paired-end reads to an early long read assembly with Bowtie2 (Langmead and Salzberg 2012).

##### 4.1.2 Assembling the short reads

A total of 44 short read assemblies were reported on in this study: 40 prior to initial evaluations and contamination analysis and filtering, and 4 thereafter using the top performing assembler given the evaluations.

###### 4.1.2.1 ABYSS

A total of 8 Abyss assemblies were made from Short Read Sets #1-4 (commands for each represented below), each with two different kmer sizes.

For all: K=k, where k was either 55 or 77

```
abyss-pe name=Sciara j=16 k=${k} in='R1.fastq  R2.fastq'

abyss-pe name=Sciara j=16 k==${k} lib='pe432' pe432='R1.q5.fastq
R2.q5.fastq' se='SE.q5.fastq'

abyss-pe name=Sciara j=16 k==${k} lib='pe432' pe432='R1.bh.fastq.gz
R2.bh.fastq.gz' se='SE.bh.fastq.gz'

abyss-pe name=Sciara j=16 k==${k} lib='pe432' pe432='R1.q5.bh.fastq.gz
R2.q5.bh.fastq.gz' se='SE.q5.bh.fastq.gz'
```

###### 4.1.2.2 Megahit

Megahit was run on Short Read Sets #1-4, generating four assemblies. Megahit iterates over kmer sizes such that no specific kmer size need be provided. It was run on Short Read Sets #1-4 as follows.

```
megahit -1 R1.fastq -2 R2.fastq

megahit -1 R1.q5.fastq -2 R2.q5.fastq -r SE.q5.fastq

megahit -1 R1.bh.fastq -2 R2.bh.fastq -r SE.bh.fastq

megahit -1 R1.q5.bh.fastq -2 R2.q5.bh.fastq -r SE.q5.bh.fastq
```

###### 4.1.2.3 Platanus

Platanus was first run on Short Read Sets #1-4, generating 4 assemblies. Platanus iterates over kmer sizes such that no specific kmer size need be provided. Evaluations showed that Platanus was a top performing short read assembler given our data, so we subsequently used Platanus with the contamination filtered read sets as well (Short Read Sets #5-8). For all Short Read Sets (#1-8) described in section 4.1.1.3, the commands were as represented below.

```
PREFIX=out
contig=${PREFIX}_contig.fa
bubble=${PREFIX}_contigBubble.fa
kmerfreq=${PREFIX}_kmerFrq.tsv
scaffold=${PREFIX}_scaffold.fa
scafbubble=${PREFIX}_scaffoldBubble.fa
scafcomp=${PREFIX}_scaffoldComponent.tsv
T=16
M=45
PE1 = R1 fastq file for given read set
PE2 = R2 fastq file for given read set
SE = orphaned reads from given read set if applicable

platanus assemble -f $PE1 $PE2 [$SE] -o $PREFIX -t $T -m $M
platanus scaffold -c $contig -b $bubble -IP1 $PE1 $PE2 -n1 83 -a1 432 -
d1 21 -t $T -o $PREFIX
platanus gap_close -o $PREFIX -c $scaffold -IP1 $PE1 $PE2 -t $T
```

###### 4.1.2.4 SGA

SGA has its own error-correction so only the raw reads and quality-filtered reads were used with it (i.e. Short Read Sets #1-2).

```
SGA -- unfiltered:
#SAMtools 0.1.19 and ABYSS 1.9.0 in PATH.
PE1=R1.fastq
PE2=R2.fastq
See SGA recipe below.

SGA -- filtered:
#SAMtools 0.1.19 and ABYSS 1.9.0 in PATH.
PE1=R1.q5.fastq
PE2=R2.q5.fastq
See SGA recipe below.
Note: SE.q5.fastq not used.

SGA recipe -- same for unfiltered and filtered reads:
# Largely drawn from example here:
https://github.com/jts/sga/blob/master/src/examples/sga-celegans.sh

# SGA has its own read correction, so did not use with BayesHammer
corrected reads

NAME=Sciara-male
OL=75
T=16
D=4000000
CK=51
COV_FILTER=2
MOL=55
R=10
MIN_PAIRS=5
MIN_LENGTH=200
SCAFFOLD_TOLERANCE=1
MAX_GAP_DIFF=0
```

```
CTGS=assemble.m$OL-contigs.fa
GRAPH=assemble.m$OL-graph.asqg.gz
SGA_ALIGN=~/software/sga/sga/src/bin/sga-align

sga preprocess --pe-mode 1 $PE1 $PE2 > $NAME.fastq

sga index -a ropebwt --no-reverse -t $T $NAME.fastq

sga preqc -t $T ${NAME}.fastq > $NAME.preqc

sga correct -k $CK --discard --learn -t $T -o ${NAME}.ec.k$CK.fastq
${NAME}.fastq

sga index -a ropebwt -t $T ${NAME}.ec.k$CK.fastq

sga filter -x $COV_FILTER -t $T --homopolymer-check --low-complexity-
check ${NAME}.ec.k$CK.fastq

sga fm-merge -m $MOL -t $T -o ${NAME}.merged.k$CK.fa
${NAME}.ec.k$CK.filter.pass.fa

sga index -d 1000000 -t $T ${NAME}.merged.k$CK.fa

sga rmdup -t $T ${NAME}.merged.k$CK.fa

sga overlap -m $MOL -t $T ${NAME}.merged.k$CK.rmdup.fa

sga assemble -m $OL -g $MAX_GAP_DIFF -r $R -o assemble.m$OL
${NAME.merged.k$CK.rmdup.asqg.gz

sga assemble -m $OL -g $MAX_GAP_DIFF -r $R -o assemble.m$OL
${NAME}.merged.k$CK.rmdup.asqg.gz

$SGA_ALIGN --name ${NAME}.pe $CTGS $PE1 $PE2

sga-bam2de.pl -n $MIN_PAIRS --prefix libPE ${NAME}.pe.bam

sga-astat.py -m $MIN_LENGTH ${NAME}.pe.refsort.bam > libPE.astat

sga scaffold -m $MIN_LENGTH --pe libPE.de -a libPE.astat -o
scaffolds.n$MIN_PAIRS.scaf $CTGS

sga scaffold2fasta -m $MIN_LENGTH -a $GRAPH -o scaffolds.n$MIN_PAIRS.fa
-d $SCAFFOLD_TOLERANCE --use-overlap --write-unplaced
scaffolds.n$MIN_PAIRS.scaf
```

###### 4.1.2.5 SOAPdenovo2

A total of 8 SOAPdenovo2 assemblies were made from Short Read Sets #1-4 (commands for each represented below), each with two different kmer sizes (K=55 or K=77).

```
PE1=R1.fastq
PE2=R2.fastq

Or

PE1=R1.q5.fastq
PE2=R2.q5.fastq
SE=SE.q5.fastq
```

```
Or

PE1=R1.bh.fastq
PE2=R2.bh.fastq
SE=SE.bh.fastq

Or

PE1=R1.q5.bh.fastq
PE2=R2.q5.bh.fastq
SE=SE.q5.bh.fastq
```

Config file for all:

```
#For example config file: http://soap.genomics.org.cn/soapdenovo.html
#maximal read length
max_rd_len=100

 [LIB]
#average insert size
avg_ins=432
#if sequence needs to be reversed
reverse_seq=0
#in which part(s) the reads are used
asm_flags=3
#use only first 100 bps of each read
rd_len_cutoff=100
#in which order the reads are used while scaffolding
rank=1
# cutoff of pair number for a reliable connection (at least 3 for short
insert size)
pair_num_cutoff=3
#minimum aligned length to contigs for a reliable read location (at
least 32 for short insert size)
map_len=32

#a pair of fastq file, read 1 file should always be followed by read 2
file
q1=${PE1} ## dependent on which set of reads being used
q2=${PE2}  ## dependent on which set of reads being used

#fastq file for single reads
q=${SE} ## dependent on which set of reads being used (some do not
have)
```

Commands:

```
SOAPdenovo-127mer pregraph -s $config_file -K $K -R -o $graph_prefix -p
$P
SOAPdenovo-127mer contig -g $graph_prefix -R -p $P
SOAPdenovo-127mer map -s $config_file -g $graph_prefix -p $P
SOAPdenovo-127mer scaff -g $graph_prefix -F -p $P
```

###### 4.1.2.6 SPAdes

The inputs to SPAdes were Short Read Sets #1-2. Since SPAdes used BayesHammer error-correction by default prior to assembly, this actually means it was run only on Short Read Sets #3-4. For a given read set, we error-corrected the input reads (SRS#1 or #2) in only one of the SPAdes runs and subsequently used the error-corrected reads (SRS #3 or #4) produced in that run with the “--only-assembler” option in subsequent runs. We tried SPAdes in three different ways for each set (see commands below), generating 6 SPAdes assemblies.

[square brackets only applicable when single-end read file used]

```
spades.py --pe1-1 $PE1 --pe1-2 $PE2 [--pe1-s $SE] -o default -t 64 --
only-assembler --cov-cutoff auto

spades.py -k 21,33,55,77 --pe1-1 $PE1 --pe1-2 $PE2 [--pe1-s $SE] -o
k2177 -t 64

spades.py -k 21,33,55,77 --pe1-1 $PE1 --pe1-2 $PE2 [--pe1-s $SE] -o
k2177auto -t 64 --cov-cutoff auto

## Allowing SPAdes to pick kmer size values automatically (it chose
K=21,33,55)
spades.py --careful --pe1-1 $PE1 --pe1-2 $PE2 [--pe1-s $SE] -o k213355
-t 64 [--only-assembler]

## Specifying kmer size values to go up to k=77 (K=21,33,55,77)
spades.py --careful -k 21,33,55,77 --pe1-1 $PE1 --pe1-2 $PE2 [--pe1-s
$SE] -o k21335577 -t 64 [--only-assembler]

## Specifying kmer size values to go up to k=77 (K=21,33,55,77) and
enabling the automatic coverage cutoff.
spades.py --careful -k 21,33,55,77 --pe1-1 $PE1 --pe1-2 $PE2 [--pe1-s
$SE] -o k21335577auto -t 64 --cov-cutoff auto [--only-assembler]
```

###### 4.1.2.7 Velvet

Velvet was run on Read Sets #1-4 using two different kmer sizes each (K=55 or K-77), generating 8 assemblies.

[square brackets only applicable when single-end read file used]

```
K=55

Or

K=77

DIR=hash${K}

velveth ${DIR} ${K} -fmtAuto -shortPaired -separate $R1 $R2 [-short
$SE]

velvetg ${DIR} -clean yes -exp_cov auto -cov_cutoff auto
```

#### 4.1.3 Evaluations of the Initial 40 Short Read Assemblies

To evaluate the short read assemblies, we used the percentage of mapped Illumina reads, probabilisitic scores from LAP (Ghodsi et al. 2013) and ALE (Clark et al. 2013), the number of potential misassembled regions reported by $FRC^{bam}$ (Vezzi et al. 2012), the percentage of error-free bases (or mean base scores) from REAPR (Hunt et al. 2013), and BUSCO (Simão et al. 2015). Metrics from LAP, ALE, FRC, and REAPR reflect the consistency of each assembly with the Illumina dataset and BUSCO estimates the completeness of each assembly in terms of gene content. Note that the percentage of mapped reads was a great predictor of how assemblies would be ranked by LAP and ALE.

LAP and ALE provide probability measures that a given set of reads came from each assembly. LAP requires finding all mappings for each read, which can be time-consuming and demanding of computational resources. However, the authors note that fairly small sample sizes of reads tend to correlate well with larger samples for eukaryotic genome assemblies, which can drastically differ from one assembler to the next (Ghodsi et al. 2013). Therefore, we first tried two independent samples of ~15,000 paired reads, followed by a sample of ~150,000, and finally ~1.5 million. All samples largely agreed with each other and with the other metrics, so we used the final sample size of 1.5 million.

$FRC^{bam}$ flags potential errors (called features) throughout each assembly given a set of reads, with lower numbers being better than higher ones. We also checked to see if normalizing to the number of features per megabase changed the conclusions, but found that it was highly correlated with the total number of features (Spearman's rho = 0.88) and that the rank leaders remained the same. Therefore, we used only the number of features in calculating mean rankings.

REAPR outputs the percent of bases in the assembly that are error-free as well as a score for each base in the assembly, which can be used to calculate the mean base score. We found that the percent of error-free bases was correlated with the mean base score (Spearman = 0.62), so chose to use only the former when calculating mean rankings.

BUSCO reports the percentage of complete single-copy orthologs (BUSCOs) found in each assembly given a set of BUSCOs expected to be present in the genome. The higher the percentage of complete BUSCOs found, the more complete and correctly put together an assembly is likely to be. For short read assembly evaluations, we used BUSCO v1.22 and the 2,675 SCOs from arthropods.

**4.1.3.1 Size statistics:**
Size statics (such as NG50) of the resulting contigs/scaffolds were calculated with asm-stats.py from SciaraTools: https://github.com/JohnUrban/sciara-project-tools

**4.1.3.2 Mapping the reads back to each assembly:**
With: Bowtie 2 version 2.2.9 (Langmead and Salzberg 2012), samtools v1.3 (Li et al. 2009).

The raw reads (Short Read Set #1) were mapped back to each assembly, and subsequently used to identify the percent that mapped and in ALE and $FRC^{bam}$. Note: LAP and REAPR have their own mapping strategies.

```
FILE=ASM.name.fasta
BASE=ASM.name
```

```
bowtie2-build $FILE $BASE

bowtie2 -p $P -q --very-sensitive -N 1 -x $BASE -1 $R1 -2 $R2 2>
${BASE}.err | samtools sort -o reads.bam

samtools index reads.bam
```

###### 4.1.3.3 Percent of Illumina reads that mapped:
This was taken from the output report of Bowtie2.

###### 4.1.3.4 ALE
With: boost 1.55, ALE (version from Scott Clark 2010)

ALE was used with the raw reads (Short Read Set #1) – see "Mapping reads back to each assembly" above.

```
FILE=ASM.name.fasta
BASE=ASM.name

ALE reads.bam $FILE ${BASE}.ALE.txt >> ${BASE}.err
```

Note: ABySS outputs contigs with non-ACGT IUPAC letters, which was breaking ALE. Therefore, each non-ACGT IUPAC letter was converted to one of the ACGT bases it represented at random with respect to the frequency ACGT occurs in the assembly. This was done using IUPAC-to-ACGT.py from: https://github.com/JohnUrban/sciara-project-tools

```
IUPAC-to-ACGT.py --f abyss.asm.fasta > abyss.asm-acgt.fasta
```

###### 4.1.3.5 FRC$^{bam}$
FRC$^{bam}$ was used with the raw reads (Short Read Set #1) – see "Mapping reads back to each assembly" above.

```
FILE=ASM.name.fasta
BASE=ASM.name

FRC --pe-sam reads.bam --pe-max-insert 800 --genome-size 292000000 --
output ${BASE}.frc
```

For short read assemblies, normalizing the number of features reported by FRC by the assembly size did not have much of an effect on rankings. Note that an expected genome size of 292Mb was used here rather than 280 Mb based on an earlier estimate.

###### 4.1.3.6 LAP
We used the LAP scores generated for each assembly given the same set of input reads to rank them relative to each other. The raw read set (Short Read Set #1) was down sampled with sampleFastq.py (from: https://github.com/JohnUrban/sciara-project-tools). We tried lap using different proportions (p) of the full dataset, where p was specified during down-sampling as either 0.0001, 0.001, and 0.01 for approximately 15k, 150k, and 1.5 M sampled paired reads respectively. Note: sampleFastq.py returns "downsampled.1.fastq" and "downsampled.2.fastq". Ultimately, we used the results for 1.5 M reads.

```
R1=R1.fastq
```

```
R2=R2.fastq
FILE=ASM.name.fasta
BASE=ASM.name
P=16
PROPORTION=p

sampleFastq.py -1 R1.fastq -2 R2.fastq -wo -p $PROPORTION -o
downsampled

bowtie2-build $FILE $BASE

calc_prob.py -p $P -a $FILE  -q -1 $R1 -2 $R2 -X 800 -I 0 -o fr -m 432
-t 75 -b $BASE > ${BASE}.prob

sum_prob.py -i ${BASE}.prob > ${BASE}.lapscore
```

###### 4.1.3.7 REAPR

REAPR was used with the raw reads (Short Read Set #1). For short read assemblies, we looked at the reported percent error-free bases. We also calculated the mean of the scores over every base in the genome, which gave similar ranking results.

```
FILE=ASM.name.fasta
BASE=ASM.name

reapr facheck ${FILE} ${BASE}_renamed
reapr perfectmap ${BASE}_renamed.fa $R1 $R2 432 perfect
reapr smaltmap ${BASE}_renamed.fa $R1 $R2 mapped.bam  -n $P
reapr pipeline ${BASE}_renamed.fa mapped.bam output_directory perfect
```

###### 4.1.3.8 BUSCO v1:

Short read assemblies were evaluated based on the number of Arthropoda BUSCOs detected in each.

```
LINEAGE=/Path/to/lineages/arthropoda
MODE=genome
BUSCO_v1.22.py -in $ASM -o $OUT -l $LINEAGE -m $MODE --cpu $CPU
```

#### 4.1.4 Contamination analyses for the selected Platanus short read assembly

It has been shown that filtering out the contaminating reads can improve an assembly (Kumar et al. 2013). Therefore, to remove reads from contaminating species, we adopted a procedure similar to that used for the Tardigrade genome (Koutsovoulos et al. 2016) with the help of BlobTools (Laetsch and Blaxter 2017). We focused on the Platanus assembly produced from quality filtered, error-corrected reads. Instead of using the final gap-closed scaffolds, we used the Platanus contigs (largest contig ~77 kb) to allow each contig to be characterized separately and avoid discarding data due to misjoins.

Ideally one could use coverage and/or GC content information associated with annotated bacterial contigs to also eliminate the likely bacterial contigs that were not annotated. Unfortunately, the annotated bacterial contigs were of equivalent coverage compared with those annotated as arthropod (see Supplemental Figure S3). Moreover, the GC content of the bacterial and eukaryotic contig clusters overlapped enough to limit its usefulness. Therefore, we used an alternative set of reads from pre-amplification stage salivary glands (Urban et al. 2016), reasoning that since this dataset was from a different tissue, from a different stage, and prepared by a different person, the contaminating contigs would have much lower coverage. Over 95% of these reads mapped to the assembly, suggesting most or all of the somatic *Sciara* genome is represented in the Platanus contigs. As expected, the BlobPlot demonstrates that the coverage of the bacterial cluster is much lower in this dataset (see Supplemental Figure S3). We therefore marked for removal non-Arthropod-labeled contigs (and associated reads) from the assembly that were either labeled as super-kingdom "Bacteria" or had coverage < 0.1. To filter out contaminating reads for re-assembly, we removed all paired-end reads where both mates mapped to contigs marked for removal, and retained pairs when at least one mate mapped inside a retained contig or when both mates did not map to the assembly. The retained paired-end reads were used for assembly with Platanus in four ways: (i) no quality filtering nor error-correction, (ii) no quality filtering but error-correction with BayesHammer, (iii) quality filtering but no error-correction, and (iv) both quality filtering and error correction.

Overall, the contiguity statistics of the re-assembled Platanus contigs did not improve over simply eliminating the contaminating contigs from the original assembly, with the longest contig being ~28 kb in both scenarios. The scaffold contiguity statistics were slightly smaller than the original Platanus assemblies due to the absence of the very large bacterial scaffolds. Evaluation scores were similar among the four Platanus re-assemblies from different sets of filtered reads. REAPR's percent error-free bases metric improved drastically from ~56% error-free bases in the original assemblies to ~80% error-free bases in the assemblies produced from contamination-filtered reads. The assembly from quality-filtered, error-corrected reads had the highest percentage of error-free bases. We attempted a second round of contamination-filtering on this assembly, but found few additional benefits. There were only 15 contigs labeled as Bacteria, all *Rickettsia* at the genus level, summing in length to only 2,277 bp.

##### 4.1.4.1 Obtaining taxonomy IDs from BLAST hits for each short read Platanus contig:

The Platanus fasta file containing contigs (not scaffolds) from the assembly of quality filtered, error-corrected reads (Short Read Set #4) was subdivided into 1000 different fasta files and submitted to SLURM as a batch script array that ran the following on each:

```
NT=/Path/to/entire/nucleotide/database
blastn -task megablast -query $Q -db $NT \
 -outfmt '6 qseqid staxids bitscore std sscinames sskingdoms stitle' \
 -culling_limit 5 \
 -num_threads $P \
 -evalue 1e-25 \
 -out ${BLASTDIR}/${PRE}.${SLURM_ARRAY_TASK_ID.blastout}
```

After ensuring all jobs completed successfully (and re-running if not), final blast results were all combined into a single file:

```
platanus.q5.bh.blast.results
```

The NCBI Taxonomy database containing nodes.dmp and names.dmp was downloaded into a subdirectory referred to as "./tax/" in commands below. This was used with BlobTools and custom taxonomy scripts as described below to classify contigs given the taxonomy IDs associated with each.

**4.1.4.2 Using BlobTools and custom scripts to label each contig with a taxonomy:**
Blob plots using platanus kmer coverage from the assemblies:

```
B=platanus.q5.bh.blast.results
BASE=platanus.q5.bh.contig
REF=${BASE}.fasta
DB=${BASE}.BlobDB.json

# Create blob database using kmer coverage from the assemblies
blobtools create -i ${REF} -y platanus --nodes tax/nodes.dmp --names
tax/names.dmp -o ${BASE} -t $B

# Plot
blobtools blobplot -i ${DB} [--format pdf]
blobtools blobplot -i ${DB} -r superkingdom [--format pdf]
blobtools blobplot -i ${DB} -r order -p 10 [--format pdf]
blobtools blobplot -i ${DB} -r family -p 12 [--format pdf]

# Make Table with Taxon Summaries for each contig
blobtools view -o  ${BASE}.table -i $DB -r all
```

Blob plots using read coverage:
The GC content for bacterial contigs was similar to the eukaryotic contigs. Moreover, the Bacterial and Eukaryotic contigs had similar coverage levels given the male embryo dataset used in the assemblies. Therefore, we tried a separate dataset generated from dissected out pre-amplification stage larval salivary glands (Urban et al. 2016). Reads were aligned with Bowtie 2 (Langmead and Salzberg 2012).

```
PFX=pre-amp-sal-gland-reads
READS=${PFX}.fastq.gz
BASE=platanus.q5.bh.contig
REF=${BASE}.fasta
B=platanus.q5.bh.blast.results
DB=${PFX}.${BASE}.BlobDB.json

# Map reads
bowtie2 -p $P -q --very-sensitive -N 1 -x ${BASE} -U ${READS} 2>
${PFX}.${BASE}.bt2.err | samtools sort --threads $P -T ${PFX}.tmp -o
${PFX}.bam

# Get contig coverage levels
blobtools bam2cov -i ${REF} -b ${PFX}.bam -o bam2cov --mq 0
grep -v ^# bam2cov.${PFX}.bam.cov | cut -f 1,3 > ${PFX}.cov

# Create Blob database
blobtools create -i ${REF} --cov ${PFX}.cov --nodes tax/nodes.dmp --
names tax/names.dmp -o ${PFX}.${BASE} -t $B

# Plot
blobtools blobplot -i ${DB} [--format pdf]
```

```
blobtools blobplot -i ${DB} -r superkingdom [--format pdf]
blobtools blobplot -i ${DB} -r order -p 10 [--format pdf]
blobtools blobplot -i ${DB} -r family -p 12 [--format pdf]
```

We also used custom scripts (https://github.com/JohnUrban/sciara-project-tools/tree/master/taxon) that segregate sequence names based on the closest taxonomy level (species, genus, family, order, class, phylum, kingdom, superkingdom, othersuperkingdoms) to our target species (Bradysia coprophila, Bradysia, Sciaridae, Diptera, Insecta, Arthropoda, Metazoa, Eukaryota, Bacteria/Archaea) that is found in the BLAST hits for each sequence. We required only 1 BLAST hit for a given level for it to be considered the closest level (i.e. if 1 BLAST hit says "Bradysia coprophila", then it will be assigned to the species level as the closest level even if other levels have more hits).

```
NODES=tax/nodes.dmp
NAMES=tax/names.dmp
BFILE=platanus.blobfilt1.q5.bh.contig.blast.results
TAX=$BFILE.taxonomy.out
PRE=platanus.blobfilt1.q5.bh.contig.taxsum
ALL=q5-bh/platanus.blobfilt1.q5.bh_contig.names
taxonomyFromTaxID.py -i $BFILE -nc 1 -tc 2 -no $NODES -na $NAMES > $TAX
taxonomy-summarizer.py -i $TAX -o $PRE -a $ALL
```

This reproduced the BlobTools approach, returning the same contigs labeled as Bacterial.

**4.1.4.3 Obtaining contamination-filtered reads for re-assembly**:
The set of male embryo paired-end reads (Short Read Set #1) was filtered as follows, making use of BEDtools (Quinlan and Hall 2010), SAMtools (Li et al. 2009), and linux commands.

```
# Make Table with Taxon Summaries for each contig
blobtools view -o  ${PFX}.${BASE}.table -i $DB -r all

## Grab contigs that are not labeled Bacterial at Super Kingdom level
that are above a coverage level of 0.1
grep -v ^#  ${PFX}.${BASE}.table | \
      awk '$6 != "Bacteria" && $5 >= 0.1' | \
      awk 'OFS="\t" {print $1,0,$2}' | \
      sortBed -i - > contigs.to.keep.bed

## Add to those, contigs below the coverage cutoff of 0.1 that are
explicitly labeled as Arthropoda sequences
grep -v ^#  ${PFX}.${BASE}.table | \
      awk '$5 < 0.1' | \
      grep Arthropoda | \
      awk 'OFS="\t" {print $1,0,$2}' | \
      sortBed -i - >> contigs.to.keep.bed

# Get read pairs from male embryo dataset (SRS#1 above) where at least
one read in the pair maps to one of the contigs-to-keep
BAM=${PFX}.bam
BED=contigs.to.keep.bed
bedtools pairtobed -abam ${BAM} -b ${BED} | \
      samtools sort -n | \
      samtools fastq -1 ${PFX}.m.R1.fq -2 ${PFX}.m.R2.fq -n -

#Also identify reads where neither maps to the assembly that might be
rescued in a subsequent assembly
samtools view -bS -f12  ${BAM} | \
```

```
    samtools sort -n | \
    samtools fastq -1 ${PFX}.u_u.R1.fastq -2 ${PFX}.u_u.R2.fastq -n -

# Combine ${PFX}.m.R1.fq and ${PFX}.u_u.R1.fastq
# Combine ${PFX}.m.R2.fq and ${PFX}.u_u.R2.fastq
```

**4.1.4.4 Platanus contamination-filtered re-assemblies**:

Four new Platanus assemblies were made using the set of contamination-filtered reads as well as quality-filtered and/or error-corrected derivatives of it (Short Read Sets #5 - #8).

(i) Contamination-filtered reads (Short Read Set #5) were assembled as described in Platanus sub-section of Section 2.1.2.

(ii) Contamination-filtered reads that were quality-filtered using the Trimmomatic command below (Short Read Set #6) were subsequently assembled as described in Platanus sub-section of Section 2.1.2.

```
java -jar $TRIMMOMATIC_BASE/Trimmomatic-0.32/trimmomatic-0.32.jar PE -
trimlog trim.log $IN $OUT ILLUMINACLIP:${PRIMERS}:2:30:10 LEADING:5
TRAILING:5 SLIDINGWINDOW:4:5 MINLEN:100
```

(ii) Contamination-filtered reads that were then error-corrected with BayesHammer using the SPAdes command below (Short Read Set #7) were subsequently assembled as described in Platanus sub-section of Section 2.1.2.

```
spades.py --pe1-1 $PE1 --pe1-2 $PE2 -o assembly -t 16 --only-error-
correction
```

(iv) Contamination-filtered reads that were already quality-filtered (SRS#6) were then subsequently error-corrected with the following SPAdes command to produce Short Read Set #8, which was subsequently assembled as described in Platanus sub-section of Section 2.1.2.

```
spades.py --pe1-1 $PE1 --pe1-2 $PE2 --pe1-s $SE -o assembly -t 16 --
only-error-correction
```

**4.1.4.5 Evaluations of contamination-filtered re-assemblies:**

The four Platanus assemblies (gap-closed scaffolds) from contamination-filtered datasets were evaluated in the same way as the evaluations of the first 40 assemblies were as described in **Section 4.1.3** (Size statistics, Percent reads mapped, ALE, FRC$^{bam}$, ALE, REAPR, BUSCOv1) except that the set of contamination filtered reads (Short Read Set #5) was used for these evaluations. This means some of the scores based on reads are not directly comparable to the first 40 assemblies (and do not appear in the ranking matrices, etc). Nevertheless, for our datasets, this re-assembly approach yielded assemblies that were extremely similar the original set of Platanus assemblies after simply removing bacterial labeled contigs/scaffolds. It did not improve upon the size statistics.

Assembly scores were similar among the four assemblies in **Section 4.1.4.4** (“i”, “ii”, “iii”,”iv”). Assembly “ii” (quality filtered) received the best LAP and ALE scores. Assembly “i” had the fewest number of features detected by $FRC^{bam}$. Assembly “iv” had the highest percentage of error-free bases as determined by REAPR. Since “iv” had the highest REAPR score and since the combination of quality-filtering and error-correction also won out in the previous set of Platanus evaluations before contamination filtering, we chose this assembly as the final/only Platanus assembly to take into a second round of contamination filtering. The contigs and scaffolds were separately BLAST for taxonomy information as above. BlobTools databases and plots were constructed as above for kmer coverage as well as coverage from pre-amplification salivary gland reads. However, there was no additional contamination filtering needed.

#### 4.2 Long Read Assemblies using PacBio and Oxford Nanopore Data

**Summary:**
We generated five hybrid assemblies (that combined short and long read technologies) starting with our final Illumina-based Platanus contigs using DBG2OLC (Ye et al. 2016) as well as 45 long-read-only assemblies with several long read assemblers including Canu (Koren et al. 2017), Falcon (Chin et al. 2016), Miniasm (Li 2016), ABruijn (Lin et al. 2016), and SMARTdenovo (https://github.com/ruanjue/smartdenovo). Assembling the *Sciara* genome with HINGE (Kamath et al. 2016) was also attempted, but it required more computational resources than we had available. For each assembler we tried different inputs (see supplemental figure S4 as well): (i) quality-filtered PacBio subreads , (ii) all PacBio subreads, (ii) quality-filtered PacBio subreads with all ONT 2D reads, (iii) all PacBio subreads with one ONT read per molecule, and (iv) only for Miniasm, all PacBio subreads and all ONT reads for each molecule ("PBall+ONTall") where each molecule can have up to three reads (template, complement, 2D). In total, there were 50 assemblies chosen to polish and evaluate after selecting several of the best candidates from each assembler for each dataset using size statistics.

Canu and Falcon follow similar paradigms where long reads are first error-corrected by mapping them against each other to construct higher quality consensus reads. They both then use the error-corrected reads to find overlaps and ultimately generate contigs. Miniasm, ABruijn, and SMARTdenovo skip an initial error-correction step and find overlaps in the raw long reads instead. ABruijn, Canu, Falcon, and SMARTdenovo have consensus steps built into the end of their assembly pipelines. RaCon, a rapid consensus caller (Vaser et al. 2016), was used with Miniasm assemblies to generate a consensus and the consensus approach using pbdagcon (https://github.com/PacificBiosciences/pbdagcon) (Chin et al. 2013) was used for Platanus+DBG2OLC assemblies as recently described (Chakraborty et al. 2016). All evaluation metrics for Miniasm, which skips both pre-assembly error-correction and post-assembly consensus steps, improved with RaCon as expected. For example, BUSCO found 0 BUSCOs in Miniasm assemblies before RaCon and 29-32% afterward. Similarly, only 29-31% of Illumina reads mapped and 0% of bases were judged error-free by REAPR in Miniasm assemblies prior to RaCon whereas 94-95% of Illumina reads mapped and 70-71% of bases were judged error-free after RaCon.

The number of assemblies from each assembler was mostly determined by how easy, fast, or resource-efficient a given assember was to run different combinations of parameters or reads with. The number for each was:

Abruijn = 3
Canu = 17
DBG2OLC (hybrid) = 5
Falcon = 12
Miniasm = 8
SMARTdenovo = 5

Total = 50

##### 4.2.1 Long read inputs to assemblers

We tried different levels of quality filtering of the long reads. For example, we either used all or a filtered subset of the PacBio reads, and we either blended in a read from every molecule sequenced on the MinION, which included 1D reads when 2D reads were not present, or blended in only 2D reads. Details are below.

###### 4.2.1.1 Long read set naming

Long Read Set #1 (PBall)
= All PacBio subreads

Long Read Set #2 (PBfilt)
= PacBio reads filtered to be have minReadScore > 0.75

Long Read Set #3 (PBall+ONTmol)
= All PacBio subreads (LRS#1)
+ Oxford Nanopore "molecule" reads (one read per molecule)

Long Read Set #4 (PBall+ONT2d)
= All PacBio subreads (LRS#1)
+ Oxford Nanopore 2D reads

Long Read Set #5 (PBfilt+ONT2d)
= PacBio reads filtered to be > 0.75 (LRS#2)
+ Oxford Nanopore 2D reads

Long Read Set #6 (PBall+ONTall)
= All PacBio subreads (LRS#1)
+ All Oxford Nanopore reads (one, two, or three per molecule)
Note: Only used by Miniasm and RaCon

Long Read Set #7 (PBfilt+ONTall)
= PacBio reads filtered to be > 0.75 (LRS#2)
+ All Oxford Nanopore reads (one, two, or three per molecule)
Note: Only used by Miniasm

Long Read Set #8 (PBfilt+ONTmol):
= PacBio reads filtered to be > 0.75 (LRS#2)
+ Oxford Nanopore "molecule" reads (one read per molecule)
Note: Only used by Miniasm

##### 4.2.1.2 Obtaining fasta/fastq files from PacBio .bax.h5 files using bash5tools

Bash5tools.py from pbh5tools (https://github.com/PacificBiosciences/pbh5tools) was used. Either all PacBio sub-reads were used, or quality filtered subreads with a cutoff of 0.75.

```
# Long Read Set #1 (PacBioAll)
bash5tools.py --readType subreads \
      --outType fasta \
      --outFilePrefix $PREFIX file.bax.h5

# Long Read Set #2 (PacBioFilt)
bash5tools.py --readType subreads --outType fasta \
      --minReadScore 0.75 \
      --outFilePrefix $PREFIX file.bax.h5
```

##### 4.2.1.3 Obtaining fasta/fastq files from MinION fast5 files using fast5tools

In summary, Fast5Tools (https://github.com/JohnUrban/fast5tools) was used to extract and analyze reads from the many different MinION libraries we generated. Examples of how one molecule per read or only 2D reads were extracted are show here and described more below:

```
# Extract all reads (up to three per molecule) from fast5s
# This gives ONTall as used with Long Read Set #6

fast5tofastx.py -r all -o fasta --tarlite minIONreads.tar.gz >
ONTall.fa

# Get one read per molecule(Urban et al 2015, and described below)
# Can derive from the fast5 files as above or from ONTall.fa
# This gives ONTmol as used in Long Read Set #3

fast5tofastx.py -r molecule -o fasta --tarlite minIONreads.tar.gz >
ONTmol.fa
# OR:
filterFast5DerivedFastx.py -r molecule ONTall.fa > ONTmol.fa

# Extract only the 2D reads from that previous dataset (or from fast5s)
# This produces ONT2d as used in Long Read Sets #4 and #5

fast5tofastx.py -r 2d -o fasta --tarlite minIONreads.tar.gz > ONTmol.fa
# OR:
filterFast5DerivedFastx.py -r 2d ONTmol.fa > ONT2d.fa
```

##### 4.2.1.4 A more detailed description of Fast5Tools and MinION read analyses:

Fast5 files were base-called with Oxford Nanopore's Metrichor cloud base-caller at the time the data was produced. See information on libraries elsewhere for dates and MAP kit versions. Metrichor (ONT base-caller) returned updated fast5 files into two folders: "pass" and "fail". "Pass" contained only fast5 files where 2D base-calling was successful and the mean quality of the 2D read was >9. Everything else (including other fast5s containing 2D reads with Q<9, fast5s with only 1D base-calling, and fast5s that failed base-calling) was put into the "fail" folder by Metrichor. We made use of data from both folders where possible. For libraries 01, 02, 03, 04, 05, 06, 07,

08, and 09, our toolset called poreminion (https://github.com/JohnUrban/poreminion) as used previously (Urban et al. 2015) to categorize the fast5s in the "fail" folder that were not base-called due to various reasons: no template, too few events, too many events, or contained an error where blocks of events are artificially repeated. The latter error was present for a minority of fast5 files in the early days of the MinION Access Program and was fixed by ONT before sequencing of Library09. Library09 and all subsequent libraries did not have any of these errors as expected.

```
#To categorize and rescue reads from the fail folder
poreminion uncalled -m -o fail-filter fail/

# To remove reads from early libraries that contained artificial
repeats of events
poreminion timetest -m -o fail-filter fail/
```

For all libraries, the pass and fail folders were individually archived and compressed with:

```
tar -xzf pass.tar.gz pass/
tar -xzf fail.tar.gz fail/
```

As this data was collected over the course of many months (2014-2016) and ONT was rapidly developing and changing the innards of the HDF5 fast5 files, we were presented with the need for a tool that can handle any version of a fast5 file given to it. At this point, we created another toolset called Fast5Tools (https://github.com/JohnUrban/fast5tools) that is able to deal with all of the different fast5 versions used in this study. Fast5Tools maintains all or most of the functionality of poreminion, although pre-filtering is no longer necessary.

For all MinION libraries, Fast5Tools was used to extract fasta and fastq information from the tar archived pass.tar.gz and fail.tar.gz files. Using the "--tarlite" option, Fast5Tools is able to work on files directly from the gzipped tar archive without extracting it all at once. This prevents issues of disk space that may arise from fully extracting the contents of the giant gzipped tar files before performing the needed operations. First to extract either all fasta or all fastq reads from every fast5 file, fast5tofastx.py was used:

```
fast5tofastx.py -r all -o fasta --tarlite pass.tar.gz > pass.all.fa

fast5tofastx.py -r all -o fasta --tarlite fail.tar.gz > fail.all.fa
```

The fast5 files can be filtered for length and quality information in the previous step (`fast5tofastx.py`). However, it can be done in subsequent steps as well as the headers of the resulting fasta/fastq returned from "`fast5tofastx.py`" contain all the information necessary to filter for reads using length, mean quality score, and other information using "`filterFast5DerivedFastx.py`". Therefore, when the "`-r all`" option is used as above, filtering can be done on the fasta/fastq files exactly as from fast5 files. When one does not need access to the signal level of the data, this can be used to keep the smaller fasta/fastq file while storing the larger tarchives in longer term storage if necessary. The basic header structure for each fasta entry looks like this:

```
>readtype|len:LEN|Q:Q|channel:CHANNEL|Read:READNUMBER|asic:ASIC_ID|run:RUN_ID
|device:DEVICE_ID|model:BASECALL_MODEL
```

An example of the 3 read types (template, complement, and 2d) from a passing molecule:

```
>template|len:2781|Q:4.45259884791|channel:508|Read:80|asic:3372918206|
run:609a1b4408f2200edefabc0d97b202bcaccf94db|device:MN15216|model:r7.3_
e6_70bps_6mer
```

```
>complement|len:2783|Q:4.43810484237|channel:508|Read:80|asic:337291820
6|run:609a1b4408f2200edefabc0d97b202bcaccf94db|device:MN15216|model:r7.
3_e6_70bps_6mer

>2d|len:3248|Q:10.8914446375|channel:508|Read:80|asic:3372918206|run:60
9a1b4408f2200edefabc0d97b202bcaccf94db|device:MN15216|model:r7.3_e6_70b
ps_6mer
```

The fasta/fastq derived from the fast5 files with Fast5Tools can then be filtered using filterFast5DerivedFastx.py. We obtained all 2D reads from the fasta/fastq file by:

```
filterFast5DerivedFastx.py -r 2d libX.pass.all.fa > libX.pass.2d.fa

filterFast5DerivedFastx.py -r 2d libX.fail.all.fa > libX.fail.2d.fa
```

High quality pass 2D reads can be obtained by filtering just the pass.all.fa file for each library (using the above command) or if all reads are combined in a single fasta, doing:

```
filterFast5DerivedFastx.py -r 2d libX.all.fa --minq 9 > libX.pass.2d.fa
```

We selected one read per molecule using the "molecule" definition that we used previously (Urban et al. 2015). This takes the 2D read if one is present, the longer of template or complement if 2D is not present (but complement is), or the template read if it is the only one present.

```
filterFast5DerivedFastx.py -r molecule libX.pass.all.fa >
libX.pass.molecule.fa

filterFast5DerivedFastx.py -r molecule libX.fail.all.fa >
libX.fail.molecule.fa
```

To obtain statistics from each fast5 file on the reads present, Fast5Tools can work directly with the fast5 files (or tarchive of fast5 files) as well as from the fast5-derived fasta files (particularly when all reads are dumped out from each file as above). Both approaches give the same results concerning length and mean quality scores (more information about events can be obtained directly from the fast5 file):

```
fast5stats.py --standard --errfile libX.errfiles.txt --tarlite
libX.pass.tar.gz libX.fail.tar.gz > stats.txt

fast5DerivedFastxMoleculeStats.py libX.pass.all.fa libX.fail.all.fa >
stats.txt
```

Note that “--errfile errfiles.txt” is populated with information on what files had errors or were not base-called for various reasons (as stated above). There is no option for this when grabbing the statistics from fasta files as any erroneous or un-basecalled fast5 is ignored during extraction with “`fast5tofastx.py`”. The output of both commands (`fast5stats.py`, `fast5DerivedFastxMoleculeStats.py`) is just one line per molecule (i.e. one line per original fast5 file) even if the information is coming from a fasta multiple reads per molecule. This is possible because `fast5DerivedFastxMoleculeStats.py` unites different reads from the same molecule by using their shared molecule names derived from the channel, the read number, the asic ID, the run ID, and device ID). The output of both commands also share the first 11 columns:

1 = molecule name
2 = molecule length
3 = molecule mean q score
4 = has complement
5 = has 2d
6 = 2d seq len
7 = template seq len
8 = complement seq len
9 = 2d mean q score
10 = template mean q score
11 = complement mean q score

The first 11 columns of the output table was summarized with fast5standardSummary.py:

```
fast5standardSummary.py -V -f libX.stats.txt
```

At the moment, this outputs 264 pieces of information, including molecule length statistics (minimum, maximum, mean, median, N25, N50, N75, expected length) on molecules as well as the various read types. One can simply combine all stats.txt tables from each library to compute the summary on all molecules from all libraries.

###### 4.2.1.5 MarginStats for percent identity of MinION Reads

The BAM file containing the alignments of all ONT reads mapped to the highest ranked PacBio-only assembly was filtered to remove unmapped reads using SAMtools. Then the resulting BAM files was partitioned into 400 smaller files using splitSAM.py from https://github.com/JohnUrban/sciara-project-tools. This enabled us to compute percent identities in parallel in the next step. MarginStats (Jain et al. 2015) was then used to calculate the end-to-end percent identities of the reads in each file. Optimized re-alignment was not performed, which can increase the percent identity reported. For this reason and since the assembly may have some errors itself, the percent identities reported here are conservative estimates. Since reads were extracted using our set of tools for working with fast5 files from ONT called fast5tools (https://github.com/JohnUrban/fast5tools), read names contained mean quality scores, which could then be paired with the percent identities from MarginStats.:

```
samtools view -bh -F4
canu10.minrl500.dip3x.pball.quiverfinal1.pilon2x/ont2d.bam >
reads/ont.bam
splitSAM.py --bam reads/ont.bam --nfiles 100 --nreads 1382636
```

On each BAM:

```
marginStats --noStats --printValuePerReadAlignment --identity $BAM $fq
$ref > wd/${pre}.txt
```

On each MarginStats output text file:

```
awk `{sub(/\ /,"\n"); gsub(/\t/,"\n"); print\' wd/${PRE}.txt | grep -v
\^{ValuesIdentity | paste - <(grep -v \^{@ <(samtools view
splitfiles/${PRE.bam)) | awk `OFS="\t" {print $2,$1\' >
pairs/${PRE}.txt
```

#### 4.2.2 Hybrid-assembling long reads with short reads

##### 4.2.2.1 DBG2OLC with Platanus short read contigs and PBDAGCON

We used the contigs from our "final" platanus assembly, which were assembled with a set of reads that went through one round of contamination-filtering, followed by quality filtering with Trimmomatic, and error-correction with BayesHammer from SPAdes.

Five hybrid assemblies were generated using Long Read Sets #1-5 (in place of longreads.fasta in commands below).

```
./DBG2OLC k 17 KmerCovTh 2 MinOverlap 20 AdaptiveTh 0.002 LD1 0 MinLen
200 Contigs platanus.contigs.fa RemoveChimera 1 f longreads.fasta

cat platanus.contigs.fa longreads.fasta > ctg_pb.fasta

mkdir consensus_dir

split_and_run_pbdagcon.path.sh backbone_raw.fasta
DBG2OLC_Consensus_info.txt ctg_pb.fasta consensus_dir >
consensus_log.txt
```

#### 4.2.3 Assembling the long reads alone

##### 4.2.3.1 ABRUIJN
Three ABruijn assemblies were generated using Long Read Sets #1, #3, and #5.

```
abruijn.py longreads.fasta Sciara $COV -t 32 --iterations 1
```

PBall (Long Read Set #1) : COV=42
PBall+ONTmol (Long Read Set #3) : COV=54
PBfilt+Ont2d (Long Read Set #5) : COV=47

At the time of this work, ABruijn runs required allocating 512 GB RAM and 32 CPUs, and 7-8 days of running time. Thus, it was impractical to try exhaustive combinations of reads, filtering, etc.

##### 4.2.3.2 Canu
Canu is very sophisticated in automatically requesting appropriate resources and distributing/parallelizing its jobs across as many CPUS/nodes available on our compute cluster. It therefore finishes within a day or 2, and it was possible to test many different combinations of reads, filtering, parameters, etc. Indeed, we have also tested many earlier versions of Canu (not reported here). We report on 18 Canu assemblies in this study, generated using Long Read Sets #1-5. Note that an expected genome size of 292Mb was used here rather than 280 Mb based on an earlier estimate. That will have no effect on assemblies that used all of the data (when setting an extremely high coverage), and little effect on the amount of data used otherwise.

Below, variables that show up in the Canu commands are defined. Not all variables are used in all commands.

```
PBall=all_subreads.fasta  # LRS#1/PBall
PBfilt=subreads.gt0.75.fasta  # LRS#2/PBfilt
ONTmol=ontmolecule.fasta #Combine w/ PBall for LRS#3
ONT2d=ont2d.fasta #Combine w/ PBall for LRS#4 or w/ PBfilt for LRS#5
```

```
PBONT2D=allPBsubreads_ont2d.fasta # LRS#4/PBallONT2d
PBFILTONT2D=pbsubreads.gt0.75_ont2d.fasta # LRS#5/PBfiltONT2d
PBMOL=allPBsubreads_ontmolecule.fasta # LRS#3/PBallONTmol
COROUTCOV=500 # excessively high to include all reads
T=24:00:00
G=292m
```

Canu_1 = default.pball = Canu 1.3 using only PacBio reads (LRS#1/PBall):

```
NAME=g292-default-all
canu  -p $NAME -d $NAME  genomeSize=$G  \
      -pacbio-raw $PBall  "gridOptions=--time $T"  oeaMemory=8
```

Canu_2 = minrl500.pball = Canu 1.3 using only PacBio reads (LRS#1/PBall) and specifying a minimum read length of 500 bp:

```
NAME=g292-default-all-min500
canu -p $NAME -d $NAME genomeSize=$G \
      -pacbio-raw $PBall "gridOptions=--time $T" \
      minReadLength=500  oeaMemory=8
```

Canu_3 = corcov500.pball = Canu 1.3 using only PacBio reads (LRS#1/PBall) and fixing coverage parameters to use all reads (not just a subset of the longest sampled by Canu):

```
NAME=g292-default-all-corcov500
canu -p $NAME -d $NAME genomeSize=$G -pacbio-raw $PBall \
      "gridOptions=--time $T" oeaMemory=8 \
      corOutCoverage=$COROUTCOV
```

Canu_4 = corcov500minrl500.pball = Canu 1.3 using only PacBio reads (LRS#1/PBall), specifying a minimum read length of 500 bp, and fixing coverage parameters to use all reads (not just a subset of the longest sampled by Canu):

```
NAME=g292-default-all-corcov500-min500
canu -p $NAME -d $NAME genomeSize=$G \
      -pacbio-raw $PBall "gridOptions=--time $T"  \
      minReadLength=500 oeaMemory=8 corOutCoverage=$COROUTCOV
```

Canu_5 = corcov500.pbfilt = Canu 1.3 using only filtered PacBio reads (LRS#2/PBfilt) and fixing coverage parameters to use all reads (not just a subset of the longest sampled by Canu):

```
NAME=g292-default-q0.75-corcov500
canu -p $NAME -d $NAME genomeSize=$G
      -pacbio-raw $PBfilt "gridOptions=--time $T" \
      oeaMemory=8 corOutCoverage=$COROUTCOV
```

Canu_6 = corcov500minrl500.pbfilt = Canu 1.3 using only filtered PacBio reads (LRS#2/PBfilt), specifying a minimum read length of 500 bp, and fixing coverage parameters to use all reads (not just a subset of the longest sampled by Canu):

```
NAME=g292-default-q0.75-corcov500-min500
canu -p $NAME -d $NAME genomeSize=$G \
      -pacbio-raw $PBfilt "gridOptions=--time $T"  \
      minReadLength=500 oeaMemory=8 corOutCoverage=$COROUTCOV
```

Canu_7 = corcov500minrl500.pball.ontmol = Canu 1.3 using all PacBio reads and one read per molecule from ONT (LRS#3/PBall+ONTmol), specifying a minimum read length of 500 bp, and fixing coverage parameters to use all reads (not just a subset of the longest sampled by Canu):

```
NAME=g292-default-pball-ontmol-corcov500-min500
canu -p $NAME -d $NAME genomeSize=$G -pacbio-raw $PB \
      -nanopore-raw $ONTmol  "gridOptions=--time $T" \
      minReadLength=500 oeaMemory=8 corOutCoverage=$COROUTCOV \
      corMemory=40
```

Canu_8 = corcov500minrl500.aspb-e025.pball.ontmol = Canu 1.3 using all PacBio reads and one read per molecule from ONT (LRS#3/PBall+ONTmol), specifying a minimum read length of 500 bp, fixing coverage parameters to use all reads (not just a subset of the longest sampled by Canu), and combining the PB and ONT data into a single file treated by Canu as PacBio data:

```
NAME=g292-default-pball-ontmol-corcov500-min500-aspb
canu -p $NAME -d $NAME genomeSize=$G errorRate=0.02 \
      -pacbio-raw $PBall_ONTmol "gridOptions=--time $T" \
      minReadLength=500 oeaMemory=8 \
      corOutCoverage=$COROUTCOV corMemory=40
```

Canu_9 = corcov500minrl500.pball.ont2d = Canu 1.3 using all PacBio reads and 2D nanopore reads (LRS#4/PBall+ONT2d), specifying a minimum read length of 500 bp, and fixing coverage parameters to use all reads (not just a subset of the longest sampled by Canu):

```
NAME=g292-default-pball-ont2d-corcov500-min500
canu -p $NAME -d $NAME genomeSize=$G -pacbio-raw $PB \
      -nanopore-raw $ONT2d "gridOptions=--time $T" \
      minReadLength=500 oeaMemory=8  corOutCoverage=$COROUTCOV \
      corMemory=30
```

Canu_10 = corcov500minrl500.aspb-e02.pball.ont2d = Canu 1.3 using all PacBio reads and 2D nanopore reads (LRS#4/PBall+ONT2d), specifying a minimum read length of 500 bp, fixing coverage parameters to use all reads (not just a subset of the longest sampled by Canu), and combining the PB and ONT data into a single file treated by Canu as PacBio data:

```
NAME=g292-default-pball-ont2d-corcov500-min500-aspb
canu -p $NAME -d $NAME genomeSize=$G errorRate=0.02 \
      -pacbio-raw $PBall_ONT2d "gridOptions=--time $T" \
      minReadLength=500 oeaMemory=8 \
      corOutCoverage=$COROUTCOV corMemory=30
```

Canu_11 = corcov500.pbfilt.ont2d = Canu 1.3 using filtered PacBio reads and 2D nanopore reads (LRS#5/PBfilt+ONT2d), specifying a minimum read length of 500 bp, fixing coverage parameters to use all reads (not just a subset of the longest sampled by Canu):

```
NAME=g292-default-pbfilt-ont2d-corcov500-min500
canu -p $NAME -d $NAME genomeSize=$G -pacbio-raw $PBFILT \
      -nanopore-raw $ONT2d  "gridOptions=--time $T" minReadLength=500 \
      oeaMemory=8 corOutCoverage=$COROUTCOV corMemory=30
```

Canu_12 = corcov500minrl500.aspb-e02.pbfilt.ont2d = Canu 1.3 using filtered PacBio reads and 2D nanopore reads (LRS#5/PBfilt+ONT2d), specifying a minimum read length of 500 bp, fixing coverage parameters to use all reads (not just a subset of the longest sampled by Canu), and combining the PB and ONT data into a single file treated by Canu as PacBio data:

```
NAME=g292-default-pbfilt-ont2d-corcov500-min500-aspb
```

```
canu -p $NAME -d $NAME genomeSize=$G errorRate=0.02 \
      -pacbio-raw $PBfilt_ONT2d "gridOptions=--time $T" \
      minReadLength=500 oeaMemory=8 \
      corOutCoverage=$COROUTCOV corMemory=30
```

Canu_13 - 17 were run in similar ways as above with earlier versions of Canu. We used optimization scripts for the bogart step for a diploid genome: [https://github.com/JohnUrban/sciara-project-tools/blob/master/canu-utilities/optimization-loop-2.sh](https://github.com/JohnUrban/sciara-project-tools/blob/master/canu-utilities/optimization-loop-2.sh)

Canu_13 = canu10.minrl500.dip3x.pball = Canu 1.0 using PBall and a minimum read length of 500 bp.

Canu_14 = canu10.minrl1000.dip3x.pbfilt = Canu 1.0 using PBfilt, a minimum read length of 1000 bp, and the diploid 3X optimization.

Canu_15 = canu11.corcov80minrl500.dip3x.pball.ontmol = Canu 1.1 with PBall+ONTmol specifying a high coverage cutoff to use all or most of the data from reads >500 bp. The diploid optimization was run and we selected the B=3X assembly.

Canu_16 = canu12.corcov80minrl500.dip3x.pball.ontmol = Canu 1.2 with PBall+ONTmol specifying a high coverage cutoff to use all or most of the data from reads >500 bp. The diploid optimization was run and we selected the B=3X assembly.

Canu_17 = canu12.minrl500.dip3x.pball.ontmol = same as Canu_16, but with the default coverage cutoff.

###### 4.2.3.3 Falcon

Assemblies were generated using Long Read Sets #1-5. Similar to Canu, since Falcon was able to launch jobs on SLURM in an efficient way, we were able to test many different sets of parameters and combinations of reads. Twelve assemblies are reported in this study that could be referred to as:

With LRS#1/PBall:
- Falcon_1: default.PBall
- Falcon_2: seed25.PBall
- Falcon_3: seed30.PBall

With LRS#2/PBfilt:
- Falcon_4: default.PBfilt
- Falcon_5: seed25.PBfilt
- Falcon_6: seed30.PBfilt

With LRS#3/PBall+ONTmol
- Falcon_7: seed25.PBall.ontmol
- Falcon_8: seed25.relaxed.PBall.ONTmol

With LRS#4/PBall+ONT2d
- Falcon_9: seed25.PBall.ONT2d
- Falcon_10: seed25.relaxed.PBall.ONT2d

With LRS#5/PBfilt+ONT2d
- Falcon_11: seed25.PBfilt.ONT2d

Falcon_12: seed25.relaxed.PBfilt.ONT2d

The headers for fast5-derived fasta files from the MinION data are not appropriate for use with falcon, whereas PacBio headers are. To use our MinION data with the Falcon assembler, we used Fast5Tools to generate Falcon-acceptable headers as such:

```
filterFast5DerivedFastx.py -o falcon file.fa > file.forfalcon.fa
```

All assemblies were initiated via:

```
fc_run.py fc_run.cfg logging.ini
```

The "logging.ini" file for all Falcon assemblies contained these lines:

```
[loggers]
keys=root,pypeflow,fc_run

[handlers]
keys=stream,file_pypeflow,file_fc_run,file_all

[formatters]
keys=form01,form02

[logger_root]
level=NOTSET
handlers=stream,file_all

[logger_pypeflow]
level=NOTSET
handlers=file_pypeflow
qualname=pypeflow
propagate=1

[logger_pwatcher]
level=NOTSET
handlers=file_pwatcher
qualname=pwatcher
propagate=1

[logger_fc_run]
level=NOTSET
handlers=file_fc_run
qualname=fc_run
propagate=1

[handler_stream]
class=StreamHandler
level=INFO
formatter=form02
args=(sys.stderr,)

[handler_file_pypeflow]
class=FileHandler
level=DEBUG
formatter=form01
```

```
args=('pypeflow.log',)

[handler_file_pwatcher]
class=FileHandler
level=DEBUG
formatter=form01
args=('pwatcher.log',)

[handler_file_fc_run]
class=FileHandler
level=DEBUG
formatter=form01
args=('fc_run.log',)

[handler_file_all]
class=FileHandler
level=DEBUG
formatter=form01
args=('fc.log',)

[formatter_form01]
format=%(asctime)s - %(name)s:%(lineno)d - %(levelname)s - %(message)s

[formatter_form02]
format=[%(levelname)s] %(message)s
```

Falcon Default Configuration:
The “default” configuration file (fc_run.cfg) for “default.PBall” and " default.Pfilt " assemblies (Falcon_1 and Falcon_4) contained the following lines:

```
[General]

use_tmpdir = false
job_type = slurm
jobqueue = production
#stop_all_jobs_on_failure = true

# list of files of the initial bas.h5 files
input_fofn = input.fofn
input_type = raw

# The length cutoff used for seed reads used for initial mapping
length_cutoff = -1
genome_size = 292000000

# The length cutoff used for seed reads usef for pre-assembly (>0, was
not able to do -1)
length_cutoff_pr = 500

#Pre-Assembly
sge_option_da = --cpus-per-task 8 --mem 30g --time 48:000:00 --
qos=ccmb-condo
sge_option_la = --cpus-per-task 2 --mem 30g --time 48:00:00 --qos=ccmb-
condo
pa_DBsplit_option = -a -x500 -s200

pa_HPCdaligner_option = -v -B128 -e0.70 -M24 -l1000 -s100
pa_concurrent_jobs = 1000
```

```
#consensus for error correction
sge_option_cns = --cpus-per-task 8 --mem 60g --time 48:00:00 --
qos=ccmb-condo
falcon_sense_option = --output_multi --min_idt 0.70 --min_cov 4 --
max_n_read 200 --n_core 8
cns_concurrent_jobs = 1000

# overlap detection for assembly
sge_option_pda = --cpus-per-task 8 --mem 30g --time 48:00:00 --
qos=ccmb-condo
sge_option_pla = --cpus-per-task 2 --mem 30g --time 48:00:00 --
qos=ccmb-condo
ovlp_concurrent_jobs = 1000
ovlp_DBsplit_option = -x500 -s200
ovlp_HPCdaligner_option = -v -B128 -e.96 -M16 -l500 -s100

# overlap filtering
overlap_filtering_setting = --max_diff 40 --max_cov 80 --min_cov 2 --
n_core 12

sge_option_fc = --cpus-per-task 16 --mem 30g --time 48:00:00 --
qos=ccmb-condo
```

<u>Falcon Seed=25 Configuration</u>:
The “seed25” configuration files (fc_run.cfg) for “seed25.PBall”, “seed25.PBfilt”, “seed25.PBall.ontmol”, “seed25.PBall.ont2d”, “seed25.PBfilt.ont2d” (falcon assemblies #2, 5, 7, 9, and 11) were similar to the “default” configuration with the following line added:

```
seed_coverage = 25
```

<u>Falcon Seed=30 Configuration</u>:
The “seed25” configuration files (fc_run.cfg) for “seed25.PBall”, “seed25.PBfilt”, “seed25.PBall.ontmol”, “seed25.PBall.ont2d”, “seed25.PBfilt.ont2d” (falcon assemblies #2, 5, 7, 9, and 11) were similar to the “default” configuration with the following line added:

```
seed_coverage = 30
```

<u>Falcon Relaxed Seed=25 Configuration</u>:
The “relaxed seed 25” configuration file (fc_run.cfg) for “seed25.relaxed.PBall.ONTmol”, “seed25.relaxed.PBall.ONT2d”, and “seed25.relaxed.PBfilt.ONT2d” (Falcon assemblies 8, 10, and 12) was almost the same as “seed25” configuration above, but with the following lines added/changed:

### specified a length cut-off
```
length_cutoff = 7500
```

### In section titled “consensus for error correction”, changed --min_idt from 0.70 to 0.65:
```
falcon_sense_option = --output_multi --min_idt 0.65 --min_cov 4 --
max_n_read 200 --n_core 8
```

### In section titled "overlap detection for assembly", changed `-e` from `.96` to `.70`
```
ovlp_HPCdaligner_option = -v -B128 -e.70 -M16 -l500 -s100
```

###### 4.2.3.4 Miniasm and RaCon

Miniasm required < 64 GB RAM, 8 threads, and < 3 hours to complete. It was therefore practical to try many combinations of reads, filtering, etc. Since Miniasm does not have its own consensus step, RaCon was used. RaCon required < 150 GB RAM, 8 threads, and < 4.5 hours to finish. In total, Miniasm+RaCon took < 7.5 hours and is by far the fastest combination of assembly and consensus steps we tested. We generated eight miniasm assemblies using different combinations of reads.

Miniasm was used with Long Read Sets #1-8

RaCon used LRS#1/PBall for assemblies that used only PacBio data.
RaCon used LRS#6/PBallONTall for assemblies that used both PacBio and ONT data.

All Miniasm assemblies and RaCon polishing followed these commands:

```
minimap -Sw5 -L100 -m0 -t8 longreads.fasta longreads.fasta | \
      gzip -1 > reads.paf.gz

miniasm -f longreads.fasta reads.paf.gz > reads.gfa

awk '/^S/{print ">"$2"\n"$3}' reads.gfa > asm.fasta

minimap asm.fasta longreads.fastq > overlaps-for-racon.paf

racon -t 8 reads.fastq overlaps-for-racon.paf asm.fasta asm.racon.fasta
```

###### 4.2.3.5 SMARTdenovo

Five SMARTdenovo assemblies were generated using Long Read Sets #1-5.

```
smartdenovo.pl -c 1 longreads.fasta > wtasm.mak
make -f wtasm.mak
```

SMARTdenovo required <48 GB RAM, 8 threads, and <15 hours to complete. It was therefore practical to try more combinations of reads, filtering, etc.

#### 4.2.4 Polishing the Long Read Assemblies

Quiver polishing (Chin et al. 2013) was performed iteratively to take advantage of the possibility of more (and higher quality) alignments to the updated consensus sequence from each previous round under the assumption that the number of variants reported by Quiver in each round reflected, in part, the number of errors left to correct in addition to true variants.

In the first round, Quiver dropped out contigs that have no coverage from the raw PacBio reads and we chose to leave them out. We noticed that, although contigs were dropped from Canu, DBG2OLC, Falcon, and Miniasm assemblies, there were no contigs dropped from ABruijn and SMARTdenovo assemblies. Moreover, Quiver reports lower-case letters for bases that do not have enough coverage (< 5 here) to compute a consensus on whereas consensus-computed bases are uppercase. Most assemblers contained contigs that were 100% lower-case, though these were typically quite small (e.g. $\leq$ 20 kb). ABruin and SMARTdenovo did not contain any contigs that were reported as 100% lower-case. These initial observations suggested that ABruijn and SMARTdenovo these assemblers already dispensed with low coverage contigs whereas the other assemblers were more conservative and retained some.

For the 6th and 7th round of Quiver polishing we lowered the minimum coverage required for Quiver to compute a consensus and identify variants from 5 to 3. This did not seem to impact most metrics for most assemblies in a negative or positive way. However, ALE measures for Canu and Falcon seemed to degrade whereas the measures for all metrics except REAPR continued to improve for PlatanusDBG2OLC assemblies. Therefore, for each assembly (for all 50 assemblies), we compared the evaluation metrics from each Quiver round and selected the assembly version from the Quiver round with the highest average rank, opting for later rounds in cases of a tie. This resulted in selecting assemblies from Quiver rounds 3-7, with 13 and 19 of the 50 assemblies from rounds 5 and 7, respectively. Canu and Falcon assemblies tended to be selected for earlier rounds whereas ABruijn, Miniasm, PlatanusDBG2OLC, and SMARTdenovo assemblies tended to be selected in later rounds, seemingly reflecting whether or not the assemblers had pre-assembly read correction steps. After selecting the best version of each assembly from the Quiver rounds, all 50 assemblies were subject to 2 rounds of diploid-aware Pilon polishing with our Illumina dataset.

##### 4.2.4.1 Iterative polishing with signal-level PacBio data using Quiver

The basic approach was to align all raw PacBio reads using PBalign, merging and sorting the alignments, and using Quiver. Each round used the previous round's polished output assembly as the input assembly. The number of variants in the GFF output file was counted after each round.

```
pbalign $BAX $REF $OUTPRE.cmp.h5 --forQuiver --tmpDir $TMPDIR --nproc
$THREADS

cmph5tools.py merge --outFile ${MERGEDCMP} ${CMPDIR}/*.cmp.h5

cmph5tools.py sort --deep ${IN_CMP} --tmpDir ${TMP_SORT}

samtools faidx $REF

quiver -j${THREADS} ${INPUT} -r ${REF} -o $OUTGFF -o $OUTFASTQ -o
$OUTFASTA --noEvidenceConsensusCall=lowercasereference --verbose
```

Prior to all alignment/polishing steps, for Canu assemblies, contigs in the unassembled file that consisted of at least 2 reads were added back to the assembly, and for both Canu and Falcon assemblies, associated bubble contigs were added to the assembly. All contigs that were added back to the assemblies were given names to identify them after polishing for optional removal. For all assemblies that had spaces in the contig names, the first "word" before the first space was used as the contig name. All of these operations were accomplished with python scripts found at https://github.com/JohnUrban/sciara-project-tools:

"filterCanuFasta.py" and "fasta_name_changer.py".

This pipeline was automated for SLURM to process all 50 assemblies:
https://github.com/JohnUrban/sciara-project-tools/tree/master/slurmgear/pacbiotools/quiver

###### 4.2.4.2 Evaluations during Quiver Polishing

Before Quiver polishing and after each round, the assemblies were evaluated with BUSCO as well as the percentage of Illumina reads that mapped to the assembly, REAPR, LAP, $FRC^{bam}$, and ALE with the same input Illumina reads used for evaluating the short read assemblies to directly compare scores.

###### 2.2.4.3 Iterative polishing with Illumina short reads using Pilon

The basic approach was to build a bowtie2 index (Langmead and Salzberg 2012) for each assembly, map the paired-end illumina reads to each assembly with bowtie2, index the BAM alignment files with SAMtools, mark duplicates with Picard Tools, and use Pilon (Walker et al. 2014) to polish. Each round used the previous round's polished output assembly as the input assembly.

```
bowtie2-build $ASM $BASE

bowtie2 -p $P --very-sensitive -N 1 --minins 0 --maxins 1000 -x $BT2 -1
$R1 -2 $R2 | samtools sort --threads $P -o ${PRE}.bam

samtools index ${PRE}.bam

java -Xmx${JX} -jar $JAR MarkDuplicates INPUT=${PRE}.bam
OUTPUT=${PRE}.markdup.bam METRICS_FILE=${PRE}.metrics.txt
REMOVE_DUPLICATES=false ASSUME_SORTED=true

samtools index ${PRE}.markdup.bam

java -Xmx${JX} -jar $PILONJAR --genome $ASM --output $PRE --changes --
frags ${READS} --diploid --fix bases --nostrays
```

This Pilon pipeline was automated for SLURM to process all 50 assemblies using shell scripts also found at:
https://github.com/JohnUrban/sciara-project-tools/tree/master/slurmgear/pilon

#### 4.2.5 Evaluations of the Initial 50 long read assemblies

We used the same evaluations as for short read assemblies as well as additional evaluations with the long read and optical maps datasets. These metrics included the percent of reads that aligned, the average number of split alignments per read, and the number and size of structural variations (SVs) detected in the assemblies by Sniffles (Sedlazeck et al. 2018). SVs serve as a proxy for the number and extent of mis-assemblies since, although all assemblies may share some real SVs, increasing numbers of mis-assembled regions will result in higher numbers of SVs detected. We aligned the raw optical maps to the assemblies, and used alignment statistics (number, scores, coverage, and span) as proxies for evaluating the long-range structural integrity of each assembly as described below.

##### 4.2.5.1 Size statistics

Used asm-stats.py from https://github.com/JohnUrban/sciara-project-tools

We looked at four statistics:

- The number of contigs
- The maximum contig length
- The NG50 given the expected genome size of 280 Mb
    - Half of the expected genome size is contained on contigs of this length and longer
- The LG50
    - Half of the expected genome size is on this many contigs
- The expected contig length (E) given the expected genome length
    - The expected contig length statistic is an alternative to NG50 (Salzberg et al. 2012)

##### 4.2.5.2 BUSCO completeness metrics

###### 4.2.5.3.1 BUSCOv1

```
BUSCO_v1.22.py -in $ASM -o $OUT -l $BV1_LINEAGE -m $BV1_MODE --cpu $CPU
```

###### 4.2.5.3.2 BUSCOv3 with ODB9

```
LINEAGE=/Path/to/diptera_odb9
run_BUSCO.py --in $ASM -o $OUT -l $LINEAGE -m $BV3_MODE --cpu $CPU --limit
$REGIONLIMIT
```

We also tried Eukaryota, Metazoa, Arthropoda, Insecta, and Endopterygota. Results were similar in all cases. We used the Diptera results as it had the largest set of BUSCOs.

###### 4.2.5.3 Illumina Metrics

###### 4.2.5.3.1 Mapping Illumina reads back to each assembly
Same as for Illumina assemblies.
Percent Illumina reads mapped taken from Bowtie2 output.

###### 4.2.5.3.2 ALE

```
ALE ${BAM} $REF ${BASE}.ALE.txt
```

###### 4.2.5.3.3 FRC$^{bam}$

```
FRC --pe-sam $BAM --pe-max-insert 800 --genome-size 292000000 --output
${BASE}.frc
```

We looked at both the number of features detected as well as the rate, number of features per megabase to normalize by assembly size.

###### 4.2.5.3.4 LAP

```
calc_prob.py -p $P -a $REF  -q -1 $R1 -2 $R2 -X 800 -I 0 -o fr -m 432 -
t 75 -b $BT2 > ${BASE}.prob

sum_prob.py -i ${BASE}.prob > ${BASE}.lapscore
```

###### 4.2.5.3.5 REAPR

```
reapr facheck $REF ${BASE}_renamed
reapr perfectmap ${BASE}_renamed.fa $R1 $R2 432 perfect
reapr smaltmap ${BASE}_renamed.fa $R1 $R2 mapped.bam  -n $P
reapr pipeline ${BASE}_renamed.fa mapped.bam output_directory perfect

zcat 03.score.per_base.gz | awk '{s+=$3}END{print s/NR} > per-base-
mean-score.txt
```

We looked at:
- Percent error-free bases
- Mean base score

###### 4.2.5.3.6 Pilon
We looked at the number of changes made in the final round of Pilon polishing (Walker et al. 2014) with the assumption that fewer changes needed to polish the assembly, the higher the quality of the assembly consensus. We found that re-running Pilon to count the number of variants was correlated with the number of changes made (assemblies with fewer variants generally needed fewer changes). For subsequent evaluations we also looked at the number of confirmed bases from Pilon's output in each assembly as a percent of the expected genome size (292 Mb).

###### 4.2.5.4 PacBio and MinION long read metrics

We used the long reads from PacBio and Oxford Nanopore both separately and in combination to evaluate the 50 assemblies. We used the long read aligner BWA (Li and Durbin 2009) and the structural variation (SV) algorithm called "Sniffles" (Sedlazeck et al. 2018) [https://github.com/fritzsedlazeck/Sniffles] to detect the number SVs in each assembly. Reasoning that the number of SVs is a combination of both real variants and errors in the assemblies, assemblies were ranked higher for having lower numbers of reported SVs. We also separately considered the sum of the interval lengths reported to be involved in SVs and refer to this as SV span. For the same reasoning as above, assemblies were ranked higher for having shorter SV spans. In some cases, assemblies can have short spans, but high numbers of reported SVs and vice versa. We called SVs separately for PacBio and ONT reads as well as when combining them.

```
# Index assembly with BWA
bwa index $ASM -p $BASE

# Align reads to assembly with BWA
## TYPE=pacbio or TYPE=ont2d
bwa mem -t $MTHREADS -M -x $TYPE $BWAIDX $FASTQ | samtools sort -T
$TYPE --threads $MTHREADS -o $TYPE.bam

# Optionally merge pacbio and ont alignments
samtools merge --threads $P combined.bam $PBBAM $ONTBAM

# Call SVs with Sniffles
# BAM=pacbio.bam or BAM=ont.bam or BAM=combined.bam
sniffles -m $BAM -b $BEDPE.bedpe
```

We additionally considered (i) the percentage of PacBio and Oxford Nanopore reads that aligned to each assembly, ranking assemblies higher for having a higher percentage, and (ii) the ratio of the number of alignments to the number of unique reads represented in those alignments. The latter can be thought of as the expected number of split alignments per read. Since reads are sometimes split to align in separate places or in different orientations, the ratio is nearly always greater than 1. Similar to the more formal SV results from Sniffles above, we reasoned that the number of split alignments for a given read is a result from true biological variation, errors in the reads or alignments, and mis-assemblies. Since the first two are constant across all conditions, the only variable is the number of mis-assemblies. Thus, the closer to 1 this ratio is, the lower the rate of split alignments, and the higher a given assembly was ranked. This pipeline was automated using shell scripts found at https://github.com/JohnUrban/sciara-project-tools.

###### 4.2.5.5 BioNano optical map metrics

We used the restriction map aligner called "maligner" (Mendelowitz et al. 2015) to align the raw BioNano optical maps to all 50 assemblies. Maligner reports an "M-score" for each alignment. We ranked assemblies using combined M-scores, total span across the assembly from alignments (sum of coordinates with at least 1 alignment over it), total coverage as determined by summing the reference interval lengths for each alignment, and the number of BioNano maps that could be aligned.

Aligning BioNano Maps with Maligner

### Merge all RawMolecule BNX files (all.bnx) using RefAligner utility from BioNano Genomics

```
./RefAligner -bnx -merge -i scia_copr_2013_031_1P_2016-04-
13_15_45/Detect\ Molecules/RawMolecules.bnx -i
Scia_copr_2013_032_1P_2016-04-12_10_39/Detect\
Molecules/RawMolecules.bnx -i Scia_copr_2013_032_1P_2016-04-
12_15_55/Detect\ Molecules/RawMolecules.bnx -i
Scia_copr_2013_032_1P_2016-04-13_11_27/Detect\
Molecules/RawMolecules.bnx -i Scia_copr_2013_032_1P_2016-04-
14_11_49/Detect\ Molecules/RawMolecules.bnx -o all -minSNR 2.75 -minlen
150 -minsites 8 -MaxIntensity 0.6
```

### Convert BNX to Maligner input (bnx2maligner.py from https://github.com/JohnUrban/sciara-project-tools )

```
bnx2maligner.py -b all.bnx > bionano.maps
```

### smooth the maps for maligner alignment using utilities from Maligner

```
smooth_maps_file -m 1000 bionano.maps > bionano.smoothed.maps
```

### Split up maps to align them to assemblies in parallel (split into 48 files)

```
mkdir split
cd split
split -l 4079 -d ../bionano.smoothed.maps bionano.smoothed.maps
```

### Convert assembly to smoothed map for Maligner using Maligner utilities (BssSI = CACGAG)

```
	make_insilico_map -o $ASM_OUT_PFX $ASM_FASTA CACGAG
smooth_maps_file -m 1000 ${ASM_OUT_PFX.maps > ${ASM_OUT_PFX.smoothed.maps
```

###### 4.2.5.6 Selection for BioNano scaffolding

To select a final subset of assemblies for BioNano hybrid scaffolding, we sorted the assemblies by taking mean ranks across combinations of all 27 evaluation metrics used on the assemblies. Since it was possible that some assemblies have higher mean ranks due to a intrinsic weighting biases present across this set of 27 evaluations, we tried 40 different combinations of the 27 metrics with as few as 6 metrics in one combination and with as many as all 27 in another. The first twenty combinations did not include the size statistics ranks of NG50, LG50, expected size, and longest contig. This was done to uncover assemblies that ranked high without giving preference to more contiguous assemblies. The second set of 20 combinations did include size rankings. For both the first and second sets of twenty, the trend was roughly to use fewer metrics from the first to last combination. Moreover, since different categories (BUSCO, Illumina, PacBio, ONT, BioNano, size statistics) tended to favor different assemblers (mainly Canu or Falcon), the trend was to converge to more similar numbers of representatives from each category. At least one metric from each category was represented in all combinations except in the first twenty where size statistics were excluded.

###### 4.2.5.7 Automating the battery of metrics

As there were many assemblies and a battery of metrics, JMU created scripts to automate launching jobs in SLURM to do them. This eventually evolved into a tool called "Battery" (https://github.com/JohnUrban/battery) that works with the SLURM job manager to launch all the jobs needed for each evaluation metric (with jobs further along a pipeline given dependencies on upstream jobs completing successfully) given a list of assemblies. The 'battery' tool includes the metrics we used in this paper in addition to exploratory options, such as using REAPR with long reads, that we introduced later. This tool (Battery) was designed only to facilitate our own analyses and will not necessarily work out of the box on other systems where SLURM is configured differently. Others interested in using Battery may also need to change some hard-coded defaults. The outputs from Battery can be collected, then analyzed and visualized in R.

##### 4.2.6 BioNano Scaffolding

**4.2.6.1 BioNano CMAP assembly**:

BioNano genomic consensus maps (CMAPs) were assembled using an overlap layout consensus method with either all molecules >100 kb or all molecules >150 kb as input. A p-value threshold of 2.6e-9 was used with BioNano Pipeline Version 2884 and RefAligner Version 2816 (BioNano Genomics). The genome-wide optical maps assembled with the molecules >150 kb were slightly longer than when using all molecules >100 kb, with a map N50 of 712 kb and a cumulative length of 325.5 Mb. Thus, we chose to use only molecules >150 kb. There were 585 consensus genome maps (CMAPs) constructed from the BioNano image data, using the Canu assembly as a reference to estimate noise parameters during construction. Canu long read assembly contigs were converted into in silico maps and aligned to the BioNano CMAPS revealing a 266 Mb breadth of alignment (unique alignments) and 278 Mb total length of alignment (which includes redundant alignments of possibly repetitive DNA). A majority of unique alignments as seen here typically indicates a high quality sequence assembly.

**4.2.6.2 BioNano hybrid scaffolding:**

In hybrid scaffolding, BioNano CMAPs (assembled above) are linked when supported by long read contig overlaps, and long read contig maps are linked when supported by BioNano CMAP overlaps. The result is a genome-wide map, or hybrid scaffold, that is typically more contiguous than either of the input maps. The long read sequence assemblies and BNG optical maps were used with hybridScaffold.pl version 4741 (BioNano Genomics) to create genome-wide hybrid scaffolds. BssSI restriction maps of long read sequence contigs were generated in silico. In silico Consensus Maps (in silico CMAP) were only created for scaffolds > 20 kbp that contained > 5 BssSI sites. A p-value of 1e-10 was used as a minimum confidence value to output initial alignments (BNG CMAP to in silico CMAP) and final alignments (in silico CMAP to final hybrid CMAP). A p-value of 1e-13 was used as minimum confidence value to flag chimeric/conflicting alignments and to merge alignments. The final assemblies include all super-scaffolds from hybridScaffold.pl and contigs that were not super-scaffolded.

**4.2.6.3 Obtaining gap intervals for gap sizing**:

We used a custom script from https://github.com/JohnUrban/sciara-project-tools to define gap intervals. Since gap intervals from BNG scaffolding include recognition sequences we merged gaps were within 50 bp of each other using BEDTools.

```
scf-N-to-BED.py $f | sortBed -i - | mergeBed -i - -d 50
```

###### 4.2.6.4 Scaffold comparisons and evaluation

Scaffolded assemblies were evaluated with metrics as was done for the initial assembly evaluations above (automated with Battery as described above). Assemblies were aligned together using Minimap2:

```
minimap2 --secondary=no -x asm20 ${A} ${B} B-to-A-asm20-nosecondary.paf
```

Overall, the metrics and assembly alignments indicated that scaffolds derived from the same assembler were very similar. Only one representative of each was chosen to move forward with. We also noted that many of the BioNano joins unique to one assembly were already present as a single longer contig in the other supporting the BioNano join.

##### <u>4.2.7 Assembly refining</u>: Gap-filling, polishing, contamination removal, and haplotig identification

###### 4.2.7.1 Final polishing, gap-filling, and meta-scaffolding

Scaffolds were iteratively polished and gap-filled as described below. Polishing steps were used to polish newly filled gaps and facilitate further gap filling.

Upper case note: Some steps, such as Quiver polishing, output assemblies that include both lower- and upper-case letters. Since some programs (e.g. BLAST) may treat upper- and lower-case letters differently, we converted assemblies to 100% uppercase prior to BNG scaffolding and between all rounds of polishing, gap-filling, and meta-scaffolding, and before any/all evaluations:

```
fastaCaseMaker.py -f input.fasta > output.fasta
```

**First set of Quiver rounds on hybrid scaffolds**:

We worked with two sets of BioNano hybrid-scaffolded long read assemblies: one from Canu, and one from Falcon. Both were iteratively polished and gap-filled in the following way. Bionano scaffolds were polished with Quiver and the PacBio datasets five times to help correct the consensus near gaps. In the first round, Quiver detected 2177 and 3167 in the Canu and Falcon assemblies respectively, whereas there were 1723 and 2338 variants detected in each after the fifth round. Thus, the number of variants were similar to that found prior to BioNano scaffolding as one might expect.

Operations were the same as in section "**4.2.4.1 Iterative polishing with signal-level PacBio data using Quiver**".

**First set of PBJelly rounds on Quiver-polished hybrid scaffolds**:

The hybrid scaffolds were then subject to three rounds of gap filling with PBJelly (PBSuite_15.8.24) (English et al. 2012), using the combined set of PacBio and MinION reads in a fastq file. PBJelly was instructed to use only BLASR alignments with 75% identity or greater, and to only fill N-gaps (i.e. not attempt to further scaffold the assemblies).

BLASR instructions in Protocol.xml for all 3 rounds:

```
-minMatch 8 -sdpTupleSize 8 -minPctIdentity 75 -bestn 1 -nCandidates 10
-maxScore -500 -nproc 48 -noSplitSubreads
```

Commands:

```
Jelly.py setup Protocol.xml
Jelly.py mapping Protocol.xml -x "-n 20"
Jelly.py support Protocol.xml -x "--capturedOnly"
Jelly.py assembly Protocol.xml -x "--nproc=20"
Jelly.py output Protocol.xml
```

**Second set of Quiver rounds**:
To polish the consensus of the filled-gaps, we again ran Quiver for 3 rounds (Canu) or 4 rounds (Falcon). In the first round, Quiver detected 4369 and 8360 in the Canu and Falcon assemblies respectively, with the increases detected here resulting from new alignment regions within filled and partially-filled gaps. There were 1564 and 2962 variants detected in each after the final rounds, back to numbers similar to those pre-gap filling.

Operations were the same as in section "**4.2.4.1 Iterative polishing with signal-level PacBio data using Quiver**".

**Second set of PBJelly rounds**:
Next, given that the filled-in and and partially-filled-in gaps were now polished, we did three extra rounds of PBJelly gap filling. In the first round, PBJelly was instructed exactly as above (to use only BLASR (Chaisson and Tesler 2012) alignments with 75% identity or greater, and to only fill N-gaps). In the second round, that was adjusted to use BLASR alignments with 65% identity or greater, but otherwise the same. In the third round, 65% identity was again used as the cutoff, but we allowed PBJelly to link contigs together in this round (i.e. gap filling and scaffolding). In this last round, 83 of the 88 "filled" were contig joins for the Canu-derived assembly and 27 of the 45 "filled" were contig joins for the Falcon-derived assembly.

BLASR options in round 1:
```
-minMatch 8 -sdpTupleSize 8 -minPctIdentity 75 -bestn 1 -nCandidates 10
-maxScore -500 -nproc 48 -noSplitSubreads
```

BLASR options in rounds 2 and 3:
```
-minMatch 8 -sdpTupleSize 8 -minPctIdentity 65 -bestn 1 -nCandidates 10
-maxScore -500 -nproc 48 -noSplitSubreads
```

Commands for rounds 1 and 2:
```
Jelly.py setup Protocol.xml
Jelly.py mapping Protocol.xml -x "-n 20"
Jelly.py support Protocol.xml -x "--capturedOnly"
Jelly.py assembly Protocol.xml -x "--nproc=20"
Jelly.py output Protocol.xml
```

Commands for round 3 differed only at the "support" step where "--capturedOnly" was left off to allow meta-scaffolding:
```
Jelly.py support Protocol.xml
```

**Final Rounds of Quiver and Pilon Polishing**:
The updated gap-filled scaffolds were polished with another three rounds of Quiver and twelve rounds of Pilon. Generally speaking, for both Quiver and Pilon, the polishing rounds after the first round resulted in relatively few additional changes. In the first round, Quiver detected 4408 and 10992 variants in the Canu and Falcon assemblies respectively, with the increases detected here again resulting from new alignment regions within filled and partially-filled gaps. Again, there were similar numbers of variants in the final Quiver round to the quantities seen prior to any gap-filling (1652 and 2488 for Canu and Falcon, respectively). We also monitored the number of changes

to the consensus sequence in each Pilon round. In the first, there were 23361 and 20638 changes to the consensus for Canu and Falcon, but only 18 and 27 in the final round. This translates to approximately 1 change per 16.9 Mb and per 11 Mb of non-gap sequence in the final round for Canu and Falcon, respectively.

**Quiver**: Operations were the same as in section “2.2.4.1 Iterative polishing with signal-level PacBio data using Quiver”. Three rounds were performed.

**Pilon:** Operations were the same as in section “2.2.4.2 Iterative polishing with Illumina short reads using Pilon”. Twelve rounds were performed.

Overall, we observed that various metrics improved from the scaffolds to the meta-scaffolds. For example, the number of “translocations” detected by Sniffles as well as the number of short and long reads spanning more than one contig all went down.

###### 4.2.7.2 Identifying contaminating contigs and scaffolds

**Labeling bacterial contigs:**
Bacterial and other contaminating contigs and scaffolds were independently identified in both the final Canu and Falcon scaffolds in the following way. Here we will refer to all separate sequences in the final scaffolded assembly as “scaffolds” even though some contigs were not scaffolded, and might more correctly be still called contigs.

The final scaffolds were first split up into one scaffold per fasta file:

```
multiFasta2manyFastas.py -f ${ASM} -d ${TIGDIR}
```

This was done such that issues with BLAST (Altschul et al. 1990) on one contig, such as hanging on one contig until it was killed for exceeding an allotted time limit on SLURM, would not affect the the analysis of subsequent contigs in a file.

Next each scaffold was broken up into 1000 bp windows, and the windows were partitioned into their own separate files with a maximum of 292 windows per FASTA file. This was done for similar reasons as were provided above for splitting the assembly into its component scaffolds. Specifically, it was done to prevent situations where one area of a scaffold causing BLAST issues prevents adequate taxonomy information being collected for that scaffold. Moreover, it exhaustively collects taxonomy information in discrete 1 kb intervals across the scaffold, allowing us to confirm that each contig component of a scaffold was in agreement with taxonomy identification later on. For example, we could make sure that a scaffold labeled as BlobTools (Laetsch and Blaxter 2017) as Arthropod/Dipteran did not contain a bacterial contig within it that was erroneously joined during BioNano or long read scaffolding steps.

For each scaffold file, the following was done:

```
# There were giant N-gap sequences that were pre-filtered out of the
windows
# First gaps were found
scf-N-to-BED.py ${ASM} > gap.bed
```

```
# Then the scaffold was windowed, and only windows that had less than
75% overlap with a gap were retained (allowing the windows nearest to
gap edges to have no less than 250 bp) using BEDTools.
bedtools makewindows -w 1000 -s 1000 -g scaf.size | intersectBed -f
0.75 -v -a - -b gap.bed > windows.bed

# The sequence for each retained window was then obtained in a single
FASTA (with BEDTools).
fastaFromBed -fi ${ASM} -bed windows.bed > windows.fasta

# The windowed scaffold FASTA was then split into multiple files such
that they contained no more than 292 window sequences for the given
scaffold.
splitFastA.py -f windows.fasta -n 292 -o winFaDir
```

Finally, all of the windowed fastas from all of the scaffolds were subject to independent BLASTN runs against the entire nucleotide database:

```
blastn -task blastn -query $Q -db $NT \
 -outfmt '6 qseqid staxids bitscore std sscinames sskingdoms stitle' \
 -culling_limit 5 \
 -num_threads 8 \
 -evalue 1e-10 \
 -out ${BLASTDIR}/${BASE}.blastout
```

The BLASTN results from all were then combined into a single file and processed to have the correct scaffold names and coordinates:

```
# Combine raw results
cat blast_results/* > all_windows.blastout

# Process results into a taxonomy file for BlobTools

awk 'OFS="\t" {gsub(/\ /,"_"); gsub(/__/,""); sub(/:/,"\t"); sub(/-
/,"\t"); print $1,$4,$5}' all_windows.blastout | sort -k1,1 -k2,2n -
k3,3nr > all_windows_blastn_NT_qseqTaxBit.txt

# Process results into a BED file of taxonomy annotations in 1 kb
windows across the assembly (for visual inspection, etc)

awk 'OFS="\t" {gsub(/\ /,"_"); gsub(/__/,""); sub(/:/,"\t"); sub(/-
/,"\t"); print $1,$2+$11-
1,$2+$12,$6";"$4";"$5";"$7";"$8";"$9";"$10";"$13"-
"$14";"$15";"$16";"$17";"$18}' all_windows.blastout | sortBed -i - >>
all_blastn_NT_annotation.bed
```

BlobTools was then run the same as for short read assemblies described in section 2.1.4 (*Contamination analyses for the selected Platanus short read assembly*). We looked at the taxonomy information when using all of the BLAST results as well as when removing any BLAST hit labeled as Bradysia coprophila (*Sciara*). Both of gave essentially identical taxonomy-labeling results at the phylum level with the only differences being that some contigs that only had Bradysia hits were now unlabeled. For both approaches and for both the final Canu and Falcon scaffolds, 96.6-98% of the assembly was labeled as Arthopoda, ~98% of which was also labeled as Dipteran. We also looked at coverage levels from our male embryo Illumina, PacBio, and MinION data. All gave similar BlobPlot results. Moreover, we looked at male pupal genome coverage levels from BioNano optical maps, noting that there were no bacterial-labeled contigs with any BioNano coverage (consistent with no bacterial-labeled contigs being scaffolded by this dataset). Finally, we also performed these BlobTools analyses on the component contigs of the scaffolds independently to ensure that there weren't scaffolds labeled as Arthropod that had a bacterial contig within them nonetheless. There were no instances of this. Bacterial and arthropod-labeled sequences scaffolded independently of each other.

Overall, we used only the taxonomy information to filter scaffolds. Bacterial-labeled scaffolds (predominantly labeled as Rickettsia) were removed from the final scaffolded assemblies. Everything else was kept in with the caveat that only arthropod-labeled scaffolds were used to train gene predictors and to find motifs associated with DNA modification (see those sections below for more details).

The custom python scripts used above and SLURM scripts to automate the analysis can be found at: https://github.com/JohnUrban/sciara-project-tools

###### 4.2.7.3 Identifying primary and associated sequences

**Identifying "remove_tigs" in the final assemblies**

As discussed previously (**Section 4.2.4.1 Iterative polishing with signal-level PacBio data using Quiver**), both Canu and Falcon assemblies were fed back additional contigs to ensure all haplotypes/etc were present to avoid forcing incorrect alignments that lead to errors in polishing. For Canu the additional contigs were contigs composed of 2 or more reads. For Falcon, these were the associated bubble contigs. While their names were marked for removal in early steps, subsequent scaffolding steps obfuscated the naming scheme. To re-identify them, we chose to align the original contigs marked for removal back to the final assemblies with minimap2 (Li 2018) to re-identify them therein:

```
X=asm10
minimap2 --secondary=no -x ${X} -c ${ASM} rmseq_tigs.fasta >
rmseq2step01scaffs-${X}.paf
```

Adjacent query alignments from the same query mapping to the same target were merged using our custom script: filterpaf.py (https://github.com/JohnUrban/sciara-project-tools), and further merged with BEDtools (Quinlan and Hall 2010).

```
filterpaf.py -i rmseq2step01scaffs-${X}.paf --merge > merge-${X}.paf

awk 'OFS="\t" {print $6,$8,$9,$7,$1,$3,$4,$2, $5, $10,$11,$12}' merge-
${X}.paf | sortBed -i - | mergeBed -d 1000 -i - -c 4,5,6,7,8,9,10,11,12 -o
distinct,collapse,min,sum,sum,collapse,sum,sum,mean | awk '{OFS="\t"; print
$5,$8,$6,$7,$9,$1,$4,$2,$3,$10,$11,$12}' > targetmerged-${X}.paf
```

We classified the merged alignments as being the "same" contig (i.e. it did not change much through scaffolding steps) or as a scaffold containing the contig. To be called "same" we required either (i) the smaller of the query or target contig sizes to be at least 80% the size of the longer one and required at least 50% identity overall or (ii) at least 60% for size and at least 60% for identity.

```
awk '($2/$7 >= 0.8 && $2/$7 <= 1.2 && $10/$2 >=0.5 && $10/$11 >= 0.5) ||
($2/$7 >= 0.6 && $2/$7 <= 1.66 && $10/$2 >=0.75 && $10/$11 >= 0.75)'
targetmerged-${X}.paf > same-${X}.paf
```

Other alignments were marked as containments:

```
cut -f 1 same-${X}.paf | tr "," "\n" | sort | uniq > found-${X}.txt
grep -w -f <(grep -v -w -f found-${X}.txt <(grep ">" rmseq_tigs.fasta ) | awk
'{sub(/>/,""); print}') targetmerged-${X}.paf | sort -k1,1 -k3,3n >
containments.paf
```

All 339 and 184 contigs marked for removal in the initial Canu and Falcon assemblies, respectively, were found in the final scaffolds mostly unchanged. For both, the majority of contigs remained as singletons through the scaffolding steps (i.e. were not part of longer scaffolds).

**Identifying candidate redundant contigs/haplotigs using Minimap2 alignments.**

Long read assemblies tend to output many relatively short contigs that map within longer contigs, and these might be considered redundant. Moreover, assemblies of diploid genomes can produce two contigs for heterozygous loci. Often associated/bubble contig are shorter than the primary contig they map inside (though these bubbles can also span > 100 kb). Falcon marks most of these in a separate file. Although we added them back, we re-identified them in the previous step. Therefore, we expect fewer ‘redundant contigs’ to be found in this step for Falcon than Canu, the latter which outputs all primary and associated contigs in a single fasta file.

We used Minimap2 to find mappings from all contigs to all other contigs. In latter steps we eliminated any self-self mappings present, merged nearby alignments from the same query on the same strand, and focused on shorter contigs that mapped inside longer contigs. For simplicity, we settled on “-x ava-ont” below. We found that this was the most sensitive way to identify candidate redundant contigs. However, “-x asm5”, “-x asm10”, and “-x asm20” combined yielded 3 additional hits. Manually inspecting them showed that they were typically on the borderline of what one might accept as redundant. We also tried other parameters such as both “-N 1000” and “-N 100 -p 1e-5 -f 0.001” with each “–x” value from above, and “-N 10000 -p 1e-5 -f 0.001” without specifying a “-x” value. Overall, we went with the option (`-x ava-ont`) that appeared to give our downstream steps the highest sensitivity, missing only 3 hits across all conditions combined.

For mapping:

```
minimap2 -c -t ${T} -N 1 -X -x ava-ont ${ASM} ${ASM} > ${B}
```

Alignments were then re-written such that every line had the longer contig first and shorter second, then clustered in a way to facilitate alignment merging:

```
awk '$1!=$6 {OFS="\t"; R=$2/$7; if (R>=1)
print $0; else print
$6,$7,$8,$9,$5,$1,$2,$3,$4,$10,$11,$12,$13,$14,$15,$16}' ${B} | sort -
k5,5 -k1,1 -k6,6 -k3,3n -k8,8n > sort.${B}

##rename
mv ${B} del.${B}
mv sort.${B} ${B}
```

A first pass of obtaining “redundant contig” names was performed by looking for alignments of shorter to longer contigs where at least 80% of the shorter contig was represented with at least 70% identity:

```
awk '($9-$8)/$7 >= 0.8 && $10/$11 >= 0.7' ${B} | cut -f 6 | sort | uniq >
names-pass-1.${B}
```

As a second pass, alignments were merged given the sort order above. The shorter contig alignments needed to occur on the same target. Gaps in the alignment along the query or target could not exceed 10 kb. Backtracking in a subsequent alignment, where the subsequent alignment starts at a position prior to the previous alignment's end position, could not exceed 10 kb, nor was it allowed to beyond the previous start site. To be considered as redundant, alignments (whether from merging or not) needed to span at least 80% of the shorter contig with at least 70% identity. Moreover, the ratio of the alignment size along the target and along the query had to be such that the shorter alignment was at least 70% the size of the longer alignment.

```
filterpaf.py --targetmerge --presorted --merge \
      --merge_rules 10e3,10e3,10e3,10e3,1 \
      -i ${B} > tmerge-10k-${B}

awk '($9-$8)/$7 >= 0.8 && $10/$11 > 0.7' tmerge-10k-${B} \
      > filt-tmerge-10k-${B}

awk '($9-$8)/($4-$3) >=0.7 && ($4-$3)/($9-$8) >=0.7' \
      filt-tmerge-10k-${B} | cut -f 6 |  sort | \
      uniq > names-similar-spans.filt-tmerge-10k-${B}
```

**Size statistics for redundant contigs found by this method:**

| | **Canu** | **Falcon** |
|---|---|---|
| **Number of contigs** | 336 | 348 |
| **Total length** | 5,571,839 | 5,111,051 |
| **Max contig size** | 53,932 | 109,991 |
| **Min contig size** | 1,134 | 889 |
| **Mean contig size** | 16,582.9 | 14,686.9 |
| **Median contig size** | 15,404.5 | 12,879.5 |
| **Contig N50** | 18,877 | 18,688 |
| **Contig L50** | 110 | 92 |
| **Expected contig size** | 20,371 | 23,840 |

For Canu, 190 of the 336 redundant contigs were also labeled as "remove tigs" from above, and these made up 45.6% of the total length.
For Falcon, 164 of the 348 redundant contigs were also labeled as "remove tigs" from above, and these made up 54.1% of the total length.

**Further exploration of candidate redundant contigs with incorporation of coverage from each dataset using "purge_haplotigs"**

a. PacBio reads were aligned to the assembly with Minimap2:

```
minimap2 -a -t ${THREADS} -x map-pb ${ASM} ${PACBIO} | \
samtools view -hF 256 | \
samtools sort -m 1G -T pacbio --threads $THREADS -o pacbio-minimap2.bam
```

b. Purge_haplotigs (Roach et al. 2018) was used to obtain coverage data:

```
purge_haplotigs  readhist  -b ${BAMPATH}  -g ${ASM} -t 12
```

c. The coverage data was used to define the means and standard deviations of the mixture of normal distributions seen in the data using mixtools in R. With these analyses we settled on appropriate coverage cutoffs for the next step.

d. Purge_haplotigs was used to flag contigs:
```
purge_haplotigs  contigcov  -i ${GENCOV}  -l 4  -m 33  -h 75  \
      -o coverage_stats.csv -j 80  -s 80
```

e. Purge_haplotigs was used to classify contigs as KEEP, HAPLOTIG, REPEAT, JUNK, and UNCLASSIFIED. Providing BED file with repeat locations from repeat annotation step did not change results.
```
purge_haplotigs  purge  -g ${ASM}  -c ${COVSTATS} -t ${THREADS} \
   -b ${BAM} -d [-repeats repeats.bed]
```

This approach largely agreed with our original minimap2 approach in step 3. Specifically, for both Canu and Falcon, 93-94% of the contigs called redundant in step 3 were identified as a category other than "KEEP" by purge_haplotigs (i.e. HAPLOTIG, JUNK, or REPEAT). As the "remove tigs" (Step 2) from Falcon were all associated haplotigs output in a separate FASTA file as part of the Falcon assembly process, we expected them to have been marked for removal by purge_haplotigs. In agreement with this expectation, of the 173 "remove tigs" not contained within a larger scaffold, 169 were identied as a class other than "KEEP", and all 173 were identified as "redundant" in step3.

**Assigning contigs to the primary or associated assembly.**
Overall, contigs labeled as "bacterial" are considered bacterial contigs and were not deposited on NCBI. Contigs that were not bacterial, not "remove contigs", not redundant, and were labeled as KEEP by purge_haplotigs are considered the primary set of contigs (primary assembly). All other non-bacterial sequences are considered "associated contigs". All primary and associated sequences were deposited on NCBI as a single assembly. Please contact us for a text file that partitions the contigs into primary and associated assemblies if interested.

**4.2.7.4 Anchoring sequences into chromosomes using known, localized, and/or homologous sequences**
The following previously known nucleotide sequences were aligned to the genome with BLAST:
- DNA Puff II/9A (DiBartolomeis and Gerbi 1989; Bienz-Tadmor et al. 1991; Urnov et al. 2002; Foulk et al. 2006)
- RNA Puff III/9B (Wu et al. 1993; Foulk et al. 2006)
- Ecdysone Receptor/minor DNA puff IV/12A (Foulk et al. 2013)
- Ultraspiracle/RNA puff IV/10A (Foulk et al. 2013)
- Hsp70 (Mok et al. 2001)
- ScoHet1 (Greciano et al. 2009)
- ScoHet2 (Greciano et al. 2009)
- rDNA (Pardue et al. 1970; Gerbi 1971; Crouse et al. 1977; Kerrebrock et al. 1989)
- B4/F7/Sccr (Escribá et al. 2011)
- Drosophila Mitochondrial Sequence (chrM from dm3/BDGP Release 5)

Regarding Hsp70:
In previous work (Mok et al. 2001), *Sciara* Hsp70 was cloned, though not sequenced, and hybridized to two sites on chromosome IV: regions 4A and 17C. All *Drosophila melanogaster*

Hsp70 protein sequences were downloaded from FlyBase and aligned to the *Sciara* genome assemblies with tblastn, and all identified the same genomic regions and mapped across at least 90% of their lengths. There were four Hsp70 regions on three different primary contigs in both the Canu and Falcon assemblies rather than the expected two regions. Both assemblies have two unanchored contigs with an Hsp70 site and two other Hsp70 regions on a single contig associated with chromosome II (identified with DNA puff II/9A sequence), the latter of which was surprising given the hybridization results showed only two regions on chromosome IV. This result was not due to a misjoin introduced in any of the scaffolding steps since puff II/9A and the two flanking Hsp70 regions were part of the same contig prior to scaffolding in both assemblies. It also seems unlikely to be a mis-assembled contig as the genome structure around this locus is consistent in long read assemblies from each algorithm we used. Since the hybridization results that mapped Hsp70 to two sites on chromosome IV would have been dependent on nucleotide sequence similarity to the probe and to each other, we reasoned that the two chromosome IV regions should cluster together by similarity in sequence alignments. To test this, the coordinates for the four Hsp70 genomic regions were used to extract the associated Hsp70 sequences with BEDtools (fastaFomBed) (Quinlan and Hall 2010). These were aligned to each other with BLAST using "-task megablast –qcov_hsp_perc X" where "X" was 75, 90, or 99. Under those conditions only sequences from the two unanchored two regions mapped to each other. Under more sensitive conditions (using "-task blastn" instead of "-task megablast and "–qcov_hsp_perc 75") we were able to get all to align. The two unanchored loci had the highest similarity, aligning across their full lengths at ~88.2% similarity. The two putative-chromosome-II Hsp70 sites each shared less than 70% DNA sequence similarity with any of the other three Hsp70 regions, one sharing only up to ~65% similarity to other Hsp70 sites across ~75% of its length. Since the hybridization results showed only two loci that must have been more similar to both the probe and to each other than to other Hsp70 sites, the most likely candidate locations in the genome assembly for the two loci seen in the hybridization results are the two yet-unanchored regions that share over 88% similarity with each other across their full lengths, and 70% or less similarity with other sites. Moreover, the lower similarity of the Hsp70-like sites anchored to chromosome II explains their absence in the hybridization results.

For rDNA, only small subsequences from *Sciara* are known. Therefore, in addition to these, a single unit of Drosophila rDNA (NW_007931121.1, positions 43508-55989) was used to identify all rDNA-containing contigs with BLAST (-evalue 1e-10 –qcov_hsp_perc 5).

Similarly, the Drosophila mitochondrial sequence (dm3) was used with Minimap2 (Li 2018) and BLAST (Altschul et al. 1990) to identify mitochondrial contigs. For the Canu assembly, Minimap2 identified a single candidate whereas BLAST identified four, one being the Minimap2 candidate. Two of the others were extraneous small contigs marked for removal and one was a tiny hit to a massive contig anchored to chromosome II. Of the three chrM candidates, all were considered JUNK by purge_haplotigs due to extremely high coverage. The minimap2 candidate was added back to primary assembly. For Falcon, Minimap2 identified three candidates whereas BLAST identified many more mitochondrial hits on many small contigs. We also inferred there must be a high number of mitochondrial sequences during repeat modeling when we noticed some de novo repeats from the Falcon assembly contained mitochondrial genes. We focused on the three Minimap2 hits. One was considered redundant and marked for removal in the above pruning analyses. Of the remaining two, one was more consistent with the structure from the Canu assembly. The other seemed to have inversions when compared to the Canu structure as well as to the other Falcon candidate. Therefore, the Falcon candidate most consistent with the Canu candidate was chosen to put back in the primary assembly.

###### 4.2.7.5 Classifying/anchoring assembled sequences as autosomal or X based on diploid vs haploid coverage levels

X-linked contigs were contigs from the primary set in the pipeline below that had haploid level coverage across 80% of their length after (i) using Minimap2 to map PacBio reads back to the given assembly, (ii) using BEDtools genomecov to get coverage distributions for each contig, and (iii) defining cutoffs for haploid and diploid coverage levels based on the genomic distribution of coverage levels. Otherwise, contigs/scaffolds were called as autosomal. We tried this on the full set of scaffolds as well as the filtered "primary" set of contigs and scaffolds. There were a few examples of contigs that appeared to have mostly haploid level coverage in the full set, but diploid level coverage when reads were mapped only to the primary set after removing haplotigs, etc. However, the majority of the total length of sequence labeled as X-linked at the end of the pipeline was the same in both cases.

```
# Map reads
minimap2 -a -t ${THREADS} -x map-pb ${PRIMARY_ASM} ${PACBIO} | samtools
view -hF 256 | samtools sort -m 1G -T pacbio --threads $THREADS -o
pacbio-minimap2.bam

# Get coverage
bedtools genomecov -split -ibam pacbio-minimap2.bam > cov-split.hist
```

The genomic coverage distribution was viewed in R. We observed what appeared to be approximately 3 modes between the coverage levels of 0 and 65 (not unlike in step 4 above). We used the normalmixEM() from the "mixtools" package in R to learn the means and standard deviations of the three mixed distributions. The diploid coverage mean was between 40-42 (standard deviation ~5) and haploid coverage mean was approximately half the diploid mean as expected at ~20-22 (sigma ~8.2). The third mode was near 3-4x coverage, and was modeled to have mean of 3.2 (sigma ~2.2). We chose to set the lower cutoff to approximately three standard deviations below the haploid mean (7), the mid cutoff between haploid and diploid to 32 (~2 sigma above haploid mean and a little over 1 standard deviation under the diploid mean), and the high cutoff to around 4 standard deviations above the diploid mean (74).

The output from "bedtools genomecov" was parsed through to determine the proportion of sequence bases that had 0 coverage (0 coverage due to gaps was subtracted out), that were between 1 and the low cutoff, that were between the low cutoff and mid cutoff (haploid), that were between the mid cutoff and high cutoff (diploid), and that were above the diploid cutoff (collapsed repeat). If the majority of sequence bases were attributed to the haploid coverage level, we consider it a putative X. If $\geq$ 80% of the sequence bases for a contig were attributed to haploid coverage, we considered it a confident X call. We only considered confident X calls as X-linked/anchored moving forward.

### 4.3 Gene and Repeat annotation

#### 4.3.1 Transcriptome Assemblies

##### 4.3.1.1 De novo Transcriptome assembly using Trinity

A total of 18 strand-specific, 100 bp paired-end RNA-seq datasets were collected (see Supplemental Table S2). For each sex there were three mixed stage embryo replicates, and two replicates each of 4$^{th}$ instar larvae, mixed stage pupae, and mixed adults. All 18 samples were combined to generate a single transcriptome assembly with Trinity (version: trinityrnaseq_r20140413p1) (Grabherr et al. 2011) that we refer to as the “general transcriptome”.

```
Trinity --seqType fq --JM $JM --left $L --right $R --SS_lib_type RF --
CPU $P --trimmomatic --quality_trimming_params
"ILLUMINACLIP:/Path/to/neb_primers.fa:2:30:10 LEADING:5 TRAILING:5
MINLEN:36" --bflyHeapSpaceMax 28G --bflyCPU 8 --bflyHeapSpaceInit 10G
```

This transcriptome assembly was used both in genome assembly evaluations and in gene annotation.

##### 4.3.1.2 Genome-guided transcriptome assembly using HiSat2 and Stringtie

We had 18 RNA-seq samples as described in Supplemental Table S2. For assembling with StringTie (Pertea et al. 2015), we tried a few approaches and used BUSCO (Simão et al. 2015), RSEM-Eval (Li et al. 2014), and Transrate (Smith-Unna et al. 2016b), to help choose an assembly for use with Maker2 (Holt and Yandell 2011) gene annotation below. First, each RNA-seq sample was independently aligned to either the the Canu or Falcon genome assembly (with all scaffolds and contigs present) with HiSat2 (Kim et al. 2019) and assembled with StringTie in the following way:

```
hisat2 --rna-strandness RF -p ${THREADS} --dta -x ${HIDX} -1 ${R1} -2
${R2} -S ${R}.sam
samtools sort -@ ${THREADS} -o ${R}.bam ${R}.sam
rm ${R}.sam
stringtie --rf -p ${THREADS} -o ${R}.gtf -l ${R} ${R}.bam
```

The independent assemblies were merged as following (approach 1):

```
readlink -f *.gtf > mergelist.txt
stringtie --merge -p ${THREADS} -o approach_1.gtf mergelist.txt
```

We also tried merging alignments prior to assembly the following way (approach 2):

```
samtools merge -@ ${THREADS} -b bam.fofn merged.bam
stringtie --rf -p ${THREADS} -o approach_2.gtf -l merged merged.bam
```

Finally, we also tested merging independent assemblies into the assembly from merged reads (approach 3):

```
readlink -f independent_asms/*.gtf > mergelist.txt
readlink -f asms_from_merged_reads.gtf >> mergelist.txt
stringtie --merge -c 0 -F 0 -T 0 -f 0 -p ${THREADS} -o approach_3.gtf
mergelist.txt
```

We tested these three approaches on both the Canu and Falcon assemblies, generating six total genome-guided transcriptome assemblies that we then pared down to only 2 (one for each assembly) using transcriptome evaluations.

Transcript sequences were obtained from the outputs by:

```
gffread ${MERGED} -w ${TRANS} -g ${ASM}
```

When needed, GTFs were converted to GFFs by:

```
gffread -E file.gtf -o- > file.gff
```

#### 4.3.2 Evaluations of transcriptome assemblies

We generated seven transcriptome assemblies: the de novo Trinity assembly and the six StringTie assemblies of which three were guided by the Canu genome assembly, and three were guided by the Falcon assembly. We evaluated the transcriptome assemblies using BUSCO (Simão et al. 2015), RSEM-Eval (Li et al. 2014), and TransRate (Smith-Unna et al. 2016) in the following ways.

**4.3.2.1 BUSCO** :
We determined transcriptome completeness by searching for the 2799 Dipteran BUSCOs expected.

```
run_BUSCO.py --in ${ASM} -o busco -l ${DIPTERA} -m transcriptome --cpu
9 -sp fly -z --augustus_parameters="--progress=true"
```

**4.3.2.2 RSEM-Eval**:
Transcriptomes from Drosophila melanogaster and Anopheles were used to get transcript length parameters:

```
rsem-eval-estimate-transcript-length-distribution anopheles-gambiae-
pesttranscriptsagamp46.fa agam.txt

rsem-eval-estimate-transcript-length-distribution dmel-all-transcript-
r6.14.fasta dmel.txt

cat agam.txt dmel.txt | awk '{s+=$1; e+=$2}END{print s/NR"\t"e/NR}' >
agam_dmel_means.txt
```

**Parameters from each**:

| | MEAN TRANSCRIPT LENGTH | STANDARD DEVIATION |
|---|---|---|
| **DROSOPHILA** | 2881.22 | 2921.29 |
| **ANOPHELES** | 1860.19 | 1912.83 |
| **MEAN** | 2370.71 | 2417.06 |

SAMtools (Li et al. 2009) was used to estimate the average insert sizes for each RNA-seq sample from properly aligned read pairs, from reads aligned with HiSat2 for StringTie assemblies as described above. Given the set of average insert sizes from all samples, the average of averages (330 bp) was used.

```
samtools stats ${reads.bam} > ${b}.stats
```

Finally RSEM-Eval was run given (i) the Read1/Left and Read2/Right reads for all paired-end RNA-seq samples, (ii) the transcript length parameters estimated from Drosophila and Anopheles transcriptomes, and (iii) the insert size average estimated from all RNA-seq samples.

```
rsem-eval-calculate-score -p 9 --transcript-length-parameters
agam_dmel_means.txt --paired-end ${LEFT} ${RIGHT} ${ASM} Sciara 330
```

###### 4.3.2.3 TransRate:

TransRate (Smith-Unna et al. 2016b) can be provided a reference proteome to provide a completeness measure. For example, it reports the number and proportion of transcripts or reference proteins that contain Conditional Reciprocal Best BLAST (CRBB) hits. It also reports the proportion of amino acids in the reference proteome covered by transcripts. For direct comparison between transcriptome assemblies, we focused on the proportion of reference proteins with CRBB hits given each transcriptome assembly (similar to the number or proportion of complete BUSCOs) as well as the proportion of the reference proteome covered by transcripts from each assembly. As a reference proteome we tried both the entire set of proteins that we provided Maker2 during the annotation step as part of the homology evidence (see below) and a subset of that, which included all reviewed UniProt arthropoda proteins. Both references yielded the same trends. TransRate also calculates an assembly score based on alignments of the RNA-seq data. The Read1/Left and Read2/Right reads for all RNA-seq samples were provided for this.

```
transrate --assembly ${ASMS} --left ${LEFT} --right ${RIGHT} --threads
${NTHREADS} --reference ${REF}
```

#### 4.3.3 Building a comprehensive repeat library for use in Maker2

##### 4.3.3.1 *De novo* repeat libraries with RepeatModeler

Building a species-specific repeat library with RepeatModeler (Smit and Hubley 2008):
Repeat libraries were built on both the final Canu and Falcon scaffolds independently using RepeatModeler-open-1.0.11. The libraries were then combined. For each, the following commands were used.

```
BuildDatabase -name ${DBPRE} -engine ncbi ${ASM}

RepeatModeler -engine ncbi -pa 24 -database ${DBPRE} 2>&1 | tee
repeatmodeler.log
```

The resulting RepeatModeler libraries from both assemblies were then combined. Canu contained 2695 repeat families (2,267 Unknown), and Falcon contained 2,661 repeat families (2254 Unknown).

##### 4.3.3.2 Known *Sciara* repeats

Adding previously known repeat sequences from Bradysia (*Sciara*) coprophila:
To ensure the completeness of the species-specific repeat library, the following 27 known repeat sequences for Bradysia coprophila (including centromeric, rDNA, tandem repeat, and retrotransposon sequences) were added:

gi|160853|gb|L00951.1|SCIRTRANSB Sciara coprophila retrotransposable element R2 reverse transcriptase gene, 3' end
gi|160851|gb|L00945.1|SCIRTRANSA Sciara coprophila retrotransposable element R1 reverse transcriptase gene, 3' end
gi|332655545|gb|HQ883513.1| Bradysia coprophila Sccr15 centromeric repeat
gi|332655542|gb|HQ883512.1| Bradysia coprophila Sccr14 centromeric repeat
gi|332655541|gb|HQ883511.1| Bradysia coprophila Sccr13 centromeric repeat
gi|332655540|gb|HQ883510.1| Bradysia coprophila Sccr12 centromeric repeat
gi|332655537|gb|HQ883509.1| Bradysia coprophila Sccr11 centromeric repeat
gi|332655536|gb|HQ883508.1| Bradysia coprophila Sccr10 centromeric repeat
gi|332655532|gb|HQ883507.1| Bradysia coprophila Sccr09 centromeric repeat
gi|332655531|gb|HQ883506.1| Bradysia coprophila Sccr08 centromeric repeat
gi|332655530|gb|HQ883505.1| Bradysia coprophila Sccr07 centromeric repeat
gi|332655526|gb|HQ883504.1| Bradysia coprophila Sccr06 centromeric repeat
gi|332655525|gb|HQ883503.1| Bradysia coprophila Sccr05 centromeric repeat
gi|332655524|gb|HQ883502.1| Bradysia coprophila Sccr04 centromeric repeat
gi|332655520|gb|HQ883501.1| Bradysia coprophila Sccr03 centromeric repeat
gi|332655519|gb|HQ883500.1| Bradysia coprophila Sccr02 centromeric repeat
gi|332655518|gb|HQ883499.1| Bradysia coprophila Sccr01 centromeric repeat
gi|332655517|gb|HQ883498.1| Bradysia coprophila X.01F4 non-LTR retrotransposon ScRTE, partial sequence
gi|332655514|gb|HQ883497.1| Bradysia coprophila X.01F7 tandem repeat
gi|332655513|gb|HQ883496.1| Bradysia coprophila X.01B4 tandem repeat
gi|332655512|gb|HQ883495.1| Bradysia coprophila X.01G2 centromeric repeat
gi|332655506|gb|HQ883494.1| Bradysia coprophila clone lambda DASH II phage clone 8.4.3.2 non-LTR retrotransposon ScRTE, complete sequence
gi|10115|emb|V01340.1| Sciara coprophila genes for 5.8S and 2S ribosomal RNAs
gi|10114|emb|X02482.1| Sciara coprophila 28S-alpha/beta rDNA gap region
gi|2190334|emb|X93402.1| B.coprophila 28S rRNA gene, expansion segment D7
gi|2190333|emb|X93385.1| B.coprophila 28S rRNA gene, expansion segment D1
gi|2190332|emb|X93391.1| B.coprophila 28S rRNA gene, expansion segment D3-5

##### 4.3.3.3 Known Arthropod Repeats

Adding the arthropoda library from the RepeatMasker (Smit et al. 2013) Combined Database: Dfam_Consensus-20181026 (Hubley et al. 2016) and RepBase-20181026 (Bao et al. 2015)

In an early Maker trial run, despite specifying that Maker use “arthropoda” as the “species” argument to RepeatMasker, we noticed that only “Simple” and “Low Complexity” repeats were being identified and masked. There were no arthropoda repeats identified in the assembly although we could identify many repeats from the arthopoda library generated by RepeatMasker (RepeatMasker/Libraries/dc20181026-rb20181026/arthropoda/specieslib) in the assemblies using Minimap2. We tried running RepeatMasker directly and were able to reproduce the problem.

```
RepeatMasker ${ASM} -species arthropoda -dir soft -pa 1 –xsmall
```

When inspecting the arthropoda library at RepeatMasker/Libraries/dc20181026-rb20181026/arthropoda/specieslib, we noticed the sequences were entirely lower-case. We found that converting the sequences to upper-case resolved the issue completely. Therefore, we converted the arthropoda library to uppercase and added it to our species-specific library described above. When we ran Maker (described later), we provided this comprehensive combined species-specific and arthropoda library instead of specifying a species. We also generated soft-masked versions of the final Canu and Falcon scaffolds:

```
RMLIB=arthropoda_and_species-specific-repeat-library.fasta

RepeatMasker ${ASM} -lib ${RMLIB} -dir ./masked -xsmall -pa
${SLURM_NTASKS} -gff -html -source
```

#### 4.3.4 Maker2 gene annotation

##### 4.3.4.1 Overview:

Maker2 (Holt and Yandell 2011) was used for protein-coding gene annotation of the two final genome assembly candidates. The gene set for a given assembly was produced using gene predictors trained only on the given assembly. In all cases, training was performed using only contigs labeled as Arthropoda in the BlobTools analysis. GeneMark-ES (Ter-Hovhannisyan et al. 2008) was trained on soft-masked contigs. Augustus (Hoff and Stanke 2019) was trained using BUSCO v3 using the 2799 genes in the Dipteran lineage from OrthoDB v9. SNAP (Korf 2004; Campbell et al. 2014) was first trained using gene models output from the first round of Maker2 that used only transcript and protein evidence (est2genome=1, protein2genome=1), and then re-trained on the output of the second Maker round that included gene predictors as described below.

There were essentially three Maker2 rounds. Transcript evidence was provided to Maker2 as the Trinity de novo transcriptome and StringTie transcriptome constructed on the given genome assembly being annotated. Alternative transcript evidence was provided from fly species spanning the Nematocera and Brachycera suborders of Diptera, including the transcriptomes from *Drosophila melanogaster* and *Anopheles gambiae*. Protein homology evidence was provided using all Swiss-Prot proteins from Arthropoda (The UniProt Consortium 2019), as well as from proteomes spanning the holometaboulous insects, including *Drosophila melanogaster* (fruit fly), *Anopheles gambiae* (mosquito), *Apis mellifera* (honey bee), *Bombyx mori* (silkmoth), and *Tribolium castaneum* (red flour beetle). Maker2 was first run to create gene models from the transcript evidence alone to use as an initial training set for gene prediction with SNAP. For the second round of Maker2, gene models were created from the three gene predictors (SNAP, Augustus, GeneMark-ES) using the RNA-seq and homology evidence. SNAP was re-trained using the output of the second round, and a third round of Maker was run. In the third round, Maker2 was instructed to output all models and predictions (keep_preds=1). InterProScan (Quevillon et al. 2005) was subsequently run on all of the protein models to annotate them with protein domain information. Models were kept if they had an Annotation Edit Distance (AED) between 0 and 1 (inclusive) and/or had a recognizable protein domain as recommended (Campbell et al. 2014).

To rescue gene models buried inside masked repetitive regions, we also re-ran the third round of Maker2 exactly as above, but using a repeat library that was filtered to remove repeat models that contained genes, such as Odorant Receptor or Histone genes. We compared the Maker2 transcriptome outputs from using the original repeat library and the filtered repeat library by looking at their AED distributions, and using BUSCO, RSEM-Eval, and TransRate (see Supplemental Table S15A-B). For both Canu and Falcon, the gene sets produced using the original repeat library were higher quality as they had more genes, better AED distributions, more complete (e.g. more BUSCOs), and better evaluations. Therefore, we chose to use the transcript models generated using the original repeat library (stringent masking) as the primary set of annotations, and included transcript models generated when using the filtered repeat library (less masking) only if those gene structures had no overlaps with the primary set (using BEDtools intersect). This resulted in adding 1132 and 607 gene models to the primary Canu and Falcon gene model sets, respectively. We found that of these additional genes added back to the Canu set, for example, 30 were labeled as “Core histone H2A/H2B/H3/H4” and 11 were labeled as “7tm Odorant receptor”. Thus, we were able to rescue many genes originally modeled as repeats. Moreover, this resulted in an increased number of BUSCOs detected as well as higher scores in other metrics such as the RSEM-Eval Score.

The combined gene sets were then finalized for both the Canu and Falcon assemblies by removing gene models corresponding to contaminating (bacterial) contigs to give the final sets specific to *Sciara coprophila* (*Bradysia coprophila*). Gene models were named Bcop_v1_gXXXXXX and isoforms tagged with -RA/RB/RC/etc. Proteins were functionally annotated, where possible, by using InterProScan (Quevillon et al. 2005) for protein domain information and BLASTp against the entire UniProt/SwissProt protein database (Consortium 2019). The steps described here are detailed further below.

##### 4.3.4.2 Training GeneMark-ES for the first gene prediction round of Maker

GeneMark-ES (Ter-Hovhannisyan et al. 2008) was trained separately on the final Canu and Falcon scaffolds. For both, the scaffolds labeled as Arthropod in the BlobTools analysis were extracted from the soft-masked assemblies (soft-masked using the arthropoda and species-specific library as described above). These sequences made up over 96.6% of each assembly and the majority was also labeled as Dipteran (>98%). This was done to avoid training on any bacterial or any other contaminating sequences present. GeneMark-ES was trained on the soft-masked arthropod-labeled scaffolds with the following command:

```
gmes_petap.pl --ES --max_contig 50000000 --cores 9 --max_intron 500000
--sequence ${ASM} --soft_mask 5000
```

The output file produced located under the working directory at "./output/gmhmm.mod" was used as the GeneMark HMM in all Maker runs involving gene prediction (see below).

##### 4.3.4.3 Using BUSCO to train Augustus for the first gene prediction round of Maker

Augustus (Hoff and Stanke 2019) was trained separately on the final Canu and Falcon scaffolds using BUSCO (Simão et al. 2015). For both, we used BUSCOv3 with the Dipteran lineage from ODB9, specifying the "--long" option, and starting with "fly" as the species (for Augustus). "Canu" is used in the examples below – the same was done for Falcon.

```
run_BUSCO.py --in ${ASM} -o Sciara_canu -l ${DIPTERA} -m genome --cpu 9
--limit 10 --long -sp fly -z --augustus_parameters="--progress=true"
```

The optimized re-training parameters were then copied into the species directory for Augustus in subdirectory named "Sciara_long_unmasked_rl10_canu":

```
cp run_Sciara_canu/augustus_output/retraining_parameters/*
/Path/to/augustus-3.2.2/config/species/Sciara_long_unmasked_rl10_canu/
```

The files therein were then re-named such that their prefixes matched the directory name (Sciara_long_unmasked_rl10_canu) instead of the prefixes given by BUSCO. Finally, the innards of the "*_parameters.cfg*" files that pointed to those files needed to be changed from the BUSCO prefix to the new prefix as well.

The name "Sciara_long_unmasked_rl10_canu" was provided as the Augustus species in the first round of gene prediction in Maker (see below).

###### 4.3.4.4 Using Maker est2genome/protein2genome to train SNAP for the first gene prediction round of Maker

SNAP (Korf 2004) was trained separately on the final Canu and Falcon scaffolds starting with evidence-only gene models from Maker2 (Holt and Yandell 2011; Campbell et al. 2014). Maker2 is an annotation tool that can be run iteratively (with training and re-training of gene predictors) to improve annotations. In the first round of Maker (described in more detail in the Maker section below), we did not use any gene predictors, instead generating gene models from EST and protein evidence alone. We used our Trinity and StringTie assemblies for EST evidence and a comprehensive set of arthropod/insect proteins (detailed in the Maker section) as protein homology evidence. Maker gene models were then extracted from only scaffolds labeled as Arthropod/Diptera and used to train SNAP in the following way:

In a directory outside of the directory Maker was launched in:

```
ln -s /Path/to/*maker.output/*datastore .
grep -w -f arthropod_scaffs.txt
/Path/to/*.maker.output/*_master_datastore_index.log > arthropod.log
gff3_merge -o Sciara_rnd1.arthropod.gff -d arthropod.log

maker2zff Sciara_rnd1.arthropod.gff
fathom genome.ann genome.dna -gene-stats > gene-stats.log 2>&1
fathom genome.ann genome.dna -validate > validate.log 2>&1
fathom -categorize 1000 genome.ann genome.dna > categorize.log 2>&1
fathom -export 1000 -plus uni.ann uni.dna > uni-plus.log 2>&1
mkdir pars
cd pars
forge ../export.ann ../export.dna > ../forge.log 2>&1
cd ..
hmm-assembler.pl Sciara_rnd1 pars > Sciara_rnd1.hmm
```

The file “Sciara_rnd1.hmm” was provided to Maker for the first round of gene prediction (second Maker round).

###### 4.3.4.5 Repeat Libraries
Repeat masking inside Maker was performed with the combined set of repeats from (i) all Arthropoda in Dfam/RepBase (Bao et al. 2015; Hubley et al. 2016), (ii) the species-specific repeat library generated from RepeatModeler, and (iii) known repeat sequences. The “te_proteins.fasta” that comes with Maker2 was used as the set of repeat proteins for repeat masking.

###### 4.3.4.6 EST evidence
Both the Trinity and StringTie assemblies constructed from both sexes across multiple life stages were passed as EST evidence, Trinity as a fasta and StringTie as a GFF.

###### 4.3.4.7 Alternative EST evidence
The following 68,299 Dipteran transcript sequences were passed to Maker as alternative EST evidence.

- Nematocera suborder of Diptera
  - Bibionomorpha infraorder
    - Known *Bradysia coprophila* mRNA sequences from NCBI nucleotide database (txid38358; downloaded on Sep 01, 2016)
      - There were 14 mRNA sequences
    - All 22,152 mRNA sequences under Bibionomorpha from NCBI nucleotide/EST database (txid43784, downloaded on Dec 15, 2018)

            - Cecidomyiidae family (13,582 sequences; 5 species)
                - 449 Mayetiola avenae (Oat midge)
                - 458 Mayetiola hordei (Barley midge)
                - 1546 Sitodiplosis mosellana (Wheat midge)
                - 9866 Mayetiola destructor (Hessian fly)
                - 1263 Orseolia oryzae (Asian rice gall midge)
            - Sciaridae family
                - 8570 Rhynchosciara Americana
    - Culicomorpha infraorder
        - *Anopheles gambiae* transcriptome (downloaded May 4, 2017):
            - https://www.vectorbase.org/download/anopheles-gambiae-pesttranscriptsagamp46fagz
            - 15,648 sequences
- Brachycera suborder of Diptera
    - *Drosophila melanogaster* transcriptome (Downloaded Jan 20, 2015):
        - ftp://ftp.flybase.net/genomes/Drosophila_melanogaster/current/fasta/dmel-all-transcript-r6.03.fasta.gz
        - 30,485 sequences

##### 4.3.4.8 Protein homology evidence

The following protein sequences from the Arthropoda phylum were passed to Maker as protein homology evidence:

UniProt Arthropoda:
All protein sequences available for Arthropoda (the phylum in which the class Insecta resides) from UniProt (12,408 fasta entries) were passed to maker for protein homology evidence.

- https://www.uniprot.org/uniprot/?query=reviewed:yes%20taxonomy:6656
- This spanned 843 species from 463 genera with a median of 2 sequences per species and 3 per genus. There were 3541 sequences from Drosophila melanogaster, an order of magnitude higher than that from the next highest (Drosophila pseudoobscura). The UniProt Arthropoda dataset contained at most 7812 differently-named protein sequences, about half of which must have had an entry from Drosophila.

All other protein evidence came from representatives from four major orders of the Insecta class.

- Diptera order
    - Nematocera suborder
        - Bibionomorpha infraorder
            - Known *Bradysia coprophila* mRNA sequences from NCBI nucleotide database (txid38358;  downloaded on Sep 01, 2016)
                - There were 16 protein sequences
            - Rhynchosciara proteins
                - 20
        - Culicomorpha infraorder
            - *Anopheles gambiae* proteome (downloaded May 4, 2017):
                - https://www.vectorbase.org/download/anopheles-gambiae-pestpeptidesagamp46fagz
                - 14,909 sequences
    - Brachycera suborder
        - *Drosophila melanogaster* proteome (Downloaded Jan 20, 2015):
            - ftp://ftp.flybase.net/genomes/Drosophila_melanogaster/current/fasta/dmel-all-translation-r6.03.fasta.gz
            - 30,485 sequences

    - Drosophila represents a selection from the Culicomorpha infraorder under the Nematocera suborder of the order Diptera
- Hymenoptera order
    - *Apis mellifera* protein sequences
        - https://www.ncbi.nlm.nih.gov/protein; txid7460[Organism:exp]; RefSeq entries only; downloaded Dec 15, 2018
        - 23,491 protein sequences
- Lepidoptera order
    - *Bombyx mori* protein sequences
        - https://www.ncbi.nlm.nih.gov/protein; txid7091[Organism:noexp]; RefSeq entries only; downloaded Dec 15, 2018
        - 22,590 protein sequences
- Coleoptera order
    - *Tribolium castaneum* proteins
        - https://www.ncbi.nlm.nih.gov/protein?LinkName=bioproject_protein&from_uid=15718; clicking RefSeq; RefSeq entries only; downloaded Dec 15, 2018
        - 22,597 protein sequences

###### 4.3.4.9 Other parameter choices

- The maximum DNA length was set to 2 Mb so contigs and scaffolds would mostly not be split up.
- The minimum contig size was kept as default (1 bp).
- Maker2 was instructed to look for alternative splicing transcripts.
- Our choices for the minimum exon size (200 bp) and maximum intron size (500 kb) that Maker2 was instructed to expect were guided by the minimum exon and maximum intron sizes found in the StringTie transcriptome assembly.
- Similarly, Maker2 was instructed to use single exon ESTs as evidence given the that a fair number of transcripts appeared to be single exon (~3500; ~10%) from the StringTie assemblies.
- est2genome and protein2genome were both turned on in the first round of Maker2 to generate gene models strictly from EST and protein homology evidence for training SNAP prior to the second round of Maker2.

###### 4.3.4.10 Maker2 Round 1

##### The maker_opts.ctl file for Maker Round 1:

```
#-----Genome (these are always required)
genome=/path/to/asm.fasta #genome sequence (fasta file or fasta embeded
in GFF3 file)
organism_type=eukaryotic #eukaryotic or prokaryotic. Default is
eukaryotic

#-----Re-annotation Using MAKER Derived GFF3
maker_gff= #MAKER derived GFF3 file
est_pass=0 #use ESTs in maker_gff: 1 = yes, 0 = no
altest_pass=0 #use alternate organism ESTs in maker_gff: 1 = yes, 0 = no
protein_pass=0 #use protein alignments in maker_gff: 1 = yes, 0 = no
rm_pass=0 #use repeats in maker_gff: 1 = yes, 0 = no
model_pass=0 #use gene models in maker_gff: 1 = yes, 0 = no
pred_pass=0 #use ab-initio predictions in maker_gff: 1 = yes, 0 = no
other_pass=0 #passthrough anyything else in maker_gff: 1 = yes, 0 = no

#-----EST Evidence (for best results provide a file for at least one)
```

```
est=/Path/To/Trinity.fasta #set of ESTs or assembled mRNA-seq in fasta
format
altest=/Path/To/combined_transcript_sequences.fasta #EST/cDNA sequence
file in fasta format from an alternate organism
est_gff=/Path/To/stringtie_merged_Sciara_general_transcriptome.gff
#aligned ESTs or mRNA-seq from an external GFF3 file
altest_gff= #aligned ESTs from a closly relate species in GFF3 format

#-----Protein Homology Evidence (for best results provide a file for at
least one)
protein=/Path/To/combined_protein_sequences.fasta  #protein sequence file
in fasta format (i.e. from mutiple oransisms)
protein_gff=  #aligned protein homology evidence from an external GFF3
file

#-----Repeat Masking (leave values blank to skip repeat masking)
model_org= #select a model organism for RepBase masking in RepeatMasker
rmlib=/Path/to/combined_arthopoda_species-specific-library.fa #provide an
organism specific repeat library in fasta format for RepeatMasker
repeat_protein=/Path/To/maker/data/te_proteins.fasta #provide a fasta
file of transposable element proteins for RepeatRunner
rm_gff= #pre-identified repeat elements from an external GFF3 file
prok_rm=0 #forces MAKER to repeatmask prokaryotes (no reason to change
this), 1 = yes, 0 = no
softmask=1 #use soft-masking rather than hard-masking in BLAST (i.e. seg
and dust filtering)

#-----Gene Prediction
snaphmm= #SNAP HMM file
gmhmm= #GeneMark HMM file
augustus_species= #Augustus gene prediction species model
fgenesh_par_file= #FGENESH parameter file
pred_gff= #ab-initio predictions from an external GFF3 file
model_gff= #annotated gene models from an external GFF3 file (annotation
pass-through)
est2genome=1 #infer gene predictions directly from ESTs, 1 = yes, 0 = no
protein2genome=1 #infer predictions from protein homology, 1 = yes, 0 =
no
trna=0 #find tRNAs with tRNAscan, 1 = yes, 0 = no
snoscan_rrna= #rRNA file to have Snoscan find snoRNAs
unmask=0 #also run ab-initio prediction programs on unmasked sequence, 1
= yes, 0 = no

#-----Other Annotation Feature Types (features MAKER doesn't recognize)
other_gff= #extra features to pass-through to final MAKER generated GFF3
file

#-----External Application Behavior Options
alt_peptide=C #amino acid used to replace non-standard amino acids in
BLAST databases
cpus=1 #max number of cpus to use in BLAST and RepeatMasker (not for MPI,
leave 1 when using MPI)

#-----MAKER Behavior Options
max_dna_len=100000000 #100mil to notdivide up -- length for dividing up
contigs into chunks (increases/decreases memory usage)
min_contig=1 #skip genome contigs below this length (under 10kb are often
useless)
pred_flank=200 #flank for extending evidence clusters sent to gene
predictors
pred_stats=0 #report AED and QI statistics for all predictions as well as
```

```
models
AED_threshold=1 #Maximum Annotation Edit Distance allowed (bound by 0 and 1)
min_protein=0 #require at least this many amino acids in predicted proteins
alt_splice=1 #Take extra steps to try and find alternative splicing, 1 = yes, 0 = no
always_complete=0 #extra steps to force start and stop codons, 1 = yes, 0 = no
map_forward=0 #map names and attributes forward from old GFF3 genes, 1 = yes, 0 = no
keep_preds=0 #Concordance threshold to add unsupported gene prediction (bound by 0 and 1)
split_hit=500000 #length for the splitting of hits (expected max intron size for evidence alignments)
single_exon=1 #consider single exon EST evidence when generating annotations, 1 = yes, 0 = no
single_length=200 #min length required for single exon ESTs if 'single_exon is enabled'
correct_est_fusion=0 #limits use of ESTs in annotation to avoid fusion genes
tries=4 #number of times to try a contig if there is a failure for some reason
clean_try=0 #remove all data from previous run before retrying, 1 = yes, 0 = no
clean_up=0 #removes theVoid directory with individual analysis files, 1 = yes, 0 = no
TMP= #specify a directory other than the system default temporary directory for temporary files
```

**Obtaining outputs from Maker Round 1:**

```
fasta_merge -o Sciara_rnd${N} -d ../../*.maker.output/*_master_datastore_index.log

gff3_merge -o Sciara_rnd${N}.all.gff -d ../../*.maker.output/*_master_datastore_index.log

gff3_merge -n -g -o Sciara_rnd${N}.makermodels.gff -d ../../*.maker.output/*_master_datastore_index.log
```

**Running subsequent rounds of Maker2 with gene predictors:**

As mentioned above, SNAP was trained using the output of Maker round1; Augustus was trained with BUSCO; and GeneMark-ES was self-trained. As we ran this annotation pipeline on the two final assembly candidates, all training was specific to the assembly being annotated. Thus, the Canu assembly annotations were trained on the Canu assembly, and the Falcon assembly annotations were trained on the Falcon assembly.

##### 4.3.4.11 Maker2 Round 2

The maker_opts.ctl file for Round 2 was the same as Round 1 except for the following:

```
#-----Gene Prediction
snaphmm=/Path/to/Sciara_rnd1.hmm #SNAP HMM file
gmhmm=/Path/to/gmhmm.mod #GeneMark HMM file
augustus_species=busco_trained_Sciara #Augustus gene prediction species model
est2genome=0 #infer gene predictions directly from ESTs, 1 = yes, 0 = no
```

```
protein2genome=0 #infer predictions from protein homology, 1 = yes, 0 =
no
```

Outputs were obtained as in round 1.

To be confident the BUSCO-trained Augustus model was most appropriate, we also compared AED distributions when specifying:

```
augustus_species=fly
```

and:

```
augustus_species=aedes
```

The overall AED distributions were similar with a slight advantage to using the BUSCO-trained *Sciara* model. However, when looking at the AED distributions of only Augustus predicted genes, the *Sciara* model definitively out-performed the “fly” and “aedes” models. The BUSCO-trained *Sciara* model was used going forward.

##### 4.3.4.12 Maker2 Round 3 (keep_preds=1):

SNAP was trained one more time with the output from Maker Round 2 to create a new SNAP HMM for Maker Round 3 (Sciara_rnd2.hmm). Thus, the maker_opts.ctl file for Maker Round 3 was the same as for Maker Round 2, except:

```
#-----Gene Prediction
snaphmm=/Path/to/Sciara_rnd2.hmm #SNAP HMM file
```

In previous rounds, we had keep_preds=0. However, some predictions that are not supported by available RNA-seq and homology evidence may have recognizable protein domains. Thus, Maker can be run such that predictions are retained, and gene sets are constructed afterward with a post-hoc script that filtered based on AED and/or containment of protein domains (Campbell et al. 2014). The initial output containing everything is the “max” set. Filtering for only genes that are supported by the evidence within the Maker run (AED in 0 to 1) gives the default set. The standard set is intermediate and contains the default set plus any additional genes from the max set with recognizable protein domains. To take advantage of assessing the three different levels of gene sets from Maker (default, standard, and maximum: (Campbell et al. 2014), Maker Round 3 was re-run with:

```
#-----MAKER Behavior Options
keep_preds=1
```

Outputs were obtained as in round 1 and with these additions:

```
# InterProScan was run on all Maker Proteins
PROT=Sciara_rnd3.all.maker.proteins.fasta
interproscan.sh -dp -cpu ${SLURM_NTASKS} \
      -appl PfamA -iprlookup -goterms -f tsv -i ${PROT}

GFF=Sciara_rnd3.all.gff
TSV=${PROT}.tsv
OUT=Sciara_rnd3.max.functional_ipr.gff

# Update GFF to new GFF
ipr_update_gff ${GFF} ${TSV} > ${OUT}

## Make default build (should be same as rnd3 run)
quality_filter.pl -d ${OUT} > Sciara_rnd3.default.functional_ipr.gff

## Make standard build
quality_filter.pl -s ${OUT} > Sciara_rnd3.standard.functional_ipr.gff
```

```
## Count number of genes between both files to find out how many were
rescued
DEF=$( awk '$3=="gene"' Sciara_rnd3.default.functional_ipr.gff | wc -l
)
STD=$( awk '$3=="gene"' Sciara_rnd3.standard.functional_ipr.gff | wc -l
)
DIFF=$( echo $STD - $DEF | bc )
H="Default\tStandard\tDifference\n${DEF}\t${STD}\t${DIFF}"
echo -e ${H} > def-std-diff.txt

## Get names of transcripts in default and standard sets
awk '$3=="mRNA" {print $9}' Sciara_rnd3.default.functional_ipr.gff |
awk '{gsub(/=|;/,"\t"); print $2}' | sort | uniq > default-
transcripts.txt
awk '$3=="mRNA" {print $9}' Sciara_rnd3.standard.functional_ipr.gff |
awk '{gsub(/=|;/,"\t"); print $2}' | sort | uniq > standard-
transcripts.txt

# Get number of rescued transcripts
python -c "default=set([e.strip() for e in open('default-
transcripts.txt').readlines()]); std=set([e.strip() for e in
open('standard-transcripts.txt').readlines()]); ans =
std.difference(default); print( len(ans) )"

# Get names of rescues
python -c "default=set([e.strip() for e in open('default-
transcripts.txt').readlines()]); std=set([e.strip() for e in
open('standard-transcripts.txt').readlines()]); ans =
std.difference(default); print( '\n'.join(list(ans)) )" > rescues.txt

# Get transcripts and proteins from default and standard sets
extractFastxEntries.py -n standard-transcripts.txt -f
Sciara_rnd3.all.maker.transcripts.fasta > standard-transcripts.fasta
extractFastxEntries.py -n default-transcripts.txt -f
Sciara_rnd3.all.maker.transcripts.fasta > default-transcripts.fasta
extractFastxEntries.py -n standard-transcripts.txt -f
Sciara_rnd3.all.maker.proteins.fasta > standard-proteins.fasta
extractFastxEntries.py -n default-transcripts.txt -f
Sciara_rnd3.all.maker.proteins.fasta > default-proteins.fasta

# Get rescue transcripts
extractFastxEntries.py -n rescues.txt -f
Sciara_rnd3.all.maker.transcripts.fasta > rescued-transcripts-for-
analysis.fasta

# Get functions of rescues
grep -f rescues.txt Sciara_rnd3.all.maker.proteins.fasta.tsv  > rescue-
functions.txt
```

###### 4.3.4.13 Maker2 Round 3 with gene-filtered repeat library (keep_preds=1)

We noticed that the histone genes, odorant receptors, and other genes were part of the Repeat Modeler species-specific libraries, and were potentially under-represented in our gene annotation set after the original "Round 3". Therefore, we set out to re-run Maker Round 3 using a filtered repeat library where repeat models with matches to protein genes were excluded (unless they were transposon related).

To construct the filtered set of repeats, we used BLASTX to compare the protein database we constructed for Maker (described above) against the two Repeat Modeler libraries that we created separately on the final Canu and Falcon assemblies (-culling_limit 5 -qcov_hsp_perc 1 -evalue 1e-5). We then manually went through the results to determine whether repeats with protein hits were related to transposons/retrotransposons or not. Transposon hits were kept in the repeat libraries. This effort was aided using NCBI's online blast suite, the full nucleotide database, non-redundant protein database, and RNA refseq database. After splitting the repeat libraries into "repeats for masking" and "contains genes", we used BLAST iteratively to identify repeats in the libraries that did not have initial BLASTX hits, but that were highly similar to repeats regarded as "contains genes" (at least 50% of the query or subject was overlapped). We used BLASTX/BLASTN against the nucleotide and RNA RefSeq databases to ensure that the majority that had hits, had hits to genes, not transposons/retrotransposons. These filtered Repeat Modeler libraries were then combined with Arthropoda in Dfam/RepBase and known repeat sequences as done above, and subsequently used in Maker2 for less aggressive repeat masking.

InterProScan annotation and all outputs of this "alternative round 3" from Maker were obtained as in the section above, "Maker Round 3 keep_preds=1".

##### 4.3.4.14 Maker2 Round 3 combined standard gene sets and finalization

We evaluated the various Maker Round 3 outputs with AED distributions, BUSCO, RSEM-Eval, and TransRate as described elsewhere. The gene sets produced using the original Repeat Library had more genes, better AED distributions, more BUSCOs, and better evaluations otherwise. Therefore, we used only gene annotations generated when using the filtered repeat library only if they had no overlaps with the set of annotations generated when using the original repeat library (using BEDtools). This resulted in adding 1132 and 607 gene models to the original Canu and Falcon gene model sets, respectively. We found that of these additional genes added back to the Canu set, for example, 30 were labeled as "Core histone H2A/H2B/H3/H4" and 11 were labeled as "7tm Odorant receptor". Thus, this increases the representation of those gene families in our final set as we set out to do in the first place as described above.

The combined gene sets were then finalized by removing gene models corresponding to contaminating (bacterial) contigs to give the final set specific to *Sciara* coprophila (Bradysia coprophila). The Maker2 gene names were changed to have a consistent naming scheme (e.g. for Canu all were given the prefix: Bcop_v1_g for Bradysia coprophila, version 1, gene number). In addition to InterProScan and Gene Ontology information obtained previously, proteins were functionally annotated, where possible, by using BLASTp against the entire UniProt/SwissProt protein database. For every *Sciara* protein, we kept only the BLASTp hit with the highest bitscore. Best hits were only retained if they had a bitscore >= 50 or an expected value <= 5e-4. The GFF and Fasta files were updated with the BLASTp hit and all other functional information obtained. For further evaluation of the Maker gene sets for the Canu and Falcon assemblies, we also found the number of genes with BLASTp hits (performed as above) in the *Drosophila melanogaster* and *Anopheles gambiae* proteomes. Gene statistics (on isoforms, exons, introns, etc) were obtained from both the final Maker2 gene model sets as well as the input StringTie transcriptome assemblies using a custom script.

Commands used to find and combine non-overlapping gene models:

```
## Note: find the following at: https://github.com/JohnUrban/sciara-
project-tools
##    parseFamiliesfromGFF.py
##    fasta_name_changer.py
##  Other scripts:
##    gff3sort.pl (https://github.com/billzt/gff3sort; Zhu et al, 2017)
##    BEDtools (Quinlan and Hall, 2010)
```

```
# VARIABLES
PRE=sciara_rnd3
ORIGDIR=../rnd3
ALTDIR=../../with_filtered_repeats/outputs_keeppred1/rnd3
ORIGGFF_PRECLEAN=${ORIGDIR}/${PRE}.standard.functional_ipr.gff
ORIGGFF=cleaned-original-${PRE}.standard.functional_ipr.gff
ORIGTRANS=${ORIGDIR}/standard-transcripts.fasta
ORIGPROT=${ORIGDIR}/standard-proteins.fasta
ORIGIPS=${ORIGDIR}/sciara_rnd3.all.maker.proteins.fasta.tsv
ALTGFF_PRECLEAN=${ALTDIR}/${PRE}.standard.functional_ipr.gff
ALTGFF=cleaned-alt-${PRE}.standard.functional_ipr.gff
ALTTRANS=${ALTDIR}/standard-transcripts.fasta
ALTPROT=${ALTDIR}/standard-proteins.fasta
NONGFF=${PRE}.nonoverlapping-genes.gff
OUTGFF=${PRE}.nonoverlapping-families.gff
OUTTRANS=${PRE}.nonoverlapping-families.transcripts.fasta
OUTPROT=${PRE}.nonoverlapping-families.proteins.fasta
OUTGFFMOD=${PRE}.nonoverlapping-families-modifiedNames.gff
OUTTRANSMOD=${PRE}.nonoverlapping-families.transcripts-
modifiedNames.fasta
OUTPROTMOD=${PRE}.nonoverlapping-families.proteins-modifiedNames.fasta
COMBGFF=${PRE}.combined.standard.gff
COMBTRANS=${PRE}.combined.standard.transcripts.fasta
COMBPROT=${PRE}.combined.standard.proteins.fasta
COMBIPS=${PRE}.interproscan.results.tsv

#0. Clean up Parent attribute in Maker Standard GFF (it has references
to mRNAs that did not make the cut, which can cause issues for some
programs) - cleanParentSlotInGFF.py found at
https://github.com/JohnUrban/sciara-project-tools

cleanParentSlotInGFF.py -g ${ORIGGFF_PRECLEAN} --notfound cleaned-
original-notfound.txt > ${ORIGGFF}

cleanParentSlotInGFF.py -g ${ALTGFF_PRECLEAN} --notfound cleaned-alt-
notfound.txt  > ${ALTGFF}

#1. Obtain gene locations from alternative gene set that have no
overlaps with the original set

intersectBed -v -a <(awk '$3=="gene"' ${ALTGFF} ) -b <( awk
'$3=="gene"' ${ORIGGFF} ) > ${NONGFF}

#2. Create a text file with just the “parents” (gene names) of the non-
overlapping add-ons from alternative gene set.

awk '$3=="gene" {print $9}' ${NONGFF} | awk '{gsub(/=|;/,"\t"); print
$2}' | sort | uniq > parents.txt

#3. Take all GFF entries that correspond to the given parents from
alternative GFF

parseFamiliesfromGFF.py -i ${ALTGFF} -p parents.txt > ${OUTGFF}

#4. Obtain all non-overlapping transcript names, and extract the
transcript and protein sequences corresponding to them
```

```
awk '$3=="mRNA" {print $9}' ${OUTGFF} | awk '{gsub(/=|;/,"\t"); print $2}' | sort | uniq > transcripts.txt

extractFastxEntries.py -n transcripts.txt -f ${ALTTRANS} > ${OUTTRANS}

extractFastxEntries.py -n transcripts.txt -f ${ALTPROT} > ${OUTPROT}

#5. Modify the names of the genes/transcripts/proteins so they do not collide with names in the original gene set

parseFamiliesfromGFF.py -i ${OUTGFF} -p parents.txt --namechanger addons_ > ${OUTGFFMOD}

fasta_name_changer.py -f ${OUTTRANS} -F addons_ > ${OUTTRANSMOD}

fasta_name_changer.py -f ${OUTPROT} -F addons_ > ${OUTPROTMOD}

#6. COMBINE the name changed "Add-ons" with the original gene set and sort

cat ${ORIGGFF} ${OUTGFFMOD} > tmp.gff

gff3sort.pl --precise --chr_order natural tmp.gff > ${COMBGFF}

#7. COMBINE the name changed "Add-on" transcripts and proteins with original sets

cat ${ORIGTRANS} ${OUTTRANSMOD} > ${COMBTRANS}

cat ${ORIGPROT} ${OUTPROTMOD} > ${COMBPROT}

#8. COMBINE the InterProScan domain/functional information

grep –w -f parents.txt ${ALTDIR}/sciara_rnd3.all.maker.proteins.fasta.tsv  | awk '{print "addons_"$0}' > parent-functions-modifiedNames.txt

cat ${ORIGIPS} parent-functions-modifiedNames.txt > ${COMBIPS}
```

Commands used to finalize gene sets:

```
## Note: find the following at: https://github.com/JohnUrban/sciara-project-tools
##    grep.py : This utility was created for simple tasks one might normally perform with grep. However, we noticed that some grep commands/options had different behaviors/results when performed on Mac OS vs Linux. This utility had stable results across platforms for the tasks below.
##    extractFastxEntries.py : Utility to manipulate FASTA files
##    tableFilter.py : Utility to extract the best BLAST hit for each gene model
##    pfammer.py : Utility to add PFAM descriptions to GFF/Fasta
```

#1. Make copies of final files into new directory such that in-place name-changing will be fine.

```
mkdir -p newdir && cd newdir
cp ../*combined* .
cp ../sciara_rnd3.interproscan.results.tsv  .
```

#2. Partition Maker GFF into bacterial and nonbacterial (Sciara) given the set of contigs classified as bacterial

```
grep.py -p bacterial.txt -f sciara_rnd3.combined.standard.gff -c 1 -C 1
> bacterial.sciara_rnd3.combined.standard.gff

grep.py -p bacterial.txt -f sciara_rnd3.combined.standard.gff -c 1 -C 1
-v > nonbacterial.sciara_rnd3.combined.standard.gff
```

#3. Identify mRNA names in Maker GFF associated with bacterial contigs

```
awk '$3=="mRNA" {print $9}' bacterial.sciara_rnd3.combined.standard.gff
| awk '{gsub(/;|=/,"\t"); print $2}' > bacterial-mRNA-names.txt
```

#4. Partition InterProScan Results

```
grep.py -p bacterial-mRNA-names.txt -f
sciara_rnd3.interproscan.results.tsv -c 1 -C 1 >
bacterial.sciara_rnd3.interproscan.results.tsv

grep.py -p bacterial-mRNA-names.txt -f
sciara_rnd3.interproscan.results.tsv -c 1 -C 1 -v >
nonbacterial.sciara_rnd3.interproscan.results.tsv
```

#5. Partition Protein Fasta

```
extractFastxEntries.py -n bacterial-mRNA-names.txt -f
sciara_rnd3.combined.standard.proteins.fasta >
bacterial.sciara_rnd3.combined.standard.proteins.fasta

extractFastxEntries.py --exclude -n bacterial-mRNA-names.txt -f
sciara_rnd3.combined.standard.proteins.fasta >
nonbacterial.sciara_rnd3.combined.standard.proteins.fasta
```

#6. Partition Trascript Fasta

```
extractFastxEntries.py -n bacterial-mRNA-names.txt -f
sciara_rnd3.combined.standard.transcripts.fasta >
bacterial.sciara_rnd3.combined.standard.transcripts.fasta

extractFastxEntries.py --exclude -n bacterial-mRNA-names.txt -f
sciara_rnd3.combined.standard.transcripts.fasta >
nonbacterial.sciara_rnd3.combined.standard.transcripts.fasta
```

#7. Change the names to consistent formatting: Bcop version 1 gene number

```
#7a. Make kev:value maps between old names and new names for Sciara
models (nonbacterial)
maker_map_ids --prefix Bcop_v1_g --justify 6
nonbacterial.sciara_rnd3.combined.standard.gff >
nonbacterial.sciara_rnd3.combined.standard.map

#7b. Make kev:value maps between old names and new names for
contaminating/removed models (bacterial)
maker_map_ids --prefix Bcop-bacterial_v1_bg --justify 6
bacterial.sciara_rnd3.combined.standard.gff >
bacterial.sciara_rnd3.combined.standard.map

#7c. Rename in place using Maker-provided utilities
for PRE in bacterial nonbacterial; do
      map_gff_ids ${PRE}.sciara_rnd3.combined.standard.map
      ${PRE}.sciara_rnd3.combined.standard.gff

      map_fasta_ids ${PRE}.sciara_rnd3.combined.standard.map
      ${PRE}.sciara_rnd3.combined.standard.transcripts.fasta

      map_fasta_ids ${PRE}.sciara_rnd3.combined.standard.map
      ${PRE}.sciara_rnd3.combined.standard.proteins.fasta

      map_data_ids ${PRE}.sciara_rnd3.combined.standard.map
      ${PRE}.sciara_rnd3.interproscan.results.tsv

done

#8. Perform BLASTP of proteins against entire UniProt/SwissProt
Database(Optionally also do on contaminating/bacterial gene models);
These commands were also used to look at the Maker proteins compared to
Drosophila and Anopheles proteomes

#8a. Download uniprot_sprot FASTA and make BLAST database
wget
ftp://ftp.uniprot.org/pub/databases/uniprot/current_release/knowledgeba
se/complete/uniprot_sprot.fasta.gz

gunzip uniprot_sprot.fasta.gz

makeblastdb -in uniprot_sprot.fasta -dbtype prot -out uniprot_sprot

#8b. BLASTp – the BLAST command used is given here although we broke
the gene sets up to compute the results in parallel. (Below PRE is
either nonbacterial or bacterial corresponding to files made above)

blastp -query $Q -db $DB -num_threads $P -outfmt 6 -evalue 1e-2 -out
output.blastp

#8c. Take only the best hit for each gene model
tableFilter.py -n 1 -s 12 output.blastp > ${PRE}.filtered-
sciara_rnd3.combined.standard.proteins.all.output.blastp

#8d. Keep hits only if they meet bitscore or e-value thresholds
(bitscore >= 50 or evalue <= 5e-4)
awk '$12>=50 || $11 <= 5e-4' ${PRE}.filtered-
sciara_rnd3.combined.standard.proteins.all.output.blastp >
```

```
${PRE}.final-filtered-
sciara_rnd3.combined.standard.proteins.all.output.blastp

#9. Update GFF and FASTA with BLAST info -- finalize

for PRE in nonbacterial bacterial; do
      ## variables
      UNIPROT=uniprot_sprot.fasta
      BLASTP=${PRE}.final-filtered-
      sciara_rnd3.combined.standard.proteins.all.output.blastp
      GFF=${PRE}.sciara_rnd3.combined.standard.gff
      BASE=$( basename ${GFF} .gff )
      PROT=${BASE}.proteins.fasta
      TRANS=${BASE}.transcripts.fasta
      NEWGFF=${BASE}.putative_function.gff
      NEWPROT=${BASE}.proteins.putative_function.fasta
      NEWTRANS=${BASE}.transcripts.putative_function.fasta

      ## Annotate Maker GFF with BLASTp information
      maker_functional_gff ${UNIPROT} ${BLASTP} ${GFF} > ${NEWGFF}

      ## Annotate Protein Fasta with BLASTp information
      maker_functional_fasta ${UNIPROT} ${BLASTP} ${PROT} > ${NEWPROT}

      ## Annotate Transcript Fasta with BLASTp information
      maker_functional_fasta ${UNIPROT} ${BLASTP} ${TRANS} >
      ${NEWTRANS}

      ## Create GFF annotating genome with InterProScan Domain/GO
      Information
      iprscan2gff3 ${PRE}.sciara_rnd3.interproscan.results.tsv
      ${PRE}.sciara_rnd3.combined.standard.putative_function.gff >
      ${PRE}.sciara_rnd3.combined.standard.putative_function.visible_ip
      rscan_domains.gff

      ## Annotate Protein Fasta with InterProScan Domain/GO/Pfam info
      pfammer.py -p ${PRE}.sciara_rnd3.interproscan.results.tsv -f
      ${NEWPROT} > final.${NEWPROT}

      ## Annotate Transcript Fasta with InterProScan Domain/GO/Pfam
      info
      pfammer.py -p ${PRE}.sciara_rnd3.interproscan.results.tsv -f
      ${NEWTRANS} > final.${NEWTRANS}

      ## Annotate Maker GFF with InterProScan Domain/GO/Pfam info
      pfammer.py -p ${PRE}.sciara_rnd3.interproscan.results.tsv -g
      ${NEWGFF} > final.${NEWGFF}
done
```

Commands used to get gene statisitcs from Maker and StringTie GFFs:

```
## Note: find the following at: https://github.com/JohnUrban/sciara-
project-tools
## annotationGeneStats.py

annotationGeneStats.py –m /path/to/maker.gff
annotationGeneStats.py –s /path/to/stringtie.gff
```

###### 4.3.4.15 Maker2 gene set evaluations

Maker2 transcriptome evaluations were performed the same as for transcriptome assemblies in "4.3.2 Evaluations of transcriptome assemblies". We also compared the distributions of Annotation Edit Distances (AED) to determine the number of genes with AEDs less than or equal to given cut offs. The Maker2 proteome was also analyzed with BUSCO. We also looked at the Pfam domains, GO terms, and BLASTp results (to UniProt-SwissProt) from Maker2 analyses above in the Maker2 proteomes as well as compared them to known proteomes for *Drosophila melanogaster* and *Anopheles gambiae*.

##### 4.3.5 Dosage compensation analysis

Transcripts Per Million (TPM) and estimated read counts per transcript were quantified for each RNA-seq sample using Salmon (Patro et al. 2017).

```
salmon quant -i ${SIDX} -l A \
        -1 ${R1} \
        -2 ${R2} \
        -p 1 --validateMappings -o quants/${PRE}_quant
```

Reproducibility amongst replicates was determined by clustering the samples on log10(TPM+1), which showed the replicates of each stage and sex clustering together. Differential expression analyses were performed with EdgeR (Robinson et al. 2010).

```
x <- read.delim("quants/all-salmon-counts-for-
EdgeR.txt",row.names="Name")

group <- factor(c(rep("FE",3),rep("FL",2),rep("FP",2), rep("FA",2),
rep("ME",3),rep("ML",2),rep("MP",2),rep("MA",2)),
                levels = c("FE","FL","FP","FA","ME","ML","MP","MA"))
y <- DGEList(counts=x,group=group)
keep <- filterByExpr(y, min.count = 1, min.total.count=10,
min.prop=0.25)#; summary(keep)
y <- y[keep, , keep.lib.sizes=FALSE]
y <- calcNormFactors(y);
design <- model.matrix(~ 0 + group)
colnames(design) <- levels(group)
y <- estimateDisp(y, design, robust=TRUE)
fit <- glmQLFit(y,design, robust=TRUE)

conE <- makeContrasts(ME - FE, levels=design)
qlfE <- glmQLFTest(fit, contrast=conE)
conL <- makeContrasts(ML - FL, levels=design)
qlfL <- glmQLFTest(fit, contrast=conL)
conP <- makeContrasts(MP - FP, levels=design)
qlfP <- glmQLFTest(fit, contrast=conP)
conA <- makeContrasts(MA - FA, levels=design)
qlfA <- glmQLFTest(fit, contrast=conA)
```

##### 4.3.6 Characterization of the Lambda Phage Insert Containing ScRTE

We characterized the repeat content of the 13.8 kb lambda phage insert sequence (called Escribá insert here) that contains two copies of the non-LTR retrotransposon named ScRTE (Escribá et al. 2011). The Escribá insert sequence was aligned to itself with BLAST to find direct and inverted repeats within it (see Figure 5E). It was also BLAST against the Canu scaffolds to determine the copy number over each base. The Escribá insert sequence was aligned to the entire NCBI nucleotide database and nonredundant protein databases to annotate it. Our Illumina 100 bp paired-end DNA dataset was aligned with Bowtie2 to indexes of the Canu scaffolds that were masked with the Escribá insert sequence followed by adding one copy of the Escribá insert sequence back in. RepeatMasker was used to classify repeats along the Escribá insert using the Arthropod repeat database. BLAST was used with our de novo repeat families from RepeatModeler to determine what percent of the top ten most highly abundant repeat sequences in the genome aligned to the Escribá insert by at least 75% across their lengths.

### 4.4 DNA modifications

Both PacBio SMRT and Oxford Nanopore single molecule sequencing technologies come with raw signal information that can be used to identify base modifications (Flusberg et al. 2010; Clark et al. 2012; Loman et al. 2015; Simpson et al. 2017; Rand et al. 2016; Suzuki et al. 2016) and even secondary structure formation (Sawaya et al. 2015; Urban et al. 2015). Since we had high coverage from PacBio, we used that dataset for site-specific modification calls in the assembly for 5mC, 4mC, and 6mA. We subsequently used those sites to learn sequence motifs associated with each modification. Our 2D read strand coverage from MinION datasets was too low to reliably perform methylation calls at specific sites. Thus, to make use of the low coverage nanopore data, we used Nanopolish (Simpson et al. 2017) to learn kmer models specific to the *Sciara* dataset and determined kmers with observed shifted signal distributions compared to the expected models from Oxford Nanopore. Such a difference between models for a given kmer can arise from the effects of modifications within a subset of observed kmers. The set of kmers determined to have different means than the expected model were used to find sub-motifs within them to compare to the set of motifs obtained by PacBio. For both PacBio and MinION analyses, motifs were analyzed only for data from contigs in the Primary assembly labeled as Arthropoda in the blobtools analysis.

#### 4.4.1 PacBio Analysis

##### 4.4.1.1 PacBio Modification Prediction

PacBio reads were aligned in the presence of the entire unfiltered assemblies to avoid forcing reads to align incorrectly. Pbalign 0.2.0.141024 (https://github.com/PacificBiosciences/pbalign) was used with BLASR v2 to map reads in bax.h5 files, outputting cmp.h5 files for each. Pbh5tools v0.8.0 (https://github.com/PacificBiosciences/pbh5tools) was used to merge and sort the mapped reads in cmp.h5 files, outputting a single cmp.h5 file that retained the raw signal information. "ipdSummary" from kineticsTools v0.6.0 (https://github.com/PacificBiosciences/kineticsTools) was used to predict base modifications across the Canu genome assembly. At the ipdSummary step, each contig was processed separately in parallel. The output from ipdSummary includes a CSV with kinetics statistics for each position and a GFF with DNA modification predictions (both deposited on NCBI for others to use). The CSV output for each contig was also used with AgIn (Suzuki et al. 2016; https://github.com/hacone/AgIn) to look at CpG methylation. AgIn makes the assumptions that CpG methylation status is the same on both strands and that neighboring CpGs are likely to have the same methylation status. These assumptions allow it to increase the effective coverage used to detect CpG methylation status by aggregating data from both strands and multiple neighboring CpG dinucleotides to determine methylation statuses of blocks of CpGs. Note methylation sites do tend to cluster in our dataset (see spacing analysis), which supports these assumptions.

```
## Align
pbalign $BAX $REF $OUTPRE.cmp.h5 --forQuiver --tmpDir $TMPDIR --nproc
$THREADS --metrics
DeletionQV,IPD,InsertionQV,PulseWidth,QualityValue,MergeQV,Substitution
QV,DeletionTag

## Merge alignments into single cmp.h5 file
cmph5tools.py merge --outFile ${MERGEDCMP} ${CMPDIR}/*.cmp.h5

## Sort merged alignments
cmph5tools.py sort --deep ${IN_CMP} --tmpDir ${TMP_SORT}
```

```
## Predict base modifications on each contig (command applied
separately for each contig)
ipdSummary ${MERGED_SORTED_CMPH5} -v -v -v --reference ${CANU_ASM} --
identify m6A,m4C,m5C_TET --methylFraction --gff basemods.gff --csv
kinetics.csv --pvalue 0.01 --minCoverage 3 --methylMinCov 10 --
identifyMinCov 5 -j ${THREADS} --maxAlignments 1000000 --ms_csv
multisite.csv --bigwig ipd.bigWig --refContigs $contigName

## A minority of contigs have a window or windows that causes the
algorithm to not finish the classification/identification step. These
windows were found through iteration and processed separately to skip
the classification step in the following way:

# Windows that have no issues are processed as usual with the
identification step
ipdSummary ${INPUT} -v -v -v --identify m6A,m4C,m5C_TET --
methylFraction --gff basemods-identified.gff --csv kinetics-
identified.csv --bigwig ipd-identified.bigWig --reference ${REF} --
pvalue 0.01 --minCoverage 3 --methylMinCov 10 --identifyMinCov 5 -j
${THREADS} --maxAlignments 1000000 --referenceWindows ${REFWINDOWS}

# Windows that with issues that cause the algorithm to freeze are
processed without identification step
ipdSummary ${INPUT} -v -v -v --gff basemods-unidentified.gff --csv
kinetics-unidentified.csv --bigwig ipd-unidentified.bigWig --reference
${REF} --pvalue 0.01 --minCoverage 3 --methylMinCov 10 --identifyMinCov
5 -j ${THREADS} --maxAlignments 1000000 --referenceWindows
${PROBLEM_REFWINDOWS}

# AgIn was also used with author-recommended settings for P5C3
chemistry on the kinetics.csv files for each contig to predict
methylated/unmethylated blocks of CpGs
/path/to/AgIn/target/dist/bin/launch -i kinetics.csv -f $REF -o
${OUT_PREFIX}-b P5C3_HdrR -g -0.88 -l 40 -c predict
```

###### 4.4.1.2 DNA sequence motifs found near sites of predicted DNA modifications

Sequence motifs associated with PacBio modifications were searched for in two ways: an enriched kmer approach as well as using MEME (Bailey and Elkan 1994). For all analyses on modifications, we used only modifications from primary contigs labeled as Arthopoda in the Blobtools analysis to ensure that we were working with *Sciara* sequences.

###### 4.4.1.3 Kmers enriched for DNA modifications

For each modification type, we looked for the dimer and trimer that were most enriched in predicted modifications with respect to their frequency in the genome sequence. We also generated motifs from all 7-mers that were enriched with predicted modifications for more sequence context. Kmer enrichment was determined by comparing observed and expected kmer frequencies as described below:

```
Let:
K = kmer size chosen as 2, 3, 4, 5, 6, or 7

B = base of interest that can be modified; either A or C

J = position where base B occurs in kmer (i.e. where a modification may
occur on B)
```

```
S = set of all kmers size K that have base B in position J

F_background = the expected background frequency of each kmer in S (for
a given set S) obtained by counting the number of occurrences of each
kmer in the genome sequence from both strands (as either strand can be
modified).

P_background = the expected background proportions of each kmer in
F_background obtained from dividing the frequency of each kmer in
F_background by the sum of the frequencies of all kmers in
F_background.

F_modifications = the observed modification frequency of each kmer in S
obtained by counting the number of times it was classified as modified
in the genome sequence on either strand.

P_modifications = the observed modification proportions of each kmer in
F_modifications obtained by dividing each the frequency of each kmer in
F_modifications by the sum of all kmer frequencies in F_modifications.
```

For a given set, **S**, that contains the subset of all kmers of size **K** with a given position, **J**, where the target base **B** occurs (e.g. C for mC or A for 6mA ), we counted the number of times each kmer in S appeared in the genome on both strands (**F_background**), and defined the proportion of F_background that each kmer count made up as described above (**P_background**). Then for each kmer in S, we counted the subset of occurences in the genome on both strands where base B at position J of the kmer was determined to be modified (**F_modifications**), and defined the proportion of F_modifications that each kmer count made up as described above (**P_modifications**). We tested whether the observed proportions of the kmers with modifications to base B at position J (P_modifications) differed from the expected proportions (P_background). If modifications occurred with kmers at random, then the proportions of kmers in the set of kmers that are modified should be the same as the background proportions of those kmers. If DNA modifications occur on some kmers more than others, then the proportions of each kmer in the modified set will differ from background, and it would be possible to determine which kmers are over- and under-represented in the modified set. We used a Chi-square goodness of fit test and used the Chi-square standardized residuals that are interpreted similar to z-scores to determine which kmers occurred at much higher frequencies in F_modifications than expected given the background frequencies from F_background:

```
Standardized Residual for a kmer x
      = ((Observed_x-Expected_x)/square_root(Expected_x))
```

For each modification type, we determined the most prevalent dimers and trimers. Specifically, we focused on the set of dimers with the target base B at position J=1 and the set of trimers with the target base B at position J=2. For each modification type, we also filtered the sets of 7-mers with the target base B at position J=3 for enriched kmers using P-value and residual value cutoffs ($p < 0.000001$, residual > 50). The enriched 7-mers, all with base B at position J=3, were used with WebLogo (Crooks et al. 2004) to construct a position weight matrix (PWM) motif using the frequencies at which the kmers occurred (weighted) as well as with uniform weighting, both yielding nearly identical results. These logos show what bases tend to be nearest neighbors to base B, and the results reflect the enriched dimers and trimers we found with slightly more context in each direction.

###### 4.4.1.4 MEME motifs in the highest scoring sites

The 9 bp centered around the modified bases were extracted for the top 500 scoring modifications for each modification type. These were used with MEME 5.0.2 (Bailey and Elkan 1994) to identify motifs using a second order Markov model trained on the primary arthropod scaffolds.

```
fasta-get-markov -m 2 -dna ${ASM} > bg.txt

for M in m6A m5C m4C; do
      SEQS=${M}.top500.primary-AND-arthropod-labeled.9mers.fasta

      meme $SEQS -norand -dna -oc ${M}_w2-5_bgcanu -time 18000 -maxsize
      60000 -mod anr -nmotifs 10 -minw 2 -maxw 5 -minsites 2 -maxsites
      600 -wnsites 0 -bfile bg.txt
done
```

###### 4.4.1.5 Enrichment/Depletion of DNA modifications in various genomic regions

We searched for relationships between modification sites and the following features:

1. Repeats as defined by the regions of the genome that were masked as repeats for annotation purposes.
2. Gene regions defined by our Maker2 protein-coding gene annotations. We looked at all gene locations as one set as well as partitioned the genes based on expression levels in our male embryo RNA-seq as quantified by Salmon (Patro et al. 2017):
    a. Not expressed (0 TPM)
    b. Lowly expressed – defined as the 50% of genes with lowest non-zero expression levels (bottom 50% of TPM values > 0).
    c. Highly expressed – defined as the 50% of genes with the highest non-zero expression levels (top 50% of TPM values > 0).
3. Exons as defined by our Maker2 annotations.
4. Introns as defined as the complement intervals with respect to exons inside gene regions defined by our Maker2 annotation.
5. Promoter regions as defined as the 500 bp upstream of genes on the positive strand or 500 bp downstream of genes on the negative strand for gene intervals defined in our Maker2 annotation. We looked at all promoter regions together as well as partitioned into expression levels as with gene bodies above.

For all features, BEDtools merge was used to ensure any interval was represented only once.

###### 4.4.1.6 Binomial tests for higher or lower modification rates in genomic regions

We determined if the DNA modification rate of a given base B was enriched in various genomic features of interest using a binomial model. Specifically, we tested whether there was a higher frequency of modifications to the B bases in the regions of interest than expected by chance given the background rate in the genome. These numbers are reflected in Supplemental Tables S20A, C, D, and E. Regions were tested for enrichment of modifications to a given base using a binomial model the following way:

```
Let:
B = base that may or may not be modified (A or C).

N_B_genome = Number of times base B occurs in the genome on both
strands.

N_mod_genome = Number of sites in genome on both strands where base B
was determined to be modified (given desired cut-offs for considering
modifications).

P_genome = expected probability of base B being modified in the genome
             = N_mod_genome/N_B_genome

R = regions of interest – i.e. intervals from the genome sequence such
as gene locations.

N_B_R = Number of times base B occurs in R, the regions of interest, on
both strands.

N_mod_R = Number of sites in R on both strands where base B was
determined to be modified.

P_R = observed probability of base B being modified in R
        = N_mod_R/N_B_R

Binomial P-value that modifications are enriched in R with respect to
the background genome probability:
      Let:
      n = N_B_R
      k = N_mod_R
      p = P_genome
```

$$P\,Value = \Pr(X \geq k) = \sum_{k}^{n} \binom{n}{k} p^k (1-p)^{n-k}$$

```
Binomial P-value that modifications are depleted in R with respect to
the background genome probability:
      Let:
      n = N_B_R
      k = N_mod_R
      p = P_genome
```

$$P\,Value = \Pr(X \leq k) = \sum_{0}^{k} \binom{n}{k} p^k (1-p)^{n-k}$$

For a given modification to base B, we counted the number of base B in the genome on both strands (**N_B_genome**) and the number of times base B was modified on either strand (**N_mod_genome**) to get the expected background probability that base B is modified (**P_genome**). We then followed the same procedure for the subset of B bases in the given regions of interest to get the number of B bases and observed modification count therein (**N_B_R and N_mod_R**). We calculated the binomial p-value as above given these parameters to test if there were significantly more modifications to base B in the regions of interest than expected by chance given the background rate of modifications to base B across the genome.

Binomial results were supported by permutation tests the following way:

1. P_R (proportion of B bases modified in R) was calculated as described above

2. The regions of interest, R, were shuffled throughout the genome with BEDtools 1000 times, and the proportion of B bases modified in the shuffled regions was calculated the same way for each (P_R_shuff).
3. P-values testing if modifications were enriched in the regions of interest (R) were approximated to be the proportion of the 1000 P_R_shuff values that were as high or higher than the observed P_R value. Alternatively, p-values testing if modifications were depleted in R were approximated to be the proportion of the 1000 P_R_shuff values that were as small or smaller than the observed P_R value.

##### 4.4.1.7 Binomial tests for enrichment/depletion of DNA modifications in genomic regions

We also used binomial statistics to look at the problem a different way. We determined if the DNA modifications were enriched (or depleted) in various genomic features of interest by testing whether there was a higher frequency of modifications to the B bases in the regions of interest than expected by chance given the prevalence of the regions of interest in the genome. These numbers are reflected in Supplemental Tables S20B and S20F as well as Supplemental Figure S15B. The tables and figures report the expected vs observed percents to simplify the interpretation.

Regions were tested for enrichment of modifications to a given base using a binomial model the following way:

```
Let:
B = target base

G = total number of bases in the genome or assembly.

R = total number of bases in the regions of interest.

N_G = total number of bases in the genome or assembly.

N_R = total number of bases in the regions of interest.

p = proportion of bases in the genome that are in regions of interest
    = N_R/N_G

n = number of modifications in the genome

k = number of modifications in the regions of interest

Binomial P-value that modifications in G are enriched in R given n
total modifications in G, k modifications in R, and the probability, p,
of randomly being in R:
```

$$P\,Value = \Pr(X \geq k) = \sum_{k}^{n} \binom{n}{k} p^k (1-p)^{n-k}$$

```
Binomial P-value that modifications in G are depleted in R given n
total modifications in G, k modifications in R, and the probability, p,
of randomly being in R:
```

$$P\,Value = \Pr(X \leq k) = \sum_{0}^{k} \binom{n}{k} p^k (1-p)^{n-k}$$

##### 4.4.1.8 Spacing between DNA modifications

BEDtools was used to obtain spacing distances between modified bases in subsets of the GFF file output by Kinetics Tools. For strand-agnostic spacing, “bedtools spacing” was used directly on the input. For strand-specific spacing, “bedtools spacing” was used on subsets of the input separated by which strand the modification was on, positive or negative. Then the two sets of strand-specific distances were combined. “shuffleBed” and “nucBed” were used to find N shuffled positions that occurred over the target base (e.g. A for 6mA or C for 4mC) in strand-agnostic and strand-specific manners, where N is the number of DNA modifications in the test set being compared. Commands used are below.

**For strand-agnostic spacing (observed and expected/shuffled)**:

```
# Get spacing between given set of DNA modifications
bedtools spacing -i <( sortBed -i ${GFF} ) | awk '$10!="."
{a[$10]+=1}END{for (e in a) print e"\t"a[e]}' | sort -k1,1n > ${NAME}

# Get spacing between the same number of shuffled “modifications” over
the same base
## N = number of GFF entries for modified base
N=$( grep -c -v "^#" ${GFF} )

## Identify nucBed column to look at
  if [ $TBASE == A ]; then COL=12; elif [ $TBASE == C ]; then COL=13;
elif [ $TBASE == G ]; then COL=14; elif [ $TBASE == T ]; then COL=15; fi

## Shuffle 4*N entries, take only N shuffled features that correspond to
the target base
cat ${GFF} ${GFF} ${GFF} ${GFF} | bedtools shuffle -chrom -noOverlapping
-g ${G} -i - | sortBed -i - | nucBed -fi ${REF} -bed - -s | awk -v
"COL=$COL" '$1!~/^#/ && $COL>0 {OFS="\t"; print
$1,$2,$3,$4,$5,$6,$7,$8,$9}' | shuf -n ${N} | sortBed -i - >
${RANDOM_NAME}

## Get random spacing
bedtools spacing -i ${RANDOM_NAME} | awk '$10!="." {a[$10]+=1}END{for (e
in a) print e"\t"a[e]}' | sort -k1,1n > ${RANDOM_SPACING_NAME}
```

**For strand-specific spacing (observed and expected/shuffled)**:

```
# Get spacing between given set of DNA modifications
awk '$7=="+"' ${GFF} | sortBed -i - > ${POS_GFF}
awk '$7=="-"' ${GFF} | sortBed -i - > ${NEG_GFF}

## Positive strand
bedtools spacing -i ${POS_GFF} | awk '$10!="." {a[$10]+=1}END{for (e in
a) print e"\t"a[e]}' | sort -k1,1n > ${POS_NAME}

## Negative strand
bedtools spacing -i ${NEG_GFF} | awk '$10!="." {a[$10]+=1}END{for (e in
a) print e"\t"a[e]}' | sort -k1,1n > ${NEG_NAME}

## Combine
cat ${POS_NAME} ${NEG_NAME} | awk '{a[$1]+=$2}END{for (e in a) print
e"\t"a[e]}' | sort -k1,1n > ${NAME}
```

```
# Get spacing between the same number of shuffled "modifications" over
the same base
## N = number of GFF entries for modified base
NPOS=$( grep -c -v "^#" ${POS_GFF})
NNEG=$( grep -c -v "^#" ${NEG_GFF})

## Identify nucBed column to look at
  if [ $TBASE == A ]; then COL=12; elif [ $TBASE == C ]; then COL=13;
elif [ $TBASE == G ]; then COL=14; elif [ $TBASE == T ]; then COL=15; fi

## POS STRAND
## ## Shuffle 4*N entries, take only N shuffled features that correspond
to the target base
cat ${POS_GFF} ${POS_GFF} ${POS_GFF} ${POS_GFF} | bedtools shuffle -chrom
-noOverlapping -g ${G} -i - | sortBed -i - | nucBed -fi ${REF} -bed - -s
| awk -v "COL=$COL" '$1!~/^#/ && $COL>0 {OFS="\t"; print
$1,$2,$3,$4,$5,$6,$7,$8,$9}' | shuf -n ${N} | sortBed -i - >
${POS_RANDOM_NAME}

## ## Get random spacing
bedtools spacing -i ${POS_RANDOM_NAME} | awk '$10!="." {a[$10]+=1}END{for
(e in a) print e"\t"a[e]}' | sort -k1,1n > ${POS_RANDOM_SPACING_NAME}

## NEG STRAND
## ## Shuffle 4*N entries, take only N shuffled features that correspond
to the target base
cat ${NEG_GFF} ${NEG_GFF} ${NEG_GFF} ${NEG_GFF} | bedtools shuffle -chrom
-noOverlapping -g ${G} -i - | sortBed -i - | nucBed -fi ${REF} -bed - -s
| awk -v "COL=$COL" '$1!~/^#/ && $COL>0 {OFS="\t"; print
$1,$2,$3,$4,$5,$6,$7,$8,$9}' | shuf -n ${N} | sortBed -i - >
${NEG_RANDOM_NAME}

## ## Get random spacing
bedtools spacing -i ${NEG_RANDOM_NAME} | awk '$10!="." {a[$10]+=1}END{for
(e in a) print e"\t"a[e]}' | sort -k1,1n > ${NEG_RANDOM_SPACING_NAME}

## Combine random spacing from both strands
cat ${POS_RANDOM_SPACING_NAME} ${NEG_RANDOM_SPACING_NAME} | awk
'{a[$1]+=$2}END{for (e in a) print e"\t"a[e]}' | sort -k1,1n >
${RANDOM_SPACING_NAME}
```

We observed approximately 10 bp periodicities in the distribution of distances between adjacent DNA modifications. To formally identify the periodicity, using the Fast Fourier Transform (FFT) provided in Python's Numpy package, we ran a discrete fourier transform (DFT) analysis on the distributions of distances between DNA modifications. Prior to computing the FFT, the data was first mean-subtracted and tapered with a Hanning window. The spectral power over 100 discrete frequency bins was then computed by squaring the modulus of the transformed data. For each set of modifications, we recorded the bp spacing periodicity (< 100 bp) with the maximum power. Spectral power was fairly monochromatic at the 10-bp periodicity across all subsets of the DNA modification datasets (all modified A, 6mA, all modified C, 4mC, 5mC) and was robust to various filtering strategies from using all sites to requiring 10X, 15X, 20X, or 25X coverage per strand and modification scores greater than 20. The python code for automating this can be found at: https://github.com/JohnUrban/fftDnaMods.

##### 4.4.2 Orthogonal support for PacBio modification results using the MinION data

For the MinION analysis, we only used the subset of our MinION data generated using the MkI with the SQK-MAP006 kit and R7.3 70 bps 6mer pore model (libraries 14, 15, 16, 20, 21). We used Nanopolish (Simpson et al. 2017) to learn updated pore models given the data. Briefly, in MinION sequencing, the raw ionic current data is segmented at sites where the picoamp (pA) level shifts thus compressing the data into a sequence of "events", each of which includes a mean, standard deviation, and duration. Each event roughly corresponds to a stretch of 6 bases called a 6-mer. Oxford Nanopore has expected models of the mean pA level and standard deviation for each 6-mer that are used for base-calling. With Nanopolish, the basecalled MinION reads are first aligned to a reference, subsequently allowing the ionic current events to be aligned to 6-mers in the reference. The collection of events aligned over any site of a given 6-mer in the reference is used to update the models and re-align the events in an iterative process (Simpson et al. 2017). The final updated model and event distributions for a given 6-mer can then be compared to the original model, and differences may imply the presence of DNA modifications in a subset of the data. Conversely, there should not be differences (or only small differences) when performing this procedure on PCR amplified genomic DNA that would only contain canonical bases. We were interested in (i) testing whether short 2-5 bp motifs consistently arose in the 6-mers that differed from the ONT models, and (ii) looking at the event distributions from 6-mers suggested to be modified often in the PacBio analysis.

We compared the event distributions for each of the 4096 6mers in our *Sciara* dataset to the expected ONT kmer models extracted from the basecalled fast5 files using our Fast5Tools scripts, and to a MinION dataset generated from whole genome PCR on *E. coli* genomic DNA using the same kit and pore model (BioProject PRJEB13021; Run ERR1309547; www.ebi.ac.uk/ena; Simpson et al. 2017). As PCR was used for the latter dataset, the signal distributions for each kmer should closely match the expected ONT kmer model. BWA (Li and Durbin 2009) was used to align MinION reads. For our *Sciara* data, although reads were initially aligned to the unfiltered assembly to avoid forcing incorrect alignments, only reads aligned to primary contigs annotated as Arthropoda were used for further analysis. Nanopolish (Simpson et al. 2017) was used to learn updated kmer models from the native *Sciara* and *E. coli* PCR MinION datasets. For mapping *E. coli* reads, the K12 MG1655 reference was used (Accession: NC_000913.3). Example commands for *Sciara* data:

```
G=assembly.fasta
R=reads.fasta
B=reads.bam
samtools faidx $G
bwa index $G
for lib in lib14  lib15  lib16  lib20  lib21; do
 nanopolish extract $lib/pass/ -o $lib.pass.fa -t 2d
 nanopolish extract $lib/fail/ -o $lib.fail.fa -t 2d
done

cat lib*pass.fa > $R   # Or cat lib*pass.fa lib*fail.fa > $R

bwa mem -M -x ont2d $G $R -t $T | samtools sort -T $lib.pass --threads
$T | samtools view -F 4 -q ${Q} -bSh - > $B

samtools index $B
```

```
nanopolish methyltrain --reads $R --bam $B --genome $G -t $T --models-
fofn=ont.alphabet_nucleotide.R7.fofn --train-kmers all --rounds=5 --
progress
```

MEME 5.0.2 (Bailey and Elkan 1994) was used with a second order Markov model trained on the primary arthropod scaffolds to find 3 bp, 6 bp, and 2-5 bp motifs from the sets of 6-mers that that differed from the expected ONT model mean by more than 0.7-1 pA for comparison to the MEME motifs learned from the PacBio results. When we tested this procedure on a native *E. coli* dataset (BioProject PRJEB7385; Run ERR1147227; file MAP006-1.tar; www.ebi.ac.uk/ena), 32 of 34 6-mers that had differences in the emission means >1 pA contained GATC, the known motif for adenine methylation. When we learned new models given the *Sciara* data using only the passing quality 2D reads, there were 196 6-mers in the template strand model and 108-111 6-mers in each of the two complement models that had means that differed by more than 1 pA from the ONT model. If the motifs from PacBio or from the MinION results were random, we did not expect to find shared motifs. If DNA modifications were present non-randomly in both datasets, we expected to find shared motifs.

```
## Trimer motifs
SEQS= # all 6mers with absolute difference > 0.7
      # or all 6mers without CG in them with absolute difference > 0.7
meme $SEQS -bfile canu01.m2.txt -dna -maxsize 60000 -mod zoops -nmotifs
10 -minw 3 -maxw 3 -minsites 96 -maxsites 400

## 6 bp motifs
SEQS= # all 6mers with absolute difference > 0.8; 6mer sequences were
padded with NNN on each side to allow flexibility in 6 bp windows

meme $SEQS -bfile canu01.m2.txt -dna -maxsize 60000 -mod anr -nmotifs 10
-minw 6 -maxw 6 -minsites 30 -maxsites 600

## 2-5 bp motifs
SEQS= # all 6mers with absolute difference > 1
meme $SEQS -bfile canu01.m2.txt -dna -maxsize 60000 -mod anr -nmotifs 20
-minw 2 -maxw 5 -minsites 2 -maxsites 600
```

We also visualized the final event distributions from Nanopolish to interrogate the motifs learned from the PacBio medication results. We selected 6mers defined by the PWM WebLogo motifs from enriched 7-mers in the PacBio analysis. We also selected 6-mers that were not enriched to visualize for comparison. We compared the event distributions from our *Sciara* data to the expected Oxford Nanopore model and MinION data generated from PCR of E. coli genomic DNA that used the same reagents, kit, MinION, and pore model (Simpson et al. 2017): BioProject PRJEB13021; Run ERR1309547; www.ebi.ac.uk/ena. Kmers shown to be enriched with modifications in the PacBio analysis of *Sciara* were expected to have shifts in the distribution whereas other kmers from *Sciara* and all kmers from the PCR E. coli dataset were expected to have the same distribution as the expected model.

##### 4.4.3 Calculating observed:expected ratios of dimers and trimers

Observed:expected (O:E) ratios were computed on each Maker2-defined transcript that had a minimum length of 500 bp and at most 5% N bases. Genome-wide, O:E ratios were computed on all 500 bp bins with <5% N bases as defined by BEDtools.

The observed:expected ratio for dimers and trimers were computed the following way(s). The observed probability of base b in the genome was calculated simply as the number of b bases divided by the sum of all bases, B:

$$O(b) = b/B$$

The expected probability of a dimer with bases b1 and b2 was calculated as:

$$\begin{aligned} E(b1b2) &= O(b1)*O(b2) \\ &= (b1/B) * (b2/B) \\ &= (b1*b2)/(B^2) \end{aligned}$$

The observed probability of a dimer with bases ordered as b1 and b2 was calculated as the number of the specific dimer, b1b2, divided by the total number of dimers, D, which works out to be the total number of bases minus 1 (B-1):

$$\begin{aligned} O(b1b2) &= (b1b2)/D \\ &= (b1b2)/(B-1) \end{aligned}$$

Thus, the observed:expected ratio of a dimer was:

$$\begin{aligned} OE(b1b2) &= O(b1b2)/E(b1b2) \\ &= ((b1b2)/(B-1)) / ((b1*b2)/(B^2)) \\ &= ((B^2)/(B-1)) * (b1b2)/(b1*b2) \end{aligned}$$

The expected probabilities for trimers, b1b2b3, were calculated two ways, but the resulting distributions of observed:expected ratios were virtually the same. The first way assumes complete independence of the bases in the trimer:

$$\begin{aligned} E_1(b1b2b3) &= O(b1)*O(b2)*O(b3) \\ &= (b1/B) * (b2/B) * (b3/B) \\ &= (b1*b2*b3)/(B^3) \end{aligned}$$

The second way models in the observed dimer probabilities to determine the expected probability of a trimer, b1b2b3, given the observed probabilities of the b1b2 dimer and b3 monomer:

$$\begin{aligned} E_2(b1b2b3) &= O(b1b2)*O(b3) \\ &= (b1b2/(B-1)) * (b3/B) \\ &= (b1b2*b3)/((B-1)*B) \end{aligned}$$

The observed probability of a trimer with bases ordered as b1, b2, and b3 was calculated as the number of the specific trimer, b1b2b3, divided by the total number of trimers, T, which works out to be the total number of bases minus 2 (B-2):

$$\begin{aligned} O(b1b2b3) &= (b1b2b3)/T \\ &= (b1b2b3)/(B-2) \end{aligned}$$

Thus, the observed:expected ratios for trimers were calculated as:

$$OE_1(b1b2b3) = ((b1b2b3)/(B-2)) / ((b1*b2*b3)/(B^3)) = (B^3/(B-2)) / (b1b2b3/(b1*b2*b3))$$

Or:

$$OE_2(b1b2b3) = ((b1b2b3)/(B-2)) / ((b1b2*b3)/((B-1)*B)) = ((B*(B-1))/(B-2)) / (b1b2b3/(b1b2*b3))$$

#### 4.5 Candidate L sequences

L chromosomes are expected to be of relatively low abundance in the male embryo datasets used to assembly the genome. Therefore, if L sequences are present at all we expected them to (i) be assembled contigs in the primary or associated sequences (not bacterial) with extremely low coverage compared to the genomic average, or (ii) not be assembled at all, but potentially be present in reads that do not map to the assembly. Approach #1 was very similar to identifying X candidates, although here we required contigs to have 0-3X PacBio coverage across 80% or more of their lengths. Approach #2 involved assembling short and long reads that did not map to the assembly. Long reads were mapped with minimap2 (-x map-ont, -x pacbio) to identify reads that did not map whereas short reads were mapped with bowtie2 to identify pairs where neither mate mapped. Long reads were assembled using Canu with settings for very low coverage. Short reads were assembled using Platanus as described above. The input long and short reads were mapped back to these mini-assemblies with minimap2 (-x map-ont, -x sr) for coverage information to use with BlobTools. BLAST was used against the entire NT database to identify taxonomy information, and BlobTools was used to summarize the taxonomy of each contig. Repeats were identified with RepeatMasker as above. The contigs were used with BLASTx against a database of proteins from the Bcop_v1 Maker2 set described above (-culling_limit 1) to identify the best protein matches over any given locus. Highlighted proteins either aligned across 90% of their lengths or had at least 3 consecutive significant hits along the protein and contig. BLASTx against the UniProt-SwissProt database gave similar results.

#### Section 5: Software versions used

**ABruijn v0.3b**
https://github.com/fenderglass/ABruijn
Git commit: 250761bac5589d83eeb1e00e19ba7fe78cbc1738

**ABYSS v1.9.0**
https://github.com/bcgsc/abyss.git
Git commit: 21106a4d0ad4b550da6f5b77b00620e530ca5037

**AgIn**
git commit 22e347cbbc34280ee085acdf41761f401f864609
https://github.com/hacone/AgIn.git

**ALE**
(C) 2010 Scott Clark
https://github.com/JohnUrban/sciara-project-tools/tree/master/alenano

**Augustus v3.2.2 (used with Maker, BUSCO evaluations, and with BUSCO for training)**
http://bioinf.uni-greifswald.de/augustus/binaries/augustus-3.2.2.tar.gz

**BAMtools v 2.4.1 (used with Augustus and BUSCO)**
git://github.com/pezmaster31/bamtools.git

**BEDtools v2.26.0**
https://github.com/arq5x/bedtools2.git

**BLAST v 2.2.30+ (for evaluations and used with Maker)**
ftp://ftp.ncbi.nlm.nih.gov/blast/executables/blast+/2.2.30/

**BLASR v2 (git commit 8cc8621, 2015-10-22)**
https://github.com/PacificBiosciences/blasr

**BlobTools v0.9.17**
https://github.com/DRL/blobtools.git
383552447b2015620917aab8120699ae2d9d44be

**Bowtie2 v2.2.9**
https://sourceforge.net/projects/bowtie-bio/files/bowtie2/2.2.9/bowtie2-2.2.9-linux-x86_64.zip

**BUSCO**
v3.0.1 https://gitlab.com/ezlab/busco.git
ODB9: http://busco.ezlab.org/datasets/diptera_odb9.tar.gz

v.1.22 http://busco.ezlab.org/files/BUSCO_v1.22.tar.gz
http://busco.ezlab.org/files/arthropoda_buscos.tar.gz

**BWA v 0.7.14-r1136**
https://github.com/lh3/bwa.git
ee730f3832314fb70d369526275f2aadab064470

**Canu v1.0, 1.1, 1.2 and several commits after v1.3 released**
https://github.com/marbl/canu.git

4d07faca1d54852966e8ca9f4bbd4bf607b521d3

**Detonate 1.11 (used RSEM-Eval for transcriptome comparisons)**
http://deweylab.biostat.wisc.edu/detonate/detonate-1.11.tar.gz

**DBG2OLC (last update: Jun 11, 2015)**
Original: https://github.com/yechengxi/DBG2OLC.git
3e577397c0c2ad2a4b61da29be59572cef6e3086
JMU Version: I forked DBG2OLC and made/used a branch -- obtain by:
git clone -b split_reads_by_backbone_minimize_open_files
https://github.com/JohnUrban/DBG2OLC.git

**Dfam_Consensus-20181026**
- used with RepeatMasker/RepeatModeler
http://www.dfam.org/web_download/Current_Release/Dfam.hmm.gz

**EdgeR 3.28.0**
https://bioconductor.org/packages/release/bioc/html/edgeR.html

**Exonerate v2.2.0 (used with Maker)**
http://ftp.ebi.ac.uk/pub/software/vertebrategenomics/exonerate/exonerate-2.2.0-x86_64.tar.gz

**Falcon v 0.7.3**
git://github.com/PacificBiosciences/FALCON-integrate.git
c275aaac952ee1fd830dbe32ddc3bfb4e929fcb0

**Fast5Tools**
https://github.com/JohnUrban/fast5tools
c6e6026769457cbde57636f7044e288302d22946

**FRC$^{bam}$ v1.3.0**
https://github.com/vezzi/FRC_align.git
5b3f53e01cb539c857fd4230ec9410d76220fe22

**GeneMark-ES Suite v4.38 (used with Maker and for training)**
http://topaz.gatech.edu/GeneMark/tmp/GMtool_72DIF/gm_et_linux_64.tar.gz

**gffcompare v0.10.6**
https://github.com/gpertea/gffcompare

**HINGE**
https://github.com/fxia22/HINGE.git
86db7bc630d2956031c4116739a7998f824738b9

**HISAT2 v2.1.0 (for genome evaluations and StringTie assemblies)**
https://github.com/infphilo/hisat2.git

**HMMer v3.2.1 (for use with BUSCO and Maker)**
http://eddylab.org/software/hmmer/hmmer.tar.gz

**InterProScan v5.32-71.0**
ftp://ftp.ebi.ac.uk/pub/software/unix/iprscan/5/5.32-71.0/interproscan-5.32-71.0-64-bit.tar.gz

**KineticsTools v0.6.0**

git commit da2286bbd5a6fb2f524c70c6c5616b9715317b83
https://github.com/PacificBiosciences/kineticsTools.git

**LAP v1.1**
http://www.cbcb.umd.edu/~cmhill/files/lap_release_1.1.zip

**Maker v2.31.10**
http://yandell.topaz.genetics.utah.edu/maker_downloads/AA48/FC36/CF23/C760F5B508D12D14AFA82FC83AE5/maker-2.31.10.tgz

**Maligner**
Original: https://github.com/LeeMendelowitz/maligner.git
Fork used: https://github.com/JohnUrban/maligner.git
Commit: 21b70bd7f8615942a65f746dc5934ddd430b32e4
I only changed a hard coded default to allow much longer molecules to align:
static int max_query_frags = 50000 ##Originally 50

**MarginAlign**
git commit cc28457352d3b9e65ecb46a04a45903ba3f55c61
https://github.com/benedictpaten/marginAlign.git

**MaSuRCA v3.1.3**
http://www.genome.umd.edu/masurca.html

**MEME 5.0.2**
http://meme-suite.org/doc/download.html

**Megahit v1.0.5**
https://github.com/voutcn/megahit.git
a851582f8a6e4ca9cbe41b946dc522336033adeb

**Miniasm 0.2-r137-dirty**
https://github.com/lh3/miniasm
17d5bd12290e0e8a48a5df5afaeaef4d171aa133

**Minimap 0.2-r124-dirty**
https://github.com/lh3/minimap
1cd6ae3bc7c7a6f9e7c03c0b7a93a12647bba244

**Nanopolish 0.5.0**
https://github.com/jts/nanopolish.git
methyltrain version 0.5.0 for R7 data

**NSEG (used with RepeatModeler)**
ftp://ftp.ncbi.nih.gov/pub/seg/nseg/

**PBalign 0.2.0.141024**
https://github.com/PacificBiosciences/pbalign

**PBDAGCON**
Git commit f19aed1668d6ace0ab3ab4eb0e1e7d81139492a8
https://github.com/PacificBiosciences/pbdagcon.git

**PBJelly from PBSuite v15.8.24**

https://sourceforge.net/p/pb-jelly/wiki/Home/

**Platanus v1.2.4**
http://platanus.bio.titech.ac.jp

**Pbh5tools v 0.8.0**
https://github.com/PacificBiosciences/pbh5tools
2679c17687a868690595ae0d9e856c2fead28ff9

**Picard Tools 2.1.1**
https://github.com/broadinstitute/picard/releases/tag/2.1.1

**Pilon v1.18**
https://github.com/broadinstitute/pilon/releases/download/v1.18/pilon-1.18.jar

**Poreminion v 0.4.4**
https://github.com/JohnUrban/poreminion
c6a1bc8d7cfc7da675fdd0979622c4e200a16657

**Quiver - GenomicConsensus_v1.1.0 (pbcore-1.2.4, ConcensusCore 1.0.1)**
https://github.com/PacificBiosciences/GenomicConsensus

**RaCon**
https://github.com/isovic/racon
c342fc65f6fd686e7975238c207fb6f3aea671d1

**REAPR v1.0.18**
ftp://ftp.sanger.ac.uk/pub/resources/software/reapr/Reapr_1.0.18.test_data.tar.gz

**RECON v1.08 (used with RepeatModeler)**
http://www.repeatmasker.org/RepeatModeler/RECON-1.08.tar.gz

**RepBase Repeat Masker Edition 20181026**
- used with RepeatMasker/RepeatModeler
https://www.girinst.org/repbase/

**RepeatMasker** v1.332 (used on its own and within Maker)
- version development-$Id: RepeatMasker,v 1.332 2017/04/17 19:01:11 rhubley Ex

**RepeatModeler v1.0.11**
http://www.repeatmasker.org/RepeatModeler/RepeatModeler-open-1.0.11.tar.gz

**RM BLAST 2.2.28**
**-** used with RepeatMasker/RepeatModeler
ftp://ftp.ncbi.nlm.nih.gov/blast/executables/rmblast/2.2.28/ncbi-rmblastn-2.2.28-x64-linux.tar.gz
ftp://ftp.ncbi.nlm.nih.gov/blast/executables/blast+/2.2.28/ncbi-blast-2.2.28+-x64-linux.tar.gz

**Salmon 1.1.0**
https://combine-lab.github.io/salmon

**SAMtools v1.3**
https://github.com/samtools/samtools/releases/download/1.3/samtools-1.3.tar.bz2

**SAMtool (HTSlib) v1.3**

https://github.com/samtools/htslib/releases/download/1.3/htslib-1.3.tar.bz2

**SciaraTools**
https://github.com/JohnUrban/sciara-project-tools

**SGA v0.10.14**
https://github.com/jts/sga.git
2ab3eae4d3d58b8d0dea3d60f2684c1181049a8f

**SMARTdenovo (2016-03-07):**
commit 61cf13dcaed6bb561129b60eaa833fa9f976f9b1
https://github.com/ruanjue/smartdenovo

**SNAP (Semi-HMM-based Nucleic Acid Parser) gene prediction tool**
- used with Maker and training
http://korflab.ucdavis.edu/Software/snap-2013-11-29.tar.gz

**Sniffles v1.0.8**
commit 51a44a23b01b006cd1241540105376ec934f1512
https://github.com/fritzsedlazeck/Sniffles

**Soapdenovo2 v2.04**
https://github.com/aquaskyline/SOAPdenovo2.git
dd6a98ba19bb21c3513a46ad5047d08e57583ab0

**SPAdes v3.8.0 (and BayesHammer)**
http://spades.bioinf.spbau.ru/release3.8.0/SPAdes-3.8.0-Linux.tar.gz

**StringTie v1.3.5**
https://github.com/gpertea/stringtie

**TransRate v1.0.3**
https://bintray.com/artifact/download/blahah/generic/transrate-1.0.3-linux-x86_64.tar.gz

**Tandem Repeats Finder (TRF) v 4.09**
**-** used with RepeatMasker/RepeatModeler
http://tandem.bu.edu/trf/trf.download.html

**Trimmomatic v0.32**
http://www.usadellab.org/cms/?page=trimmomatic

**Trinity r20140413p1**
https://sourceforge.net/projects/trinityrnaseq/files/PREV_CONTENTS/previous_releases/trinityrnaseq_r20140413p1.tar.gz

**Velvet 1.2.08**
https://www.ebi.ac.uk/~zerbino/velvet/
